# BBX transcription factor evolution in the green plant lineage

**DOI:** 10.1101/2025.09.03.673946

**Authors:** Huiting Jin, Sitaram Rajaraman, Jarkko Salojärvi, Michael Wrzaczek, Aleksia Vaattovaara

**Author notes:** **Corresponding authors:** Aleksia Vaattovaara, University of Helsinki, Organismal and Evolutionary Biology Research Programme, Faculty of Biological Sciences, Viikinkaari 1, POB 65, FI-00014 Helsinki, Michael Wrzaczek, Institute for Plant Molecular Biology, Biology Centre CAS, Branišovská 1160/31, 370 05 České Budějovice, Czech Republic. These authors contributed equally (Huiting Jin, Aleksia Vaattovaara).

## Abstract

BBX transcription factors play versatile roles in integrating environment cues and regulating plant development. The model plant Arabidopsis continues to be a primary tool for the molecular investigation of protein function, but comprehensive evolutionary studies across different plant species can support translational research. Here, we constructed robust maximum likelihood phylogenies of full-length BBX proteins, the B-box domain, and the CCT domains by using diverse plant species ranging from algae to eudicots and monocots. Our analysis resolved five major structural groups (I–V) with BBXs from algae at the root of major clades. Our results suggest that Group II emerged after the split between Chlorophytes and Streptophytes. Our results also highlight subgroupings in the different BBXs clades. The evolution of the B-box domains showed a dynamic domain losses and acquisitions overall correlating with the BBX phylogeny. While the CCT domain was overall highly conserved across all groups, we identified secondary loss of the CCT domain in select lineages. We also detected significant expansions and contractions among angiosperm BBX orthogroups. Overall, the BBX family demonstrates ancient origins, structural plasticity via domain gain, loss and duplication, and lineage-specific innovations that underpin their diverse roles in light signaling, development and stress responses.

## Introduction

Transcriptional regulation controls the abundance of mRNA production by regulating its production rate from the DNA template, directly influencing protein abundance (Vogel and Marcotte 2012). Transcription factors (TFs) positively and negatively regulate or modulate the expression or repression of genes. General TFs are involved in fundamental cellular processes and are necessary for the basal transcription machinery to function properly (Carlberg and Molnár 2020). Specialized TFs target and regulate the expression or repression of particular genes or gene sets, often in response to specific cellular signals or environmental cues (Lee and Young 2000). The genome of the model plant *Arabidopsis thaliana* encodes approximately 1500 TFs, of which ∼ 45% belong to plant specific families (Riechmann et al. 2000). Generally, TFs include a DNA binding domain for the recognition of binding sites but also additional motifs or domains which can facilitate multimerization of recruitment of other proteins for co-regulation of gene expression (Luscombe et al. 2000). TFs are typically classified into different families according to their DNA binding domain type.

One group of TFs are the zinc-finger-containing B-box (BBX) proteins. BBXs are characterized by one or more B-box domains that are frequently followed by a CCT domain and a valine-proline (VP) motif in the C-terminus (Khanna et al. 2009). BBX TFs are not plant-specific but are conserved in multicellular organisms (Sardiello et al. 2008). The B-box is considered to be critical for protein-protein interactions and has been subdivided into two types, B-box1 and B-box2 (B1, B2), based on the consensus sequence and the spacing of zinc-binding residues (Crocco and Botto 2013). Both types contain five cysteine (C) and two histidine (H) residues involved in binding two Zn^2+^ ions in a RING-like domain. The CCT domain is typically located at the C-terminal region of B-box proteins belonging to structural groups I-III (Gendron et al. 2012). The CCT domain has been shown to be involved in protein-protein interactions, nuclear localization, and transcriptional regulation, facilitating BBX proteins to interact with other transcription factors, co-regulators, and DNA binding (Wenkel et al. 2006; Yan et al. 2011). These interactions are crucial for the regulatory functions of BBX proteins in various biological processes. BBX proteins containing a CCT domain are known to act in various physiological processes such as flowering time control, photomorphogenesis, circadian clock regulation, and stress responses (Gendron et al. 2012).

The Arabidopsis genome contains 32 BBX TF family members (AtBBX1 to AtBBX32). They have been previously (Khanna et al. 2009) subdivided into five distinct groups based on their domain composition (Figure S1). Members of group I contain two B-boxes, B1 and B2, and a CCT domain followed by a conserved valine proline (VP) pair. Group II contains B1, B2, and CCT domains, similar to group I, but the B2 domain displays different features. Group III contains a single B-box, with a CCT domain. Members of group IV have B1 and B2 but lack the CCT domain. Members of group V have one B-box but no other detectable domains. This domain-based grouping is also reflected in the phylogenetic clustering of Arabidopsis BBX proteins (Huang et al. 2012). Similar to TF families in general, whole-genome duplications (WGDs) have been found to be the main driver of the BBX gene family expansion, followed by segmental duplication. These duplications tend to have more introns and are subject to purifying selection (Cao et al. 2017).

BBX proteins play pivotal roles in controlling flowering time, particularly in response to photoperiodic cues. CONSTANS (CO/AtBBX1), the first identified BBX protein, is a central regulator of flowering by directly stimulating *FLOWERING LOCUS T* (*FT*) expression via the CCT domain (Putterill et al. 1995; Samach et al. 2000; Suárez-López et al. 2001; Valverde et al. 2004). Overall, BBXs exhibit both positive and negative control over flowering by controlling the expression of BBX1 (Liu et al. 2020). BBXs also participate in light signaling pathways and photomorphogenesis. They interact with key components of light perception and signaling, including phytochromes, CONSTITUTIVE PHOTOMORPHOGENESIS 1(COP1)/ SUPRESSOR OF PHYA1 (SPA1) complexes, and ELONGATED HYPOCOTYL5 (HY5) transcription factor (Laubinger et al. 2006). The COP1-SPA1 complex targets the transcription factor HY5for its degradation and various BBX proteins modulate this process. In the HY5 signaling pathway, BBX proteins act as positive or negative regulators of photomorphogenesis (Indorf et al. 2007; Datta et al. 2007; Chang et al. 2008; Holtan et al. 2011; Fan et al. 2012; Xu et al. 2016; Job et al. 2018; Luo et al. 2018; Lin et al. 2018; Heng et al. 2019; Zhao et al. 2020). BBX4 and BBX11 also promote photomorphogenesis by repressing negative regulators like PHYTOCHROME-INTERACTING FACTORS (PIFs) PIF3 and PIF4 (Heng et al. 2019; Song et al. 2021).

In addition to their roles in the developmental regulation, BBX proteins play critical roles in abiotic stress responses, including responses to drought, cold, heat, and salt stress. *Chrysanthemum morifolium* CmBBX19, CmBBX22, CmBBX24 and *Malus domestica* MdBBX10 influence plant drought tolerance by regulating the expression of ABA-responsive genes and interacting with key transcription factors, e.g., HY5 (Yang et al. 2014; Riboni et al. 2016; Liu et al. 2019; Xu et al. 2020). AtBBX7 and AtBBX8 act downstream of the CRY2-COP1-HY5 module to positively regulate cold acclimation by modulating the expression of cold-responsive genes (Li et al. 2021b). Thermomorphogenesis involves AtBBX18 and AtBBX23, which promote hypocotyl growth at elevated temperatures by reducing ELF3 levels and releasing PIF4 activity (Ding et al. 2018). BBX18 also reduces thermotolerance by downregulating heat-shock proteins including HSP70 and 101 (Wang et al. 2013). STO/AtBBX24 and *Ginkgo biloba L.* GbBBX25 are involved in the response to salt stress by regulating genes related to osmotic stress and ion homeostasis promoting root growth under salt stress (Nagaoka 2003; Yang et al. 2014; Huang et al. 2021). In addition, BBX proteins can act in conjunction with other BBX family members and different plant B-box proteins. For example, AtBBX32 interacts with STH2/AtBBX21 to negatively regulate light signaling (Holtan et al. 2011), and AtBBX32 physically interacts with soybean BBX62 (GmBBX62) through its B-box domain, and transgenic soybean lines expressing AtBBX32 show increased grain yield in multiple environments tested (Qi et al. 2012). The B-box domain has a crucial role in regulating transcription, but the specific function remains unclear.

Taking into an account the important regulatory functions of BBXs, identified mostly in model species such as Arabidopsis, there is an interest to gain deeper insights into the evolution of BBXs throughout the plant kingdom. Phylogenetic analyses will allow identification of homologous proteins between model species as well as plant species having agricultural importance. Evolutionary reconstruction can be challenging, especially in large gene families, which frequently display lineage-specific expansions or contractions of subgroups. The BBX gene family is highly conserved and widely distributed in plants, with the highest number of homologs found in angiosperms (e.g., Arabidopsis: 32, rice: 30). The frequency of BBX homolog identification varies across different organism groups, reflecting the evolutionary and functional diversity of these genes. Currently most information on the function of BBX transcription factors is derived from studying the model plant species *Arabidopsis thaliana*. However, for translational research extrapolating information from Arabidopsis to other plants with agricultural and economic importance it is imperative to obtain a detailed understanding of the evolution of the BBX family in the plant lineage.

## Methods and materials

### Sequence identification and annotation

17 plant species with 3 red algae species and 7 green algae species (Supplementary Table 1) were selected to cover the major plant lineages for BBX phylogenetic analyses. Primary nucleotide and protein sequences were retrieved from Phytozome 13 (Goodstein et al. 2012), Plaza 4.5 (Proost et al. 2010), NCBI (O’Leary et al. 2024). Norway spruce (*Picea abies*) was obtained from Plantgenie (Sundell et al. 2015) and *Klebsormidium nitens* NIES-2285 V1.0 from its genome project BioProject ID: PRJDB718 (Hori et al. 2014). To ensure comprehensive identification of BBX homologs, genomic sequences were re-scanned with TBLASTN (Camacho et al. 2009) to ensure no BBX homologs were missed as previously described (Vaattovaara et al. 2017, 2019).

Each predicted BBX protein sequence was confirmed by SMART (Simple Modular Architecture Researcher Tool) (Letunic et al. 2021) to contain B-box domain. In addition, HMMER tool search for PFAM domain (Finn et al. 2010, 2016) for B-box domain ID PF00643 (B-box zinc finger) was carried out among BBX protein sequences from all study species. Same method was used to identify CCT domain (PFAM ID PF06203). The sequences for the domain-specific analyses were extracted based on this information. If the domain annotation was uncertain based on SMART and HMMER, B-box motif identifications were manually extracted based on multiple sequence alignment done with MAFFT (version 7.505) (Katoh and Standley 2013) in Wasabi (http://wasabiapp.org/) (Veidenberg et al. 2015). If the gene model was found to be erroneous, it was corrected using Fgenesh+ (Salamov and Solovyev 2000). Detailed annotation information can be found from in the Supplementary Table 2 and complete set of amino acid sequences used in this study are provided in Supplemental Data 1 and Supplemental Data 2.

### The sequence logos of conserved domains

The sequence logos of B-box (for all BBXs and each group separately) and CCT (for group I, II, III and all of them together) domains were generated with MEME suite software (Bailey et al. 2015) (https://meme-suite.org/meme/tools/meme) using the extracted sequences of B-box and CCT domains predicted manually or by SMART.

### Phylogenetic analyses

Multiple sequence alignments of BBX proteins and domain sequences were generated using the MAFFT. Maximum likelihood (ML) phylogenetic trees (for the full-length BBX protein, B-box domain and CCT domain sequences) were constructed by using RAxML (version 8.2.12) (Stamatakis 2014) with 1000 bootstrap replicates. Bootstrapped trees are presented in Supplementary Data 3. The neighbor-joining (NJ) of the BBX protein sequences from Arabidopsis was built by MEGA 11 (Molecular Evolutionary Genetics Analysis) (Tamura et al. 2021) with 1000 bootstrap replicates. Phylogenetic trees were visualized by program FigTree (version 1.4.4).

### BBX protein mean distance calculation

Using MEGA 11 software (Tamura et al. 2021), we calculated evolutionary distances between sequences. Aligned BBX sequences in FASTA format were loaded into MEGA. Guided by the protein sequence tree, sequences were grouped, and mean distances were computed within and between these groups using default settings.

### CAFE5 analyses

Gene family evolutionary analysis was carried out using CAFE5 software (Mendes et al. 2021) for eight species *Amborella trichopoda*, *Aquilegia coerulea*, *Arabidopsis thaliana*, *Populus trichocarpa*, *Vitis vinifera*, *Brachypodium distachyon*, *Oryza sativa* and *Spirodela polyrhiza*. The orthogroup estimation was done with OrthoMCL (version 2.0.9) (Li et al. 2003) using an inflation parameter of 1.5 for the clustering phase. A rooted ultrametric tree and a table containing orthogroup gene counts per species were passed as input and CAFE5 was run using default parameters. The BBXs were analyzed in two separate runs: as five groups based on the phylogenetic results and separately as one group were all BBX subgroups were combined as one group. The results were visualized with CafePlotter (https://github.com/moshi4/CafePlotter) with option “--ignore_branch_length”.

## Results

### Identification and evolution of BBX protein models

In order to elucidate the evolution of BBX transcription factors we analyzed the genomes of representative species from major plant clades: the charophyte *Klebsormidium nitens* (Hori et al. 2014), the liverwort *Marchantia polymorpha* (Bowman et al. 2017), the gymnosperm *Picea abies* (Nystedt et al. 2013), the basal angiosperm *Amborella trichopoda* (Albert et al. 2013; Chamala et al. 2013), the monocots *Oryza sativa* (Sakai et al. 2013), *Sorghum bicolor* (McCormick et al. 2018), *Spirodela polyrhiza* (Wang et al. 2014), *Brachypodium distachyon* (The International Brachypodium Initiative 2010), and the eudicots *Arabidopsis thaliana* (Cheng et al. 2017), *Aquilegia coerulea* (Filiault et al. 2018), *Populus trichocarpa* (Tuskan et al. 2006), *Medicago truncatula* (Tang et al. 2014), *Prunus persica* (The International Peach Genome Initiative et al. 2013), *Vitis vinifera* (The French–Italian Public Consortium for Grapevine Genome Characterization 2007), *Theobroma cacao* (Motamayor et al. 2013), *Solanum tuberosum* (The Potato Genome Sequencing Consortium 2011), *Nelumbo nucifera* (Li et al. 2021a) and *Solanum lycopersicum* (Hosmani et al. 2019). The majority of protein sequences in databases are annotated using computational methods but have not been experimentally validated. The level of erroneous annotations in large databases is currently unknown and has not been analyzed in depth. We manually reviewed the predicted BBX protein sequences and identified several annotation errors, including low quality annotations and misannotated gene models. Inaccurate gene models might cause problems in downstream analyses, and therefore low quality gene models were manually reannotated using Fgenesh+ (Salamov and Solovyev 2000). Through manual curation and exclusion of partial gene models, we obtained high quality amino acid models for subsequent analyses. In total, we obtained 425 high-quality BBX gene models and protein coding sequences (Supplementary Data 1) from the genomes of the selected plant species (Table 1). Additionally, to trace the early evolution of BBX proteins, we also included 17 BBX sequences from red algae *Chondrus crispus* (Collén et al. 2013), *Galdieria sulphuraria* (Schönknecht et al. 2013)*, Porphyridium purpureum* (Nelson et al. 2021), and green algae *Volvox carteri* (Prochnik et al. 2010)*, Chlamydomonas reinhardtii* (Merchant et al. 2007)*, Ostreococcus lucimarinus* (Palenik et al. 2007) and *Coccomyxa subellipsoidea* (Blanc et al. 2012). These lineages represent key transitions in photosynthetic eukaryotes, and their genomes can provide insights into the origin and diversification of BBX transcription factors. The amino acid sequences of the algal BBXs are provided as Supplementary Data 2 and the genome information in Table S1.

**Table 1:**
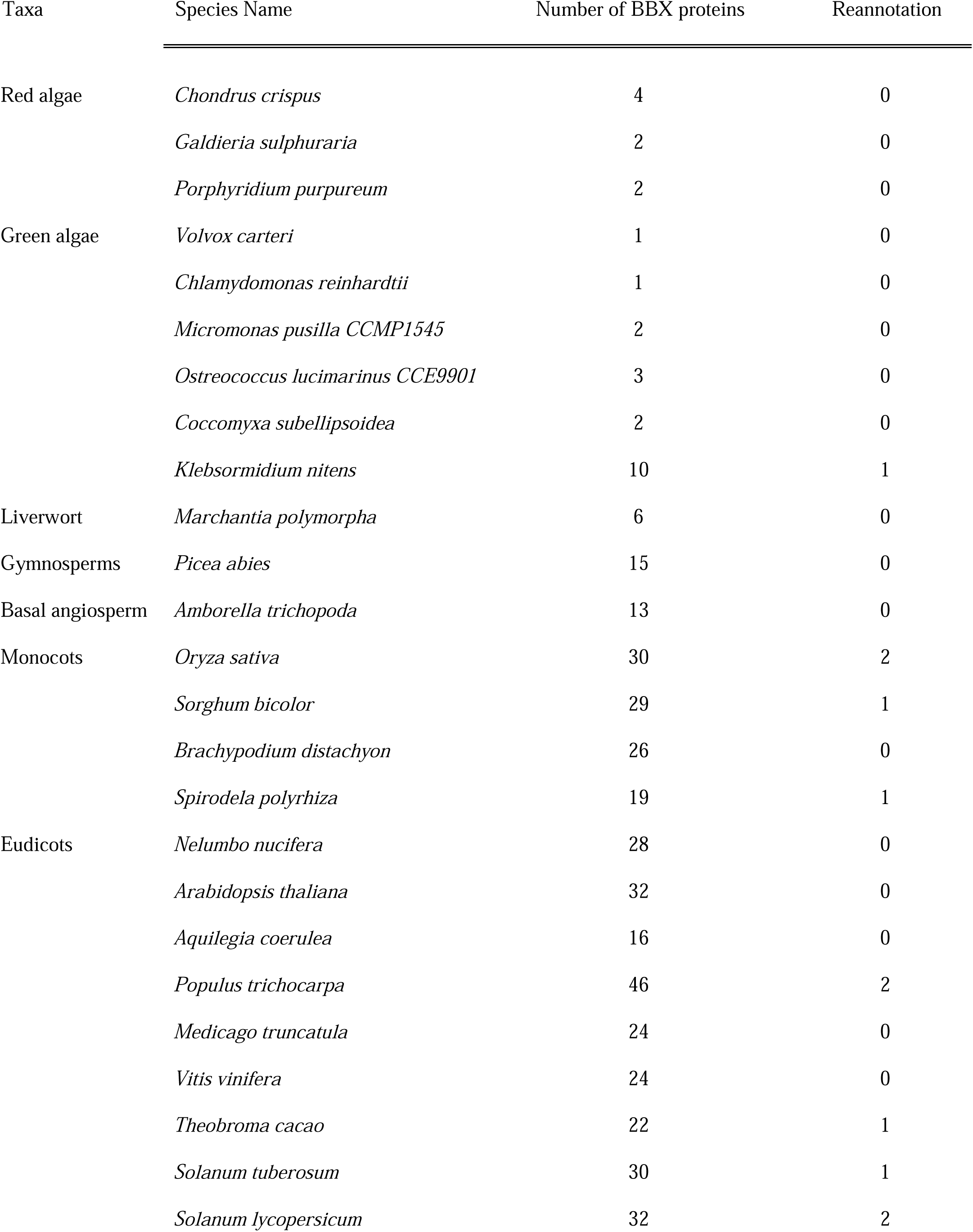
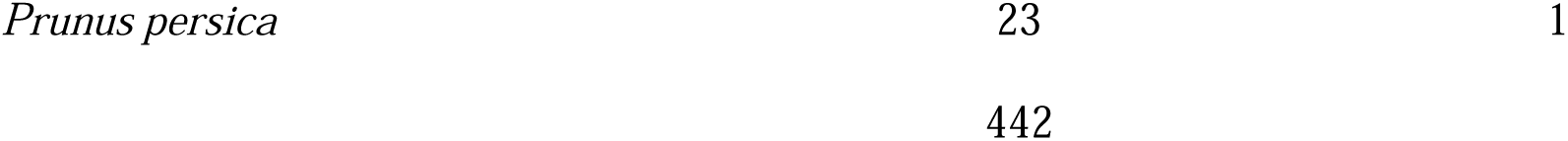
The number of the reannotated gene models contain manually curated gene annotations.

### Phylogenetic analysis of BBX proteins in the plant kingdom

Previous works explored the phylogenetic relationships of BBX proteins in Arabidopsis and other plants using Neighbor-Joining (NJ) phylogenetic inference. Previous analyses have identified 32 BBX TFs in Arabidopsis referred to as AtBBX1 to AtBBX32 (Khanna et al. 2009). We reanalyzed the evolutionary relationships and domain structures of the BBX gene family in the model plant Arabidopsis and constructed both a NJ and a maximum likelihood (ML) protein sequence tree for AtBBX proteins (Fig. S2). Our NJ phylogeny showed five distinct clades consistent with previous studies using Neighbor-joining analysis and clustering based on domain architecture. The ML tree demonstrated an overall similar grouping as the NJ tree, with the exception of AtBBX26 and AtBBX27. Previously described as members of group V based on domain composition and NJ phylogeny (Khanna et al. 2009), the ML tree placed AtBBX26 in a long branch close to root of group III while AtBBX27 was placed in group II. Manual validation of the protein domains in AtBBX26 identified a B2 based on sequence alignment, which was not identified earlier or by SMART, while AtBBX27 was found to contain both B1 and B2 domains with a partial CCT domain that is approximately half of the full-length CCT domain (Fig. S1).

To investigate the structural diversification and evolution of BBX proteins, which primarily arises from variations in their domain composition and arrangement, such as B-box domains, CCT domain, and other regulatory domains, we conducted a global phylogenetic analysis with the full-length amino acid sequence of 425 BBXs from land plant species and the charophyte algae *K. nitens* (Table 1). In the multispecies ML phylogenetic tree (Fig 1, S3 and S4), *K. nitens* BBXs placed near the root of each respective clade. Overall, the identified clades were consistent with observed subfamily divisions based only on Arabidopsis BBX sequences (Fig. S2). However, AtBBX26 and AtBBX27 were still positioned differently from previous classifications. AtBBX26 formed a long branch close to the root of group IV, while AtBBX27 was located in group II similarly to its placement in the ML phylogeny inferred for Arabidopsis BBXs (Fig. S2). This contrasts their placement in group V in previous studies (Gangappa and Botto 2014). In the multispecies ML tree, some groups formed clear subclades in contrast to the groupings obtained using only Arabidopsis BBX sequences. Specifically, group I was separated into two distinct subclades and group IV separated into four distinct subclades. To elucidate the ancestral origins of BBX proteins, we expanded our phylogenetic analyses to include red algae and green algae belonging to Chlorophyta (Table 1, Fig. S5). The chlorophyte green and red algae BBXs were clustered within the groups present in previous analyses with land plants and Klebsormidium (Fig. 1), but no algal sequences were clustered with group II. Group II contained only BBX sequences from streptophytes, suggesting that this clade emerged after the divergence of Chlorophyta and Streptophyta (about 725–1200 MY ago) (Leliaert et al. 2012).

**Figure 1.**
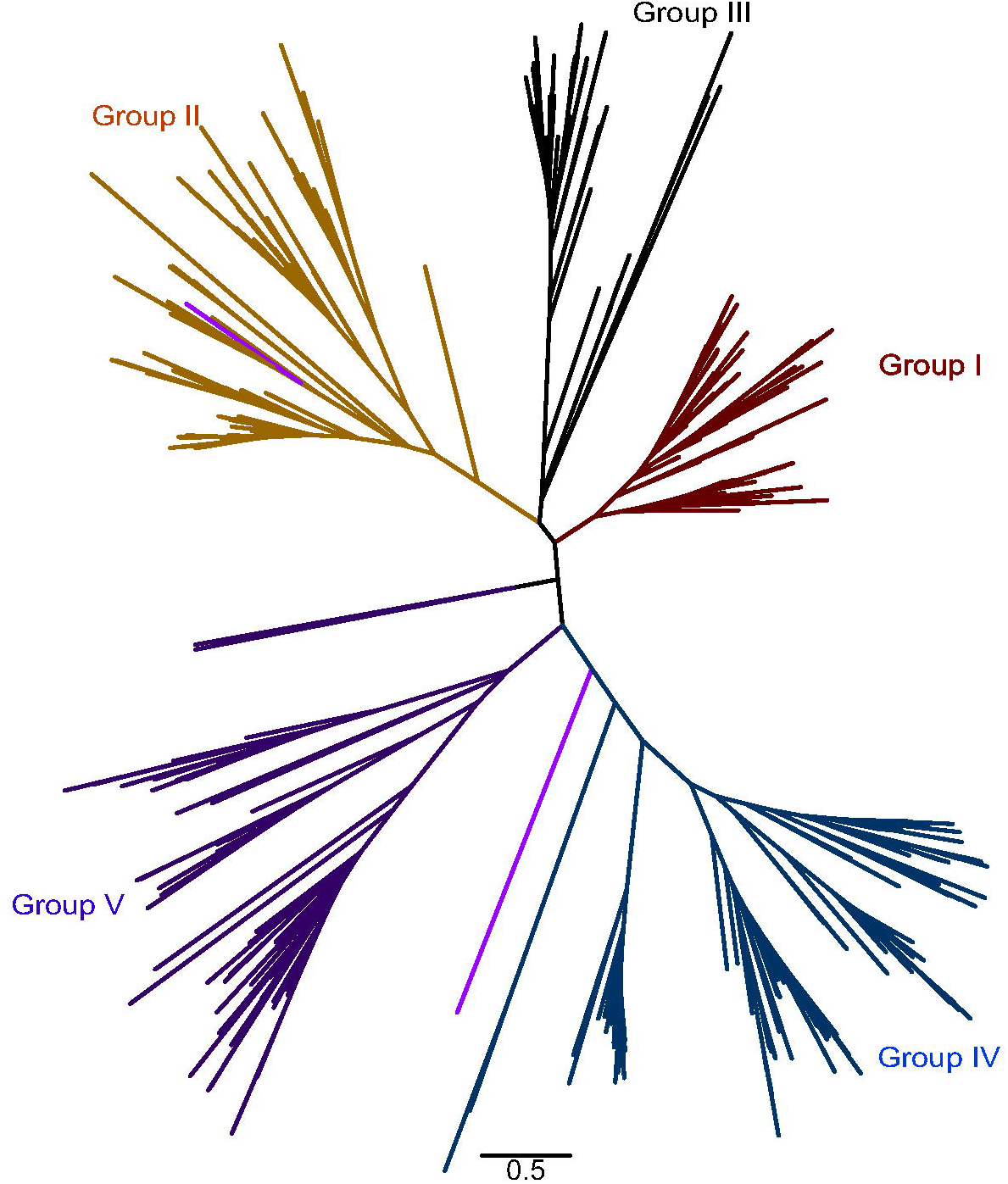
Unrooted maximum likelihood phylogenetic tree for BBX proteins. The phylogenetic tree was inferred with the maximum-likelihood method using high-quality full-length B-box containing protein sequences. Topological consistency was confirmed across multiple reconstructions with varied seed numbers, and branch robustness was evaluated using 1,000 bootstrap replicates. A radial layout illustrating the full topology and groups are color-coded as follows: group I (red), group II (yellow), group III (black), group IV (blue), and group V (dark purple). Notably, AtBBX26 and AtBBX27 are highlighted in light purple to distinguish their unique placement within the phylogeny.

### Phylogenetic analysis of B-box domain in the plant kingdom

The B-box motif plays an important role in protein-protein interactions and in transcriptional regulation (Gangappa and Botto 2014). In order to trace the evolutionary origins and diversification of B-box domains in land plants, we extracted the B-box sequences from the 425 BBXs using SMART or manual extraction based on MAFFT alignment (Table S2) and conducted a targeted phylogenetic analysis of the identified B-box 1 and B-box 2 domains (Fig. 2 and S6). The B-box consensus sequence for this data set is approximately 38 residues, based on a MEME logo alignment (Fig. 3); The length is similar to the profile HMM transcription factor binding model (TFBM) for PFAM ID PF00643 (42 residues).

**Figure 2.**
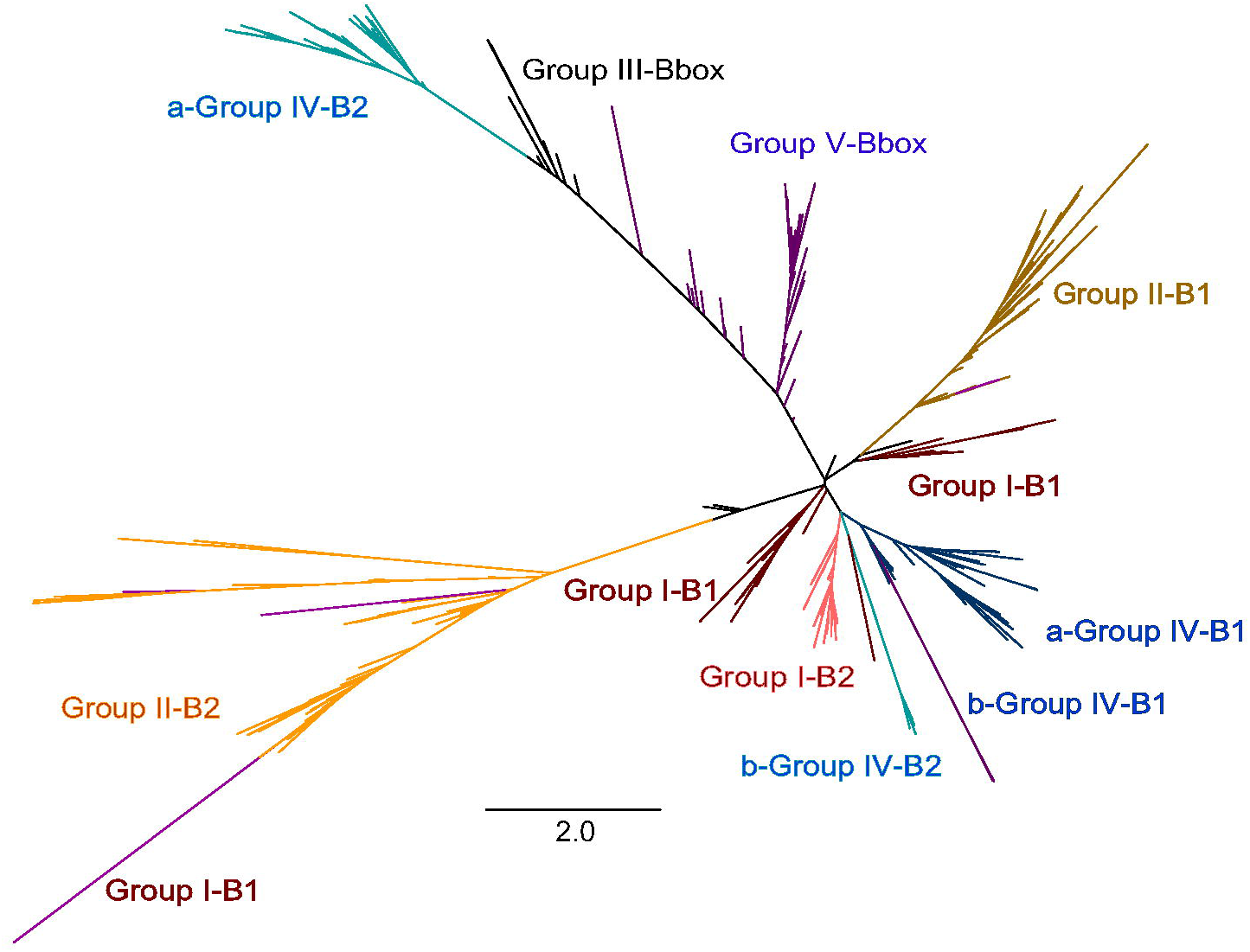
Unrooted maximum likelihood phylogenetic tree for B-box domain. Phylogenetic relationships of B-box domains were reconstructed using the maximum-likelihood method based on B-box domain sequences. Topological consistency was validated through multiple tree reconstructions with varied seed numbers, and branch support was assessed using 1,000 bootstrap replicates. A radial layout showing the full topology. groups are color-coded to reflect domain classification: group I-B1 (dark red), group I-B2 (light red), group II-B1 (dark yellow), group II-B2 (light yellow), group III (black), group IV-B1 (dark blue), group IV-B2 (light blue), and group V (dark purple). AtBBX26 and AtBBX27 are highlighted in light purple to denote their distinct phylogenetic placement.

**Figure 3.**
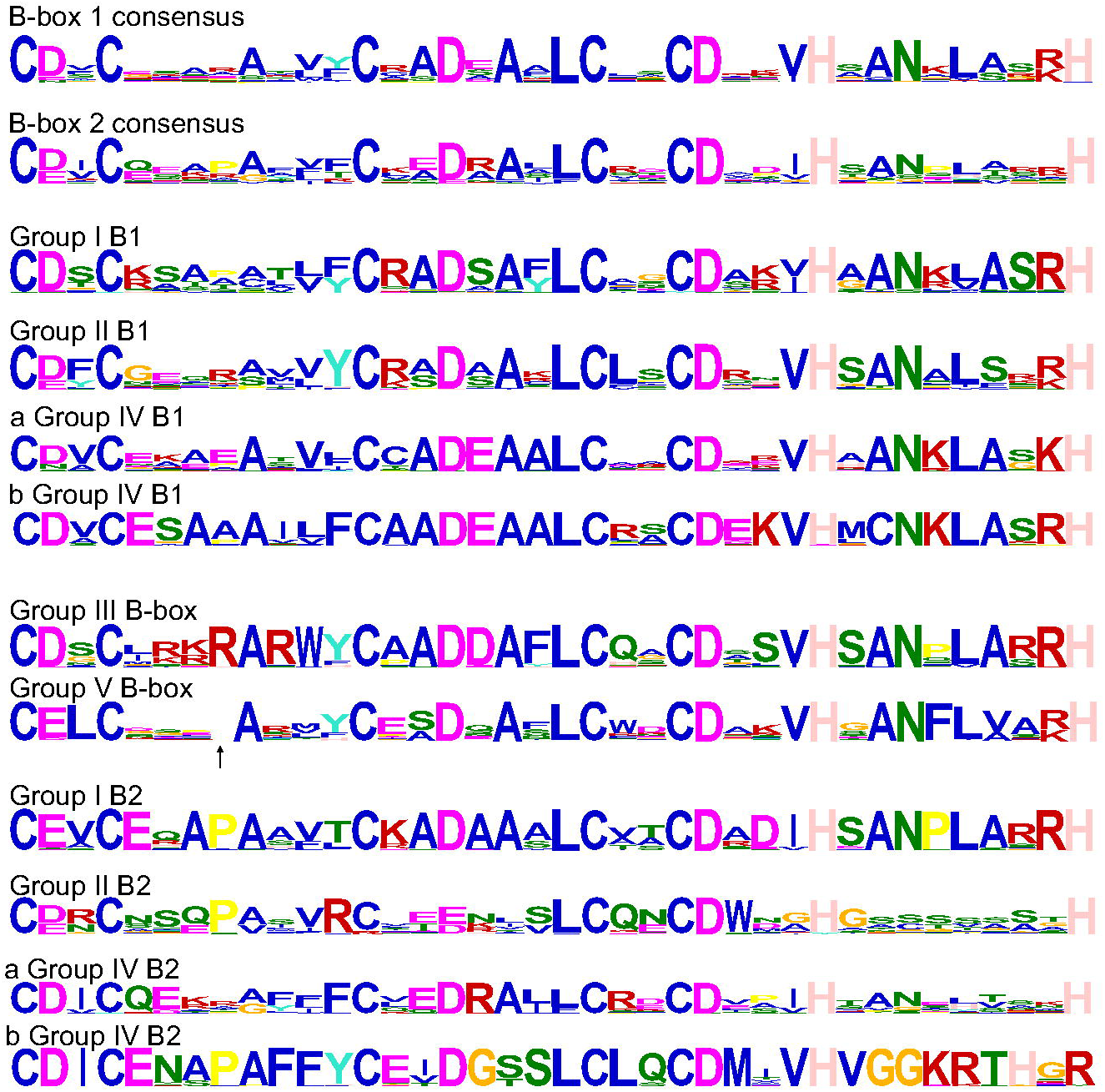
The MEME logos of B-box domain. Sequence logos generated by MEME analysis illustrate the conservation patterns of B-box domains across BBX proteins. The figure includes overall sequence logos for B-box1 (B1) and B-box2 (B2) domains B-box1: CD/EXCX3-4AX3CXA/SDXAXLCX2CDX2VHXAX2LX2R/KH; B-box2: CD/EXCX4AX3CXA/ED/EX3LCX2CDX3HX8H. Group-specific logos for group I, II, and IV B1 domains. Logos for group III and group V B-box domains, and group-specific logos for group I, II, and IV B2 domains. Notably, in the group V B-box domain logo, an arrow marks the position of a missing amino acid, highlighting a unique structural divergence within this group.

In the B-box domain tree (Fig 2. and S6), the B1 domains for group II are found in one clade, while the B1 domains for group I are divided into two separate clades, and for group IV the form two separated clusters. Unlike the B1 domains, the B2 domains for group I are all found in a single clade. Group II B2s form one cluster, and B2 of AtBBX27, as well as B1 and B2 of AtBBX26, are included in this clade. Interestingly, for group IV, while the B1 subgroups (named a and b) were only slightly diverged, the subgroups differ dramatically for the B2 domain. The a-subgroup B1 and B2 clusters are highly diverged, while b-subgroup B1 and B2 clusters are highly conserved. This pattern could implicate that in group IV the B2 for a and b subgroups have different evolutionary trajectories. In the b-subgroup the B2 domain could originate from a duplication of the ancestor of group IV B1, and in the a-subgroup share ancestry with group III and V as these groups with single BBX domain have their BBX domain sequences grouping in the same major branch as a-subgroup B2s for group IV.

The conserved residues of the B-box domains in the N-terminal region (Fig. 3) are critical for structure and function, including canonical zinc finger C3H2 conserved motif, characterized by three cysteine and two histidine residues coordinating zinc ions (Massiah et al. 2007), that likely stabilize the metal-binding core, alongside structurally conserved residues such as valine, alanine, arginine, and leucine, which contribute to hydrophobic packing, electrostatic interactions, or domain stability (Al Mughram et al. 2023). The sequence conservation of B1 and B2 for all BBXs and separate conservation of B1 and B2 of the five structural groups of BBXs are illustrated in Fig. 3. The position weight matrices (PWMs) based on B1 and B2 from all groups have similar patterns for conserved amino acids and are more diverged for other positions compared to group-specific domain illustrations. Group V BBX proteins display a slightly shorter B-box domain (∼37 residues vs. ∼38 in other groups), with an absence of one amino acid between position 4 to 9 (typically a conserved R or P in other groups). This truncation possibly disrupts the tertiary structure of the domain, potentially altering coordination of the Zn^2+^ positioning or the interaction surfaces. The B2 domain of group II shows significant sequence divergence from other group-specific B-box domain motifs which is in line with phylogenetic tree where group II B2 cluster also shows long and separated branches indicating divergence in sequence level. For group IV B2, the PWM obtained using both subgroups is also more diverged compared to most group-specific B-box domain motifs. In the b-group, hydrophobic and charged residues in positions 26–38 and the H38 substituted by R may disrupt zinc-binding. (Thomas and Emerson 2009).

Subsequently we calculated mean distances based on the differences in B-box domain sequences to estimate evolutionary distances between these sequences to provide insights into the genetic diversity within five groups and the evolutionary relationships between the groups or subgroups. We calculated the mean distance within (Fig. 4a) and between the five groups B-box (including B1 and B2) using Mega11. Group III showed the smallest within group mean distance value compared to other BBX groups, indicating relatively high genetic similarity within this group. For group V, the mean distance also suggests the relative similarity within the group. Group I and group IV show moderate to high diversity, suggesting more divergence within these groups. By contrast, group II displayed the largest within group mean distance of all groups, suggesting considerable divergence in this group (detailed within group mean distance data see Table S3). Based on the mean distance value between groups (detailed between mean distance data see Table S4), group I and group III exhibited the smallest value compared to other groups, suggesting that group I and III likely share a common ancestor. However, it cannot be ruled out that purifying selection is acting on BBXs subclades. Groups III and V, group I and V followed, showing moderately larger but still relatively close distances indicating potential shared ancestry or functional convergence. By contrast group II and group III displayed a much larger distance, indicating early divergence. Finally, group II and group IV showed the largest distance of all, these groups are the most diverged from each other, suggesting an ancient divergence, more relaxed selection, or independent evolutionary paths (detailed between mean distance data see Table S4).

**Figure 4.**
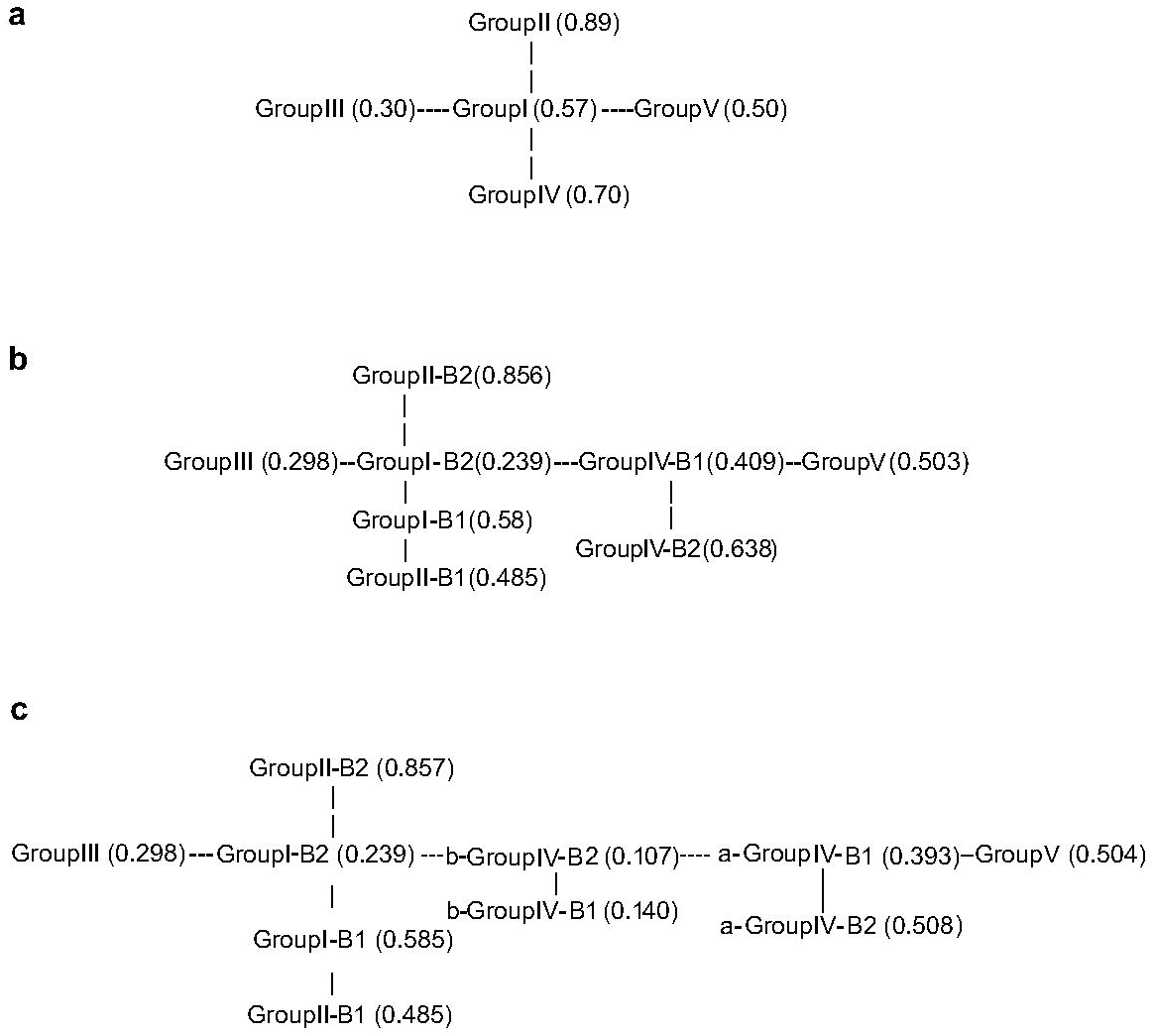
B-box domain evolutionary pathway schematic modes. **a**. Mean Genetic Distances of B-box domains in five groups. The results illustrated the relationships and diversity among five BBX protein groups based on their B-box domain sequences. Within-group mean distances, group III exhibits the smallest mean distance (0.3), indicating high conservation and strong purifying selection, likely linked to conserved roles in light signaling. Group V shows moderate diversity (0.5), reflecting partial functional specialization. Group IV (0.70) and group I (0.57) display moderate to high divergence, suggesting lineage-specific adaptations. Group II (0.89) displays the highest divergence, indicative of adaptive evolution or relaxed selection. Between-group mean distances, illustrating close relationships, group I and group III (0.673) shared ancestry and functional overlap in conserved pathways. Group III and group V (0.694), group I and group V (0.779), group II and group III (0.878) suggested functional convergence or partial shared ancestry. Most divergent pairs: group II and group IV (0.968): Ancient divergence, aligning with distinct roles. Group III and group I form the conserved regulatory hub, group IV and V evolved as later divergent groups. Group II was as an early divergent group. **b** Evolutionary dynamics of B1 and B2 domains. Within-group mean distances illustrated diversity within B-box domain. Group I B2 (0.239) and group III (0.298) exhibit the lowest distances, reflecting strong purifying selection and conserved roles in light signaling. Group IV B2 (0.638) and group II B2 (0.856) display the highest divergence, indicative of adaptive evolution. Intermediate diversity is observed in group I B1 (0.58) and group IV B1 (0.409). Between-group mean distances depicted evolutionary relationships between subgroups. Closest pairs: group I B2 and group IV B1 (0.621): Shared regulatory roles in conserved pathways. Group I B2 and group III (0.628): Functional overlap in photomorphogenesis. Most divergent pairs: group II B2 and group IV B2 (1.120): Ancient divergence aligning with distinct roles. Group II B2 and group III (1.099): Early functional separation. Evolutionary pathway schematic model of B-box domain diversification. Ancestral Core: group III and group I B2 form the conserved regulatory hub for light signaling. The ancestral lineage gives rise to group IV and group V. Early Divergence: group II B2 splits, followed by group II B1, adapting to stress responses. **c** A quantitative evolutionary analysis of B-box group IV subgroups. Within-group mean distances: The results illustrating genetic diversity within subgroups of B-box group IV and other key BBX groups. Group IV B2b (0.107) and group IV B1b (0.14) exhibit the lowest distances, indicating strong purifying selection and functional constraint. In contrast, group II B2 (0.857) displays the highest divergence, reflecting adaptive evolution or lineage-specific pressures. Intermediate diversity is observed in a-group IV B1 (0.393) and a-group IV B2 (0.508). Between-group mean distances highlighting evolutionary relationships between subgroups. Closest pairs include group I B2 and group IV B1b (0.612) group III and group IV B1b (0.621), suggesting shared ancestry or functional convergence. Most divergent pairs include group II B2 and group IV B2a (1.143) and group II B2 and group IV B2b (1.028), indicative of ancient divergence or independent evolutionary trajectories. Proposed evolutionary pathway: The evolutionary trajectory of BBX proteins begins with group III and group I B2 forming the ancestral core, group II B2 diverged early, followed by group II B1. Subsequently, group IV B2b emerged from the ancestral lineage, giving rise to group IV B1a and group V. Finally, within group IV, group IV B2b and group IV B2a diverged, with group IV B2b retaining high conservation (low mean distance = 0.107) and group IV B2a exhibiting greater divergence (mean distance = 0.508).

To further dissect evolutionary patterns, we analyzed mean genetic distances for B1 and B2 domains independently (Fig. 4b). This domain-specific approach revealed distinct conservation and divergence dynamics across groups. For the mean distances within groups, group I B2 and group III B-box exhibited the lowest distances, indicating strong purifying selection and functional constraints, potentially due to critical roles in conserved processes such as light signaling. Group I B1, group II B1, group IV B1, group V B-box showed intermediate divergence, reflecting balanced evolutionary pressures and partial functional specialization. Group IV B2 and group II B2 displayed the greatest divergence, likely driven by adaptive evolution or lineage-specific neofunctionalization (Table S5). In case of group IV B2 domain, the likely explanation for higher within group mean distance is the sequence difference between subgroups a and b. For between groups mean distances, group I B2 and group IV B1 indicated a strong evolutionary link between B2 of group I and B1 of group IV. Group I B2 and group III B-box had a close relationship, implying functional overlap in conserved pathways. Group III B-box and group IV B1 shared ancestry highlights potential co-option of B1 domains in group IV for novel roles. Group I B1 and group III B-box showed moderate linkage, possibly reflecting ancestral domain shuffling. Group IV B1 and group V suggested functional convergence in stress or developmental regulation. Group II B2 and group III B-box had minimal evolutionary relationship, underscoring early divergence and functional separation. Group II B2 and group IV B2 showed extreme divergence points to ancient splits or independent evolutionary trajectories, aligning with distinct roles (e.g., group II B2 in stress vs. group IV B2 in flowering; Table S6). Overall, the group II B2 has highest values (over 1.05) when compared to all other B-box domains suggesting extensive adaptive evolution or lowered selection pressure. This is in line with phylogenetic interference and also lower conservation based on the the PWM.

Intriguingly, our phylogenetic analysis of B-box domains revealed a distinct subclade within group IV B2 that diverged from other group IV B2 sequences (Fig. 4c), prompting its classification as b-group IV B2, with the remaining group IV B2 sequences designated as a-group IV B2. Similarly, we subdivided group IV into subgroups (group IV a-B1, b-B1, a-B2, b-B2) to refine evolutionary insights. Within-group mean distances highlighted stark conservation contrasts: b-group IV B2, b-group IV B1, group I B2, and group III B-box emerged as the most conserved subgroups, indicative of strong functional constraints or recent divergence. Moderate diversity was observed in a-group IV B1, group II B1, group I B1, group V B-box, and a-group IV B2. In contrast, group II B2 exhibited the highest divergence, suggesting extensive adaptive evolution (Table S7). The within-group mean distances were similar as when analysed with combined group IV domain subgroup sequences. Between-group mean distances (Supplementary Table S8) further clarified evolutionary ties, group I B2 and group IV B1b showed strong linkage, likely reflecting shared regulatory roles. Group III B-box and group IV B1b, group I B2 and group III B-box were also close ancestry, underscoring conserved functions in light signaling. Group IV B1a and group IV B1b were tight intra-group homology, typical of recent duplication. Group I B1 and group IV B1b and group IV B1a and group V B-box could be partial homology, suggesting potential functional redundancy. Group II B2 and group IV B2a, group II B2 and group IV B2b demonstrated ancient divergence, aligning with independent evolutionary trajectories (Table S8). These analyses suggest that the evolution of the individual B-boxes is highly dynamic and could be indicating subfunctionalization of the B-boxes and the BBX TFs.

### The CCT domain

The hallmark of BBXs categorized into group I, II and III is the presence of the conserved CCT domain. It is characterized by an approximately 42 amino acid-long PWM, and the profile HMM for PFAM ID PF06203 provides a slightly longer TFBM of 44 residues. (Fig. 5a). The high conservation of residues in the CCT domain such as R, L, I, V, A, E, F might reflect their critical functional roles for BBXs. Our analysis identified several BBXs belonging to groups I, II or III not containing an identifiable CCT domain (Table S2). In certain species like in *Nelumbo nucifera*, BBX members without the CCT domain were paired with BBXs, that retained it, suggesting secondary loss in some members after divergence from a common ancestor that originally possessed this domain (Fig. 5b). To trace the evolutionary history of the CCT domain, we calculated mean distances based on sequence divergence within groups I-III (Table S9 and S10), revealing evolutionary relationships between the CCT domains of BBX groups (Fig. 5c). Group III showed strong conservation, suggesting strong purifying selection to maintain critical functions. In comparison, group I displayed also relatively high conservation, suggesting strong functional constraints while group II exhibited lower conservation, suggesting relaxed selection, adaptive diversification, or older divergence. Additionally, phylogenetic analysis of CCT domains (Fig. 5d, S7 and S8) showed distinct evolutionary relationships, each group as conserved clade with structural features shared across species likely reflecting ancestral functional roles. Lineage-specific changes could be associated with functional diversification. For instance, *O. sativa* and *B. distachyon* classified in Group III of the global BBX phylogeny, shifted to Group I in the CCT domain tree. This phylogenetic replacement suggests either convergent evolution of the CCT domain in grasses or divergent evolutionary constraints acting on the domain compared to the full-length BBX protein.

**Figure 5.**
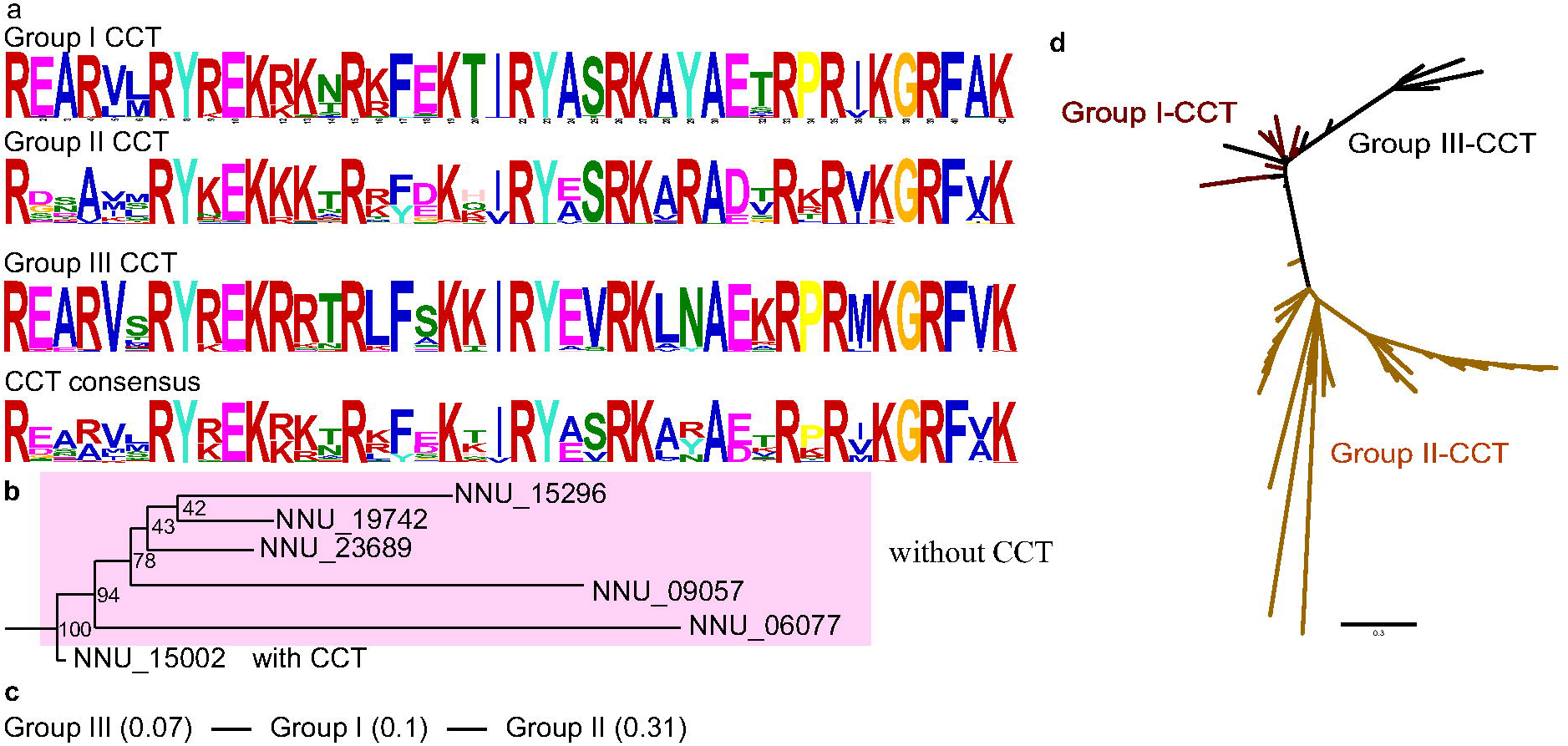
Evolutionary and Structural Analysis of the CCT Domain in BBX Proteins. A. The MEME sequence logos of CCT domain. Sequence logos generated by MEME illustrate the conservation patterns of CCT domains in BBX proteins. Overall CCT logos highlights conserved residues critical for nuclear localization and protein-protein interactions. And group-specific logos for groups I, II and III. Overall CCT: R-X5-R-Y-R/K-E-K-R/K-K/R-X-R-X3-K-X-I/V-R-Y-A/E-S/V-R-K-X2-A-E/D-X-R-X-R-X-K-G-R-F-V /A-K B. Phylogenetic tree highlighting CCT domain dynamics in group I. *Nelumbo nucifera*, members lacking the CCT domain (e.g., NNU_06077, NNU_09057, NNU_23689, NNU_19742annotation, NNU_15296) cluster with CCT-retaining homologs (NNU_15002), indicating lineage-specific domain loss. C. Evolutionary Pathway of CCT Domains. groups I and III exhibit low mean sequence divergence, forming the conserved regulatory hub. Group II showed higher mean distance, as an early divergent group. D. phylogenetic tree for CCT domain. Some BBX proteins classified in group III in the full-length BBX tree cluster within group I in the CCT domain tree.

### C-terminal motifs of BBXs

In addition to B-box and CCT domains, some BBXs contain conserved C-terminal motifs like the valine-proline motif (VP) (Holm 2001; Kirschner 2025) and Motifs 1 to 7. The VP motif critical for COP1-mediated ubiquitination, has been founded in groups I, III and IV (Fig. S1 and Table S2). Sequences from red algae species used in our analyses did not contain the VP domain, but the BBXs from green algae *Volvox* and *Coccomyxa* did contain VP motif indicating that the origin of this motif has already been present before the divergence of Chlorophyta and Streptophyta.

A previous study (Job et al. 2018) highlighted the functional importance of C-terminal domains in BBX proteins, particularly in light signaling and photomorphogenesis. The authors propose that Motif 6 (M6), a conserved sequence element within the C-terminal acts as a critical determinant of BBX protein activity by mediating interactions with HY5—a central transcription factor in light-regulated development. To investigate this hypothesis, we conducted a phylogenetic subclade-specific alignment of BBX21 homologs from group IV using MAFFT. Our analysis revealed that M6 is strikingly conserved across the BBX21 subclade (Fig. S9), with invariant residues such as glycine (G), arginine (R), aspartic acid (D), glutamic acid (E), serine (S), proline (P), isoleucine (I) tyrosine (Y) tryptophan (W) and leucine (L) forming a putative interaction interface, The presence of a mix of charged (R, D, E), polar (S, Y), and hydrophobic (I, W, L, P) residues is a classic signature of a protein–protein interaction surface, enabling electrostatic, hydrogen bonding (Teichmann 2002; Pawson and Nash 2003). This conservation suggests M6 may serve as a functional hotspot for HY5 binding or downstream signal transduction.

### BBX gene family evolution

To estimate significant expansions and contractions in BBX gene family in angiosperms, BBXs were analyzed with CAFE5 in eight species, the basal angiosperm *Amborella trichopoda*, the dicots *Aquilegia coerulea*, *Arabidopsis thaliana*, *Populus trichocarpa*, *Vitis vinifera* and the monocots *Brachypodium distachyon*, *Oryza sativa* and *Spirodela polyrhiza*. The analyses were carried out for all BBXs defined as a single orthogroup and separately for BBXs split into five subgroups based on their position in the phylogenetic ML tree (Fig. 1). In the CAFE5 analyses, multiple orthogroups were found to be significantly expanded or contracted in all species and ancestral nodes (Fig 6a). Also, BBXs had significant changes when analysed as one group. A significant expansion in *Populus trichocarpa* (P-value 1.04736e-06) was likely due to preferential retention following whole genome duplication in Salicaceae. When BBXs were analysed as five subgroups, groups I, II, III and V had significant changes. Group IV was close to significant (P-value 0.05) (Fig 6b-f). These results highlight the evolutionary plasticity of the BBX family in angiosperms, which could potentially be linked to ecological diversification and functional redundancy or innovation across subgroups. The summary of all the results of the clustering of whole BBX protein sequences together with the individual analyses for the domains within the BBXs led to the hypothesis on BBX evolution presented in Fig 7.

**Figure 6.**
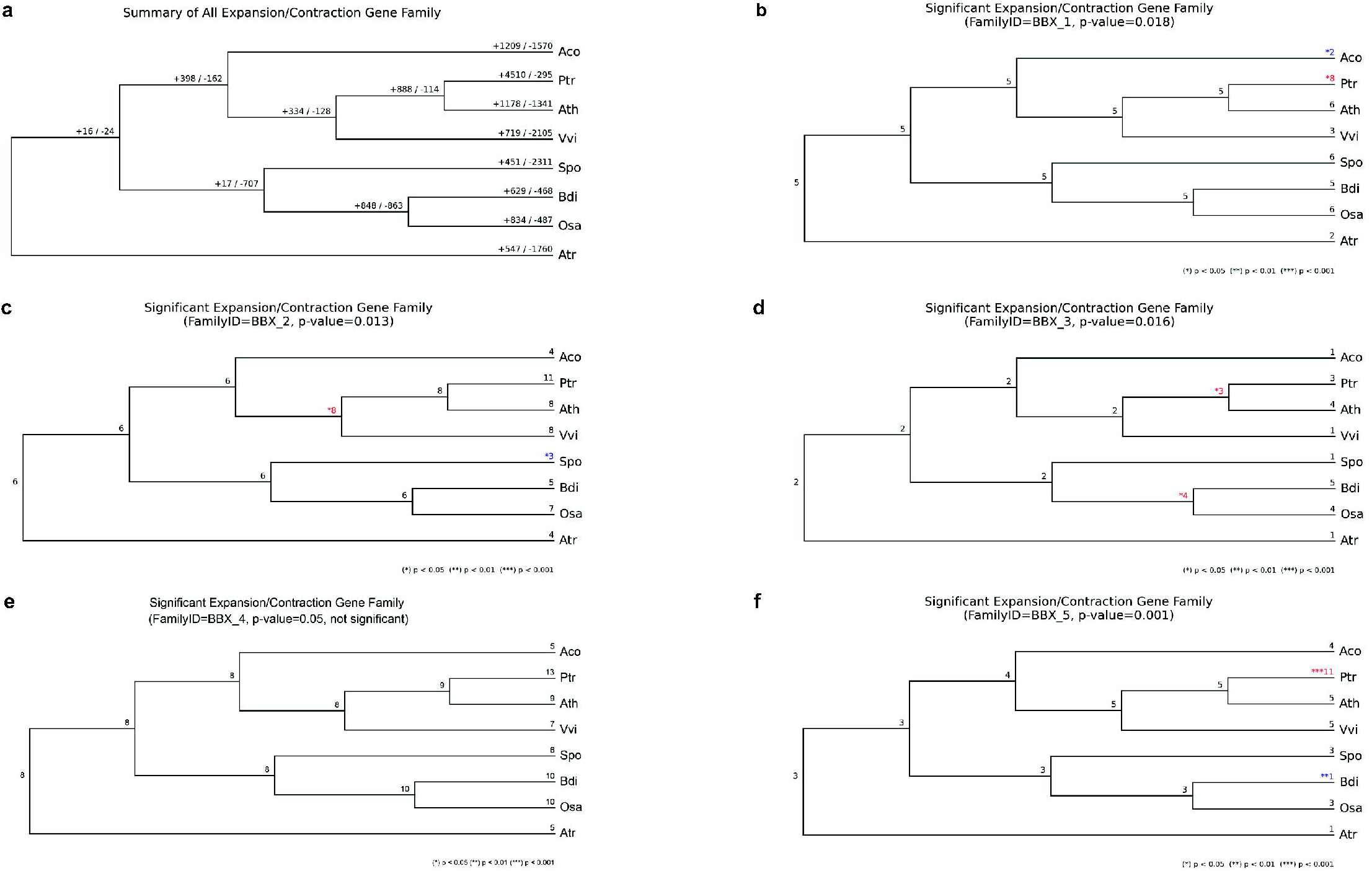
Gene family expansions and contractions based on CAFE5 analyses. **a** Overall gene family changes in eight angiosperm species. (B) Gene family changes in BBXs analysed as one group. The BBXs were also analyzed as five subgroups based on the phylogenetic grouping. The subgroup I (**b)**, II (**c**), III (**d**), IV (**e** and V (**f**) had significant changes. Species abbreviations: Aco *Aquilegia coerulea*, Ath *Arabidopsis thaliana*, Atr *Amborella trichopoda*, Bdi *Brachypodium distachyon*, Osa *Oryza sativa*, Ptr *Populus trichocarpa*, Spo *Spirodela polyrhiza* and Vvi *Vitis vinifera*.

**Figure 7.**
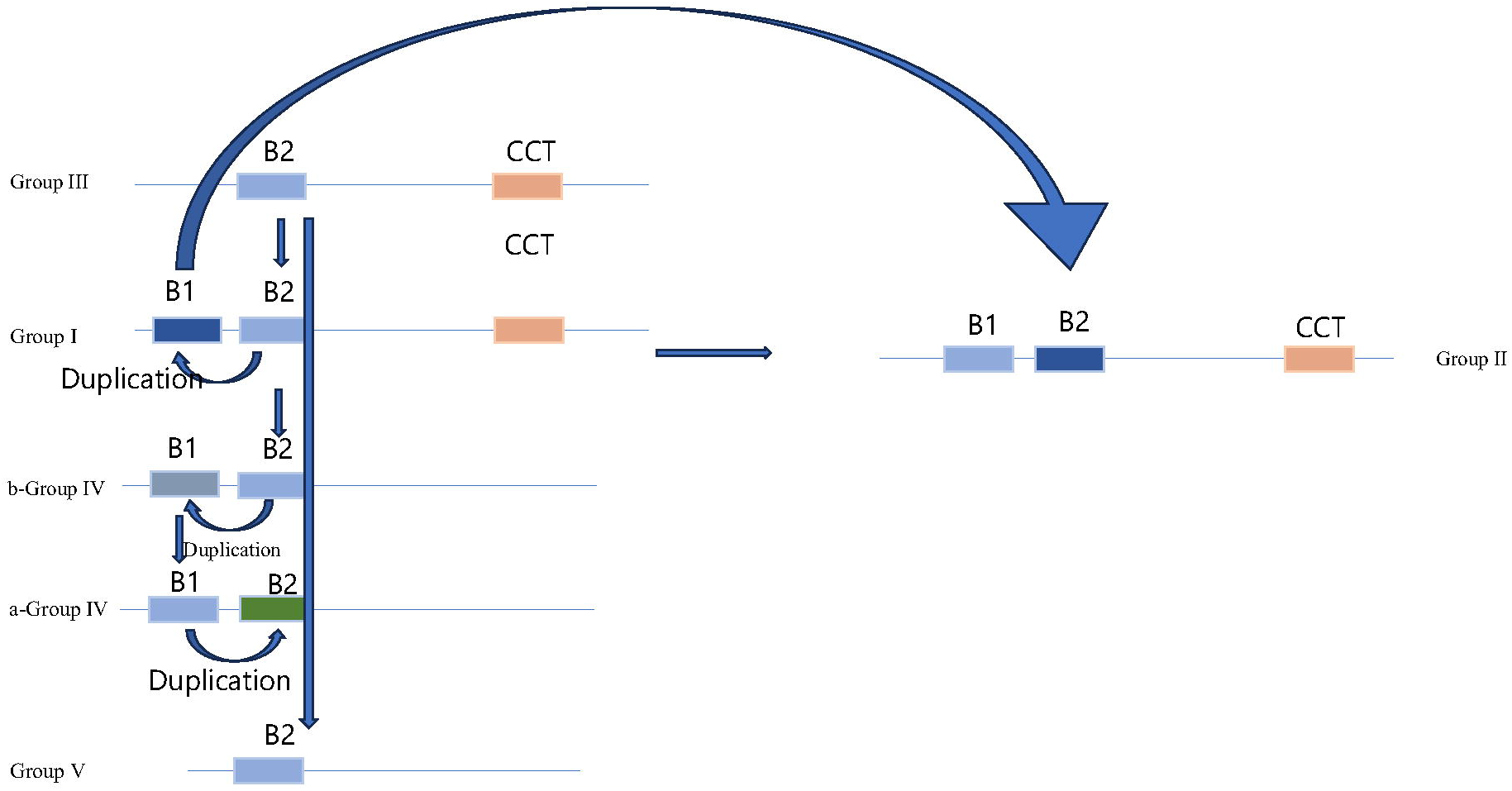
Hypothetical evolutionary model of B-box evolution history. Based on phylogenetic tree topology and mean distance analysis, this figure proposes a phylogenetically informed model for the evolutionary history of B-box domains in BBX proteins. Proposed evolutionary pathway: Group III is hypothesized as the ancestral source of the B-box domain (BBOX), representing the earliest functional form. The ancestral B-box domain from group III was inherited by group I B2 (B-box2), which subsequently diverged. From group I B2, the domain spread to b-group4 B2 (a subgroup within group IV), which later gave rise to the truncated B-box in group V. Duplication Events and Functional Diversification: group I B2 underwent duplication, generating group I B1 (B-box1). Group I B1 diverged further, contributing to: group II B2 (via functional divergence) and group II B1 (via direct inheritance). Expansion in group IV: b-group4 B2 duplicated to form b-group4 B1, which then diversified into a-group4 B1. a-group4 B1 underwent additional duplication, producing a-group4 B2, completing the expansion within group IV.

## Discussion

BBX TFs are ancient and highly conserved not only in plants but also in other multicellular eukaryotes (Burroughs et al. 2011). Based on their importance to many physiological processes in Arabidopsis model, researchers have a strong interest in identifying homologous BBX proteins in plant species of agricultural importance. Here, we analysed the evolutionary relationships in model lineages for the BBX family of plant transcription factors. The selection of these species represents a broad evolutionary spectrum from algae and early diverging land plants to more recently diverged flowering plants. These lineages represent critical transitions in eukaryotic evolution, from unicellular to multicellular forms and from aquatic to terrestrial habitats.

Our maximum likelihood (ML)-based evolutionary analyses for full-length BBX protein sequences contained five main clusters. For model species Arabidopsis, these clusters mostly corresponded to previous studies with exceptions for AtBBX26 and AtBBX27. AtBBX26 had been previously assigned to group V (single B-box) but in ML phylogenetic estimation it was reclassified into group IV (B1/B2 domains, no CCT), where it forms a long branch, while AtBBX27 (B1/B2 with a truncated CCT motif) shifted from group V to group II (B1/B2/CCT). AtBBX26 and 27 might originate from gene fusions of gene conversion events in the *Brassicaceae*. This finding underscores the dynamic evolutionary trajectory of BBX proteins and provides new insights into their functional specialization.

In the ML tree for B-box domains, a split was observed within group I BBX proteins—where B2 domains form a single conserved clade, while B1 domains separate into two distinct subclades (AtBBX1-3 and AtBBX4-6). This suggests that functional and evolutionary pressures have differentially shaped the B1 domain. This divergence likely reflects adaptations to specialized roles in plant development or stress responses. Phylogeny revealed that group II BBX proteins, absent in red algae, diverged early, coinciding with terrestrial colonization. Their specialization in photomorphogenesis regulation likely arose as plants adapted to land-based light and stress cues (Cao et al., 2023). Our evolutionary analysis of BBX proteins has unveiled significant diversification within group IV, characterized by the emergence of two distinct subgroups (a and b). The two subgroups within group IV suggest rapid functional diversification, likely driven by selective pressures to fulfill specialized roles in plant development or environmental responses. The split of B2 domains into b-group IV B2 and a-group IV B2 highlights domain-specific evolution. This divergence may reflect structural or functional innovations. The functional diversification of group IV BBX proteins in light signaling underscores their pivotal role in balancing photomorphogenesis and skotomorphogenesis. Central to this regulatory complexity might be the motif 6 (M6) within the C-terminal domain, which appears to act as a molecular switch determining antagonistic roles among group IV members (Job et al. 2018). The M6 motif is enriched in residues critical for protein-protein interactions (e.g., L and E) (Seychell and Beck 2021). Group V BBX proteins exhibit a distinct structural feature: a truncated B-box domain (∼37 residues) compared to other groups (∼38 residues), with a notable absence of a conserved arginine or proline residue between positions 4–9. This truncation could have implications for their structure and function. Group V proteins may interact with unique partners or exhibit reduced binding affinity for canonical targets. Group V BBX proteins, characterized by a single B-box domain and the absence of a CCT domain, exhibit a unique functional strategy: they rely on heterodimerization with CCT-containing BBX proteins (e.g., group I, II, or III members) to mediate DNA binding and transcriptional regulation (Shin et al. 2023). This dependency underscores the modularity and cooperative nature of BBX regulatory networks, where domain-deficient members act as co-factors rather than autonomous transcription factors. This could lead to specialized roles, such as acting as competitive inhibitors of other BBX proteins or participating in stress-specific pathways. The VP motif, critical for COP1-mediated ubiquitination, was found to be present in groups I , III and IV in our dataset, implicating possibly a broader role in light signaling (Kirschner 2025). Its conservation across multiple clades suggests an ancestral role in light signaling, which has been retained in divergent lineages to fine-tune COP1-mediated processes such as photomorphogenesis, flowering time regulation, and stress responses.

The high conservation of residues in the CCT domain might reflect essential functional roles in the regulation of photoperiodic flowering and circadian rhythm (Ballerini and Kramer 2011) (Fig. 5). Basic residues R and L are often key components of nuclear localization signals (Christophe et al. 2000). They may interact with the phosphate backbone of DNA or RNA, facilitating proper positioning of transcriptional regulators on DNA. The hydrophobic amino acids I, V, and A likely contribute to forming the hydrophobic core of the CCT domain, which is essential for maintaining the correct folding and structural stability of the domain. Residues F and E are thought to be involved in mediating protein-protein interactions, which are crucial for transcriptional regulation (Philip et al. 2011). Notably, we identified *Nelumbo nucifera* loss of the CCT motif after divergence from a common ancestor containing a CCT domain. This likely reflects adaptive evolution, where CCT-free variants diverged in function or regulatory mechanisms while retaining other roles.

The dynamic evolutionary history of the BBX gene family in angiosperms, as revealed by CAFE5 analyses, underscores its remarkable plasticity and adaptability. The significant expansions observed in *Populus trichocarpa* for groups I and V aligns with its ecological resilience and capacity to thrive in diverse environments, where stress tolerance and rapid growth are critical. Such expansions may reflect selective pressures driving gene family diversification to enhance regulatory complexity, particularly in species exposed to fluctuating abiotic or biotic stressors.

The evolutionary history of B-box domains in BBX proteins, reconstructed through phylogenetic tree topology and mean distance analyses (Fig. 7), suggests that the ancestral B-box domain was likely most similar to group III (featuring a single B-box paired with a CCT domain) as the earliest functional form. Structurally, group III B-box domains share homology with group I B2 domains, particularly in conserved zinc-coordinating residues and hydrophobic cores, indicating that group I B2 inherited this ancestral domain and diverged to form a conserved clade. Functional expansion of group I B2 proteins included roles in photomorphogenesis and flowering (e.g., CO/BBX1), while retaining the CCT domain for DNA binding and nuclear localization. A duplication event in group I B2 generated group I B1 (B-box1), which further diverged into group II B2, evolving novel roles in circadian regulation or stress responses via functional divergence and group II B1, which retained ancestral B1 structural features but diverged in interaction surfaces. From group I B2, the B-box domain expanded into the conserved subgroup IV B2b, which later gave rise to group V. Subgroup IV B2b duplicated to form subgroup a-IV B1, while subsequent duplication within group IV produced subgroup a-IV B2, which exhibited significant divergence (mean distance = 0.508) compared to the highly conserved b-IV B2 (mean distance = 0.107). Evolutionary trajectory analyses based on mean distances indicate that group III B-box and group I B2 formed the ancestral core, followed by the divergence of group II B2 and then group II B1. Subgroup IV B2b subsequently emerged, giving rise to IV B1a and group V, while within group IV, IV B2a diverged from IV B2b, with the former showing marked structural divergence and the latter retaining ancestral conservation.

## Acknowledgements

The authors wish to acknowledge CSC – IT Center for Science, Finland, for computational resources. We acknowledge support from the Institute of Plant Molecular Biology, Biology Centre CAS (to MW). HJ acknowledges support from the Chinese Scholarship Council (File No. 201606180057).

## Data availability

All data used for the preparation of the manuscript has been included in the supplementary data. Supplementary Table 1 summarizes the genome versions used with all identifiers including Genbank.

## Competing interests

The authors declare no competing interests.

## Notes

### Competing Interest Statement

The authors have declared no competing interest.

