## Supplementary Figures for "BBX transcription factor evolution in the green plant lineage"

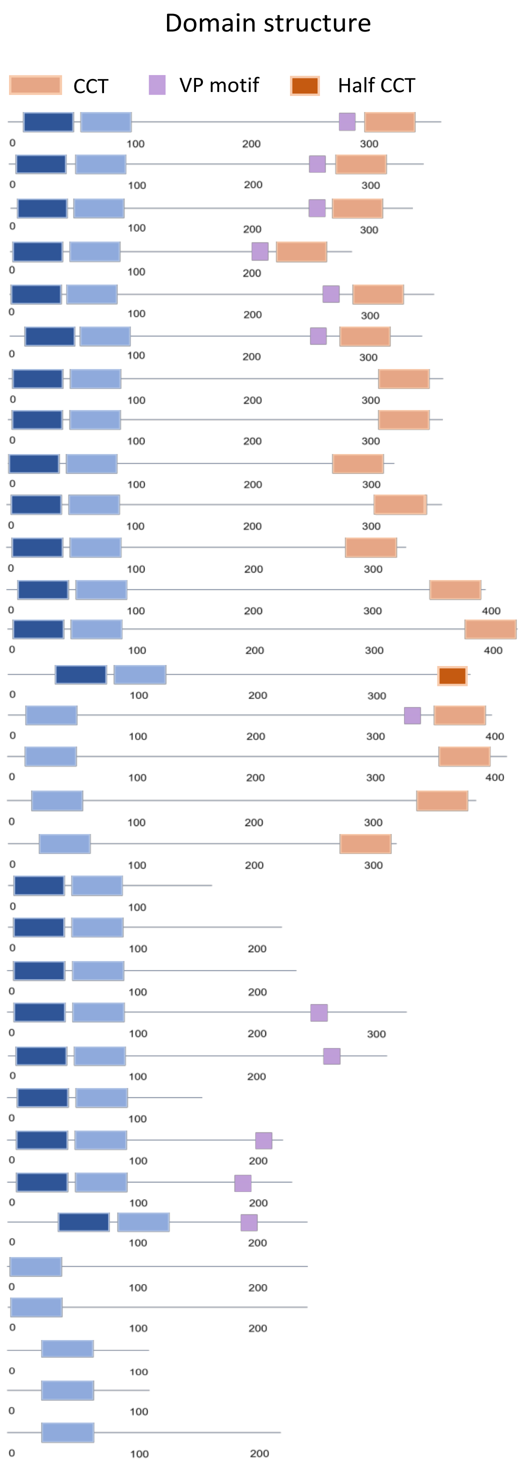


| AGI number | BBX name | Protein length | Structure group |
|---|---|---|---|
| AT5G15840 | BBX1 | 373 | I |
| AT5G15850 | BBX2 | 355 | I |
| AT3G02380 | BBX3 | 347 | I |
| AT2G24790 | BBX4 | 294 | I |
| AT5G24930 | BBX5 | 362 | I |
| AT5G57660 | BBX6 | 355 | I |
| AT3G07650 | BBX7 | 372 | II |
| AT5G48250 | BBX8 | 373 | II |
| AT4G15250 | BBX9 | 330 | II |
| AT3G21880 | BBX10 | 368 | II |
| AT2G47890 | BBX11 | 332 | II |
| AT2G33500 | BBX12 | 402 | II |
| AT1G28050 | BBX13 | 433 | II |
| AT1G68190 | BBX27 | 385 | II |
| AT1G68520 | BBX14 | 406 | III |
| AT1G25440 | BBX15 | 417 | III |
| AT1G73870 | BBX16 | 392 | III |
| AT1G49130 | BBX17 | 326 | III |
| AT2G21320 | BBX18 | 172 | IV |
| AT4G38960 | BBX19 | 226 | IV |
| AT4G39070 | BBX20 | 242 | IV |
| AT1G75540 | BBX21 | 331 | IV |
| AT1G78600 | BBX22 | 319 | IV |
| AT4G10240 | BBX23 | 162 | IV |
| AT1G06040 | BBX24 | 248 | IV |
| AT2G31380 | BBX25 | 238 | IV |
| AT1G60250 | BBX26 | 251 | IV |
| AT4G27310 | BBX28 | 223 | V |
| AT5G54470 | BBX29 | 215 | V |
| AT4G15248 | BBX30 | 117 | V |
| AT3G21890 | BBX31 | 121 | V |
| AT3G21150 | BBX32 | 225 | V |

**Figure S1** Domain architecture and classification of the Arabidopsis BBX protein family. The phylogenetic classification, domain organization, and protein lengths of Arabidopsis BBX proteins are categorized into five structural groups. Notable reclassifications include AtBBX26 (previously assigned to group V in prior studies, now placed in group IV) and AtBBX27 (originally categorized in group V, now classified in group II). Protein lengths of BBX5, BBX10, and BBX27 differ from previous reports and are highlighted in red. Abbreviations: AGI, Arabidopsis Genome Initiative.

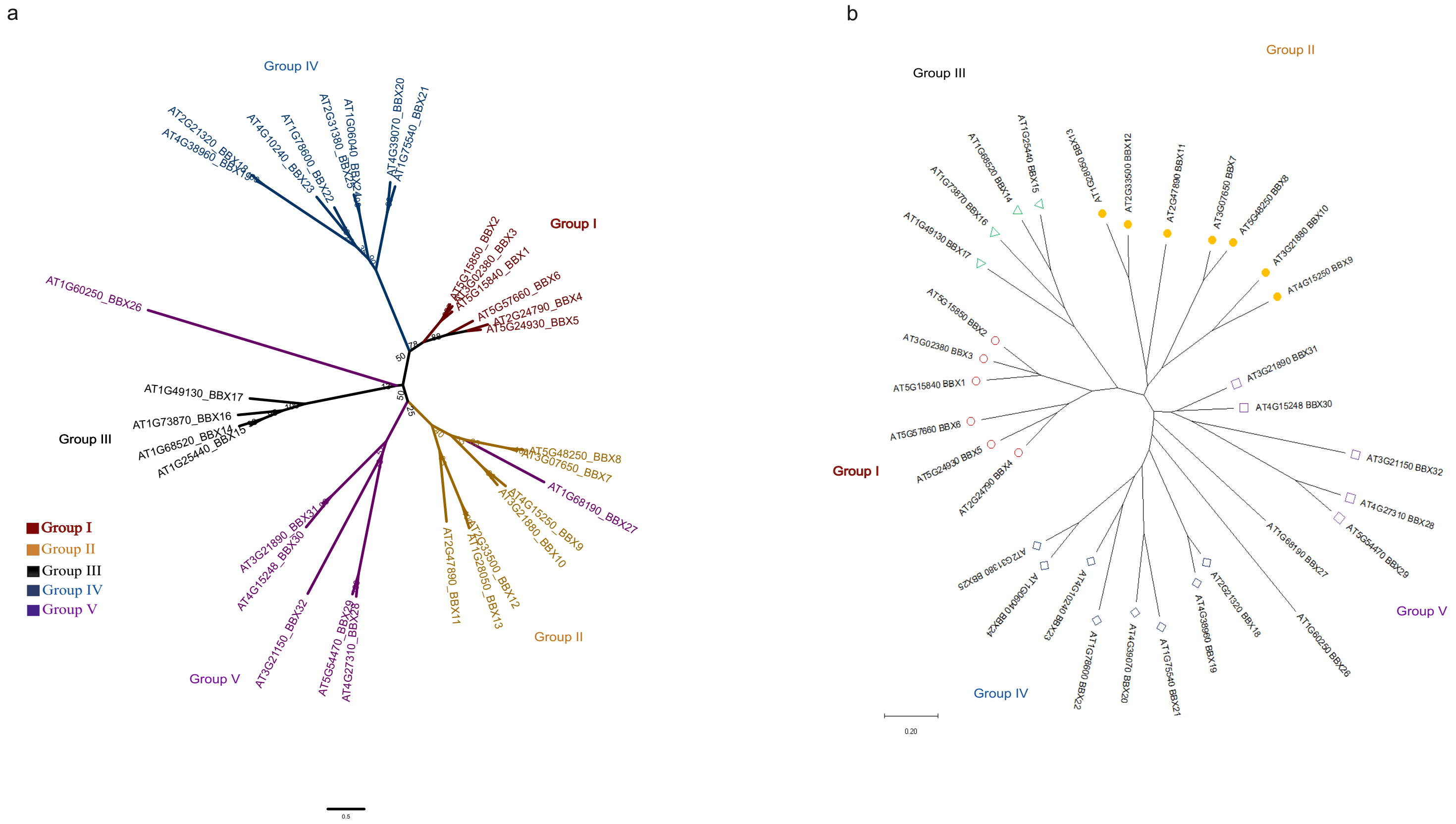


**Figure S2** Unrooted maximum likelihood phylogenetic tree and neighbor-joining (NJ) tree for Arabidopsis BBX proteins. A. Unrooted ML phylogenetic tree resolving five major clades (groups I–V), consistent with prior classifications based on domain architecture. Previously study AtBBX27 assigned to group V in domain based studies were repositioned to group II. AtBBX26 formed a long evolutionary branch within group III. B. NJ phylogeny reconstructed using the methodology from earlier studies. In contrast to the ML topology, AtBBX26 and AtBBX27 clustered near group V in the NJ tree, aligning with previous domain-based groupings.

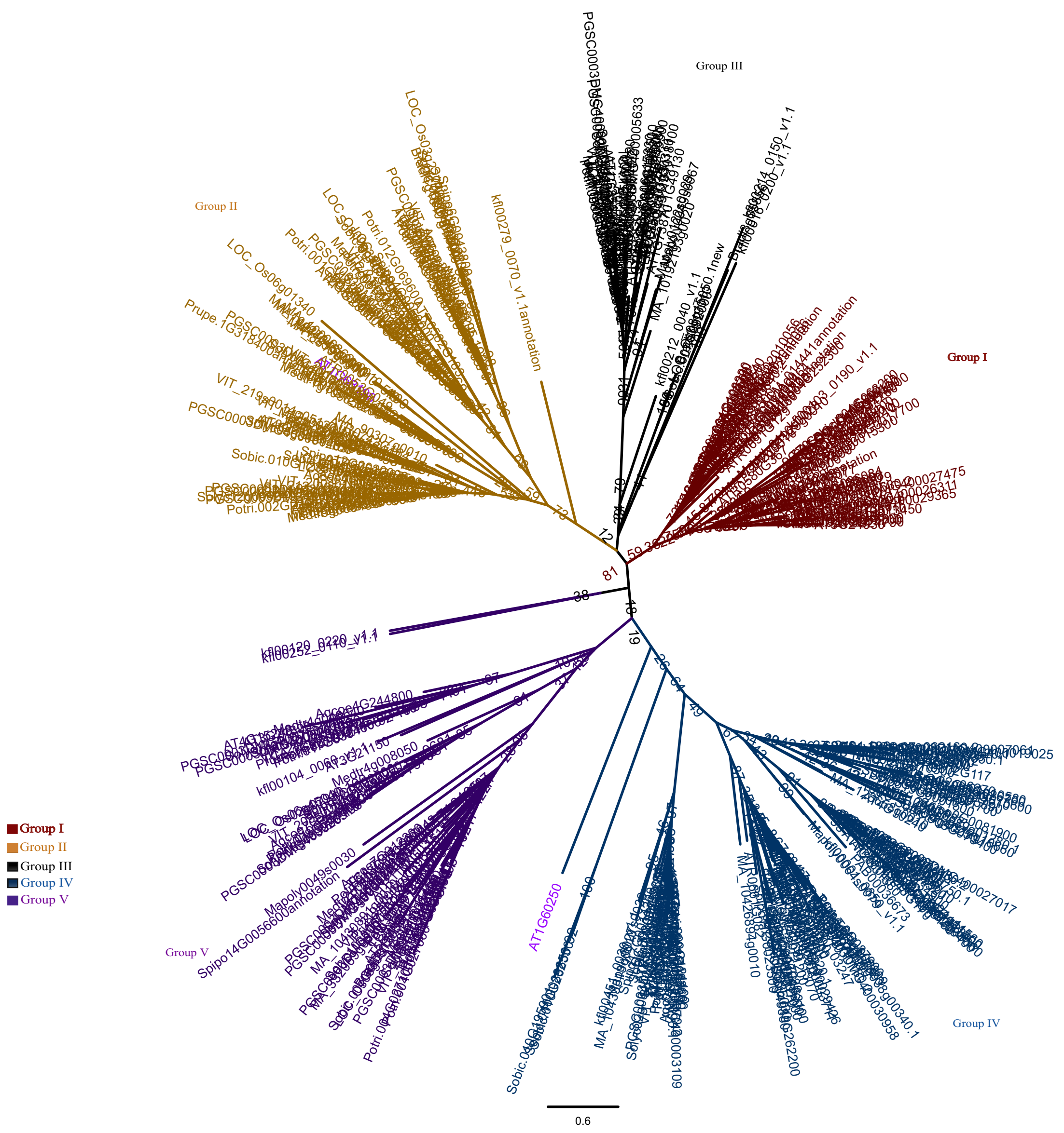


**Figure S3** Radial layout of Labeled unrooted maximum likelihood phylogenetic tree for BBX proteins. The tree replicates the topology from the global BBX tree presented in the main figure but includes terminal labels for all proteins. Phylogenetic groups (I–V) are color-coded as follows: group I (red), group II (yellow), group III (black), group IV (blue), and group V (dark purple). Notably, AtBBX26 and AtBBX27 are highlighted in light purple to distinguish their unique placement within the phylogeny.

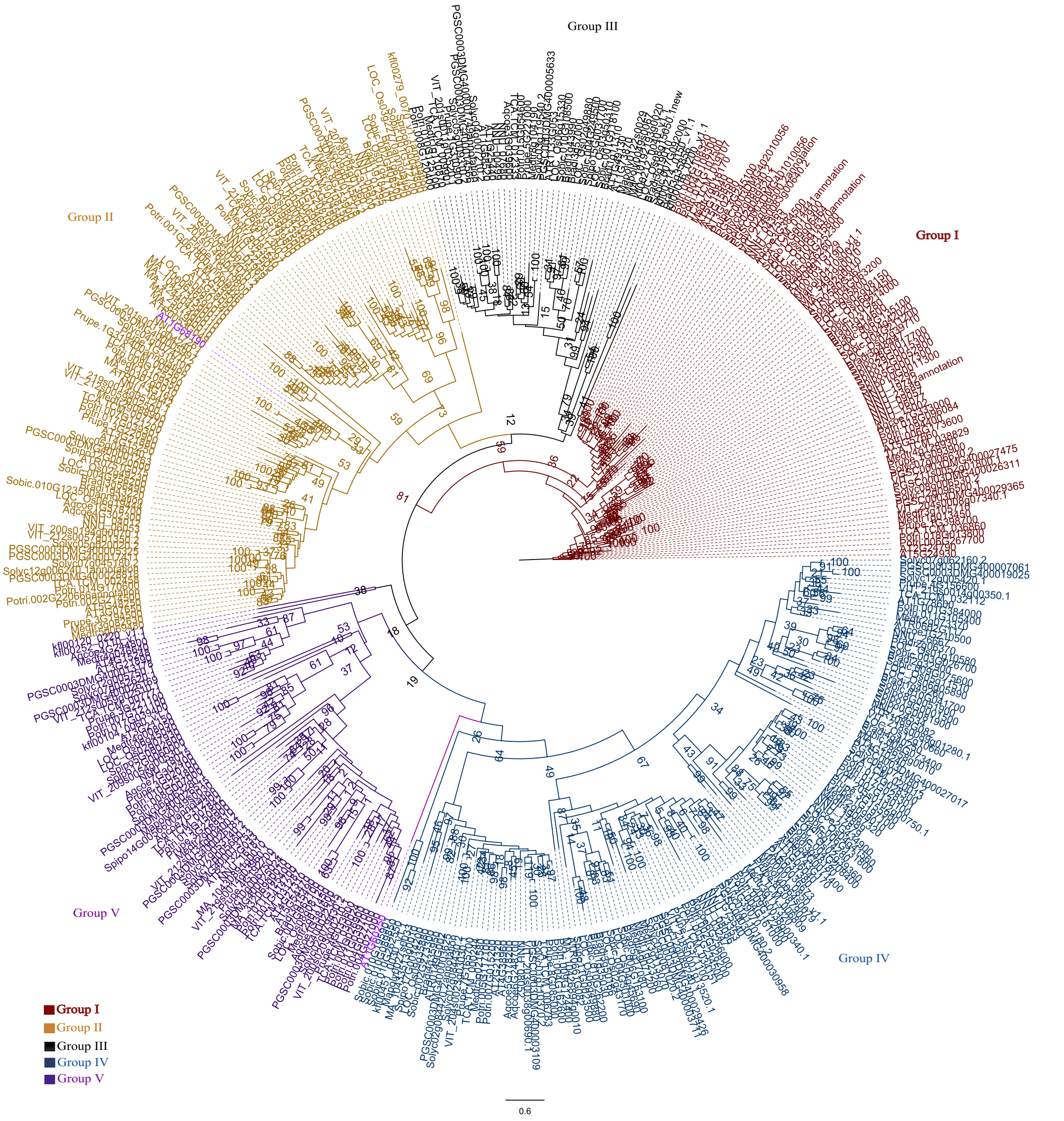


**Figure S4** Polar layout of Labeled unrooted maximum likelihood phylogenetic tree for BBX proteins. The same tree as the main figure with terminal labels for all proteins. Phylogenetic groups (I–V) are color-coded as follows: group I (red), group II (yellow), group III (black), group IV (blue), and group V (dark purple). Notably, AtBBX26 and AtBBX27 are highlighted in light purple to distinguish their unique placement within the phylogeny.

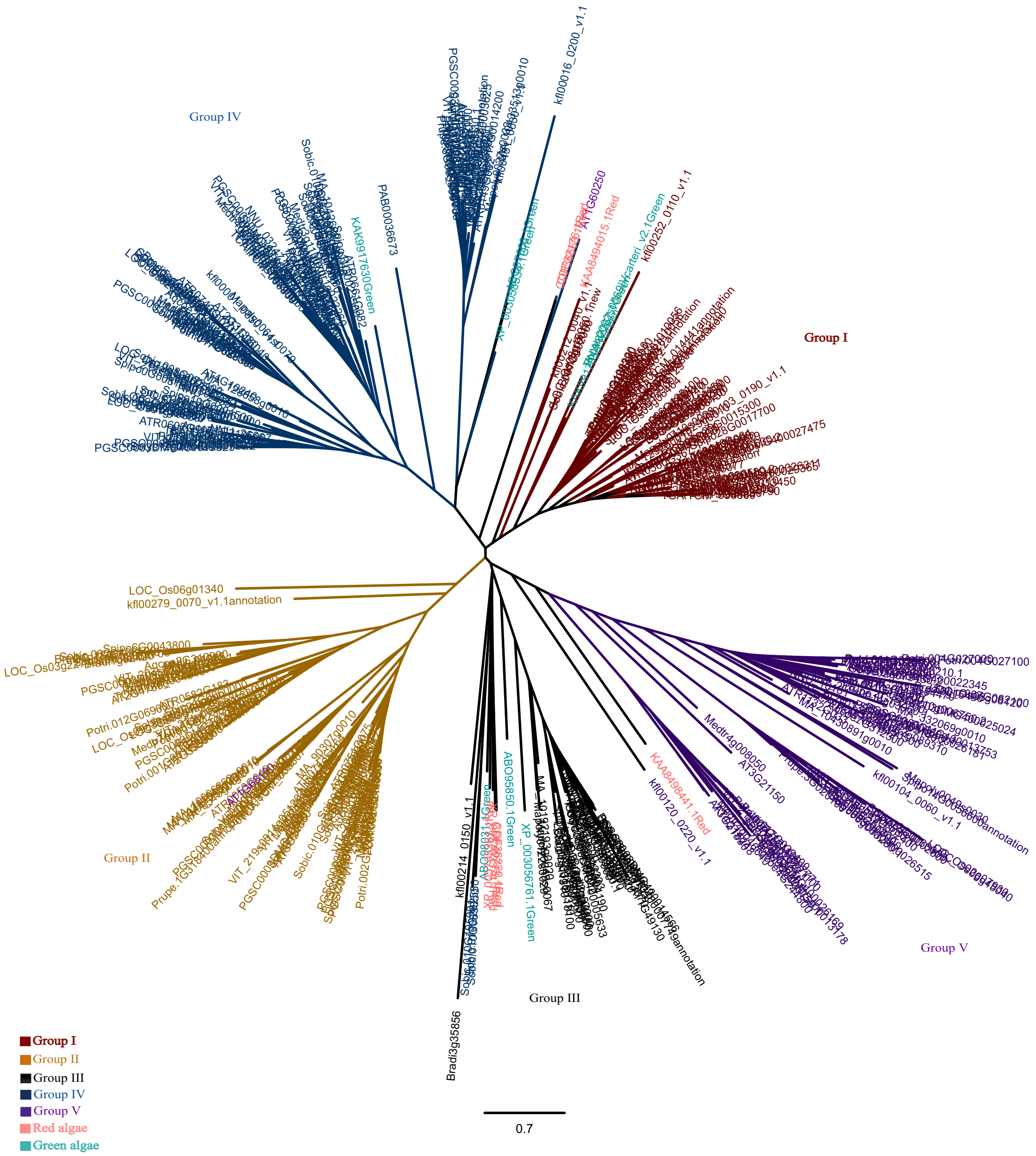


**Figure S5** Labeled unrooted maximum likelihood phylogenetic tree with algae for BBX proteins. The tree replicates the topology from the global BBX tree including algae species presented in the main figure but includes terminal labels for all proteins. Phylogenetic groups (I–V) are color-coded as follows: group I (red), group II (yellow), group III (black), group IV (blue), and group V (dark purple). AtBBX26 and AtBBX27 are highlighted in light purple to distinguish their unique placement within the phylogeny.

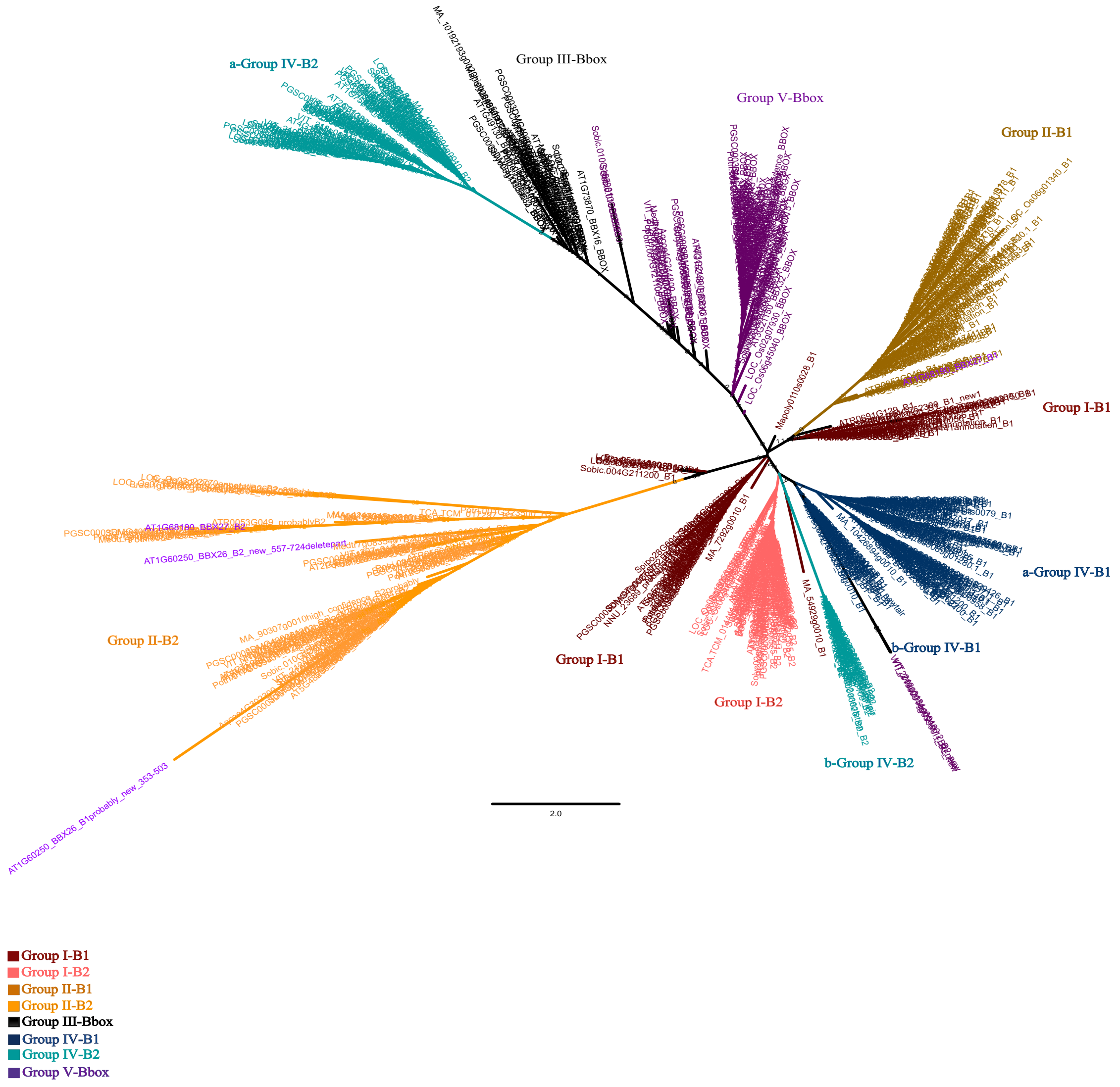


**Figure S6** Labeled unrooted maximum likelihood phylogenetic tree for B-box domain. The tree replicates the topology from the B-box domains tree presented in the main figure but includes terminal labels for all proteins. A radial layout showing the full topology. groups are colored to reflect domain classification: group I-B1 (dark red), group I-B2 (light red), group II-B1 (dark yellow), group II-B2 (light yellow), group III (black), group IV-B1 (dark blue), group IV-B2 (light blue), and group V (dark purple). AtBBX26 and AtBBX27 are highlighted in light purple to denote their distinct phylogenetic placement.

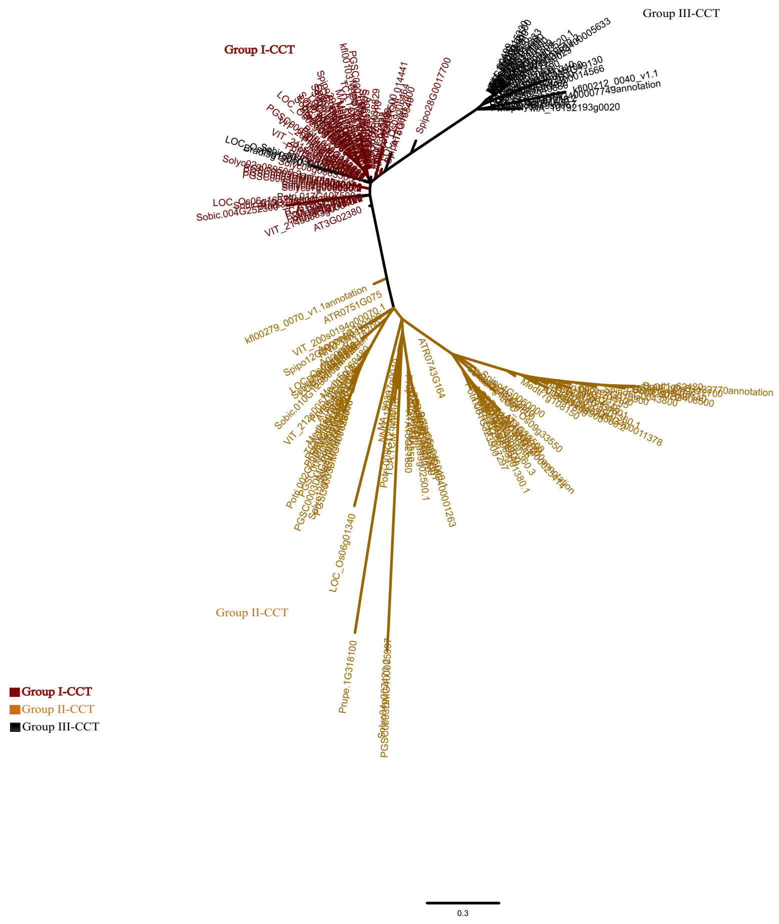


**Figure S7** Labeled unrooted maximum likelihood phylogenetic tree for CCT domain. The tree replicates the topology from the CCT domains tree presented in the main figure but includes terminal labels for group I- III proteins. A radial layout showing the full topology. Phylogenetic groups are colored as follows: group I (red), group II (yellow), group III (black). Some members in group I from Brachypodium distachyon, Oryza sativa, and Sorghum bicolor phylogenetically assigned to group I (labeled in black) cluster within group III in the global BBX tree (main text Figure).

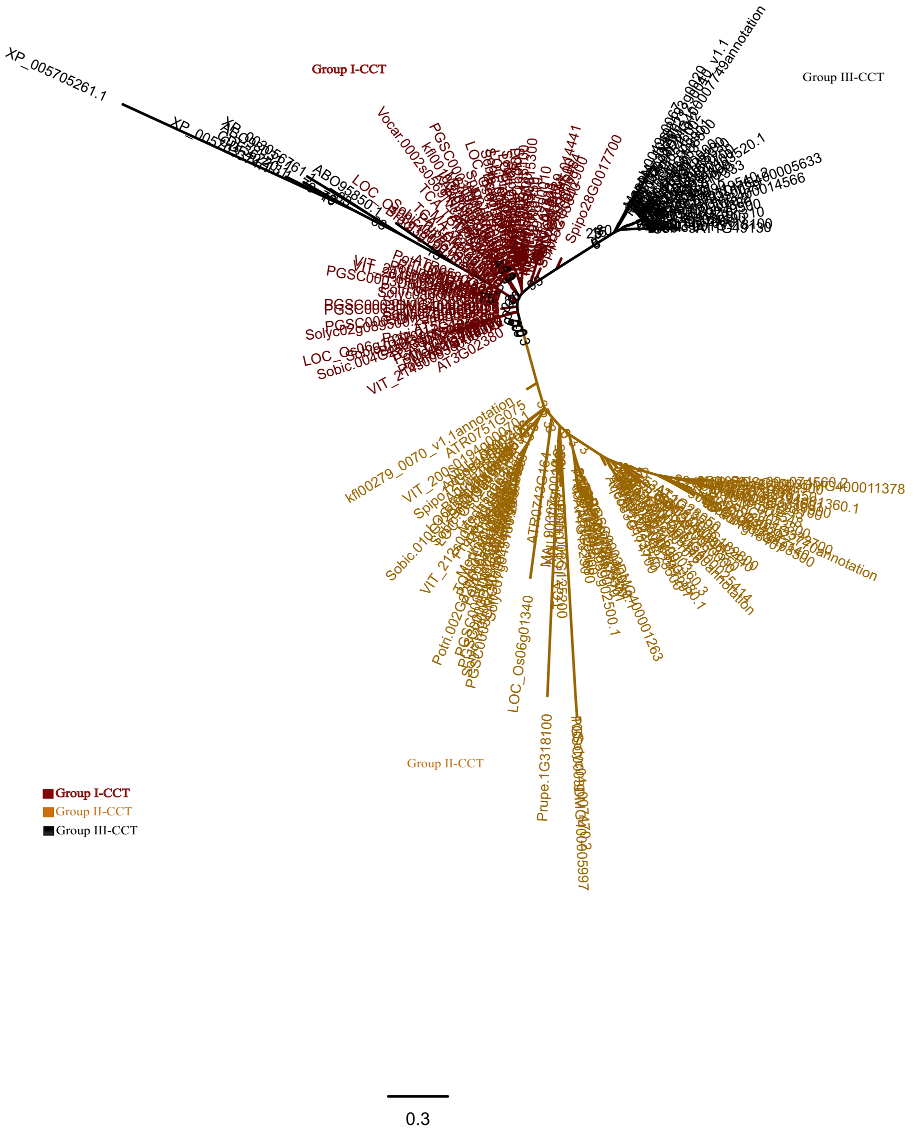


**Figure S8** Labeled unrooted maximum likelihood phylogenetic tree for CCT domain including algae. The tree replicates the topology from the CCT domains tree presented in the main figure but includes terminal labels for group I- III proteins. A radial layout showing the full topology. Phylogenetic groups are colored as follows: group I (red), group II (yellow), group III (black). Some members in group I from algae phylogenetically assigned to group I (labeled in black) cluster within group III in the global BBX tree (main text Figure).

```
Spipo11G0023500            ----------------------DSDGNT---GSTASATGGSRISEYLMKELPGWHVEDLL    169
ATR0661G082                N------------------------------SPAPLSSTSSSISEYLTHTIPGWRVDDFL    159
Medtr3g113070              ---------------------------------LHEETGNFTISEYLINTIPGWKFEDFL    141
Medtr1g023260              TTPSPT--------------------------SMEEGSGGSTISQYLIETLPGWQVDDFL    172
Medtr4g071200              --------------------SQSSFKEN----MVCDTVSTSSISEYLIETIPGYCMEDLF    175
Solyc04g081020.2           SP---VSGSVPQQ-------------QVSVAANIGENSYTSSISEYL-EMLPGWHVEELL    186
PGSC0003DMG400003711       LP---VSGSVPQ--------------QVSVAVNIGENSYTSSISEYL-EMLPGWHVEELL    185
Solyc12g089240.1           GA---VSSAVESVKVVKEKVG--GCNNNVQFVNGGGNNLTSSISEYL-EMLPGWHVEDFL    217
PGSC0003DMG400029426       GA---VSSVVESVKVVKEKVG--GCNNNVQLVNGGGNNLTSSISEYL-EMLPGWHVEDFL    218
AT1G75540                  LSAPPQSNKIQ-------PFSKINGGDA----SVNQWGSTSTISEYLMDTLPGWHVEDFL    195
Prupe.1G371100             STLTTN----------TANS--NKGGGIFVAHDGVGCGSTSSISEYLIETLPGWHVEDFL    208
Potri.005G234500           -----L--------SANTVI--NKDGDNLVTSEGFGSTTSSTISEYLMETLPGWHVEEFL    204
Potri.002G028200           -----L--------STNTEV--NKGGDNLVTNEGFGSTTSSTISEYLMETLPGWHVEDFL    205
TCA.TCM_034151             SPVSTT--------AAAVTN--KSGGDNLLANEGGGST--SSISEYLIEMLPGWHFEDFL    264
VIT_218s0001g13520.1       SSPTT----------AINSI--NKGGDASLT-SEGVST--SSISEYLIEMLPGWHVEDFL    195
AT4G39070                  SSSSTT--TSN-----CYYGIEENYHHVS--DSGSGSGCTGSISEYLMETLPGWRVEDLL    199
Solyc01g110180.2           ISSTTE--STH-----NYFHV------DY---VQEGSVSTSSISEYLTETLPGWHVEDFL    193
PGSC0003DMG400030958       ISSTTE--STP-----NYFQV------DY--HVQEGSVSTSSISEYLTETLPGWHVEDFL    193
Aqcoe6G256000              ST-----TSIPSKGGGGGSSGGHVASSTT--TTTSGDGSTSSIAEYLIETLPGWHVEDFL    197
NNU_08086                  P--------------AAATITTSKADDHP--AS--EGYSTSSITEYLMETLPGWQVEDFL    205
NNU_23353                  SPPPPP---PPPT--TTATITTHKSDDHP--AS--GGGSTSSISEYLIKMLPGWQVEDFL    211
Medtr3g117320              -------NIIPTSVSNENASSSCMVEDNM--ASDTGSVSTSSISEYLIETIPGYCFEDLL    183
VIT_203s0038g00340.1       IKPSKTSTKRPTSVSAGISNPTVKTAPAA--ASYKRDHDNQSISEYLMETLPGWRVDDFL    212
Potri.009G122000           SPPTAY-----------------SYDDNH--VSGGGSVSTSSISEYLETVVPGWRVDDFL    198
Prupe.8G087900             TSSSSY-----------------KTGENC--GSDNGSVSTSSISEYLMETLPGWHVEDFF    199
TCA.TCM_006809             LPSTTD-----------------KVEDNC--TSDTVSISTSSISEYLMETLPGWRVDDFL    200
Sobic.004G301000           --PPPS-----------------S--AAPATSHGSGSDNGSSISEYLIKTLPGWHVEDFL    196
Bradi3g50166               --------------------------------HGCGSSTSSSISEYLTKTLPGWHVEDFL    170
LOC_Os02g43170             --------------------------AAP-ATSHGGSSGSSSISEYLT-TLPGWHVEDFL    175
Sobic.006G163100           ------------------------------ATTTPSASDGSSISEYLTKTLPGWHVEDFL    181
Bradi5g17080               ------------------------------ATASASASDGSSISEYLTKTLPGWHVEDFL    174
LOC_Os04g45690             ------------------------------GTAAGSASDGSSISEYLTKTLPGWHVEDFL    170
Sobic.010G262200           ----A---------------------AKASALESGSVGGGSSISDYLTNICPGWRVDDLL    193
Bradi1g35030               ----PP---------------------VLNGV-GGGGGGGSSISEYLTNICPGWRVEDLL    181
LOC_Os06g49880             ----PL-----------------D--ASSNGAGGGGGVGGSSISDYLTTICPGWRVEDLL    193
                                                                     *::**    **: .::::
```

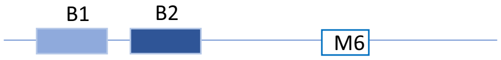


**Figure S9** Multiple sequence alignment of motif 6 in BBX group IV. The alignment highlights striking conservation of M6 within the BBX21 clade. Invariant residues including glycine (G), arginine (R), aspartic acid (D), glutamic acid (E), serine (S), proline (P), isoleucine (I), tyrosine (Y), tryptophan (W), and leucine (L) form a conserved structural motif, suggesting a functional interaction interface critical for molecular recognition or binding activity.
