## Supplementary material for "BBX transcription factor evolution in the green plant lineage": Data S1.docx

**Supplementary Data 1** Amino acid sequences (FASTA format) of all gene models analyzed without algae species.

>AT2G24790

MASSSRLCDSCKSTAATLFCRADAAFLCGDCDGKIHTANKLASRHERVWLCEVCEQAPAHVTCKADAAALCVTCDRDIHSANPLSRRHERVPITPFYDAVGPAKSASSSVNFVDEDGGDVTASWLLAKEGIEITNLFSDLDYPKIEVTSEENSSGNDGVVPVQNKLFLNEDYFNFDLSASKISQQGFNFINQTVSTRTIDVPLVPESGGVTAEMTNTETPAVQLSPAEREARVLRYREKRKNRKFEKTIRYASRKAYAEMRPRIKGRFAKRTDSRENDGGDVGVYGGFGVVPSF

>AT3G02380

MLKEESNESGTWARACDTCRSAACTVYCEADSAYLCTTCDARVHAANRVASRHERVRVCQSCESAPAAFLCKADAASLCTACDAEIHSANPLARRHQRVPILPLSANSCSSMAPSETDADNDEDDREVASWLLPNPGKNIGNQNNGFLFGVEYLDLVDYSSSMDNQFEDNQYTHYQRSFGGDGVVPLQVEESTSHLQQSQQNFQLGINYGFSSGAHYNNNSLKDLNHSASVSSMDISVVPESTASDITVQHPRTTKETIDQLSGPPTQVVQQLTPMEREARVLRYREKKKTRKFDKTIRYASRKAYAEIRPRIKGRFAKRIETEAEAEEIFSTSLMSETGYGIVPSF

>AT5G24930

MASKLCDSCKSATAALYCRPDAAFLCLSCDSKVHAANKLASRHARVWMCEVCEQAPAHVTCKADAAALCVTCDRDIHSANPLARRHERVPVTPFYDSVSSDGSVKHTAVNFLDDCYFSDIDGNGSREEEEEEAASWLLLPNPKTTTTATAGIVAVTSAEEVPGDSPEMNTGQQYLFSDPDPYLDLDYGNVDPKVESLEQNSSGTDGVVPVENRTVRIPTVNENCFEMDFTGGSKGFTYGGGYNCISHSVSSSSMEVGVVPDGGSVADVSYPYGGPATSGADPGTQRAVPLTSAEREARVMRYREKRKNRKFEKTIRYASRKAYAEMRPRIKGRFAKRTDTNESNDVVGHGGIFSGFGLVPTF

>AT5G57660

MGFGLESIKSISGGWGAAARSCDACKSVTAAVFCRVDSAFLCIACDTRIHSFTRHERVWVCEVCEQAPAAVTCKADAAALCVSCDADIHSANPLASRHERVPVETFFDSAETAVAKISASSTFGILGSSTTVDLTAVPVMADDLGLCPWLLPNDFNEPAKIEIGTENMKGSSDFMFSDFDRLIDFEFPNSFNHHQNNAGGDSLVPVQTKTEPLPLTNNDHCFDIDFCRSKLSAFTYPSQSVSHSVSTSSIEYGVVPDGNTNNSVNRSTITSSTTGGDHQASSMDREARVLRYREKRKNRKFEKTIRYASRKAYAESRPRIKGRFAKRTETENDDIFLSHVYASAAHAQYGVVPTF

>AT5G15840

MLKQESNDIGSGENNRARPCDTCRSNACTVYCHADSAYLCMSCDAQVHSANRVASRHKRVRVCESCERAPAAFLCEADDASLCTACDSEVHSANPLARRHQRVPILPISGNSFSSMTTTHHQSEKTMTDPEKRLVVDQEEGEEGDKDAKEVASWLFPNSDKNNNNQNNGLLFSDEYLNLVDYNSSMDYKFTGEYSQHQQNCSVPQTSYGGDRVVPLKLEESRGHQCHNQQNFQFNIKYGSSGTHYNDNGSINHNAYISSMETGVVPESTACVTTASHPRTPKGTVEQQPDPASQMITVTQLSPMDREARVLRYREKRKTRKFEKTIRYASRKAYAEIRPRVNGRFAKREIEAEEQGFNTMLMYNTGYGIVPSF

>AT5G15850

MLKVESNWAQACDTCRSAACTVYCRADSAYLCSSCDAQVHAANRLASRHERVRVCQSCERAPAAFFCKADAASLCTTCDSEIHSANPLARRHQRVPILPISEYSYSSTATNHSCETTVTDPENRLVLGQEEEDEDEAEAASWLLPNSGKNSGNNNGFSIGDEFLNLVDYSSSDKQFTDQSNQYQLDCNVPQRSYGEDGVVPLQIEVSKGMYQEQQNFQLSINCGSWGALRSSNGSLSHMVNVSSMDLGVVPESTTSDATVSNPRSPKAVTDQPPYPPAQMLSPRDREARVLRYREKKKMRKFEKTIRYASRKAYAEKRPRIKGRFAKKKDVDEEANQAFSTMITFDTGYGIVPSF

>AT1G28050

MSSSERVPCDFCGERTAVLFCRADTAKLCLPCDQQVHTANLLSRKHVRSQICDNCGNEPVSVRCFTDNLILCQECDWDVHGSCSVSDAHVRSAVEGFSGCPSALELAALWGLDLEQGRKDEENQVPMMAMMMDNFGMQLDSWVLGSNELIVPSDTTFKKRGSCGSSCGRYKQVLCKQLEELLKSGVVGGDGDDGDRDRDCDREGACDGDGDGEAGEGLMVPEMSERLKWSRDVEEINGGGGGGVNQQWNATTTNPSGGQSSQIWDFNLGQSRGPEDTSRVEAAYVGKGAASSFTINNFVDHMNETCSTNVKGVKEIKKDDYKRSTSGQVQPTKSESNNRPITFGSEKGSNSSSDLHFTEHIAGTSCKTTRLVATKADLERLAQNRGDAMQRYKEKRKTRRYDKTIRYESRKARADTRLRVRGRFVKASEAPYP

>AT2G33500

MGTSTTESVVACEFCGERTAVLFCRADTAKLCLPCDQHVHSANLLSRKHVRSQICDNCSKEPVSVRCFTDNLVLCQECDWDVHGSCSSSATHERSAVEGFSGCPSVLELAAVWGIDLKGKKKEDDEDELTKNFGMGLDSWGSGSNIVQELIVPYDVSCKKQSFSFGRSKQVVFEQLELLKRGFVEGEGEIMVPEGINGGGSISQPSPTTSFTSLLMSQSLCGNGMQWNATNHSTGQNTQIWDFNLGQSRNPDEPSPVETKGSTFTFNNVTHLKNDTRTTNMNAFKESYQEDSVHSTSTKGQETSKSNNIPAAIHSHKSSNDSCGLHCTEHIAITSNRATRLVAVTNADLEQMAQNRDNAMQRYKEKKKTRRYDKTIRYETRKARAETRLRVKGRFVKATDP

>AT2G47890

MEAEEGHQRDRLCDYCDSSVALVYCKADSAKLCLACDKQVHVANQLFAKHFRSLLCDSCNESPSSLFCETERSVLCQNCDWQHHTASSSLHSRRPFEGFTGCPSVPELLAIVGLDDLTLDSGLLWESPEIVSLNDLIVSGGSGTHNFRATDVPPLPKNRHATCGKYKDEMIRQLRGLSRSEPGCLKFETPDAEIDAGFQFLAPDLFSTCELESGLKWFDQQDHEDFPYCSLLKNLSESDEKPENVDRESSVMVPVSGCLNRCEEETVMVPVITSTRSMTHEINSLERNSALSRYKEKKKSRRYEKHIRYESRKVRAESRTRIRGRFAKAADP

>AT1G68520

MMKSLASAVGGKTARACDSCVKRRARWYCAADDAFLCHACDGSVHSANPLARRHERVRLKSASAGKYRHASPPHQATWHQGFTRKARTPRGGKKSHTMVFHDLVPEMSTEDQAESYEVEEQLIFEVPVMNSMVEEQCFNQSLEKQNEFPMMPLSFKSSDEEDDDNAESCLNGLFPTDMELAQFTADVETLLGGGDREFHSIEELGLGEMLKIEKEEVEEEGVVTREVHDQDEGDETSPFEISFDYEYTHKTTFDEGEEDEKEDVMKNVMEMGVNEMSGGIKEEKKEKALMLRLDYESVISTWGGQGIPWTARVPSEIDLDMVCFPTHTMGESGAEAHHHNHFRGLGLHLGDAGDGGREARVSRYREKRRTRLFSKKIRYEVRKLNAEKRPRMKGRFVKRSSIGVAH

>AT1G49130

MTSHQNIKISEKIMISKYQEDVKQPRACELCLNKHAVWYCASDDAFLCHVCDESVHSANHVATKHERVCLRTNEISNDVRGGTTLTSVWHSGFRRKARTPRSRYEKKPQQKIDDERRREDPRVPEIGGEVMFFIPEANDDDMTSLVPEFEGFTEMGFFLSNHNGTEETTKQFNFEEEADTMEDLYYNGEEEDKTDGAEACPGQYLMSCKKDYDNVITVSEKTEEIEDCYENNARHRLNYENVIAAWDKQESPRDVKNNTSSFQLVPPGIEEKRVRSEREARVWRYRDKRKNRLFEKKIRYEVRKVNADKRPRMKGRFVRRSLAIDS

>AT1G25440

MMKSLANAVGAKTARACDSCVKRRARWYCAADDAFLCQSCDSLVHSANPLARRHERVRLKTASPAVVKHSNHSSASPPHEVATWHHGFTRKARTPRGSGKKNNSSIFHDLVPDISIEDQTDNYELEEQLICQVPVLDPLVSEQFLNDVVEPKIEFPMIRSGLMIEEEEDNAESCLNGFFPTDMELEEFAADVETLLGRGLDTESYAMEELGLSNSEMFKIEKDEIEEEVEEIKAMSMDIFDDDRKDVDGTVPFELSFDYESSHKTSEEEVMKNVESSGECVVKVKEEEHKNVLMLRLNYDSVISTWGGQGPPWSSGEPPERDMDISGWPAFSMVENGGESTHQKQYVGGCLPSSGFGDGGREARVSRYREKRRTRLFSKKIRYEVRKLNAEKRPRMKGRFVKRASLAAAASPLGVNY

>AT1G73870

MVVDVESRTASVTGEKMAARGCDACMKRSRASWYCPADDAFLCQSCDASIHSANHLAKRHERVRLQSSSPTETADKTTSVWYEGFRRKARTPRSKSCAFEKLLQIESNDPLVPELGGDEDDGFFSFSSVEETEESLNCCVPVFDPFSDMLIDDINGFCLVPDEVNNTTTNGELGEVEKAIMDDEGFMGFVPLDMDLEDLTMDVESLLEEEQLCLGFKEPNDVGVIKEENKVGFEINCKDLKRVKDEDEEEEEAKCENGGSKDSDREASNDKDRKTSLFLRLDYGAVISAWDNHGSPWKTGIKPECMLGGNTCLPHVVGGYEKLMSSDGSVTRQQGRDGGGSDGEREARVLRYKEKRRTRLFSKKIRYEVRKLNAEQRPRIKGRFVKRTSLLT

>AT1G78600

MKIQCNVCEAAEATVLCCADEAALCWACDEKIHAANKLAGKHQRVPLSASASSIPKCDICQEASGFFFCLQDRALLCRKCDVAIHTVNPHVSAHQRFLLTGIKVGLESIDTGPSTKSSPTNDDKTMETKPFVQSIPEPQKMAFDHHHHQQQQEQQEGVIPGTKVNDQTSTKLPLVSSGSTTGSIPQWQIEEIFGLTDFDQSYEYMENNGSSKADTSRRGDSDSSSMMRSAEEDGEDNNNCLGGETSWAVPQIQSPPTASGLNWPKHFHHHSVFVPDITSSTPYTGSSPNQRVGKRRRRF

>AT1G75540

MKIRCDVCDKEEASVFCTADEASLCGGCDHQVHHANKLASKHLRFSLLYPSSSNTSSPLCDICQDKKALLFCQQDRAILCKDCDSSIHAANEHTKKHDRFLLTGVKLSATSSVYKPTSKSSSSSSSNQDFSVPGSSISNPPPLKKPLSAPPQSNKIQPFSKINGGDASVNQWGSTSTISEYLMDTLPGWHVEDFLDSSLPTYGFSKSGDDDGVLPYMEPEDDNNTKRNNNNNNNNNNNTVSLPSKNLGIWVPQIPQTLPSSYPNQYFSQDNNIQFGMYNKETSPEVVSFAPIQNMKQQGQNNKRWYDDGGFTVPQITPPPLSSNKKFRSFW

>AT1G06040

MKIQCDVCEKAPATVICCADEAALCPQCDIEIHAANKLASKHQRLHLNSLSTKFPRCDICQEKAAFIFCVEDRALLCRDCDESIHVANSRSANHQRFLATGIKVALTSTICSKEIEKNQPEPSNNQQKANQIPAKSTSQQQQQPSSATPLPWAVDDFFHFSDIESTDKKGQLDLGAGELDWFSDMGFFGDQINDKALPAAEVPELSVSHLGHVHSYKPMKSNVSHKKPRFETRYDDDDEEHFIVPDLG

>AT2G31380

MKIQCDVCEKAPATLICCADEAALCAKCDVEVHAANKLASKHQRLFLDSLSTKFPPCDICLEKAAFIFCVEDRALLCRDCDEATHAPNTRSANHQRFLATGIRVALSSTSCNQEVEKNHFDPSNQQSLSKPPTQQPAAPSPLWATDEFFSYSDLDCSNKEKEQLDLGELDWLAEMGLFGDQPDQEALPVAEVPELSFSHLAHAHSYNRPMKSNVPNKKQRLEYRYDDEEEHFLVPDLG

>AT2G21320

MRILCDACESAAAIVFCAADEAALCCSCDEKVHKCNKLASRHLRVGLADPSNAPSCDICENAPAFFYCEIDGSSLCLQCDMVVHVGGKRTHRRFLLLRQRIEFPGDKPNHADQLGLRCQKASSGRGQESNGNGDHDHNMIDLNSNPQRVHEPGSHNQEEGIDVNNANNHEHE

>AT4G39070

MKIWCAVCDKEEASVFCCADEAALCNGCDRHVHFANKLAGKHLRFSLTSPTFKDAPLCDICGERRALLFCQEDRAILCRECDIPIHQANEHTKKHNRFLLTGVKISASPSAYPRASNSNSAAAFGRAKTRPKSVSSEVPSSASNEVFTSSSSTTTSNCYYGIEENYHHVSDSGSGSGCTGSISEYLMETLPGWRVEDLLEHPSCVSYEDNIITNNNNSESYRVYDGSSQFHHQGFWDHKPFS

>AT4G38960

MRILCDACENAAAIIFCAADEAALCRPCDEKALHMRLDISKCSESVKRVQIVETSSLIWWIKMGTFCLQSLHLVVHMCNKLASRHVRVGLAEPSNAPCCDICENAPAFFYCEIDGSSLCLQCDMVVHVGGKRTHGRFLLLRQRIEFPGDKPKENNTRDNLQNQRVSTNGNGEANGKIDDEMIDLNANPQRVHEPSSNNNGIDVNNENNHEPAGLVPVGPFKRESEK

>AT4G10240

MKIQCEVCEKAEAEVLCCSDEAVLCKPCDIKVHEANKLFQRHHRVALQKDAASATTASGAPLCDICQERKGYFFCLEDRAMLCNDCDEAIHTCNSHQRFLLSGVQVSDQSLTENSECSTSFSSETYQIQSKVSLNSQYSSEETEAGNSGEIVHKNPSVILSP

>AT1G60250

MAQVCHTCRHVTAVIHCVTEALNFCLTCDNLRHHNNIHAEHVRYQLCDNCSMYPSILFCYEDGMVLCQSCYSHHYNCATNGHQTQVVFANMNNQHHDHAHMPHVVHHNNNNNHQQQHVGGHQRRAEMFERSCHGDNNCERWMFAMRCELCVASNSNAVVYCPTHNQILCDSCDRMIHSHEDAVPPHSRCKLCVICKRPSRRFLIGGYQFNFPPVHPPAAEGIPVTPPTELPQQDINYDYLDDVDDFSWFGR

>AT1G68190

MKKLLKTDRRSFCHQSLIFLVLLSGGIFYMLCIIIIENMERVCEFCKAYRAVVYCIADTANLCLTCDAKVHSANSLSGRHLRTVLCDSCKNQPCVVRCFDHKMFLCHGCNDKFHGGGSSEHRRRDLRCYTGCPPAKDFAVMWGFRVMDDDDDVSLEQSFRMVKPKVQREGGFILEQILELEKVQLREENGSSSLTERGDPSPLELPKKPEEQLIDLPQTGKELVVDFSHLSSSSTLGDSFWECKSPYNKNNQLWHQNIQDIGVCEDTICSDDDFQIPDIDLTFRNFEEQFGADPEPIADSNNVFFVSSLDKSHEMKTFSSSFNNPIFAPKPASSTISFSSSETDNPYSHSEEVISFCPSLSNNTRQKVITRLKEKKRARVEEKKA

>AT3G07650

MGYMCDFCGEQRSMVYCRSDAACLCLSCDRSVHSANALSKRHSRTLVCERCNAQPATVRCVEERVSLCQNCDWSGHNNSNNNNSSSSSTSPQQHKRQTISCYSGCPSSSELASIWSFCLDLAGQSICEQELGMMNIDDDGPTDKKTCNEDKKDVLVGSSSIPETSSVPQGKSSSAKDVGMCEDDFYGNLGMDEVDMALENYEELFGTAFNPSEELFGHGGIDSLFHKHQTAPEGGNSVQPAGSNDSFMSSKTEPIICFASKPAHSNISFSGVTGESSAGDFQECGASSSIQLSGEPPWYPPTLQDNNACSHSVTRNNAVMRYKEKKKARKFDKRVRYASRKARADVRRRVKGRFVKAGEAYDYDPLTPTRSY

>AT3G21880

MSPSMEPKCDHCATSQALIYCKSDLAKLCLNCDVHVHSANPLSHRHIRSLICEKCFSQPAAIRCLDEKVSYCQGCHWHESNCSELGHRVQSLNPFSGCPSPTDFNRMWSSILEPPVSGLLSPFVGSFPLNDLNNTMFDTAYSMVPHNISYTQNFSDNLSFFSTESKGYPDMVLKLEEGEEDLCEGLNLDDAPLNFDVGDDIIGCSSEVHIEPDHTVPNCLLIDKTNTSSFTGSNFTVDKALEASPPGQQMNINTGLQLPLSPVLFGQIHPSLNITGENNAADYQDCGMSPGFIMSEAPWETNFEVSCPQARNEAKLRYKEKKLKRSFGKQIRYASRKARADTRKRVKGRFVKAGDSYDYDPSSPTTNN

>AT4G15250

MEARCDFCGTEKALIYCKSDSAKLCLNCDVNVHSANPLSQRHTRSLLCEKCSLQPTAVHCMNENVSLCQGCQWTASNCTGLGHRLQSLNPYSDCPSPSDFGKIWSSTLEPSVTSLVSPFSDTLLQELDDWNGSSTSVVTQTQNLKDYSSFFPMESNLPKVIEEECSGLDLCEGINLDDAPLNFNASNDIIGCSSLDNTKCYEYEDSFKEENNIGLPSLLLPTLSGNVVPNMSLSMSNLTGESNATDYQDCGISPGFLIGDSPWESNVEVSFNPKLRDEAKKRYKQKKSKRMFGKQIRYASRKARADTRKRVKGRFVKSGETFEYDPSLVM

>AT5G48250

MGYMCDFCGEQRSMVYCRSDAACLCLSCDRNVHSANALSKRHSRTLVCERCNAQPASVRCSDERVSLCQNCDWSGHDGKNSTTTSHHKRQTINCYSGCPSSAELSSIWSFCMDLNISSAEESACEQGMGLMTIDEDGTGEKSGVQKINVEQPETSSAAQGMDHSSVPENSSMAKELGVCEDDFNGNLISDEVDLALENYEELFGSAFNSSRYLFEHGGIGSLFEKDEAHEGSMQQPALSNNASADSFMTCRTEPIICYSSKPAHSNISFSGITGESNAGDFQDCGASSMKQLSREPQPWCHPTAQDIIASSHATTRNNAVMRYKEKKKARKFDKRVRYVSRKERADVRRRVKGRFVKSGEAYDYDPMSPTRSY

>AT3G21150

MVSFCELCGAEADLHCAADSAFLCRSCDAKFHASNFLFARHFRRVICPNCKSLTQNFVSGPLLPWPPRTTCCSESSSSSCCSSLDCVSSSELSSTTRDVNRARGRENRVNAKAVAVTVADGIFVNWCGKLGLNRDLTNAVVSYASLALAVETRPRATKRVFLAAAFWFGVKNTTTWQNLKKVEDVTGVSAGMIRAVESKLARAMTQQLRRWRVDSEEGWAENDNV

>AT3G21890

MCRGLNNEESRRSDGGGCRSLCTRPSVPVRCELCDGDASVFCEADSAFLCRKCDRWVHGANFLAWRHVRRVLCTSCQKLTRRCLVGDHDFHVVLPSVTTVGETTVENRSEQDNHEVPFVFL

>AT4G15248

MCRGFEKEEERRSDNGGCQRLCTESHKAPVSCELCGENATVYCEADAAFLCRKCDRWVHSANFLARRHLRRVICTTCRKLTRRCLVGDNFNVVLPEIRMIARIEEHSSDHKIPFVFL

>AT4G27310

MGKKCDLCNGVARMYCESDQASLCWDCDGKVHGANFLVAKHTRCLLCSACQSLTPWKATGLRLGPTFSVCESCVALKNAGGGRGNRVLSENRGQEEVNSFESEEDRIREDHGDGDDAESYDDDEEEDEDEEYSDDEDEDDDEDGDDEEAENQVVPWSAAAQVPPVMSSSSSDGGSGGSVTKRTRARENSDLLCSDDEIGSSSAQGSNYSRPLKRSAFKSTVVV

>AT5G54470

MGKKKCELCCGVARMYCESDQASLCWDCDGKVHGANFLVAKHMRCLLCSACQSHTPWKASGLNLGPTVSICESCLARKKNNNSSLAGRDQNLNQEEEIIGCNDGAESYDEESDEDEEEEEVENQVVPAAVEQELPVVSSSSSVSSGEGDQVVKRTRLDLDLNLSDEENQSRPLKRLSRDEGLSRSTVVMNSSIVKLHGGRRKAEGCDTSSSSSFY

>LOC_Os02g39710

MEAVEDKAMVGVGGAVAAGYSSSSWGLGTRACDSCGGEAARLYCRADGAFLCARCDARAHGAGSRHARVWLCEVCEHAPAAVTCRADAAALCAACDADIHSANPLARRHERLPVAPFFGPLADAPQPFPFSQAAADAAAAREEDADDDRSNEAEAASWLLPEPDDNSHEDSAAAADAFFADTGAYLGVDLDFARSMDGIKAIGVPVAPPELDLTAGSLFYPEHSMAHSLSSSEVAIVPDALSAGSAAPPMVVVVASKGKEREARLMRYREKRKNRRFDKTIRYASRKAYAETRPRIKGRFAKRTADADDDDEAPCSPAFSALAASDGVVPSF

>LOC_Os02g08150

MEMELGLGRYWGVGRRRCGACAVAPAAVHCRTCDGDGGGGGYLCAGCDAEHGRAGHERVWVCEVCELAPAAVTCKADAAALCAACDSDIHDANPLARRHERVPVHPIGSSAAPPPDALLLGGENDAAAAVDGGGGGKEVKLDFLFADFMDPYLGGSPELARFPHADSVVPNHNGSAGPAMELGFAGGGGAAVKPSYSSYTAASLGNSGSSSEVGLVPDAICGGGGGGIIELDFAQSKAAYLPYASTPSHSMSSSMDMGVAAPEMSDCAAAAAGRAYAAEGRAARLMRYREKRKNRRFEKTIRYASRKAYAETRPRVKGRFAKRADDHDAAAPPPQIMLDFAGYGVVPTF

>LOC_Os04g42020

MEGDDKSAVVGGAYWGLAARACDACGGEAARLFCRADAAFLCAGCDARAHGPGSRHARVWLCEVCEHAPAAVTCRADAAALCAACDADIHSANPLARRHERLPVAPFFGALADAPKPGSGAHGGDAAAADDDGSNDAEAASWLLPEPDHGQKDGAVGATDELYADSDPYLDLDFARSMDDIKAIGVQNGPPELDITGGKLFYSDHSMNHSVSSSEAAVVPDAAAGGGAPMPVVSRGREREARLMRYREKRKSRRFEKTIRYASRKAYAETRPRIKGRFAKRTKGGAGADADADADADGEDEEMYSSAAAAVAALMAPGGSDADYGVDGVVPTF

>LOC_Os06g44450

MMELRKYWGVGGRRCGACEASPAAVHCRGCGGVYLCTACDARPGHARAAHERVWVCEVCEVAPAAVTCKADAAVLCAACDADIHDANPLARRHARVPVAPIGSAAAAAVAAEAMLFGVAAAGAEAEAVEDKAAAEHHHHQQRQQHGALNLNVEAKDMKLDYLFSDLDPYLNVEFARFPHADSVVPNGAGAGAAIELDFTCGLGVGVGGAKQSYSSYTATDLAHSGSSSEVGVVPEAMCGGGGAIDLDFTRPKPQPYMPYTATPPPSHSVVSAQMSSSVVDVGVVPERAAAMGEGREARLMRYREKRKNRRFEKTIRYASRKAYAETRPRIKGRFAKRADHDADDADADADDPAAVPSSYMLDFGYGVVPSF

>LOC_Os06g16370annotation

MNYNFGGNVFDQEVGVGGEGGGGGEGSGCPWARPCDGCRAAPSVVYCRADAAYLCASCDARVHAANRVASRHERVRVCEACERAPAALACRADAAALCVACDVQVHSANPLPAITIPATSVLAEAVVATATVLGDKDEEVDSWLLLSKDSDNNNNNNNNNDNDNNDNNNSNSSNNGMYFGEVDEYFDLVGYNSYYDNRIENNQDRQYGMHEQQEQQQQQQEMQKEFAEKEGSECVVPSQITMLSEQQHSGYGVVGADQAASMTAGVSAYTDSISNSLVVTTCHVVNVLEFILVLLSGKTFSKQKLLDHLPIPSQISFSSMEAGIVPDSTVIDMPNSRILTPAGAINLFSGPSLQMSLHFSSMDREARVLRYREKKKARKFEKTIRYETRKAYAEARPRIKGRFAKRSDVQIEVDQMFSTAALSDGSYGTVPWF

>LOC_Os09g06464

MLKLEPEFPGLPQRCDSCRSAPCAFYCLADSAALCATCDADVHSVNPLARRHRRVPMGVVAAPGAGGAFVVRPAGGVNSSWPIREGRRCDYDDDDADAAGEEDEEATSWLLFDPLKDSSDQGLPPFGDALVADFLNLGGGAGEKEDASSSKDCSSSHGKSSEGSHEFAVPGEPVPERQGFGAVSMDITDYDASNFRRGYSFGASLGHSVSMSSLENMSTVPDCGVPDITTSYLRSSKSTIDLFTAAAGSPVAAHSIMSPPQFMGAIDREARVHRYREKRKTRRFEKTIRYASRKAYAETRPRIKGRFAKRSDTDLEVDQYFSTTADSSCGVVPTF

>LOC_Os03g22770annotation

MGQDEVEVGAEKKDQELPEVEVVEEEEEEGSKKAAAGCDYCGDAAAVVYCRADAARLCLPCDRHVHGANGVCSRHARAPLCAACAAAGAVFRRGAGGFLCSNCDFSRHRHGGERDPAAPLHDRSTVHPYTGCPSALDLAALLGISYSDKAAAATAAAGGDDGGWWAIWEEPQVLSLEDLIVPTTSCHGFEPLLTPSSPKIQNSPDGKVNEEVIRQLTELANSDGGGAQIWAHREAAQAGDHQLPSWGTTTQHNTGHGNFGTANSNEVATMPTPGYESCSDRGNCIHTEIGHEKRRVEFTYEQPPASSAEACISSFVQMSELCPSMSNGSSMEETHQTNPGNGTPMQVLPKMPEFVPCPDRNLVISRYKEKRKTRRFDRQVRYESRKARADSRLRIKGRFAKVNQI

>LOC_Os06g01340

MENEVGCECQLCGGRRGVVFCGAHGGRLCLQCDRALHQAHGGAGDHPRAPLCDSCNAAAAELRLNDGATLCGPCAYPYAYAYPYTYTYVYTGCPTPLEMMRLLHAAPPPPPATCSLQQRGEGEELLPTLLSATATPNTATAAPMAMPPPPLQHHTTTSLIMMIRNIHKREERNRAKLRFSKQIKYACRKAGADARKRVKGRFAKASSSSSSSSSSSSSIDHRL

>LOC_Os07g47140

MARDDDPAKKLAVDGGVAAAARCCDFCGGLPAVVYCRADSARLCLPCDRHVHAANTVSTRHARAPLCSACRAAPAAAFHRGDGFLCSSCDFDERLRRGSIGGGGDELPLDDRAAVEGYTGCPSIGELAAILGVVGGDSDKPADDGWWSASWEEEAPQVLSLDDIIVPTTSCHGLRPLLTPPSPENQSSPDNGELDGEVVRQLGELARSEAAAQATFVAGDQLASWASPEFTSGHGDFGIEAASTTVPSCENETWIMSTDCTDPTDASKTDIAREEAPASSSAEPCLSSLVEISEICRSMSYSGSGIDNGGHDPSTLAIMPTQALPKKGVYDIAYPDRGTVISRYKEKRKNRRFDKQIRYESRKARADGRLRIKGRFAKSN

>LOC_Os08g42440

MKDGGGGGGRGQQQQWPCDYCGEAAAALHCRADAARLCVACDRHVHAANALSRKHVRAPLCAACAARPAAARVASASAPAFLCADCDTGCGGDDGAALRVPVEGFSGCPAAAELAASWGLDLPGGCGGEEEEADDAFFSALDYSMLAVDPVLRDLYVPCDPPEVVVAGGGRRLKGEALGHQLAEMARREAETAHPHTQPHSDLSPRTPRRTSAAASGRLQEKQAPPPLPHAAATAAPLPYTSLLMMAPANCTELMENNRVGDEDENVLWESTAPSVPPTQIWDFNLGKSRDHNENSALEVGFGSNNGGFMIKSYNDMLKEISSGTTKDLEDIYDSRYFAAAEDIMSTNVCQLSSKNPSTRSNKRKASSCASTIDGPTTSTSHVPAASGALGGSSNDRGSALPKEISFCDQTVVPTGADQRPCTIKIDSETLAQNRDSAMQRYREKKKNRRYEKHIRYESRKLRADTRKRVKGRFVKSNGAPDDVSNGG

>LOC_Os09g33550

MTWRSCDYCGEAAAALHCRADAARLCVACDRHVHGANALSRRHVRAPLCARCEARPAAARVAAVAGAGGCGGGGEARFLCAGCADDDGAEAARVPVVGFSGCPGAAELAASWGLDLGGGGGRDEFEEDPFFPEAGYPMLAADRVLRDMYVPCDPPPEVAAGGRGRRLKGDSLCHQLAELARREMESAPAQANSGSISPSARRGSAAAIRHEAAAAAAAQRATLPYKSTPVTEAAGCGDVGNGEQFTDDNELVWQRTAPSDPPCQIWDFNLGKSRDHDEHSALELHFGPKDGGFMIKSYNDMIEEVSSSSRKDLQYIYDSTYSFATEDIVSANIYQLTPKQLSTATSGNRRHKNEQHGLTNDGPSSSRIVDVDRTLNSSPEEVAAVLAGENCITDQTVTGADQRNSLKIDSKTIAMNRDNAMQRYREKRKTRRYDKHIRYESRKMRADTRTRVKGRFVRATDIFNVGGGDGG

>LOC_Os02g49880

MSCSSEKAAGAVGGKAARACDSCLRRRARWYCAADDAFLCQGCDTSVHSANPLARRHERLRLRVSSPPPLTARASVEEEAAAAVGTTTTTTSKREGGVTPAWSKRKARTRRPQVKSVGQLLSRRLVVPEMAVESSDERKADEDGAHEELEGQLLYRVPVFDPSLAEFCSPPPIDDAAAASSSCFKEDAADGAVEDAKYPAAAASSPVQQLPDSFVNFEPTDAELREFAADMEALLGQGLDDSNELQDSFYMETLGLITPPVEESGRVKMELDGGVASNSRVSLPSCRAHPKPEDVESADVLDIDFNCTSPDEQKSSASNGAAADSQFFHRSLDLRLNYEAIIESWGNSPWTDGRPPHGQLDDFWPNDHHYSGLWAAGGGGHGAEVGMMTVRPRMDGPGREARVTRYREKRRTRLFSKKIRYEVRKLNAEKRPRMKGRFVKRPSAAAAPCAVT

>LOC_Os03g50310

MASAAAATGAALGARTARACDGCMRRRARWHCPADDAFLCQACDASVHSANPLARRHHRVRLPSASSSPASSPRSAAAPRAGSDDPDAPAWLHGLKRRPRTPRTKPGGGGKHDASAATVAAAAASAVPDLEAEESGIVGDTDHDVGEEDDEDLLYRVPVFDPMLAELYNPVAADDEEQQIEQKPAARVVPFSEPSPEFASGSVEADGLSGFDVPDMELASFAADMESLLMGVDEGFDDLGFLDDEKPHVKLDLDMDMDFASISPAPAPEREERKRKRPEMILKLDYEGVIDSWARDGASPWFHGERPRFDPSESWPDFPAGSRGGLGAAVTAVTGGEREARVSRYREKRRTRLFAKKIRYEVRKLNAEKRPRMKGRFVKRAAALPPLPLPRHQHPPPPPPRALPPVPMMLAPRGAHGRYRF

>LOC_Os06g15330

MSSTAKAAAAGAVGAKSARACDGCLRRRARWYCAADDAFLCQGCDTSVHSANPLARRHERLRLRPSSPPPLVPPSGSGRRDEAVPAAWFKRKARTPRSHAAKSAAAFGQLLSRRLVVVPEAAAGSGGDSPEERKDEGEIVEEQEQLLYRVPIFDPALSEFCSPPPLEDAAAAVSCCNEDGAVENPTKPSMTTTTATTPPLQFFPDGQANFGPTDAELREFAADMEALLGRGLDDGNDEDSFCMETLGLIEPVDDDAGRVKVEADGDAGMTLAWCHELDTETSSGEMLDIDFDCGSPQAATTPDEKVGSSGPAAADDDAQLQQSNLALSLNYEAIIESWGTSPWTDGERPHVKLDDSWPRDYSGVWMAAAGVFGHGGEEQALTPRLGMDGGREARVSRYREKRRTRLFSKKIRYEVRKLNAEKRPRMKGRFVKRAAAAATAAVATACVA

>LOC_Os01g10580

MKVLCSACEAAEARVLCCADDAALCARCDLHVHAANRLAGKHHRLPLLSSSSSSSSPSPPTCDICQDAHAYFFCVEDRALLCRACDVAVHTANALVSAHRRFLLTGVHVGLDAAADDDDKHPPHPLSSSLPRNTAPPPQPPPKRSPSPIYSDDDVIDWATGGHDIGITGNLPDWSLVDEQFNTPALPPVVTKTPPKRASRGPVTAGTAAAVFGNLAGGSPDWPLNEFFGFADFSSGFGFAENGTSKADSGKIGSMDGSPNGGRSSSSSSSSSAAAAGGGGGGQDFFGQVPEVHWAVPELPSPPTASGLHWQRDPRYGGGATDASAVFVPDISSPENPFRCFAAAAAGDHTMKRRRRC

>LOC_Os02g43170

MKVQCDVCAAEAASVFCCADEAALCDACDHRVHRANKLAGKHRRFSLLNPSASGRSPTSTTAPLCDICQEKRGFLFCKEDRAILCRECDVPVHTASELTMRHSRYLLTGVRLSSEPAASPAPPSEEENSSSFCCSADDAVPAPAAPATSHGGSSGSSSISEYLTTLPGWHVEDFLVDDATAEAAAAAAATSSGISANGPCQVGATDRP

>LOC_Os02g39360

MKIQCDACESAAAAVVCCADEAALCAACDVEVHAANKLAGKHQRLPLEALSARLPRCDVCQEKAAFIFCVEDRALFCRDCDEPIHVPGTLSGNHQRYLATGIRVGFASASPCDGGSDAHDSDHHAPPMGSSEHHHHHQQPAPTVAVDTPSPQFLPQGWAVDELLQFSDYETGDKLQKESSPPLGFQELEWFADIDLFHNQAPKGGAAAGRTTAEVPELFASQAANDVAYYRPPTRTAAAAFTAATGFRQSKKARVELPDDEEDYLIVPDLG

>LOC_Os04g45690

MKVQCDVCAAEAASVFCCADEAALCDACDRRVHSANKLAGKHRRFSLLQPLASSSSAQKPPLCDICQEKRGFLFCKEDRAILCRECDVTVHTTSELTRRHGRFLLTGVRLSSAPMDSPAPSEEEEEEAGEDYSCSPSSVAGTAAGSASDGSSISEYLTKTLPGWHVEDFLVDEATAASSSSDGLFQGGLLAQIGGVPDGYAAWAGREQLHSGVAVAADERASRERWVPQMNAEWGAGSKRPRASPPCLYW

>LOC_Os04g41560

MRIQCDACEAAAATVVCCADEAALCARCDVEIHAANKLASKHQRLPLDAALPAALPRCDVCQEKAAFIFCVEDRALFCRDCDEPIHVPGTLSGNHQRYLTTGIRVGFSSVCSANADHLPPPAPKGNSKPPASGIAAAAAPKPAVSAAAQEVPSSPFLPPSGWAVEDLLQLSDYESSDKKGSPIGFKDLEWLDDIDLFHVQSPAKGGSTAAEVPELFASPQPASNMGLYKASGARQSKKPRVEIPDDDEDFFIVPDLG

>LOC_Os05g11510

MSPPPPPYYHHLLLLRSSPTTTGGGARVLAAAELARMKLLCSACEAAEASVLCCADEAALCARCDRDIHAANRLAGKHLRLPLLSPASSSSSSAAALAPPPPSPPKCDICQESHAYFFCLEDRALLCRSCDVAVHTANAFVSAHRRFLLTGVQVGQEQDEHSPDPPEPSPPPPPPPPASKSDHPAPLYGEGGGGFSWDAADSPAAGGLPDWSAVVDQFGSPPPPRHTDTATVTTPPPTKRSPRAPAFGGQGGMMDWPLGEFFGGFTDFTGGFGFGFGDSGTSKADSGKLGGSTDGSPYYRSSSEDDRNADELFGQVPEIQWSVPELPSPPTASGLHWQRHPAATHGGGGGGPDTTAFVPDICSPDSCFPATTSKRRRQ

>LOC_Os06g49880

MRVQCDVCAAEPAAVLCCADEAALCSACDRRVHRANRLASKHRRLPLVHPSSSSSGDGGAAAAPLCDVCREKRGLVFCVEDRAILCADCDEPIHSANDLTAKHTRFLLVGAKLSPAALAEQPLPSSDCSSDDDAAAAATEEEYHSSAASTGAAVSAPLDASSNGAGGGGGVGGSSISDYLTTICPGWRVEDLLPDDDAFAAAAAQAGKEKDERVPFLDADLFDVVAGRPEKKGGAWAPHVPHLPAWCLDEVPVVVAASAAPAATPVKAKQGHVRDSHWSDSDAFAVPEFSPPPPPAKRARPSSQFWCF

>LOC_Os06g05890

MKIQCNACGAAEARVLCCADEAALCTACDEEVHAANKLAGKHQRVPLLSDDGGAAPAAAAPAVPKCDICQEASGYFFCLEDRALLCRDCDVSIHTVNSFVSVHQRFLLTGVQVGLDPADPVPPVADKHVKSAGGSVDSATKHLQRNPTDLSGENSASLPSQNVINGNYSRQSSVTMAKTGQVNWTMSNNTIRSIDPPPKYSSEESPALLLASHTSTMAAYSSQISKDSDRIYNLPFTGGNGSDSLHDWHVDEFFSNSEFGFAEHGSSKGDNAKPGSAGGSPQCRLAEGLFVEGLLGQVPDNPWTVPEVPSPPTASGLYWQNNLLCPSYDSTMFVPEISSLENSQNNFTVSAGLKRRRRQF

>LOC_Os09g35880

MRTICDVCESAPAVLFCVADEAALCRSCDEKVHMCNKLARRHVRVGLADPNKVQRCDICENAPAFFYCEIDGTSLCLSCDMTVHVGGKRTHGRYLLLRQRVEFPGDKPGHMDDVAMQQKDPENRTDQKKAPHSVTKEQMANHHNVSDDPASDGNCDDQGNIDSKMIDLNMRPVRTHGQGSNSQTQGVDVSVNNHDSPGVVPTCNFEREANK

>LOC_Os12g10660

MKIGCDACEQAEAAVLCCADEAALCRRCDAAVHSANRLAGKHTRVALLLPSSSSAAAGDDDHHPTCDICQEKTGYFFCLEDRALLCRSCDVAVHTATAHAAAHRRFLITGVRIGGSVDAAAAADVIVSPTSSSIAPAGSASSNHAGAAGNNNGRSPAPVRFSGGDGGVEPEQQWPWSDVFAADDDDDVSAAMEQCYYHGISEPHSSSLTG

>LOC_Os02g49230

MDALCDFCREQRSMVYCRSDAASLCLSCDRNVHSANALSRRHTRTLLCDRCVGQPAAVRCLEENTSLCQNCDWNGHGAASSAAGHKRQTINCYSGCPSSAELSRIWSFSMDIPTVAAEPNCEEGINMMSINDNDVNNHCGAPEDGRLLDIASTALMSDLPTGDKFKPLIGSSSGDGMNLLPLNSDQPAEPVSTTPKAPCVTDKDMFNDGSVYGDFCVDDADLTFENYEELFGTSHVQTEQLFDDAGIDSYFEMKDVPADESNEQPKPVQPECSNVASVDSGMSNPAARADSSHCIPGRQAISNISLSFSGLTGESSAGYFQDCGVSSMILMGEPPWHPPGPESSSAGGSRDNALTRYKEKKKRRKFDKKIRYASRKARADVRKRVKGRFVKAGEAYDYDPLSQTRSY

>LOC_Os06g19444

MGALCDFCGEQRSMVYCRSDAASLCLSCDRNVHSANALSRRHTRTLLCDRCASQPAMVRCLVENASLCQNCDWNGHSAGSSAAGHKRQTINCYSGCPSSSELSKIWTFVSDIPNVAPEPNCEQGISMMSISDSGVSNQDNAAGDSSLLDIASATLMSDLGTAGKPKSLIGSSSEAGVNLLPLATDQMAGSVDSTSAKVPYTADQDMFSKDSIYEDFCVDDVDLSFENYEELFGTSHIQTEQLFDDAGIDSYFESKEIPSGNSDEPKLMQPVTSNAVSADSGMSIPGAKGDSSLCIPVRQARSSISLSFSGLTGESSAGDYQDCGVSPVLLMGEPPWHPPGPEGSFAGATRDDAITRYKEKKKRRKFDKKIRYASRKARADVRKRVKGRFVKAGEAYDYDPLCETRSY

>LOC_Os02g07930

MEVGNGKCGGGGAGCELCGGVAAVHCAADSAFLCLVCDDKVHGANFLASRHRRRRLGVEVVDEEDDARSTASSSCVSTADSASSTAAAAAAVESEDVRRRGRRGRRAPRAEAVLEGWAKRMGLSSGAARRRAAAAGAALRAVGRGVAASRVPIRVAMAAALWSEVASSSSRRRRRPGAGQAALLRRLEASAHVPARLLLTVASWMARASTPPAAEEGWAECS

>LOC_Os06g45040

MGGEAERCALCGAAAAVHCEADAAFLCAACDAKVHGANFLASRHHRRRVAAGAVVVVEVEEEEGYESGASAASSTSCVSTADSDVAASAAARRGRRRRPRAAARPRAEVVLEGWGKRMGLAAGAARRRAAAAGRALRACGGDVAAARVPLRVAMAAALWWEVAAHRVSGVSGAGHADALRRLEACAHVPARLLTAVASSMARARARRRAAADNEEGWDECSCSEAPNALGGPHSFRTTEAPAFSFIASVRWKKRLLIVLAEYKYATSNQFPTLN

>LOC_Os08g08120

MSVAAEGKEKGVGGGGGGAGAGACELCGAAARVYCGADEATLCWGCDAQVHGANFLVARHARALLCRGCARPTPWRAAGPRLGPTASLCERCVRRGGGGRGGGGGGGAAGGGGRGGGGDEEMGGEGDEEEEDEDEEVVVEEEEDEDDEDEEGEGEGENQVVPWAEEAEATPPPVASSTSSSSREAAANGANAADRVKEDQPCSTSQPSLCRYASSAHHGGGGRSDEATSSRNGGGVGGRFLASRHRKRSPSDFRRSGLAQSVSGVQGRNCSNAVVGRNDFS

>LOC_Os08g15050.1new

MMASDGSASPASCGGAACGVCGGAATVYCAADAAALCVPCDAAVHAANPLASRHDRVPLAVAMAAASSGVYDHLFAPDDDAASSWAAAAAAGAAVQGQGQGSPNDSSSSFTNDSAGGGGGGGAERSLFDLLSDVDIMSCGGGGLASSFDGAAAPPLWLHPGQLAALTPWSPADSVVVPTSAAGAVAAAAAAREERVRRYREKRKNRKFQKTIRYASRKAYAEARPRIKGRFVKRATTAAASSSSDDDSTAAAGVSGAGGAGAAATKEAKFWLSFSDDGRADGVGFYMDSTTAATAAYGVVPTF

>ATR0580G367

MSTANVSATTKPDSEPRGSGDAFSAPFAAQFPGHEAPPSAPWSVAKACEGCKATAALLFCRADSAFLCLGCDARVHGANKLASRHERVWVCEVCEQAPASVTCKADAAALCVACDADIHSANPLARRHERFPVVPFYEPPAPKSSASCLLVPDRGPKTTENAKLDDEDEDDDHNSIGADEAEAATWLLPNPKIGGGEMKSADYFFPEVDPYLDLEYGSPVDQRFQAHNGTDSVVPVQSKGMGNDQCLEFDFSRPKNGYSYTTASMSHSVSSSSLDVGVVPDGSAMTDISNPYSRTLKGGQDLVAHVPSQPSSHYPPMDREARVLRYREKRKNRKFEKTIRYASRKAYAETRPRIKGRFAKRTDIEVEVDRIYSSAAALMADTSYGIVPSF

>ATR0691G129

MVKDEDGHPLLLGSGSNAWKTRRACESCRAVPSAIFCRADGAYLCTPCDGRVHSANLLASRHQRVLLCEVCEAAPASLTCKADAAALCASCDADIHAANPLASRHHRLPILPLSLKSGIQDRENGNQEEDERESEEEANSWLNIGGGGGGGGGGGGGGERNTVSNGCFYGDVESYLDIDFGVDGINGFEGGEMGMKDGGVGECIVPVQPSNSNNSDCINNNGSQSHMNGNNNSSSCNNLEFLEMEVCKDGYGYTASLSHSVSSSSLDVGVVPEANITDFSASHLRTSRLGDITTGHPLQMSHQFTPMDREARVLRYREKRKTRKFEKTIRYASRKAYAETRPRIKGRFAKRTDVEVEVDQFYSASLVAETGYGVVPTY

>ATR1133G053

MTSSAESRRAVVAAAAGAKTVRPCDSCGKGRARWYCAADDAFLCHACDGTIHSANSVARRHDRVRLKTSSSHRCEFVLPSWQRGKARSMRPHPNMKNPIDGAKPHANPLVVPQMQEESSPDEVDEQLLYQVPVFEPMAADLRSSPARATPADCKPTIDTPAEMAVAKFSAIPSLDMDLADFAADVESLLGQGLDEDAFGIEGLGLLDNYREEEEGVASRVKVEEEEDEDGIFGCSHHPDVAEVELSSETLEFNFDYDTPTTAGEDDEEKCRIVLEKSISLRLDYEAVISAWGRSPWTTGQRPEFNPDDCWPDPFTGNCVPEMSTYSEVGNATASDGGREARVSRYREKRRTRLFSKKIRYEVRKLNAEKRPRMKGRFVKRTSFPPTPYPF

>ATR0149G002

MRTLCDACEGAPAIFFCAADEAALCRACDNKVHMCNKLASRHVRVRLADPSDVPRCDICENSPAFFYCEIDGSSLCLQCDMAVHVGGKRTHERYLLLRQRVEFIGDKPTPIEDHAWQPMEKVEYKRDQNMQPHLMMNREGPKENNINHQGPRVPTYDSNANNDKHDAKLIDLNARPHRNQGQPSNSSQKSSGMDTANNRDCASLVPVGSFKREVARNSVADAKDVDM

>ATR0602G117

MKIQCNVCEIAEAAVMCCADEAALCLSCDEKVHAANKLAGKHQRVSLMNSSSQMPKCDICQETTGFFFCLEDRALLCRKCDVAIHSTSPYVSAHQRFLLTGVKVGLEPAEPISPVVPNSPSSTKKPNARRSFEISSSGVNKEQFSREIGGNGSLALNRVSFSGGQPAECLPSWPLDEILSLPDISQSYGLVYSGSPKADASNFGESNWAPSSYPADEDIGMEEYLFQVPDVKLKTPSPPTASGIHWQRGLGLRSCYDHEISNNSVFVPDISYPEQDNYPERIALKRRRQI

>ATR0661G082

MKIQCDVCNKAEASVFCCADEAALCSACDQRVHHANKLAGKHRRFSLHHPCSDQGPRCDVCQERRAFLFCQEDRAILCRECDIAIHTANDLTYRHNRFLLTGVKLQSIAADTLENKKAKMPATSSTPQQNSPAPLSSTSSSISEYLTHTIPGWRVDDFLTGAGSLSPPHNNDGLLETDELLKLVESDFNGDCFFPSDDFILQIPKQGKGWGAERFMSSINGGEHHGFTVPQISPPTKRQRLVWQM

>ATR0743G170

MKIQCDVCEKACASVICCADEAALCTECDIEVHEANKLASKHQRLHLHFISNQLPRCDICQEKPAFIFCVEDRALFCRDCDEPIHTTGSLSGNHQRFLATGIRVALASASKDTSVPIVPEPPSRNSQKVAEKELQVHAPSAYASPSWPDDELLQLSDFDANDKRECIGFGGLDWMEDIGLFNEDVPHESFGAAQVPEISNFNNARPPTSKSLVSLKKPRLEIPQQEEDYFTVPDLGDIHT

>ATR1089G082

MKIQCDVCEKAEAAVLCCADEAALCWSCDETVHAANKLASKHQRVPLLLNSSHHANPKCDICQEKSGFFFCLEDRALLCRQCDVSIHTHSPYVSSHQRFLLTGVKVGLNHHTSPTPQDSKNENLPNSALPRDQAAGVRPQWPLDEIFSFPDFDHLDTLSDCGSSQISTRANNHHHQNHPFLHHQQN

>ATR0053G049

MLDPEAKSQNYGFQNQKMEKNCEFCGKSRSLIYCKADAAFLCLACDARVHSANALSHRHVRGLLCGSCSTEQAVLKCMDHKLFFCRTCDCDSHHNVVQHSKRLVNGYIGCPPAKDFAALWGIDLNLCREMGFAHFLKDEEGFRVDVSKGRIAGSSMSSLLDGSGASMAELNEMILGASVADCSMSSSSQNKVVHNNQWHQANYPILQQLFDLEKLQLSESISRPSSLRVQNEVFRPETQTCRQLSQDNDMNQQHSQGEHIDKQQVDNLNLHLQQQMQPFSLITSQPQPLSSSSNAGVSLQGDSFWQCKSPVQSNQVN

>ATR0582G183

METPKKTVDTLCDYCGEEKAVLYCRADSAKLCLSCDQHVHSANALSKKHSRSLLCDNCSSEPVSMRCCSENLVLCQECDWDIHGNCNSVSSHHRQSIEGFSGLPSPLELALLWGFDLSDKPSENGYCRNEPLTLNPMESMVTMDSWICKSPVNLHDLIVPSGNVGLPAYLDVPCVQIPSLSKDQNPTCGKRKQLVFQQLMALLKHDLSNSDKVHGDLESPTNLDPDTPNGNFNSRNLEPPNSRDADLLDGSLQHTPFTSLLMYPHTTNFKDEHLIDEDGMWGCSVPDQANQKCIFD

>ATR0743G164

MSSSIHLINGERRLPSFCLCFEHCLDELDWNLDFERQVNSGFENSKSSPGFGVRTMDSLCDFCGVQNAVVYCRPDAARLCLTCDRNVHSANALSSRHNRVLICEACSSHPATVRCFDDKVSLCQACDWNGFGCSMASGHNRQSLNGYSGCPSPSQIWSCVLDGSSANSHCLMTTINENGESNDCWEGEGNGGLIGLAGQSRIQELDNGNKCDLWMHASSATSGSNPAPFREEQQLEGRNESNESKHGCPQLKDTDFCDGDDLCEVLGISDDALNLENCGELFDGSESLIKDVFEDTTIDCFFMENNFSVADSNGQSDIAIEASSTAQQDCMTLPSMQFVGTPRSIQATTTCSATNQVISNPCSNSNSVNLGFPVRQFFSSISLPLSGLTTGDNTGSDYQDCGVSPSPVMFIGDLEPQCPQARDSAMMRYKEKKKNRKYEKQIRYASRKARADIRKRVKGRFVKAGEVYDYDPLNATTGC

>ATR0751G075

MGPLCDFCGDQRSMVYCRSDAACLCLSCDRKVHSANALSRRHSRTLVCDRCSSQPAIVRCLDESTSLCQNCDWNGHGAPGSASGHKRQTINCYSGCPSATELSRIWTFVMDFPSTEDTTCEQGMGLMTIDENNVSNSWGSMENGDKVDFCVGSSRIPALDSMSQVDQQVVTSDFTPPKMCSTEIKGLGICDDDIYKDFNVDDVDLDFDDYEELFGGSHNPSGYLFDDAGVDSLFMGKNASVANSNAQSENSMEASSVRPDGSNVLTPCISRHVNTIQTTTCSEPIVSGHDRCTGLDLTFPVRQACSSVSLSFSGMTGESNAGDYQDCGISSTLFMGEPPWDLSGPEGTFPQSRDSAVLRYKEKKKTRKFEKKIRYASRKARADIRKRVKGRFVKAGEAYDYDPLCQFQARSF

>ATR1132G033

MRDCELCDLPAMMYCESDQASLCLECDAKVHGANFLVARHSRCLLCQVCQCPTPWRASGSKLGSAVSLCEKCVNSRHKQQSQHEEEEDGEEEEEEEDEEEEEMDDEADNGENVSDGDSEDGENQVVPWSLTPPPPPPPPSRSVPLPSSSSEESSQGRLNMKRLRDNADLHYQEGFGSSSSSPQRGNDQARSSGEDEANSISSCRPLKDRKRESMISVRQRSYASKSDIVQALRRLQQQRSDPNSSSINCRISKDHSPVDVLSTGSESPPV

>Mapoly0110s0028

MPKPCDACQSSSAVVYCRADAAFLCAGCDTKVHGANKLAQRHERVWVCEVCEVAPAVVTCKADAAALCVACDTDIHSANPLARRHERTPVTPFYECPGQSIKVAHISALVAPQGGQGRDDDADLDADEHSVAEEAASWLLPNPKPLTENNILGDDKGSMGQGEMGSLVNDDGDDDDLEPSSFLVPSGGFSPNSTLSHKSDGMSMKHHKSKVEANAGDLFSDVDPYLDLDYATVVGANPNLGADSLVPVHSDHPAHSSPSMSTGSGSFEGEAVCKAGYGYESSLTHSMSSSSLDVGIVPDTNALSDISTPYDGSQGHGFEFPPRMLHMAQVEPPMAREARVMRYKEKRKNRKFEKTIRYASRKAYAESRPRIKGRFAKRNPTDMSLQDMYPNVPDSGFGVVPAF

>Mapoly0049s0067

MTATTEDRSMDPKTATAVMSMAGRAVRACDVCGRDRARWYCAADEAYLCGRCDNSIHCVNALASRHERVRLNPNGTACKGGERTSSKSFPGLQIRKSSKSLKHHVNPKMRMIKDEMLAQSFFSNPDDTSRVPGYPESLPEDESSRLGLPAYVPDLQASVDDLGGDGSASDGFPMDMPQELDSLERGDEDCVGGSDDEYLDFTDIVTEADFLDDKGDEGSVFSLDDIKSEHCHDAVKNEGFGQFQPVCNDGFESINVKVEKDGSLFHQTQQFSEEGQTKCLSLKLNYEDVLSAWSGRGPLWTEDGGQPQTVPILSNIDMALNLDLESYHREQLGEVPVLGMGEEVSEVGVGREARVMRYREKRRTRLFSKQIRYEVRKLNAERRPRMKGRFVKRLPP

>Mapoly0122s0029

MKSKASAIGGMAAAMAIAGRAARPCDVCGRERARWYCAADEAYLCEPCDGSVHNANALASRHERVRLSPNGAPMKIDRHRKDDTTSTTSLADQAPPDRLAISSRKKQFRDNRASQRATAPAGPPATLQATPSAQQDPGSQAFEVKLERSSPEDAQIYFDELPRDPRSPLGSPCRDSKGEPDYAHHFMVFDECEVDDVLTPDECEAEGVAGGLESSAGDFDALGEQEDSEAAGGSGMEKPEPGMFLPLTSTASTVQSSKDEEGSHVPSLRLNYADILNAWSDRGTLWTGGHAQTVPEDSTSDAAANPDHGVVPLLAGEDGNSAREARVNRYREKRRTRLFSKKIRYEVRKLNAERRPRMKGRFVKRSAGEQ

>Mapoly0027s0033

MRTLCDVCEAAPARLFCAADEAALCLTCDEKVHGCNKLASRHVRLELAEARAVPRCDICENAPAFFFCGIDGTSLCLQCDMDVHIGGKKTHERYLLMGQRVELPTKPRQDDLAAVKDGATKDGKKSKDGNEHIHHHHHHHHHHHHHHEGQQDKTKPVTSGEWNNSNAPAVAIAVGDEDLQSPTNRRVSRMIDLNTRPAHLQQLAHTSKDQELASSDDECVGVVPEMAPRSSSENSEE

>Mapoly0064s0079

MRVQCDGCERAVASIMCCADEAALCNECDTRIHAANKLANKHQRVSLMGQTESPRCDICQEKSAFFFCLEDRALLCRDCDVSIHSATALSSAHQRFLVPGTRVALEALSNSSPAPQETATSEELARHPLSSIINGTATNSSKSPSVSYKQSSYYTPTPSFTSPSSGKTSVITSNKSVGMSNTSANHQSVSSTQAKRTSITEFLSDAIPNWRVDELLNIPELGEGYSLGDIASSKADQANLGDYDWTADLSLFDEQVYAESFHEVPQMFSNPAASGMCRTGRATGAMKGKAKQDSLVPEYDDAFVVPDLGLHSAPASPPVAKRRRLLEV

>Mapoly0049s0030

MEKKCELCGDRATLYCAADDANVCWSCDAKVHSANFLVARHARSVLCGRCCQETDCKTSGPHPSPLSRLCPRCNPSTTAAVVNDCNGDSEEMENSETLSYPSATTEAADSGCTCASQESHTGEHLSRQGRRKSAVEMVGRARGSRRFHGAEIKRLWGVERGKTLQGRRIQKRSSSPTGLVIQTLQLESRSQRADRSFGYGSGVGSNSVSPTKEVCQKPVSCDLTEESSCVSYLDEEGSYNEDAV

>Bradi3g05800

MEMQGIGRYWGVGGRRCGSCGGSPATSHCRTCGGGESSYLCAGCDAAHARAGHERVWVCEVCELAPAAVTCRADAAALCASCDADIHDANPLARRHERVPVRPIGSSESEIDDAGDIGHVSHGSRLLGGGGKADAIVGGGKENAMMKMDFLFADVMAEMDPFFGPEFARFPHADSVVPSNNNHGGSGPGAVELDFGRVAAAPVSKPSSYSSYTAASLGGSGSSSEVGLVPDAICGRGGGIIELDFSLSKAAYLPYAPTPTHSVSSTVDVAAVPERGAVDGAASTAATGEMSREARLMRYREKRKNRRFEKTIRYASRKAYAESRPRVKGRFAKRTEDADADADADADADADAAATDMATARAPSCVLDFGSYYGVVPTF

>Bradi3g56260

MFMNCNFNCNLLEQEAGRRNFPWARPCDGCHATPSMVYCRVDAAYLCASCDAQVHSANRVASRHERVRVCEICESAPAVLACRADAAALCTTCDAQVHSANPIAQRHQRVPVLPLSAVAISAASGFAEVRAATIHGDKEEGEEVDSWLLLRRNSDDNNCSNSIDRYFNLVGYNPYYDNATCNPGPGEQYRLQEQQVQNRYREKEGSECVVPSQIVMASEEQESGYRIIGTEQAAFMTVGASTYTASISNSISFSSMEVGIVPDNTRPDISKTNILTTSGAMELSVHSVQMPVHFSSMDREARVLRYKEKKQARKFQKTIRYATRKAYAEARPRVKGRFAKRSDIEHEVNHMLSPPVLPESSYGTVPWF

>Bradi3g48447

MKAEEMPVVVGGAGGGCWGQQQQQQQGARPCDTCGVDAARLYCRTDGAYLCGGCDARAHGHGGAGSRHARVWLCDVCEQAPVAVTCRADAAALCAACDADIHSANPLAGRHERVPVAPFFGALAHEADAAAAHKEEDGSNEEAEAASWLLPEPGDSPEAEDTAAFFADSDAYLGLDLDFVRSMDGINAIGVPVASSELDLAAAGTLFYPDHSMNHSVSSSEVAVVPDAMSLGAAAAVVVSRGKDREARLMRYREKRKNRRFHKTIRYASRKAYAETRPRIKGRFAKRTGTGTADDDALEHDDGPFSPAVSALVASDGDYGIVPSF

>Bradi5g14600

MEGEEKSVAAGAGAYWGVGARACDACAGEAARLFCRADAAFLCTGCDARAHGHGSRHARVWLCEVCEHAPAAVTCKADAAALCAACDADIHAANPLARRHERVPVAPFFGALDVDAPNKHFVGGAGAHAPAAAGINNEEDEDDRSNDAEAASWLLPEPDQKVGGAFFADSDPYNLDLDFARSMDDIKAISVQLNGAQAELGLTGGNNKLFYSDHSMNHSVTSSEAAVVPESAPVAVVSRGREREARLMRYREKRKSRRFEKTIRYASRKAYAETRPRVKGRFAKRTGNGGAAALGEEEDEHEGLYSSAAAAVAALLQAPGAGHGHGPELDYGVDGVVPTLV

>Bradi1g31280

MMELRKYWGVGGRRCGACVGEAAAAAVHCRTCASYLCGVCDAAPEHAGRAHERVWVCEVCEASPAAVTCKADAAVLCAACDADVHRANPLAQRHVRVPISPILGFHGAAMAMRAPELEEEEEEDLALINLNVEAGKGVKLDLLFSDLVDGPYLGGGGVHDFAARFNGHADSCLVPSAGAVVEMDFACGIGAAKPPPPSSYGSYTAAAATNSLGHSGSSSEAGVVPEAPICGAAGSFELDFTRTELQYPAPYNMPMPYTAAPPPPTHCVPAAAAADNMGMVVPAAATGEEREARLTRYREKRKNRRFEKTIRYASRKAYAESRPRVKGRFAKRSSPGADDDSDEINEAAVPPSSYMLDFGYGVVPSF

>Bradi3g56490

MGALCDFCGEQRPTIYCRSDAASLCLSCDRNVHSANALSRRHMRTLLCDRCASQPAAVRCLEENTSLCQNCDWNGHGATSLAAGHKRQPINCYSGCPSSEELSRIWSFAMDTHTAADEPNCEEGISMMSINDSGVNNHYAAEQESSLLDIASTALMSHPPTVEKLKPLNSGDGMNLRPLATHQPAGSVSVTPKVQCITDENMFDDGSIYEDFCVDDADLTFENYEELFGTSHIQTEQLFDDAGIDSYFETKEMPATESKEELKPMQPECSNVVSADSSLCIPSRHAISSISLSFSGLTGESNAGDHQDCGVSPMLLMGEPPWLPPGSEGSFASGSRGSALTRYKEKKKRRKFDKKIRYASRKARADVRKRVKGRFVKAGEAYDYDPLSETRSY

>Bradi1g43220

MGALCDYCGEHRSMVYCRSDAASLCLSCDRNVHSANALSRRHTRTLLCDRCASQPAMVRCLEENTSLCQNCDWNGHSAGSPDAGHKRQNINCYSGCPSSAELSRVWSFILDIPNVAPEPNCEQVISMMSISDSAVSNEDNAPGGNSFLDIASATLSSDHNNDDKLKTVIGSSSEAGVNLLPHATGQTAVSVDSTTAKVPYTPDKHMFSKDTIYEDFSMDDIDLSYENYEELFGNSHIQTEELFDDAGIDSYFEMKEVLAGSSDEQPKPMQPAASNAVSADSGMSNPGVKDDSSLCIPVRQALSFSGFTAESNAGDYQDCGVSSLLLMGEPPWLPPGPDGSFAGIRDSAITRYKEKKKRRKFDHKIRYESRKARADVRKRVKGRFVKAGEAYDYDPLDTRSY

>Bradi1g18407

MASGVSGDAAAAACCAAAACYYCDSAASAVVYCRADAAGLCLPCDRLVHAANTVSSRHARVPLCAACRAAPASVCHPLAAPASAARFLCSGCCSNNFDDDGGAAVEGYTGCPSAGELATILGVVAHDARGHGNEAAAVNGDEEGWWLRVWEKESTVISMDDVIVPTTSCPKGRGQSSPCEDLDGEVLRQLRELARSEAAAAYVEVEVAGADQLPPWGPSGYYAADHGDLGALTFEAASMAVPSCDQLHQAWIAADLPDDVVEAVSAAGGREQSPAAADPILSPLVLEIAAGVCPSMSCSGHAHPSAVSAPARAVAKPKDDDGGHRDDHRTLSAGAPAEEARPPGRSVAFGGYDIAYPDRGTVISRYKEKRKNRRFGKQIRYESRKARADGRLRIKGRFAKSGST

>Bradi1g62420

MGQDERQDREVAPQAQMEQEERQGPETAAHEAIFSCDYCSGARAVVYCRADSARLCLPCDRHVHAANAVCSRHARAPLCAACSAAGAVFRSGATALFLCSNCDFGRNREGEQPPLHDRCTVQAYTGRPSAHDLAALLGVPDFEKPPADQGWWTIWEEPQVFSLQDLIVPTTSCHGFQPLVTPSSPKNQGSPDGKTNEEVIRQLRELAEADGGAVQIAPREEAEQGAHQLPSWEPSEHITGSGNFATENSHEVLATMPTPGYENGGWNNNSNYHALNDVCKNEYEHEQAPVGSAEACLSSFVQMSELCPSMSNGSTRDDSQQANLGIGMSMQTFPKRGGFDVVAGPDRDIVISRYKEKRRTRRFDKQVRYESRKARADSRLRIKGRFAKANQINP

>Bradi3g41500

MPSLDGFGVWIASRASAAAEQKEKKLISLSEALSLSAFDLTSPPSPLVRACVVEMKEGGGRQQWPCDYCGEAAAALHCRADAARLCVACDRHVHAANALSRKHVRAPLCAACAARPAAAARLASGSSDPEFLCSACDDDGACEGAGAARVPVEGFSGCPAASELAASWGLDLLHPLPTDGCGGGGGIGRGEQEDEEDALFFSSLDYSMLVDPEMRDLYVPCDPPDSGGRPLKGEALCQQLAEMARRETQSHPPPPPQQQQYTPDLSPRTPRRSSAGPEKQHQQPPPLPQEPPFPYTSLLMNMMPPDNLAAGNNDRLRDDEAGQQLQWEFTAPSSVPPTQIWDFNLGRSRNHNENSALEVEFGSNNGGFMIKSYNDMLKEISSGTTKDLEDIYDSGYCAAAEDIMSTNICQLSSKNVSTASNKRKVSSCTSTIDGPTTSGNYVPTSGPLGSSSQDRGAALAREISFGEQTIVPTGADRPTTRIDSETLAQNRDSAMQRYREKRKNRRYEKHIRYESRKLRADTRKRVKGRFVKSNEALNASGNGG

>Bradi3g57000

MSSSERKAGGGGGASAAVGGKAARACDGCMRRRARWYCAADDAFLCQSCDTSVHSANPLARRHERLRLRGAMPMPMPVEGGVASETTMATKTATKAKRLQGGVAWAKRKARTRRPPVKSVGQLLLSRRLVVRVPDDHQAGAGGESSDEQRAAAAEDEEEQQLLYRVPVFDPALGEFCSPPPVEDYYAIGNDNDNAFVVGEDTNKQLVGPASSSSPVQELPDCFASFGPTDAELREFAADMEALLGGQDALGGNEQLMAEDSFYMEALGLVSPLSGDHKGVDAGRPVKVETDGGARRSSPPTLVDDDDGSFEHKAATASYGEAADAGFLKRSLDLRLNYEAVIEGWGSSPWTDGRRPHGGQLDDLLLHDHYYSGMWTAAAGGGRAARPAADDGGWREARVSRYREKRRTRLFAKKIRYEVRKLNAEKRPRMKGRFVKRPAGAAPCAVVA

>Bradi1g11310

MTNAGAALGARTARACDGCMRRRARWHCAADDAYLCQACDASVHSANPLARRHHRVRLSSSASSSSSPAAASQQADPDAPAWLHGLKRRPRTPRRKPGGINGSKQQQHDALGTKAAVAASAPVPDLEAEDESLSGIIMGAGNEVDQVDEDDDLLYRVPVFDPMLAELYSHTMPDDDQALEQKPCFAPLLATDPSSDQYGGVSGLADGTDGFSGFDLVPDMELASFAADMESLLMGVDSGYDDLGFLDDEKPQMNHLGFDDMQDDFDQSTVAPPAPQEEEQQEDRKRKRPDQMILKLDYEGVISSWTHDGASPWFYGERPHLDSSDSWLDFPAGSGRGFGLGAAVTAVTGGEREARVSRYREKRRTRLFAKKIRYEVRKLNAEKRPRMKGRFVKRAALPPLPPRATPMMLAAPHGGSAHGRYRL

>Bradi1g43990

MSSSLAKAAAGAMGSKAARACDGCLRRRARWYCAADDAFLCQACDASVHSANPLARRHERVRLRPTSPLQAGAGGPRARRGDEVVPAWFKRKPRTPRGKSVIGHLLSSRRPVVVPDEASGGEGSPEEHKLFEGETDEEELLYSVPVFDPALAELCSPSSPRHLEDYASCCNEDRVVVLESPTEPAAPSLAVDQFFPDSVVSFGPTDAELMEFAADMEALLGRGLDEGTEEEPFCMEALGLIQPMVDVDGAGGRVKMETDGEARGMFLACGLEREPETEVSGNMLGIDFDDDYGSPQATPDENAASGSAARFLPRSLSLNLNYEAIIDKWEKSPWADGKRPDVKLDGCWPHDYSMSVWMMGGVLVGHGTEELRTPRTKMDGGRDARVSRYREKRRTRLFSKKIRYEVRKLNAEKRPRMKGRFVKRANGVAVATACVA

>Bradi3g50166

MKVQCDVCAADAASVFCCADEAALCDACDRRVHRANKLAGKHRRFSLLNASPSASSASSPPPLCDICQEKRGFLFCKEDRAILCRECDVPVHAVSDLTMRHTRFLLTGVRISSEPAASPAPPSDQEENNADYCCSGDNAATSHGCGSSTSSSISEYLTKTLPGWHVEDFLVDDATAAAAAAASSSSTGISAHGSYSYQGETRNSGMHEAGYASWMAQERPFCDSVDVGTLAMSNRERWVPQMYAEPAGSKRSRTSAAYPYW

>Bradi3g48180

MKIQCDSCGVAAATVVCCADEAALCGRCDVEVHAANRLASKHQRLPLDALGAGKLPRCDVCQEKAAFIFCVEDRALFCRDCDEPIHVPGTLSGNHQRYLATGIRVGFGGPVSACADADHGASHDADHHAPPMAPAAERPPAQAQQVPSPPQFLPQGWAVDELLQFSDYESSDKLHKESPLGFKELEWFTADMELFQDQTPKGGRTAIEVPEFLNSQAADDAAYYRPSRGAATAVVRQSKKARIEIPDDEDFFIVPDLG

>Bradi2g06370

MKVLCSACEAAEARVVCCADEAALCARCDRDVHAANRLAGKHHRLPLLSSSSAALQSSSSAPNCDICQEGHAYFFCVEDRALLCRSCDVAVHTANAFVSAHRRFLLTGVQVGLQPADHQDPDPEPEPEPEPEPEPQPPPCKKRRSPTPPLYSDDDIGWAAGGITGTLPDWSAVDQQFSSSPPPTPRNPAAEEPVGVIKTPPKRIPRTAPVSAALFGGSMPDWPLDEFFRFADFNSGFGFADNGTSKADSGKLGSTDGSPNRRSLSSSSSGAAATQNAQEFFGQVPEVHWSSVPELPSPPTASGLRWQGDPHYGDTAAVFVPDISSPDNAFRCFASAGCGGTQTESLKRRRRC

>Bradi2g32900

MKVLCSACEAAEACVLCCADEAALCDRCDRDVHAANRLAGKHQRLPLLSPGSASADPAPPASPPKCDICQECHAYFFCLEDRALLCRGCDVAVHTANAFVSTHRRFLLTGVQVSLDEQEDDCLPDPPEPSPAPAPPPPAPAAKRDQASLYGEGDFSWAAATADANESLPDWSIVNEQFGSPAPRHAEAASASAAASRTPPKRAPAFSGQGGMMDWPLGEFFGGFSNFNAGFGFGFGESGTSKADSGKQGGSTGGSPYYVSSSDDRNADELFGQVPEMQRSAAELPSPPTASGLHWQRRPADYGAFVPDISSPDSSLRYCFPADQTAVKRRRKC

>Bradi5g17080

MKVLCDVCAAEAASVFCCADEAALCDACDRRVHRANKLAGKHRRLSLLHPSASPSSSAQKPPPLCDICQEKRGFLFCKEDRAILCRECDVQVHTASELTRRHGRFLLTGVRVSSAPADSPAPSEEEPEDEEEENSCSGGSGSASATASASASDGSSISEYLTKTLPGWHVEDFLLDDAAAAASAAAVYLDKPYQGVQAARTGGVQLQEASYTAWAGREQRIGGVPVAAGERARRELWVPQMSAAGAGAEWTGSSKRPRAPPPYSYW

>Bradi5g14280

MRIQCDACEGEAATVVCCADEAALCARCDVQIHAANKLAGKHQRLPLHHDSPSTRSSPAPRCDVCQDKPAFVFCVEDRALFCADCDLSIHVQGALSGNHHRFLATGIRVGFAFTTVCSTHAGERRPRAAPSKPPPAVVPAPAPAAAPQEVPSSPAYLSPSGWAVEDLLQFSDYESSGDKKGSTSPLGFKELEWFADIDGLFHHDSPLAPATKGVSRTAAAEVPELFASPQPAGNAGFYKAAGARPSKKARVELPDDDEDYLIVPDLG

>Bradi4g35950

MRTICDVCESAVAVLFCAADEAALCRSCDEKVHLCNKLASRHVRVGLADPNKLVRCDICENSPAFFYCDIDGTSLCLSCDMAVHVGGKRTHGRYLLLRQRVEFPGDKPGNMDDVPMQQIESENQRDQNKAPHSVPKEQMVSHHHAYDNHASDGNCNGQGNIDSKMFDLNMRPARNHGQGSSSQTQAVDHSANNHDSSGVVPTCNLERDTNK

>Bradi4g40250

MKIGCDACGRAAAAVLCCADEAALCRRCDAAVHSANRLAGRHQRVELLSSSSTGAGAGEGDGTHPACDICQEKTGYFFCVEDRALLCRSCDVAVHTATAQASSHRRFLITGVRVGGSANAQQDRHIVSPSSSSANGSSSINVPPASSRTRGLADNDHLSAQTAARLLGVGGEDEEEAAAAGQQQWPWSDIFADDSGVGNGMDDRQFYNGLVSEPAGSSGLTG

>Bradi1g35030

MRVQCDVCGVEPATVLCCADEAALCSACNRRVHRANKLAGKHRRLALQQPSSPTNATAAGPLCDVCKERRGIVFCVEDRAILCADCDEPIHSANDLTAKHSRFLLVGAKLSAAELVDQDHQIPSPDGSPDEHENSSSCAANGAEDSAPPVLNGVGGGGGGGSSISEYLTNICPGWRVEDLLLDDAAFASKQKGRDDEVPFLDADLFDVVAAGARPEKRGGAWDQAPHVPQPAWASLEEVVPAPKAKAKQQQGRVREWYHSDSDSDVFAVPEISPTPPAKRARPSPFWCL

>Bradi1g49260

MKIQCNACGSAEARVLCCADEAALCAACDEEVHAANRLAGKHQRVPLLSDAHAPTAAAAAEPPKCDICQDASGYFFCLEDRALLCRDCDVAIHTVNSFVSVHQRFLLTGVQVGLDPADPVPPIADKHVNVSGGSVDSQMKHLPRKNPTVLFSGETSVSIPSQNAISEDYSRQSPVPNIRTGMVNWTMNNSAIRSEEPPPKYLSEGSPTLLLSSQTTTAFSNQMNKDSDRAYNLPFSGGNGSDSLPDWHVDEFFSNSEYGPNLGFAEHGSSKGDNAKLGSAGGSPQCRLAEGLFAEDLLGQVPGFVAEDPWVVPEVPSPPTASGLYWQGNLRYPVYDHAMFVPEIPSLQSSQNHFTASDGSKRRRREF

>Bradi3g15490

MAASGGRDGGKGSGKGGCELCGAAARVYCCADDATLCWGCDAQVHGANFLVARHARALLCRGCARPTPWRAAGPRLGPTASLCERCVGRGRGGGDGEEEDEEMTVEEEEEEEEEEGEGENQVVPWAEDAEATPPPVASSTSSSSREAAPNAGHRCAKEDLPCSTSQPSSCRGGQSDEATSARNSSGGRHFFASGHRKRSTSDFLSSGSAQSGSGPPGRNRSTAAISRNE

>Solyc08g006530.2

MVAESWSTTAKRCDACKATPSTVFCKADMAFLCLTCDSKIHAANKLASRHARVWVCEVCEHAPASVTCKADAAALCTTCDQDIHSANPLARRHERIPVVPFYDSASASSSRGAAADGNDDPQQHDDDTEEEEAEAESWLLQAPSTNNNTQGIEYKSVEYLFSDVDPYVEMDIIADQKPSNDIAQLHNQEEYKEDCVVPHVQNNKNDIQLQGPVVDGYPTYEIDFSGGSKPFMYNFTSQSISQSVSSSSMEVGVVPDHNTMTDVSNTFVRNSAIDGLPNPVSSLDRKARVLRYREKRKNRKFEKTIRYASRKAYAETRPRIKGRFAKRTENEVGDSLVASDASYGVVPSF

>Solyc07g006630.2

MGIFREAPNCFPGGWNIGAAARMAKSCEYCHLAAALVFCRTDNTFVCLSCDTRLHARHERVWVCEVCEQAAASVTCRADAAALCVACDRDIHSANPLARRHERVPVVPFYDPVESVVKSTAATLLVSINGTTTTATTTATITPELGKVDTCIGHHENNNDPWIPPNTITSKLPLNTEMKGMDFIFTDSENFLDFDYPACVDTQSQPHYNSSNDSVVPVQANTPIKSLPFHHQEKHFEIDFTQSHIKSYNTPSLSVSSSSLDVGIVPDGSSISEISYPYIRTMNNSNSSIDLSNSANHQGEKLLGLDREARVLRYREKKKNRKFEKTIRYASRKAYAETRPRIKGRFAKRTDGSAGAGEFDDVDGIFSGTDFIAAESRYGVVPSFLT

>Solyc02g089540.2

MLKKENSNNWARVCDSCHSATCTVYCRADSAYLCAGCDARIHTASLMASRHERVWVCEACERAPAAFLCKADAASLCASCDADIHSANPLARRHHRVPIMPIPGTIYGPPAVHTITGGSMMIGGTTGEGTEDDGFLSLNQDADDTTIDEEDEDEAASWLLLNPPVKNNNKNNNYGMLFGGEVVDDYLDLAEYGGDSQFNDQYSVNQQQQHYSVPQKSYVEDSVVPVQNGQRKSLILYQTPQQQQSHHLNFQLGMEYDNSNTGYGYPASLSHSVSISSMDVSVVPESAQSETSNSHPRPPKGTIDLFSGPPIQIPPQLTPMDREARVLRYREKKKNRKFEKTIRYASRKAYAETRPRIKGRFAKRTDVEAEVDQMFSTQLMTDSNYGIVPSF

>Solyc02g089500.2annotation

MLKNENSGVFYGSRNNWSRVCDSCRSTACAVYCRADSSFLCAGCDTRMHAANLLASRHKRVWICEACERSPAAFLCKADAASLCTSCDADIHSASPLACRHHRVPIMTILVRLDSTFQEAFMVLQLLKLLAVPEDYGFLSFTQNADDMTVNEEDEDEAASWLLLNPPVKKNNKNNFDNDNNDQNNNYGTLFGGEVVDDYLDLAEYGGDSQFNDQYGVNQQQHHYSVPQKSYHGDSVVPVQEGQGKSLILYNQQQQQQIHHLNFQLGMDYDNSNTRYDTSQLTPMDREARVLKYREKKKNRKFEKTIRYALRKVYAETRPRIKGRFAKRTDVAEEDQMLSTQLMADGIYGIVPS

>Solyc02g089520.1

MLKKENSGGLDGSSNYWARVCDSCRSVTCTIYCQADSAYLCADCDARIHAASLVTSRHKRVWVCEACERAPAAFLCKADAASLCASCDADIHSANPLAHRHHRIPIITIPGTLYGPPAVETVGGDSMMISGSTGEGTEDDGFLSLTQDADDTIIDEEDEDEDEAASWLLLNHPVKNNNKNNVNNNNNQTNNYDMLFGGEVVDDYLDLAEYGGDSQFNDQYNVNQQQQQYFVPQMSYGGDSVVPVQDGQGKPLIFYQQQQQQQQSHHQNFQLGMEYDNSNTRLGYPASMSHSVSVVSMDVSVVPESALCETSNSQPRPQKGTIELFSGHPIQIPLLTPMDREARVLRYREKKKNRKFEKTIRYASRKAYAETRPRIKGRFAKRTDVEAEVDQMFSTQLMTDSSYRIVPSF

>Solyc12g096500.1

MGTENWSLTAKLCDSCKTSPATVFCRADSAFLCLGCDCKIHAANKLASRHARVWVCEVCEQAPASVTCKADAAALCVTCDRDIHSANPLARRHERFPVVPFYDFAVAKSHGGGDTDADAVDDEKYFDSTNENPSQPEEEAEAASWILPTPKEGTDQYKSADYLFNDMDSYLDIDLMSCEQKPHILHHQQHQHNHYSSDGVVPVQNNNETTHLPGPVVDGFPTYELDFTGSKPYMYNFTSQSISQSVSSSSLDVGVVPDHSTMTDVSNTFVMNSSSGAIAGAGADVVPNAVSGLDREARVMRYREKRKNRKFEKTIRYASRKAYAETRPRIKGRFAKRTETEIDSLITVDASYGVVPSF

>Solyc07g045180.2

MGYICEYCGEQRSIVYCRSDSACLCLSCDRNIHSANALSQRHSRTLVCERCNSQPAIFRCVEERVSLCQNCDWLAHASSGTCSTHKRQALSCYTGCPSAVELSTIWSFILGDPSVCDSTCEQGMGSMSITDCQPGDSQHPQVKEKSQDISSDEAKDLHNLVKSAPFMGSSMPSLDNELPNVELLVASTNLTWSKVKNSGTKGYNDPFYDDFNMDEVDLSIENYKELFGVSIDNCDQLFKNEDIDDFFGMKDMSVAESSFQGVNAVEVIVDNGKAPGSAVGKVNAVKSACSNAASAESMMSCKTEPPTLCFAMQQSSFSFSNLTGESNAGDCQDCGASSMLRMGEPSWHPPCPESSMPSTSRSDAVSRYKEKKKTRQFDKRVRYVSRKARAEVRRRVKGRFVKDGDAYDYDPLHQTRSY

>Solyc09g074560.2

MTEVKKDEENHHQHLCDFCGNNTALLYCRADSAKLCFTCDREVHSTNQLFTKHTRWLLCNLCDSSPASILCCTETSVLCQNCDWESHNKLLSLHERRPLEGFTGCPSVSELLSILGFEDLGKKELLCGGDDGAYGFSDWVIWDAPSVVTLDDLIANNNESRHNYQAIGVPPLPKNRNAACGKHKEEILGQLRELSKLEPNSGDDQDENVPTTGFQSMEPVQNCPLRYKGSGFMQRSDQHVVPSSEGSAFHWHGDTGEFVDQGFSSSLTDCFIETKCLLPDRDSDVCDASGGGNEEQSHHPTTTETFQMVPKVVHRELNSQERETAVSRYKEKKKTRRYEKHIRYESRKARAETRTRIKGRFAKMDYRDSSVHQ

>Solyc05g024010.2

MSPGESRPCDFCNQQIAVLYCRADTAKLCLFCDQLVHSANALSKKHLRSQICDNCGSEPVSIRCDTDKLVLCQECDWDAHGSCAVSGAHDRSPVEGFSGCPSASDLASAWGLDIESKKLHQQHTVLEYPSWMSKDAPPSSVLLQDLMVPSAINSAIYSTKQTPTCGKQKQVIFKQLIELFKRDLADGVGAGAEDLVPKTPNATSDWQGNINVSIMDGVTQKPQQQQPQNVPFTSSIMPHNPKDSDQMVERNILWRGNSFDQNTQIWDFNLGQLRSHEQSSSLEADYSESDMACMMKSYGELIKGTSLATSKVLGLSGINCSVVHDDMTAFSNNSNNRAASQGPATSESNNLPRIKTSSDLGCVKPKCGGVSTDLNFMDQSIVVGGDNTGEETLKADMELLAKNRGNAMQRYKEKKKTRRYDKHIRYESRKARADTRKRVKGRFVKANESPDG

>Solyc12g006240.1annotation

MGYICEYCGEQRSIVYCRSDAACLCLSCDRNVHSANALSQRHSRTLICERCNSQPAVVRCVEERISLCQNCDWSGHASSSSGSSMHKRQALNSYTGCPSAAELSNIWSFLLDDPSIGDTCEQRMGSMSINDNRPRDGQDPQGKDNSQNVCAAVEANDMSISEKSNLLVESSMPTFDNKLHNMEPPIGSSSKGCYMGAKGSNLFEEDPYCDNLIMDAVDLSIENYEELFGDSLNYPDELFENENLDSFFGMKDIKGADYSYHGVNATEGLSNARVNTVQPTCSDAASADSIMSCKTDSILYFARQSSLSVSNQTGGECSAGDHQDCGVSPMLLMGEPPWCPPCPEISSTSTSRSNAVLRYKEKKKTRKFDKRVRYVSRKARADVRRRVKGRFIKAGDAYDYDPLPTRSY

>Solyc04g007210.2

MSSEKKLANAMGAKTARACDNCIRKRARWYCAADDAFLCQSCDSSVHSANPLARRHERVRLKTSSFKSSDDFPNLESTVSGLGSGSGSGSDSIPSWHCGFTRKARTPRYGNKHAKRVKSTEEEEEEEEMKNPIQLVVPEILSDENSHDENEEEQLLYRVPIFDPFMADGSNYGNEYSSNKVDFNQDMNTFQGLLAPSEMELAEFAADVVSLLGKGLDDEESFNYMEGLGFLEKHDEKLVKVEDEGEVGFVNMISTNNQVDYSEFDMVGETFELKFDYDSQVINNLDEDNKKVEFLEINYDSGKNNNKIMLNLDYESVLKSWGDKRFPWTTGVRPEVDFNDCWPVCMGNCGKIHSYGDIAIMNGHGGGVVDEGREARVLRYKEKRRTRLFSKKIRYEVRKLNAEKRPRMKGRFVKRTNFAPTPFPSLNK

>Solyc03g119540.2

MMTSESKTANAIGGKTARACDSCLSKRARWFCPADDAFLCQSCDVSIHSANQLASRHERVRLETCSNKSTITKLVDKTHQPSWHQGFTRKARTPRNGKKAQIRQWKKNEENRVPEIGSEENSLDENEFENEEQLLYRVPIFDPFEAELCNVPDETGSIVDLDILLNTEDACHDLNLPEFLSSDIELAEFAADVETLLGGEEEQSTQLLNADFENNKAIKIEVEDEEMRAVVACHLDPELDMEREALNWNFDEYYEETVEQKVMAADAAVTEFIASAEEYSGSSTKSDEKNRLFLSLNHEAVISAWPNQSSPWTNGIRPHFNPDDYWPDFFSETSVGNYGGHVRCGDGGREARVSRYREKRQNRLFSKKIRYEVRKLNAEKRPRLKGRFIKRSSSLFSVPGFSYMMNKR

>Solyc05g009310.2

MVSERKLASAMGGKTTRACDNCIKKRARWYCPADDAFLCQNCDASVHSANPLARRHERVRLKTSSLKQTSSPSSSSDDYFPDLESPLSISSVSVSVSVPSWHRGFTRKARTPRQGRKASKSAGDGDVIRKNPIHLVPEILSDENSLDENEEEQLLYRVPILDPFVGHLYSSSTAPTDADSEFKLESKEMTLQDDICNVDLNRFHEMLPSEMELAEFAADVESLLGKGLDDESFDMEGLGLLGVCNKEENSMISHEKVKIEDEGEMEVVTKTTSPTTHNHQYNHTHDHDIDINEDTFEFKFDYDSSINIIGDDEVVTNDENKKKILLNLDYEGVLKAWADQRCPWTNGERPELDSNESWPDCMGNYMGIMNENVTIVDRGREARVTRYREKRRTRLFSKKIRYEVRKLNAEKRPRMKGRFVKRANFVTTSTPNYPLVK

>Solyc07g062160.2

MKIQCNVCEVAEANVLCCADEAALCWSCDEKVHAANKLASKHQRVPLSGSSSSMPMCDICQETVGYFFCLEDRALLCRKCDIAIHTANPHVAAHQRFLLTGVKVGLEPVDPGGISSSGTSQSIQKVSEPESAPLSKRNASVSLDAQFNKVLPTQVSGIEDFAPTKSPFAGGSAAGSMPQWQFDEFIGLSDFNQNYGYMDDGSSKADNGKLGGESDSSSILRVEDEELDGDECLGQVPDTSWAVPQVPSPPTASGLYWPKTYQNPFDSAVFVPDISYSPSSSLQQQPPSGTRLKRRRQC

>Solyc04g081020.2

MKIQCDVCNNNEASVFCVADEAALCDSCDHRVHHANKLASKHQRFSLIQPSPKQIPVCDICQERRAFLFCQQDRAILCRECDVSIHKANEHTQKHNRFLLTGVKISANSSLYTSSESVSAASCSANQDSVTNLNKPQICTKKTSPVSGSVPQQQVSVAANIGENSYTSSISEYLEMLPGWHVEELLNASTIPTNGFCKIGDNDVFPIWDSEIESSMNSFSPENIGIWVPQAPPALTPQKNQNQVFPRNINFGGQIEFKNMKEVTSKKSSRKWRDDNSFAVPQISPSSSSISFKRSRTLW

>Solyc06g073180.2

MKIQCDVCEKAQATVICCADEAALCAKCDIEVHAANKLASKHQRLHLQCLSNKLPPCDICQDKAAFIFCVEDRALFCKDCDEAIHSASSLAKNHQRFLATGIRVALSSSCNKESVKNQLQPQPPQQNSQQVGLKMPPQQLSCITSPSWPVDDLLGFPDYESSDKKDLLELGEFEWLGGIDLFGEQTAAEVPELSVPQSSNTNIYRTTKYQMPYKKSRIEIPDDDEYFTVPDLG

>Solyc02g084430.2

MRTLCDVCESAAAILFCAADEAALCRACDEKVHMCNKLASRHVRVGIAKPNEVPRCDICENSPAFFYCEVDGSSLCLQCDMMFPGDKSGSTEELARKTLDTGENKKDHSHSPNPMIKDNQQNHRVSPILISDGSADGHGKKDKMIDLNVKPSRFHGQASNQ

>Solyc02g084420.2annotation

MRTLCDVCESAAAILFCAADEAALCRACDEKVHMCNKLASRHVRVGIAKPNEVPRCDICENSPAFFYCEVDGSSLCLQCDMMVHVGGDKSGSTEELARKTLDTGENKKDHSHSPNPMIKDNQQNHRVSPILISDGSADGHGKKDKMIDLNVKPSRFHGQASNQ

>Solyc01g110180.2

MKIQCDVCDKEEASVYCSADEATLCQSCDYQVHHANKLASKHLRFSLIHPSFKDSPLCDICQERRALLFCKEDRAILCKECDLPIHKANEHTKKHNRFLLSGVQLSSDILASNYNNNQNSISPAGSAASNAGTNNFKALSGNFGMKSNSISSTTESTHNYFHVDYVQEGSVSTSSISEYLTETLPGWHVEDFLEYPSSSSYEF

>Solyc01g110370.2

MRTLCDVCESAAAILFCAADEAALCRSCDEKVHLCNKLASRHVRVGLADPSKIQRCDICENAPAFFYCEIDGSSLCLQCDMIVHVGGKRTHGRYLLIRQRIEFPGDKLGPSNEQGLPSTEQGDVRRETAQPFKLPMIDNHQPNRETAMTAVENNVNNSVKVENELIDLNSRPQRMHGQTSNNQEQVMDMLGGSNHESVGVVPDGPFKREPEKCELLPFQISNLNYELELPAPAEFYAMSKL

>Solyc12g005420.1

MKIQCNACEVAEAKVLCCADEAALCWYCDDKVHAANKLANKHQRVALSASSSPMPKCDICQETVGFFFCLEDRALLCRKCDISIHTVNAYVSSHQRFLLTGVKVGLEPLGPCASASSRKSHEQRSPPILKITPPSEGVLPVHTSGNGNFAPSRLPIVGNIPQCQFDQYLGMPDFNQNYGYMDYGQSKAGNGKVGESASSPFLRDVDAEVAGDECFSTEVPDTCWAVPQIPSSPTASGLNWPTKSIQNPFDAALLESDASYFPLQNIQDQQSNGSGTKRSRRF

>Solyc12g089240.1

MKIQCDVCNKKEAIVFCTADEAALCDDCDHRVHHVNKLASKHQRFSLVQPSPKQAPMCDICQERRGFLFCQQDRAIMCRECDIPIHKANEHTQKHNRYLLTGIKLSANSALYSAPSQSQSQSQSAISSADSCVSNLKSKDSTSKPVAGSVFVSPAIISNSTKGGAVSSAVESVKVVKEKVGGCNNNVQFVNGGGNNLTSSISEYLEMLPGWHVEDFLDCSTPNVYSKNIGDEDMLSFWDTDLESQFSSFPPQNVGIWVPQAPPLQESKQETQIQFFPSQNLNFGGQIGLKESREVTNIKSSRKWTDDNSFAVPQMKPPSTSFKRTRTLW

>Solyc04g007470.2

MEMEKACEFCMLLKPVVYCEADAAHLCLSCDAKVHSANALSNRHPRTLVCECCGHNPAYIRCSDHQTFMCRDCDRCHHDLSSQHQRKVITSYMGSPSAKDLAALWGFGLKDLENATPPDQFISTSNGKANGVKVISKKFKRSHSSPGGSSLASELDFTGLVVSPESEVGSTSYYTKVLSLRKRRENTSLILQQILDLERLQLTEGSNNLTSGESRNNVSSLKNCTSWNMHNKFDCLQSSLDLGPELQDWGSTHESPVDESFPLPLPDGDSFWECKSPVQSSQLWPQNLQDLGAYAELERFDNSNMPDVDLTFQNFEELFGEDQYLNNTLLEEDMTCSSMENDSSIDRSDYSYVKKDISTASSVRTGHSTHFGQAHGEHIPTIKDCPPPIRTNFSSLSFSASRLSSESSGNEYVDSPAANDQEVSCNSQMDSKKKARLYEKQARNTPRRARTNFKKQHVLKAHCYETDALNMSRSF

>Solyc05g020020.2

MCEFCGEQRSIVYCRSDAACLCLSCDRNVHSANALSQRHSRTLLCERCNSQPAIVRRVEEKVSLCKNCDSIGHAGSGTGSVHNRQALSSYTGCPSAAELSTIWSFLLDNSLGGDSTCEKGMGSMSITDNRLTDSRAPQGKFNSQDASATVEVSEIHTPGKSSILVGSSMPNLGSKLNKVEHIAGSVNISSSKDCYSGVKASTIYEDDPFSQDFNMDEVDLSFENYEELFSGSLDNPNQLFENDDIDGLFGTKDMSVSDSSCQDANAVEGSSIRTVMSILPACSKAESADSLMSCKTEPTVCFARQASILSFSNLGGESNGGDYQECGASTMLLMGEPPWCHPCPESSLASSSRSDAVLRYKEKKKTRKFDKHVRYASRKARADVRRRVKGRFVKAGDAYDYDPLNETRSF

>Solyc05g046040.1

MDPLCDLCGEVRAVVYCKSDSARLCLQCDDYVHSPNLISRRHSRSFICDKCNSQPSIVRCMDEAISLCERCDWDGNGCIGTGHRLKKLNPYTGCPSPDEFTKMLSQVLEMPIGTDTNFGSFGNSLGSSLSINENNSSLENKVNEDSFVSSKLLASNYKFEAWSIPPEPNYLNSYQIDLAPFSEGSGLSKQDCPIKDLGLQEGDDLSKGVDFDDVTLDFNCSYEILPDSRQTGFSDENKELDCLVMEKNSSVTGSNNVETSHEATSSVQQEYMGLQSSQISAAASSTNLLQTMSANANCMLMNSTCNGSIGALPFLPAPIHPSMSLSLSNITGESCAEQDCGLSPGFLNEAPWDLSLENCPQKRHEAKMRYNEKKKTRTFGKQIRYASRKARADTRRRVKGRFVKAGEAYDYDPSETRDF

>Solyc07g052620.1

MCSGRREGDEKTSSTSYCKGPSKEGESIISSAITCALCSSEASVYCEADNAFLCRKCDRSVHGANFLAQRHIRCLLCSVCRKTTRRFLIGTSSELILPTIARLEQRNRSRSAESETTDYRTTPQELFLFI

>Solyc07g053140.2

MKNCELCTGLARMYCESDNASLCWDCDAKVHSANFLAARHSRSLLCQVCQSPTAWSAAGAKLGKTVSVCDKCVDGYYHRDGVEEMEESESVNDEESDTEDEETDYEEDEDEIDSGDIQVVPWSNTTPPPPASSSSSEDSSNGCRNVSSKRMREADPDLQSDDDSGSSSCQRKVHISPAMDADGGDDAAASVDSYSSKMRTPVKIQRTDRLTAPRCTPVIEFIRRNNRQRMNSGAAIAELCRLNHESRTTDLKSSETS

>Solyc06g063280.1

MELLSSKLCELCNDQAALFCPSDSAFLCFHCDAKVHQANFLVARHLRLTLCSHCNSLTKKRFSPCSPPPPALCPSCSRNSSGDSDLRSVSTTSSSSSSTCVSSTQSSAITQKINIISSNRKQFPDSDSNGEVNSGRCNLVRSRSVKLRDPRAATCVFMHWCTKLQMNREERVVQTACSVLGICFSRFRGLPLRVALAACFWFGLKTTEDKSKTSQSLKKLEEISGVPAKIILATELKLRKIMKTNHGQPQAMEESWAESSP

>Solyc10g006750.2

MMNCELCGENKSMMYCESDQANLCWDCDSKVHSANFLVAKHSRNLLCHSCQNSTPWNASGPKLSPTFSLCNSCIQNPDPIEQIGETIEENYQTEADNDEDEYVSTDSEIDDYDDENQVVPLSSSPSSDFSPSPSPVRSVSSSGSYGDLSAIGDYGGGATAADNSSPWKRVRESDCLHFEGESVPPTLHPRIEISGSE

>Solyc02g079430.2

MKKCELCSSIARVYCESDQASLCWDCDARVHTANFLVAKHSRILLCNSCQSLTPWTGSGSKLGPTVSVCQKCFHRNNNDGADQHLNDHNDTEEEEDDDDDTEEDEDDGSSDGDNQVVPLSSSTSDQPNPPSLTSSSSSSEDSTNRFNRRSIIGFSPNRTEENIQSHYCGITRPKEKIERSSWRWR

>Solyc12g005750.1

MCNGRREIDEEKIEELHNIIVCELCKSEAYVYCEADNAFLCKKCDKLVHTANFFAQRHIRCILCGICKKLTKRYLIGVSHEVILLKVVRCTNFDEQNCSTKVKEPFLFL

>Solyc12g005660.1

MKNCELCNGLARIYCESDHANLCWDCDLKVHSANFLVAKHSRSLLCNVCRSPTVWSASGAKIGRTVSVCERCVNDEETDREEEEKEEIDLEDIQVVPWSSTPPPQPESSSSSSDESLTDFFSLKRMRKDDDSGTLTSHREVNMLSPATEDHGGDEAAEFVKSFPVKMPSKIQRTGEILSGRVENIGCSATVELIGRINRGGG

>NNU_04365

MMLKEEGSSGGSRGGWARMCDSCRSAACAVFCRADSAYLCAGCDARIHAVNQVASRHERVWVCEACERAPAAFTCKADAAALCTTCDADIHSANPLARRHHRVPILPIAGCLYGPSPTNPGRRVRPDINIEDGFMTQEGDEDIDEEDEDETASWLLLNPGKNDNNQNNNGFLLGGEVDEYLDLVDYSSCTDNQYNQQQHYSVQHKNDGSDSVVPVQCLAVKEEQQQQNLHLDMDYDASKPGFSYSHSVSLSSMDASVVPDAMMNDISNSHSRPPKGTIDLFSSPLAMPPQFTPMDREAKVLRYREKRKTRKFEKTIRYASRKAYAETRPRIKGRFAKRTDVEVEVEQMFSTTVLAESGYGVVPSF

>NNU_06077

MKKREMVIDDESGKGYPASWSLSAKPCDSCKSASTLLFCQADSAFLCFNCDAKVHGANKLASQHERVWMCEVCEQAPTSVTCKTDVAALCVTYDQDIHSANPLTRRHERIPIVSFYESAASAMKSNIANLLVPDGLLKSSDNDDDHNDKEEDKPRGGRSCFLVTSKPKPYQTHGNTQFEVRRLFLLQHGSLSRSRLRVVHGDEVSPPEQRQC

>NNU_07533

MMIKEEGGSGGTGGGGGWARMCDSCCSAPCAVYCRADSAYLCAGCDARIHDQVVSQHERVWICEVCERAPAAFTCKADAAALCTTCDADIHSANPLARRHHRVPILPISGCLYGPSGMNPGRPVRPEINIEDEFMTQEADEEDEDEAASWLLLNPVKNNNQNNSGLLFGGEVDEYLDLVDYNSCTENQYNDHYYQQNQCSVQQKNEGSDSVVPIQCLAVKENQHNLQLEMDYEASKPGFSYTASLSHSVSFSSIDASVVPDGIMNDISNSHSRPPKGTIDLFSSPPLPMPPQFTPMDREAKVLRYREKRKTRKFEKTIRYASRKAYAETRPRIKGRFAKRTDVEVEVDQMFSTTVLAESGYGIVPSF

>NNU_09057

MAIDDESDKGYSASWSLSAKPCDSCKSASALLFCRADSAFLCFNCDAKVHGANKLASRHERVWMCEVCEQAPANVTYEIDAATLCVTCDRDIHSANPLAWRHKRIPVVSIYESATPAMKSDSSDDDDDRNDKEEDNDTCRAEAEAASWLLPNLAKLMETPDLKSGDYFFSDVDPYLDLDYGSSMETRYHHQNNANADSVVPVHTKPPPPPLMNTFDRCFDMGFCRSNPSYGYTTPAIIQSVSSSLLDIGGSFIIATTYAVFFDKCYSVSPRAKACKIGSEGRQKKRFLSTVHSCITELKFVKDELEALCGEYAYLVTEKDMAVKKAGEVVSAPKAIEKIVENLTLELIAL

>NNU_15002

MAIDEESGKGYPASWSLSAKPCDSCKSASALLFCRADSAFLCFNCDAKVHGANKLASRHERVWMCEVCEHAPASVTCKADAAVLCVTCDRDIHSANPLARRHERIPVVPFYESAASAMKSNVANLLVPDGLLKSSDEDDDRNDKEEDNDTCREEAEAASWLLPNPNPPKLMETPDLKSGDYFFSDVDPYLDLDYGSSMETRYHHQNSANADSVVPVHTKPPPPPLMNTSDRCFDMGFCRSKPSYGYTTPTLSQSVSSSSLDVGVVPDGNSNAISDITNPFGRKGGVDAGGGNHGAPQLSGMDREARVLRYREKRKNRKFEKTIRYASRKAYAETRPRIKGRFAKRVEIDSEVDRIYSSTAAFIADTGYGVVPSF

>NNU_15296

MAIDDESGKGYLTSWSLLAKPCDSCKSASAVLFCRANSVFLCFNYNAKVHGTNKLASRHERAWMCEVCEQALASVTCEIDAAALCVTCDRDIHSANKLAQRHE

>NNU_19742annotation

MAIDDESGKGYPASWSLLAKPCDSCKSALALLFCRADSTFLCFDCDTKVHGTNKLASRHERVWMCEVCEQAPASVTYETDAAALCVTCDRDTTRQTHSLGGTASLLVPDGLLKFSDVDDDRNDKEEDNDTYREEAEAASWLLPNPAKLMETPNLKSGDYFFSDVDPYLDLDYKSSMETRYHHQNNANADSVVPVHTKRPPSPLMNTSDRCFDMGFLVLDGNNNAVLDITNPFGRKGGVDASGGNHGAGAGGK

>NNU_23689

MAIDNESGKGYPASWSLSAKPSDFYKSASNLLFCRADSTFLCFNYDAKVHGTNKLASRHERVWMCEVCEQAPVSVTCKTGVAALYVTCDRDIHSANPPTWQHERIPVVPFYESAASAMKSNVANLLVPDGLLKSSDDDDDRNDKEEDNDTYRKEAEAASWLLPNPNLAKLMETPDLKSGDYFFSDVDPYLDLDYGSSMETRYHHQNSTNADSVVPVHTKPPPPPLMNTSNHCFDMGFCRSKLSYGYTTPTIS

>NNU_01295

MVNFRGDPGRREQQQPRYVVEEAAEQDEQLQQQQHLAAKVTLHGLCDFCGESRAPLYCRADTAKLCFSCDREVHSGNPLFGKHSRSQLCDACASKPASIFCSTDNLVLCQNCDWDAHGGCSSSSSSSLHDRRPLEGFSGCPSAVDLSSILGFEDVGNKSLLLMGDAMDDCLVVSGPYGSMGMADSFTDELVRDTPVVSFDDLIVSSDSTHNFLPMVVPPLPKNRKSACGKHKEEILRQLCKLAKLERSLSSENGVVESFLGLQPSVQERSFQTGYVDTGIEHGTGPTLAPGFDVNAFQWHGDYSEAADQVPILITFPDNRSEENFLVLGEPGEDGSSGSHTNVGCKEQSQHLIIRESLQGLQKVGQHEPTRIDRDLAISRYKEKKKARRYNKHIWYESRNVRAEGRTRIKGRFAKAND

>NNU_01969

MDQNPNKSRATQSIILCDFCNEEIAVIYCRADSAKLCLFCDQHVHSANALSRKHLRSQICDNCSSQPVSVRCSTDHLVLCQDCDWDAHGSCSLSATHERCPVEGFSGCPSALELSSIWGFNLADKKHAMQLPQSQVSQMGSDNFMMFPNWGSLNSVVSDDSWVYKSAAMTLQDLMLPSENGNSGTSMYPSVPSADIPLLSKKLSSSCGKQTQVIFKQLVELLKRDPVRGDGGGGGDGLGPGTPAQQSNMEDLELPDGGDGATDATQLLHQQTPFTSLLMLSTRMDLREINNRLIEDNIPWDCNPSDQAAQVWDFNLGKSRDHDEAGPLEAQYAASDAGFMFKSYQDLIKETPLPAKKVYDNIYDISYSMAHDNKSSQNNNSNNPAASQGPTTSESNNFPLRKPSSSSSLGKYKTYSGTRDVDFMEQSVFLRSDTLKPATTKADMELLAQNRGNAMLRYKEKKKTRRHDPPYLLYDKHIRYESRKARADTRKRVKGRFVKAGEAPDVEIKKS

>NNU_04012

MGQLCDFCGEQRSMVYCRSDAACLCLSCDRNVHSANALSRRHSRTLLCERCNSQSAIVRCVEEKISLCQSCDWNGHGSSTLGSAHKRQTINCYSGCPSAAELSRIWSSIIDSPSIDNSNCEQELGLMSIDDNSISNRWGPPEDNSMIDLSGDNILNELENMDKFGILMGSSSVPALNPIPYDVDQLAGSMDSTTPKLSCPGTKDLGLGEGDDLCEDFSVSDVDLNFENYEELFGISQNQSEQLFENGGLDNLFGIKDISGADSNYQDDFVAEGMSVGKIQAMQPTCSNAASADSMMSAKTEPNLYYPPRQAHSGLSISFSGMTGESSGGDYQDCGVSPALLMGEPPWHPPCAENSFPSTSRDSAVMRYKAKKKIRQFEKKIRYASRKARADVRRRVKGRFVKAGDAYDYDPLCQTRSY

>NNU_06053

MGQLCDFCGEQRSMVYCRSDAACLCLSCDRNVHSANALSKRHSRTLLCERCNYQSAIVRCIEEKISLCQACDWNGHGNSTSGSAHKRQTIKSYSGCPSAAELSRIWSSVLDFPSMDDSNCEQGLGLMSIDENNISNCWDHPENNNIIDFACASRLDELENADKFGIWMGSSSMPALSPMECGVDHLVGSIDSTTTKVTFDFKSVYGELNRMNS

>NNU_06912

MAEFIRDPQSPEEEQEQPINVKGKEDDEQRQQQQNRETQKCLCDFCGELRALIYCRADTAKLCISCDHDVHSGNPLFGKHSRSQLCDACASNPASIYCSTENLVLCQNCDWDSHGGSSSTLHDRRPLEGFSGCPSAIELSSILGFEDVGYKSLLLREDDPEGNSFVSVHYGSMGDKLTDDLAWDTSVVSLDNLIVSTNLSQNFQAIGIPPPPKNRKTACGKHKEEMLHQLCKLAKLESLSSMNIEPGEIESLLGLETPILEQNLQSGYMHTGIEHNTGITLLSGYDASVLQWHGDCGEPADQVPISLTYSENHLEESLLVLGKPTEDCGTVSHANAGYKEQSQCPIVTESLPILARTFQHELTRIDRGSVISRYKEKKKARRYDKHIRYESRKVRAESRTRIKGRFAKVDNIN

>NNU_16442

MDPVCDFCGVARAVVYCRSDITRLCLNCDSFVHSANAISQRHCRSLICNKCNSQPAVIGCLEDKMYLCQSCDSNGCGCTGPGHRRQAVTFYSGSPTALELSRIWPSNSNASSPPREVNASLEPPGLMSMNENCISNSWEPRDNGSSSAGLTTTNRLNELEPCTTFDPWACPSSVIQHSVSSMPYCEDKPPFFPEELKLSKAVSYDVQQGCSPLKDLEICHGGDLCEGLNMDGVELNFENSDEIFGCSESQPSHSFEDIGMDCLFMDKNFSVADSSCPSENVLEAPYISCMADNHLCSHLHQASSSSGQPDYIALQSSYGAGSATAVQGINSATDRVFLNPNHNKNINLSFPPQQVNSSLSLSFSNLTGESSVSDYQDCGVSPMFLTGESPWDSNLDTSCPQARDKAKMRYNEKKKTRIFGKQIRYASRKARADTRKRVKGRFVKAGEAYDYDPLILLLPRAFHRA

>NNU_17261

METPKEKLKQQRNVSRRANGLETKKMEKVCEFCAESKPVVYCKADAAYLCLSCDSRVHSANSLSEKHFRALLCDGCRNHPAYSRCLDHRMLMCHGCDQKLHAIDSQHKKRAMSSYRGCPSARDFAVLWGYDLNELCASGFQDQLVSASSGSLSPSVANLDIARNLCPLTEVSSLTEVESNSLQGTQRETCIILQQILDLERLQLTERKSHSSLIRAEMQADIYSSKNNNSGPLGEDLDHNFLNPQGLSIDIQQMDRMNHELQVQPFPLPFSQLELLNSSSSSEIPLHGDIYWQCKSPAQTSQLWTQTMEDLEICEETVCNDDFNIPDVDLTFQNYEELFSGDQDHIRSLFGEKDIGFSSMDKDISLEKSDNGYSRAIEDISVVPSVYVSQSSHQYKSIGHSDQSYQIGGSMDSHHLIRPSYSCLPFSVSRLSAESSASDCLDSGISPNTIQASWNSPDVENTHSEAKENAMIRYKEKKMARTYEKRIRYASRKATADVRKRVKGRFVKAEVYESDTINMTRSC

>NNU_22255

MEKNPRLTNSEGIPCDFCNEQIAVLYCRADSAKLCLFCDHHVHSANALSRKHLRSQICDNCSSEPAPIRCSTDNLVLCQECDWDAHGNCSISASHERNPLESFSGCPSATELASVWGFDLGDKKSLMPLPPSQLSQDNHMFPSWSALDSVVSVDSWVYKYAPVTLHDLMSPNENGPPLYPSVPSAEIPALSKRQNSNCGKQTQVIFKQLVELLKRDLVSGDGVVDELGPGTPGRTTQPGNIEALDLPDGGDVVADTNQLLQQQTPFTSLLMLPARLDPRESNRLIEDNIMWDCNPTDQAAQIWDFNLGKSRNHEEAGPLEVRYGSSNAGFIIKSYNDLIKGTSLTAKKVVDDIYEMNWFMTHDDISSQNNNSNNLAASQGPTTSESNNIPVLRASSSSTLAKPKACGNSKDAHFLKQSLLVSSETARAATKADMEMVAQNRGNAMLRYKEKKKSRRYDKHIRYESRKARADARKRVKGRFVKASEALNAERNS

>NNU_00499

MSSDKKVANAVGGKTARACDSCLRKRARWYCAADDAFLCQGCDASVHSANPLARRHGRVRLKTASFKPNDDSLVENYVPAWHQGFTRKARTPRHGKPTALQQSKSTDHEEPIPNPLPLVPEMGSEEASPDENEEQLLYRVPIFDPFVAELCTPATSNEASMTANDSSTAAVGTETKAFLHEGHDVTGDMDNLPGFLPSDMDLVEFAADVETLLGKGLDEDSFGMEGLGLLDCKDEEKTDCCFGSGRVKVEEEEVEAAVECHVEVELEHDMARETLDLNFDYDSPTTAEDEVEEKMVTGATMMVTNGGCKAEETTRKTSLKLDYEAIISAWANKGLPWTTGDRPLFNPDDCWPDCMGTCPMTEVHHYHPYGEMGGVGGHVTAGDGGREARVSRYREKRRTRLFSKKIRYEVRKLNAEKRPRMKGRFVKRASCTGPAAFPF

>NNU_07680

MSSDKKAANAVGGKTARACDSCLRKRARWYCAADDAFLCQGCDSSVHSANPLARRHERVRLKTASIKPMLNDGFLVENSAPAWHQGFTRKARTPRHVKPAALQPPKVEEPILLNPLPLVPEMGGEESSPDDNEEQLLYRVPIFDPFVAELCTPATFNEPAMTTNDATTAEDMGSERKALLPDGHDFDNDLTGETDKLPVFLLSDMDIAEFAADVETLLGRGLDDDSFGMEDLGLLDCKEEENMEHCFRSGRVKVEVEEVEAVVGCQQVEPGLGMTRESLDLNFNSDSPTTAEEEEVEEKAVKEAANSNVVTNMNSGYKGDKMSSKISLRLRLNYEGVICAWASKGSPWTTGERPQFNPDDDCWPDCMGTYPMEVYHHHPYGEMGGVGGHVTMGNGGREARVSRYREKRRTRLFSKKIRYEVRKLNAEKRPRMKGRFVKRTSLMGVAAPFPF

>NNU_03247

MKIQCDVCDKDEASVFCSPNEAALCYECDCRIHSANKLACKHLRFSLLHPSFKVAPHCDVYQERRAFLICQEDKAILCRECNLPIHMANDHTKKHNRFLLTGVIHLCTPPTHPPMKHALLTGHWHVDTGLRVDPIRVFSRAGIDAQRGTNGATTHGELG

>NNU_08052

MRTLCDVCESAAAILFCAADEAALCRSCDEKVHMCNKLASRHVRVGLANPSDVPRCDICENAPDGSSLCLQCDMIVHVGGKRTHGRYLLLRQRVEFPGDKPGNLEDLALQNTDPGETRREQNSLPKLTARESQQNHRVSTVQVLDANNDNHGKKDTKMIDLNARPQRVHGQASNNQAPGMDVLSGSNHEAAHIVPIGAFKKEPEK

>NNU_08086

MKIQCDVCDKDEASVFCSADEAALCDGCDRRVHHANKLAGKHLRFSLLHPSFKEAPRCDVCQERRAFLFCQEDRAILCRECDHPIHTANDHTKKHNRFLLTGVKLSSSSSLYSATNSSTNEACDSLNGKVNENKISKLSKKQKPMPVPFGEIVHPKPPMVEKPPAAATITTSKADDHPASEGYSTSSITEYLMETLPGWQVEDFLDSYSAPHGFCKGDALSPFLDMDFQSNLSFSPQDTTIWVPQVPQASYPVNCLPNQISFLSGFKELKEASNFKTSNKRWRDDSLTVPQISSPSCERSTPFW

>NNU_17879

MKIQCDVCEKAPATVICCADEAALCAKCDIEVHAANKLASKHQRLLLQCLSDKLPRCDICQEKTAFIFCVEDRALFCQDCDEPIHSANSLSANHQRFLATGIKVALRSGCAKDTNKGFLEPTNQKPQPMAIKMSSPQPSTFASPPWAVDDLLQFSDFQSSDKKESIEFGELEWFTEIGLFGEQVPQEALAAAEVPELPILQPSNAMINRPTKPHMPLKKPRIEISDEEEFFTVPDLG

>NNU_17889

MKIQCDVCEKAPATVICCADEAALCAKCDIEVHAANKLASKHQRLLLQCLSDKLPRCDICQEKTAFIFCVEDRALFCQDCDESIHSANSLSANHQRFLATGIKVALRSGCNKDNNKGFLEPPNQKPQPVAMKIPSPQPSTFASPPWAVDDLLQFSDFEPSDKQKESVEFGELEWFTDIGLFGEQVPQETLAAAEVPQLPVLQPSNATLYRPTKPHMPLKKPRIEISDDEEFFTVPDLG

>NNU_23353

MKIQCDVCNQDEASLFCSADEAALCDGCDRRVHHANKLAGKHLRFSLLHPSFKETPRCDVCQERRAFLFCQEDRAILCRECDLPIHTANEQTKKHNRFLLTGVKLSPSSSLYSTTNSSINDVCDSVKDKLNNSKLYLKKKSVPLCSGAVVHPTPPVVEKPSPPPPPPPPTTTATITTHKSDDHPASGGGSTSSISEYLIKMLPGWQVEDFLDSYSAPHGFCKGDDLSPFQIQDMDLENSWSFAAQDITICVPQVPQASTTPYCLPNEINFENGFKEASNQKTAANKRWRDDGGFTVPQISPPSCKRSTHFW

>NNU_24534

MKIQCDVCEKVEAAVLCCADEAALCWGCDEKVHAANKLAGKHQRLPLLNPSSHVPSCDICQEKVGYFFCLEDRALLCRQCDVSIHTVSPYVSSHQRFLVTGVKVALQHSSNSDNINSNGNSSNGGNYHLSPPTSNASSADLPTNKKVRLSLTRSVSTSASTPASLSDEPTLAIRPNWPLDEILNSSDYDCCYGFSDLGSSRIGSQ

>NNU_26667

MKIQCNACEAAEASVLCCADDAALCWACDEKVHAANKLASKHQRVPLSNSSSQMPKCDICQETVGYFFCLEDRALLCRKCDVAIHTANNYVSGHQRFLLTGVKVGLESEEPVASSTKEKLNPAGRDVGVVSRNPGTECRPEPMSAGVVPRNERIISRLEPSSAGMVSRNIGTISRPEPRRHISPQMAGENSGSLSSLVCGVGSLPTSTMSFSGGPMSGSSPQWPLDEFFGFTELSQNYGFMDHTSSKADSGRQEDSDWSPILRAADEELDVDECLGQVPEMSWMVPQMPSPPTASGLNWPKNLRNPPSDCAASVPDICSLSVQNLYQYQPNAAILK

>NNU_16181

MKECELCNHPARMFCDSDQASLCWDCDAKVHGANFLVARHSRSLLCHVCQSPTPWKASGAKLDGGDDDDDDLDSDGEDDDDGENQVVPWSSTPPPPVASSSSSDESSSSGGRGGDESAMPYSFKRMRENADLWYQDDLGCSSSHLNNDNLGAAADRNLADGEATSYRALLMPWRDRKRNIGESTSEALASSLKRFHRDKLASKDNSSVIFGFTRAVDLVSSSAQ

>NNU_18015

MKESVCELCSGEASLYCSSDSAFLCWNCDARVHEANFLVARHVRRTVCIKCKGFDGKRVSGVGFRPVQSVCRSCSPETGDEDPEYSSSSSSSACISSSESRAVAPPKKIDFDRRRQDKIGCLSSVTEVSGDDLSSKKTMKKRTKVPRSRAPASLDVNAEGILVNWCRKLGLKNICSVRLASHAFGICLRKLTILPFRASLAASLWFSLKLCEGRSASTCQNLKRLEEISGVPAKLILLGEAKLSRVLKIKRSQQHDLEEGWAECSD

>Spipo5G0007300

MGARECDFCRGSPALLFCRADAAFLCGGCDAKVHGANKLASRHARVWLCEVCEQAPAAVTCKADAAALCVSCDADIHSANPLARRHHRLPVAPFYDTPTAAALINPANPPAAGAAVAAELLLKPSSSLSDDDDDDDDDNQSEEAEAASWLLPDPEPLASEEGHRLHHLHLHHQQKVEEVEEEELKTSDYLSEEYPYLDLEYPSSVDAAMDSVVPEQMPAAGLVAGHPAAFISPDGGIDFDFGMSKPHLHGIYPSASCISHSFSSSEVGVVPDAGAGVDVTNPYGGRVDREARLMRYREKRKNRRFEKTIRYASRKAYAETRPRIKGRFAKRGEREGDVAGIYSSGNPAAATAAAFMVDPGYGVVPSL

>Spipo29G0015300

MAARECDSCNSAAAVVFCRADSAFLCAACDGEVHSANRFASRHERVKICEVCEQAAAAIFCKADAAMLCAACDADVHSANPLARRHDRQPVVPFRDSPPLAGVKLAGGESLSPPTPAPVPAIARDEEEEALSWLIPEENHKVLMDIPDLKEEMDFFFFSDADYASCMEDSVVPVQPAAVAGDPPAFTACFDQELGGSATMSSSEVGTVPEANATAEVTNPYGGAAAAALGMSREARLMRYREKRSSRRFEKTIRYASRKAYAETRPRVKGRFAKRAAAEEPEFTYQAAAGGAAGYLAADYDYAVVPSL

>Spipo28G0017700

MGEEMRVKSYCDACRGSPAQLFCRVHAAFLCAACDGHLHGGAARHERVWVCEICEQAPATVVCKADAAALCAACDSDVHSASTLSRRHQRFPVVPFSQSLAGDWSLDRSAAENSYAHSEELPRTSPELKAADCCYYYHRQVSDGDPYFALEEVVAAPQPVVFTTDDGCMDMNSGGPKPLYGGDAATPSAEESAALETTEHGGGWAAPLVARAVDREARVMRYREKRRTRRFDKIVRYASRKAYAESRPRVKGRFVKENTVIY

>Spipo10G0011300

MAGERVEGKSYWGMEARECDSCKSSPALLFCRADSAFLCGACDVRVHGANKLASRHERVWMCEVCEQAPAAVTCKADAAALCLTCDADIHSANPLARRHERVPVVPFFESPAVDKSGEETDLLLKPVGEEGNNDESEEAASWLLPNSNSFAGNHHQKVAVEAPEIKGGEYLFSEVDRYLDLECASSMDGSNHQADSVVPVQAKIAVPAVHLPPFFAPDGGLDYDFSRSKPSSGSYSTVPSLSHSMSSSDVGVVPDVSAMADITNPYGAAVVVQSTAQVAVGLDREARVMRYREKRKNRKFEKTIRYASRKAYAETRPRIKGRFAKRTEIDTEMDHIFSSPAATLVVDSGYGLVPSF

>Spipo12G0012900

MKKKLAGSGGGGRACDSCQSAPPAVYCRADSAYLCGGCDARVHAANWVASRHERVSLCDAAPVTCKEEEEDDDDDDGDDEVYDGDEDEVSSWLLLNSGKNSGQGLLFAGEDNDYLDLVGYGSSGETQQTQGGDGKKLDGREEQVLLQQQQLSFPMEMAAAYEASNGFYGYTGFFSHGDQPAMDARTVPDVSAGAVAHFRQPKGTIDLFAGAPAPPAATPFGPGDREARVLRYREKRKTRRFEKTIRYASRKAYAETRPRVKGRFAKRSDAELEVDQIFSAAAMAETGYGVVPSF

>Spipo12G0052800

MAARECDSCKESPALLFCRADSAFLCGTCDAKVHGANKFASRHERVWMCEVCEQAPAAVTCKADAAMLCVACDADIHSANPLARRHERFPVVPFYDSPAAATGTLKFTNGPDLLKPSDDDDDCRGNETEPSWVLPDPYPVKNDHLQKIAAEVPELKTVEYFFSDVDPFLDLDYASSVDPRYDQTDSVVPVQAKAAAVASPVPALIATDGLVDFDYCRPKPSCGGYSISSSDAGVVPDASAMADITYPYGRTNGPPPAGEPDREARLMRYREKRKSRRFEKTIRYASRKAYAETRPRIKGRFAKRTENGGEVIYGAAADLMVDSEYGVVPSF

>Spipo6G0043800

MASAAGSGCARGDIFYYLQSSQRQQQGSYSDESGSDGRRRCDFCEEAAALVYCRADSARLCLACDRLVHAANTVSSRHSRSLICDSCVSAPASLLRCSAASPGGQRRREGHPLLLCANCDFDAGEVGRAAERRPLEPYSGCPAAEEMISLLGVPVEEEKVLLVAGDGDGGGGGDGGGDEGLFGDDGWMWETPPILCMEDLLLPITPFHGFPAVGVPPPPKDRNPACGRHKEEILRQLRGMDLPEGFEPLEPDVAFHPPEGLEQGAMPSNVEDDDASPIVPLSGSQAEAASEDCRRESRWDAPSTEQQPETSVVEGKALFDGTIAGAGSVGGIGAERAGAAPPSPAAARYDKHIRYESRKARADGRLRIRGRFAKVGRTPAA

>Spipo4G0060000

MDGKIQQLPKRRADAVPCDFCGEETAVIYCRADSAKLCLSCDQHVHLANALSRKHLRTQICSNCGSQPVAFRCPADGLALCQDCDWDAHGSRGSSHERTPVEGFSGCPSALELAAAWGFDLAGKDSSDPHRFSGWLSGDSISDMDSALLQDLYVPCEDMASFSRAQQTPLRRRRHALALLRQLTEMTKRNAAAAPPCELGPETPVRNNRCRDAEDGADLRQMAYPSFLMLPPGGCPGHGERDQLPEEDILWDCDPLNHTSQIWDFHLGRSRDHHESLPADVSNDAAFMIKSFGDLANGSSFPTATAGAVGDVYESSCPSRPGDISSTDVSHRSGPPKDILSTGAQNLGAVQADGGWRAGSNSASNGASRSSHNRATVVGPSGASSGEPRPSCSSKDVSLGDQLPPAAGSGATLVAARIDGELLALNRDSAMLRYREKRKTRRLKRIRYESRKARADTRKRVRGRFVKSAEGVDVQSCG

>Spipo15G0031700

MTTLSEKRSANAVGGKTARACDSCLRRRARWYCAADDAFLCQSCDTLVHSANPLARRHERVRLKTASSSSPLPPFKQAALPTAPSWHQGFKRKARTPRARHTAMAPAAAKSEPLVPDFEASSLDETAEEEGEEEEEEQLLYRVPIFDPVLAEFCSSHVSDDSAATSSGEAKPAMNPSPKYASSPGGLVNALPGFSPSDVDMGEFSADVESLLGRVLDEDAFSMEGLGLMEAGGDGADCCFAGGGYVKLETSDGDDLNGDTLDLNFECGSPAPAPAKLAAPPAMGASVKRRRIALRLDYEGVIDAWSSTQGCSSPWTDGERPQVNPDECLPDFMGMWGEAPGLAYGGEMSGGGDGGPGGSGMGGAQGAAGDGGREARVSRYREKRRTRLFSKKIRYEVRKLNAEKRPRMKGRFVKSASFGGSAGAPAGVAPQPAFAF

>Spipo8G0051900

MKIQCNVCEAAEAKVLCCADEAALCLACDEKVHAANKLAGKHQRVPLTAAASSQVPKCDICQEAAGYFFCLEDRALLCQECDISIHSVNLHVSAHRRFLLTGVRVGTEPAGSTLSFAEQQLSPSLVKSLPKRSSSTTFSGENKEAVPSQVGWQGNQPDSRAPSTGGLLTGGITDWQFDEFFGADYGHNYGFAVNESSKGDSGKLGSSEGSPIYRPACEELEVDSVSQVPEVQWTVPEIPSPPTASGLNWSRNSWSHPLADHAAFVPDVCASSCQDTQRCRQGAGSMKRQRPR

>Spipo4G0071400

MKIQCDVCEKAPATVICCADEAALCAKCDLDVHAANKLASKHQRLRLDCLSTKLPRCDICQEKPAFIFCVEDRALFCRDCDESIHVPGSLSGHHQRLLATGIRVALTSACSHKDAAGKAFTEPPNQTPHPLAQKAPTGKQASSEFTPSAWAVDEFLQLSDYESSDKKEVPDCFGELDWFADIGLFHDQAPEHATSAAEVPELPISPPSHAAAYRPAKTTPFKKQRVEVLDADDYFTVPDLG

>Spipo17G0014200

MRTICDVCESAPAVLFCAADEAALCQSCDDKVHMCNKLASRHVRVRLADPSDVPLCDICENSPALSAFDVIGTAFFYCEIDGSSLCLQCDMTVHVGGKRTHARYLLLRQRAEFPGINKPDNSEDMADTVNDTKDGRLPNPMTRDALQMQHAASESNAFAHGSVGNKMIDLNAWPRQAHGQPSSTQSIRRNHIYPKMVMFWSLEGDGAFRARRRVPATAMGKAADLF

>Spipo0G0081900

MKVQCDVCERAEATVLCCADEAALCERCDEKVHAANKLAGKHQRVALLSRSSSHIPSCDICQERTGYFFCLEDRALLCRHCDVAVHVASPHSSSHQRFLITGVRVALQPVPSGGGSGGATSNGSMASDSPASNSSPEKKRFRAAAAAPPPPLELPLEVAAEGALGLQCPWAELLATGDLDQFCYALSSEPGSVADSH

>Spipo19G0001500

MKILCNACEAAESSVFCCADEAALCWACDQKVHAANKLASKHQRVPLTSSSSSLVPKCDICQDAPGYFFCLNDRALLCRACDVTIHSVNSLVSAHQRFLVTGVRVGIEPNGCPSSLAKEQETPQPPPKLASEVSLSMSLSRGGHKVVSSQVGRNESFPVSRAVVNGGSTTASTSGWPLNEFFGEDFNQDCGFQGNESSKVGPILSCHFL

>Spipo11G0023500

MKILCDVCRREEAAIFCCADEAALCGACDGRVHCANKLAGKHRRLSLQHPASPQQSPLCDICQEKRAFLFCQEDRAVLCRDCDSSIHSANDLTRKHSRFLLTGIKLSVSDAPTGLLTNAEVKDKNNSVNEPAAEDSDGNTGSTASATGGSRISEYLMKELPGWHVEDLLLETPSTGYGLPKMLGDDDLASFPVENLNAAAAAAAADGGLFSALWVSDDCFTVPEIGAAHTRRRRSDPPLHKRPRVAAAATGYDLPPLPSVFFR

>Spipo12G0020000

MGQSCDFCGDQRPMVYCRSDAACLCLSCDRNVHSANALSKRHSRTLLCDGCSSQPAIVRCIDEKASLCHNCDWNRHGRSCPTSGHKRQAISCYSGCPSASELSKNWSFLLDIPPTEDSHCEQGLGLMRIEEESVSKFMCPPGNSSTGEQASASNMDDIGNADRLNSWSGPSAVDAFPSSGNQPAGSVDSTEPKVLCDMGTDGFGMCENDEFYDEFSVEDVDLNFDNYNEFFGTSRNHSEHLFDDAEIDSMFDLKNPSSANENHLDEFPAEACPADQDGKIQPASSNVVSTDSVKTNPKNRSPKLCFPARHGHSSLSLSFSGLTGESSAGDYQDCGVSPTLLMGEPPWCPSGPGNPAFTTANRETAVKRYQEKKKARKFEKKIRYESRKARADVRRRVKGRFVKAGEAYDYDPLCQTRSF

>Spipo14G0056600annotation

MKQCELCDREARMYCEADQASLCWECDGKVHGANFLVARHSRSLLCSSGVPPMAATLVSVLSACAQLGALEQGRWVHRHLRTIGGGGATINVFLGTALIDMYAKCGEVDAAVAVFEAMPERNLLSWTTLIGGLAVHGRGTAALRLFEQMELSGVAPDDVACVAALSACSHAGLVDEGRRIFAAMRQRYGVEPKLEHYGCLVDMLARGGRLAEAMAAVEDMPMSPDGRIWSALMAGCRFHGDAKLAERVAERL

>Spipo5G0002600

MKKTKIGCELCDGEAALRCESDSAFLCWGCDARVHGANFLVSRHPRHLLCAECGAAEVQQQPVSGSGFRPLRFLCRSCSDDGSGGALSSDSESSSCCSSTRHASGSSCVSSAESGAAAGHQDGGGGITSISPPVGASSSIKFARSVRSVAGSSPRREKRSRRIGGGRRELNARAESALLSWCRRVGLDDPCAAAAAAAHALAVGMQRMPSAPFRALGAAALWFAVKTTTSRAAAAPLLRQVEECSGVPAKLILAAEARLSRAIRRCHVGEEGEGWAECS

>Spipo12G0060000

MEARRACELCEGEAAVHCESDSAFLCWACDAAVHGANFLVARHRRRSRCPECGVFDAVQGPLSGATLLPVLLLCRSCRSLEQDDASLSDSDSQSSARSSSSSLCSSTSSSPVSSAESGAAAEWAKPGRRRRRAAGGELDAGIDLVLANWCRRLGLEGGGAAAASHALLVGTRRMPALHLRVSMAAAVWFAAKLRFSTGGPRRRATALLRLLEECSGVPAKLISIGEAKLSRALRRCHAAEDGEGWAECS

>Sobic.010G214000

MELHKYWGVGGRRCGSCEAAPAAVHCRTCVGGSSSFLCTTCDARPAHARLAHERVWVCEVCELAPAAVTCKADAAVLCAACDADIHDANPLARRHARVPVAPIGSEAAAAAVEAMLFGTGDAAEADDQHNNAAAAAEQHQHQHHAHHAHALNLNVEAKDMKLDYLFSELDPYLSVEIPRFHHADSVVPNGAGAAGAVELDFTCGIGVKHSSYSSYTATSLDLAHSGSSSEVGVVPEAFGGGGGGGGGSFELDFTRPKPQAYMPYTATPQSHSVSSVDVEVVPERGDLPAVRPVPLMGESREARLMRYREKRKNRRFEKTIRYASRKAYAETRPRIKGRFAKRADHDGDGDADDAEAEAAVPSSYVLDFGYGVVPSF

>Sobic.010G115800

MNYNFSSNALDEEEVAGRGGEGGSCAAAPAWARPCDGCRAAPSVVYCHADAAYLCASCDVRVHAANRVASRHERVRVCEACERAPAVLACRADAAALCVVCDAQVHSANPLAGRHQRVPVLPLPVAAIPAASVLAEAAATAVAVGDKQEEEVDSWLLLTNTKDPVSDNNNCNCSSSSNNNISSSNTSTFYADVDEYFDLVGYNSYCDNHINSNPKQYGMQERQQQQQLLLQKEFGDKEGSEHVVPASQVAMANEQQQSGYGVIGVEQAASMTAAVSAYTDSITNSISFSSSMEVGIVPDNMATTTDMPNSGILLTPAEAISLFSSGSSLQMPLHLTSMDREARVLRYKEKKKSRKFAKTIRYATRKTYAEARPRIKGRFAKRSSDMEIEVDQMFSSAALSSDGSYGTVLWF

>Sobic.004G211200

MDTAVELELEQKPAVGYWSVVGARPCDACAAEPARLHCREDGAFLCPGCDARAHGAGSRHARVWLCEVCEHAPAAVTCRADAAALCAACDADIHSANPLARRHERLPVAPFFGALADAPQPFPSPAFAAAAAAGGQAQGEAAAADNDDDDGSNEAEAASWLLAEPDNSHEDSAAATAADTLFAESDAYLGVDLDFARCMDGVKAIGVPVAPPELDIAAGSFFYPEHSMNHSLSSSEVAVVPDAQAAGVPAVVSRGKEREARLMRYREKRKNRRFDKTIRYASRKAYAETRPRIKGRFAKRCSAEADDDALEHDEGACFSPAGSAHAASDGVVPSFC

>Sobic.004G063200

MEALVAGRYWGVGGRRCEACGGSPAAVHCRTCPGGGAYLCAGCDAGHARAGHERVWVCEVCERAPAAVTCRADAAALCAACDADIHDANPLARRHERVPVQPIGAAAAAPAAETLLFGAAAEENQDDDDGAAAAAKVVGVDAGKLADFLFADVMDPFFGQDFTGGTRFPHADSVVPNKGSCGGGGAVDLDFGGGVAAAAVAAKPSYSSYTAASLGHSGSSSEVGLVPDAMCGRGGSVTGGVIELDFAQSKAAYLPYAATPTHSMSSLDVGAVPERGDGVMAGRVATPPAAAAAESREARLMRYREKRKNRRFEKTIRYASRKAYAESRPRIKGRFAKRADDNDADADADFDFDAGAAAATAPARSRSQQQQPSYPYVLDFAAGYGVVPTF

>Sobic.004G252300

MRYQKNGRRYEALGRSSPTARPCDGCHAVQSVVYCHSDAKCLCVSRDKQVHSANQVAERAHVCEVCKSASTVLTCCADAPALCTTCDAKLHSANTLSQRHQRVPVLPLPAAAIQTTSSFDEGKAFVITHGIKEEEEEVDSWLLLTEDSDYSNCTNSTATANNNRNKKMGFGDVDQYFDLSGYNPYYHSNITRNPEEQYMQEQQQIQRRYLEKEWNECAVPSQLTMVYEQQQSVYGIGGAKNAVSVTSSISLSSMEAGIVPDNTIAGISNLNILTTGGVDLLPVRSFQMPVHLSPRDRAARILRYKEKRQARNFNKTIRYATRKAYAQARPRIKGRFTKISDVELKVDLMSSPPDLPNSSYGTVPWF

>Sobic.006G135100

MEGDEKSAGGAPAYWGLGARPCDACGAEAARLYCRADAAFLCAGCDARAHGAGSRHARVWLCEVCEHAPAAVTCRADAAALCASCDADIHSANPLARRHERLPVAPFFGALADAPKPFASSAAAVPPKATAGADDDGSSEAEAASWLLPEPDHGHKEEGATTEVFFADSDPYLDLDFARSMDDIKTIGVQGGPPELDLNGAKLFYSDHSMNHSVSSSEAAVVPDAAAGAAPVVAVVSRGLEREARLMRYREKRKSRRFEKTIRYASRKAYAETRPRIKGRFAKRTPGAGEDPLEEHEEMYSSAAAAVAALMAPGGADADYGVDGVVPTY

>Sobic.002G408500

MSSAAADAASGKEAPACESCTSLPAVVYCRADSARLCLPCDRHVHGANAVSTRHVRAPLCSGCRATATVTAGGGTFLCANCHFGSEEEEGRHRDGDDPQPLHHDRAAVEGYVGCPSIAELAAILGVAGYDEKAAAAGNGGWWPASAWEDPQVLRLEDVIVPTTSCHGLQPLLTPPSPENRSSGGEMADEVVRQLGELAKLEATVAAAYAEMEPADGEQLPPWASPELAIGHADFGALDAGAAWHDAATIAAVPSTEEQEAWIAAGCDVDAAGRTDEEAREHAALAPAPAEPCLSSFVEMSEICPGSVVTLSHGGVGGGGTADVDNSGKTDAETAPRPQLAPTAPVLVAVPVPVTEKMGGYDVAYPDRGTVISRYKEKRKNRRFDKQIRYESRKARADGRMRIKGRFAKSGGEV

>Sobic.001G372700

MGEDDDDQRNQMLGAGLDHEPERRPGEAEPEEGKKPAASEAEAGGDGAGTEAATCDYCGTAAAAVYCRADSARLCLPCDRLVHGANGVCSRHARAPLCADCRAAGAVFRRASSSAFLCSNCDFGRHRDGGDPPLHDRCAVQPYSGCPPASDLAALLGVPLFDKPATEDGGAWWNIWEEPQVLSLEDLIVPTTPCHGFEPLLTPSSPKNRSISPDGKMNEEILRQLGELAESDGGVQASAGREEAEQAGGDQFPSWASPQYATGHGNFGTENNHEVATMPTPLYENGRWNNCDLDALNDACKVEVAYDQVPVSSAEPCLSSFAPLSEICPSISNGNSMEDNHQVNPGIGMPMQGLPKRTGFDVVPCPDRDSVISRYKAKRKTRRFDRQVRYESRKVRADGRLRIKGRFAKANQT

>Sobic.007G189800

MKSGGGGGGGGGGGGQQWPCDYCGEAAAALHCRADAARLCVACDRHVHAANALSRKHVRVPLCAGCAARPAAARVSPVPGADPAFLCAGCCDDAASAAVRVPVEGFSGCPSAAELAASWGLDLRRAEEGKDGAGGDIDDGDPFLSVLDYSVLGVAVDPDLRDLYVPCDPPRVPAPDAAGARPLRGQALCDQLAEMARRETDTAHAHPHSDLSPRTPRRTSAASGGRLPPGKMSPPAAMPTHHPPPAAVQEVPLPYTSLLMMASANCADLIGGADRVGDDDEQLLWDCAAPSVPPTQIWDFNLGRSRDHDEKSALEVGYGSNHGGFMIKSYSDMLKEISSGTTKDLEDIYDSRYCSTAEDIMSSNICQLSSKNVSTASNKRKLSSCASTIDGPTTSGNHVPTSGPALTREISFGDQTVSTPAAERPAVRIDSETLAQNRDSAMQRYREKKKNRRYEKHIRYESRKLRADTRKRVKGRFVKSTEALNAGYGG

>Sobic.010G108500

MIATTGSSAKTAAAAAVGGKSARACDGCLRRRARWYCAADDAFLCQGCDTSVHSANPLARRHERLRLQPAAASSSPLHTPPRTGAAANNKRERHDEVVPAWFRRKARTPRGGHAKSVGGQALSRSRRLGVVVPHAAAGGGDSPDDGRSAEGEFEAEEEQLLYRVPIFDPALAEFCSPPPAPLEDAAALASSCNEDGAVEDPANSKPDPGPATPAPAPVVQFFPDSGHANFEPTDAELREFAADMEALLGHGLDDGNEEDSSFYMETLGLLDPVEVGDDATRVKVETDGGSACGEASGTLACALELLDPAEVSDEMLDIDFNYGSPLDTMMDDEKAASSDTGGADDAQFLQTSLSLTLNYEAIIQSWGSSPWTAGGERPHVKLDDSWPHTNMWVVGGVAGHGGEDLLLGTARLGMDGGREARVSRYREKRRTRLFSKKIRYEVRKLNAEKRPRMKGRFVKRATAGGSSLAIAGLA

>Sobic.001G118100

MTTSAGAAAGAALGARTARSCDGCMRRRARWHCPADDAFLCQTCDVSVHSANPLARRHHRVRLPSASCSSPPCDPDAPTWLHGLKRRPRTPRSKPGGGKHEATTPNSMAAAASAAVPDLEAEESGSGIVGDNDDHGFLVDDDEDLLYRVPVFDPMLAEFYNPVADEGEQKPLAEFYNLVADEGEQKPACLMPPLVETSPEFASGGLAEADGLSGFDVPDMELASFAADMESLLMGVHDGFDDLGFLDEEKPQVNADAYLEAMAAPVPEREDKKRKRPEMILKLNYDGVIASWVRDGGSPWFHGERPHLDPYELWSDFPAGSRGLLGGAVTAVTGGEREARVSRYREKRRTRLFAKKIRYEVRKVNAEKRPRMKGRFVKRTTLPPLPRPPPQQQQKQLPRALPHVGMVLAPPPGANGRFQF

>Sobic.004G249500

MSSSKHAAAGAGAVGGKAARACDSCLRRRARWYCAADDAFLCQGCDASVHSANPLARRHERLRLRPMTSPPDPAHSTLEAGGVGVASTSTWKKRQQQQQQVAPAWSKRKARTRRPHVKSVGQLLSRKLVVVPEVATVESSEERKVEEEDEEEEEEEEQLLYCVPTFDRALAELCTPPPPLDDPTATASSSCCRDNDVDGAVDNAKAAPPAVVVAESPVQQLPDSFAGFGPTDAELREFAADMEALLGQGLGDSNELDESFYMESLGLMTTTQQAEDVDVGRVKMEPNGSVISRSRGEGAPGFGPAELMKPEASSAEVLVLDIDFNCSSPTVMMDHEDEDSFEHKASASNGDAAAAGTQFLKRSLDLSLNYEAIIESWGSSPWTDGQRPNVQLDDFWPHAHLTGWMAGGGRLGGEAAAVSPRLGMVGGREARVTRYREKRRTRLFAKKIRYEVRKLNAEKRPRMKGRFVKRPAAAGGGGAAIAAPCAVT

>Sobic.010G041700

MKIQCNACGAAEARVLCCADEAALCVACDEEVHAANKLAGKHQRVPLLTDADAAGTAAAAPAVPKCDICQEASGYFFCLEDRALLCRDCDVAIHTVNSFVSVHQRFLLTGVQVGLDPADPVPPIAEKHVNAAGGSVNQPVKHLPRRSPTVQFSVEGSASVPSKNVTNGDYSRQNSVPTARAEVVDWTMNNSTIRSVESPPKYMSEESPTLLQSSQTTTAFNQINGNSDGPYHLSFSGGNVTDSLPDWPVEEFFSNSEYGPNFGFSEHGSSKGDNAKLGNAGGSPQCRLAEGSVAEELLGQVPGLITDEYMSRVPENSWTVPEVPSPPTASGLNWHGNLCFPAYDSTMFVPEITSLQTSQNQFAVPSSFKRRRREY

>Sobic.010G262200

MQVRCDFCGAAPAAVLCWADEAALCSACDRRVHRANKLVHKHRRIPLVQPASGNVSDADADAAAPLCDVCKERRGLVFCVEDRAILCPDCDDPIHSANDLTAKHTRFLLVGAKLSAELVDQAPASPDDDDDDDDACGRDTRAAAEPDAVPALGAQGSCAAKASALESGSVGGGSSISDYLTNICPGWRVDDLLFDDPAFSAASQKASGYSDDGHEQVPSLDADLFDVVAGGRPGKRGGVWSTGAGALGFDKATPASVVAVPTQGFVREMSWNSDSDSDVFAVPEFPHPPPAKKARPAPASTFWCF

>Sobic.002G273300

MRTICDVCESAPAVLFCAADEAALCRSCDEKVHMCNKLASRHVRVGLADPNKLARCDICENSPAFFYCEIDGTSLCLSCDMTVHVGGKRTHGRYLLLRQRVEFPGDKPGHMDDVPMETVPMETKDPENQRDQKKAPKEQMANHHNGDHPACDGNCDDQGNIDSKMIDLNMRPVRTHGQGSNSQTQGVDLSVNNHDSPGVVPTSNSERDASK

>Sobic.004G301000

MKVQCDVCAAEAASVFCCADEAALCDACDRRVHRANKLAGKHRRFSLLNPAPPSSSGSGSPAQQQAQPPLCDICQEKRGLLFCKEDRAILCRDCDVSVHTASELTMRHTRFLLTGVRLSAEPAACPAPPPPPSGSEDENSSGSGSFCCSAGGDASAAPPPSSAAPATSHGSGSDNGSSISEYLIKTLPGWHVEDFLVDEAAAGAATNIAGVSADASYQGGLARIGGLQDGYGYSAWMAPEQLFYEDSSAAGGARGIREQWVPQMAMYSSSTGLSVAGAGSKRSRATSAASSYSYW

>Sobic.004G208400

MKIQCDACEGAAATVVCCADEAALCARCDVEIHAANKLASKHQRLPLEALSARLPRCDVCQEKAAFIFCVEDRALFCRDCDEPIHVPGTLSGNHQRYLATGIRVGFASASACSSDGACDAHDSDHHAPPKATVETPQAQAAVSAAAAAQQVPSPPQFLPQGWAVDDLLQFSDYESSDKLHKESPLGFKELEWFADIDLFHEQQAPKAGRTLAEVPELFGSQAANDAAYYRPAKAAAGAGVRQSKKARIEVTDDEDYLIVPDLG

>Sobic.003G026700

MKVLCSACEAAEARLLCCADEAALCARCDRDVHAANRLAGKHHRLPLIPHADVSAPNCDICQEAHAYFFCVEDRALLCRACDVAVHTANAFVSAHRRFLLTGVQVGLQPDAAADADDPNPPTAAAASDPLQTPPPPDRKAAAAGGGSPAPLYSDDDIDWAPGADAGAGVGLPDWALVHEQFSAPPVTRPADPALARTPASKRSPRRSLAAAFTVQSGGGLAGGLPDWPLDEFFGFSEYSAGLGFAENGTSKADSGKLGSTDGSPAGRSSSDASQDFFGQVPEFHQWSVPELPSPPTASGLHWQGGPRHGATTTTDTNTAAVSVPDISSPENPFRCYAATAAGQPPAKRRRRC

>Sobic.006G131800

MRIQCDACEAAAATVVCCADEAALCARCDVEIHAANKLASKHQRLPLALGDATAASASSLPRCDVCQEKPAFIFCVEDRALFCRDCDEPIHVPGTLSGNHQRYLATGIRVGFSSVCGAGAGAEGLPPPAPPKGSSKPAAVVSAPAAGATKTTTTVKDTLPQEVPSSPFLPPSGWAVEDLLQLSDYESSDKKDSPLGFKELEWFADIDLFHAHSPAKTTTAEVPELFASPQPASNAGFYKTNGVARQSKKPRMEVPEDDEDYFIVPDLG

>Sobic.006G163100

MKVQCDVCAAEAASVFCCADEAALCDACDRRVHRANKLAGKHRRFSLLHPCSSSSSAAAQKPPLCDICQERRGFLFCKEDRAILCRECDAPVHSASDMTRRHSRFLLTGVRLSSAPVDSAAGPSEEEGEEEENSSSPCNDDSCSGGAGGAGATTTPSASDGSSISEYLTKTLPGWHVEDFLVDDASAGDVGAACSDGLYQGQRGHISGVLQEEAYTPWTGREQVLGDVADERASWELWVPQMHAEFAGDSKRPRPSPSPPCSYW

>Sobic.008G073400

MKIGCDACERAEAAVLCCADEAALCRSCDAAVHSANKLAARHHRVALLPSSTAHPPSSTSPIADDGSGSGGGGGDGHPACDICQEKTGYFFCLEDRALLCRPCDVAVHAAGVHVSSHRRFLITGVRVGDVESLSHGVPGSDGGASPSTTSSGNGSSNAPGSSSGGGNPTTTTTTTTMPDQVRPSSSSSIRATAATTAEGSPGQWQQWLWSDFLADDVGGGGGVDMEEECCHAELSEPGSSGLTRS

>Sobic.009G075600

MKVLCSACEAAEASVLCCADDAALCARCDREVHAANRLAGKHQRLPLLAPGGQSAAAVSPPKCDICQECDAYFFCLEDRALLCRSCDVAVHTANAFVSAHRRFLLTGVQVGQELESDDLSREQQPEASPPPPPSKSEPAPPPPPPLYNESDFGWAAGAGATGSLADWSAVEEEFGSPAPCLAEAAPRATPKRSPRAPPAFGAAGQGRVAGGVMDWPLGEFFRGVSDFNGGGFSFGESGTSKADSSGKLGGSAGGSPYYRSSSEDRDAANELFGQVPEIQWSVPALPSPPTASGLHWQHGGHDSNAFVPDICSPDGGGAGVRCFPTANGAAKRQRNR

>Sobic.010G123500annotation

MGALCDFCGEQRSMVYCRSDAASLCLSCDRNVHSANALSRRHTRTLLCDRCASQPAMVRCLAENASLCQNCDWNGHIAGSSAAGHKRQTINCYSGCPSSAELSRIWSFVSDIPNVAPEPNCEQGISMMSISDSGVSSQDNAAGDNNLLDIASETLISDLGTCDKPLVGSSSGAGVNLLPLATDQTAGSVDSPPDKVPYTPDKDMFSKDSIYEDFCVDDVDLAFENYEELFGTSHIQTEQLFDDAGIDSYFEVKEAPAGNSTEQSKLKQPANSNAVSADSGMSNPGVKGDSSVCIPLRQARSSLSLSFSGLTGESSAGDHQDCVVSSLLLMGEPPWQPPGPEGSIAGGSRDSAITRYKEKKKRRKFDKKIRYASRKARADVRKRVKGRFVKVGEAYDYDPLCQTRSY

>Sobic.004G256200

MASLCDFCGKQRSMIYCRSDAASLCLSCDRNVHSANALSRRHTRTLLCDRCGSQPASVRCLEDNASLCQNCDWNGHDAASGASGHKRQAINCYSGCPSSAELSRIWSFIMDIPTVPAEPNCEDGLSMMTIDDSDVTNHHDASDDKRLLEIANTTLMSDPPSADKPKPLISSSSGDGFDVLPLATDQPAGSVSVTPKVPYARDDDNFNDGMYEDLCVDDADMTFENYEELFGTSHIRTEELFDDAGIDSYFEMKETQPFDFNEEPKTMQLECSNVVSADCGMLNPGARADSSLCIPVRQVRSSISHSLSGLTGESSAGDHQDCGVSPMLLMGEPPWHSPGPEGSVAGGSRDSALTRYKEKKKRRKFDKKIRYASRKARADVRKRVKGRFIKAGEAYDYDPLSQTRSY

>Sobic.010G089850

MVGGLRLCDICDDPASCFCPADDAFLCDDCDKQIHEANFLAKKHNRVSVCHLNTPCSRRLELPNFDPGNSEIGSVLNHGSRITPIAPGDKTDPTQMEVEKERGQTNKKAGTGGSGESDRSSRVFRGKGFVGGRGEGHQGGRVQT

>Sobic.010G195901

MVAMAGEAKLCDVCDDPAKFFCPVDSAFLCGDCDVQIHEANLIACKHQRLMVGLMEDSQRTDGGVIALNCWLDSKGVMLNSIRGSSMAMGVASTPGGLGGGIREKQNQGKVVDPTHGESEEWPRLGAKTESNLSIHIQTFYNRGKTES

>Sobic.010G262550

MVGGLRLCDICDDPASCFCPADDAFLCDDCDKHVHEANFLAKKHNRISTCQLNTPCSRRLVLPNFAPGDLEIRSVLNQGSRTPIVPERMC

>Sobic.007G062100

MSSAAAGEGGKERGGAGAGPGGACELCGAAARVYCGADEATLCWGCDAQVHGANFLVARHARALLCRGCARPTPWRAAGPRLGPTASLCDRCVRRGPGAVGVGGGDEEMGGAGDGRGHEEEDHDDDGGDDDDEVVVEDDEDEEEEEEEGEGENQVVPWTEEAEATPPPVASSTSSSSREAPANGASAADCAKENMPCSTSQPGLCHHLSSAHHGGRSDEATSSRNGGRFLASSRHRKRSPSDFFSSGSAQSGSGTPARNCSNAGIGRNDFT

>PGSC0003DMG401010056

MLKKENSGGFDGSSNYWARVCDSCRLVTCTIYCQADSAYLCAGCDARIHAASLVASRHKRVWVCEACECAPAAFLCKADAASLCASCDADIHSANPLAHRHHRIPIIPIPGTLYGPPAVDTVGGDSMMIGGSTGEGTEDDGFLSLTQDADDTTIDEEDEDEAASWLLLNHPVKDNNKNNVHNNNNQTNIYGMLFAGEVVDDYLDLAEYGGDSQFNDQYNVNQQQQHYSVPQKSYGGDSVVPVQDGQGKSLFFYHHQQQSHHLNFQLGMDYDNSYTRLGYPASMSHSVSVSSMDVSVVPESALSETSNSHSRPQKGTIDLFSGPPIQIPPQLTPMDREARVLRYREKKKNRKFEKTIRYASRKAYAETRPRIKGRFAKRTDVEAEVYQMFSTQLMADSSYRIVPSF

>PGSC0003DMG402010056

MLKKEKSGGFDGSSNNWARVCDSCHSATCTVYCRADSAYLCADCDARIHAASLMASRHERVWVCEACERAPAAFLCKADAASLCASCDVDIHSANPLARRHHRVPIMPIPGTLYGPPAVHTVSGGSMMIGGTTGEGTEDDGFLSLTQDADDTTIDEEDENEAASWLLLNPPVKNNNKNNINNNNNNQNNNYGMLFGGEVVDEYLDLAEYGGDSQFNDQYSVNQQQQHYSVPQKSYVEDSVVPVQNGQRKSLILYHQPQQQQQQQQQSHHLNFQLGMEYDNSNTGYGYPASLSHSVSISSMDVSVVPESALSETSNSHPRPPKGTIDLFSGPPIQIPPQLTPMDREARVLRYREKKKNRKFEKTIRYASRKAYAETRPRIKGRFAKRTDVKAEVDQMFSTQLMTDSSYGIVPSF

>PGSC0003DMG400027475

MGIIREAPNCFPGGWNTGAAAQMAKSCEYCHLAAALVFCRTDNTFVCLSCDTRLHAHHERVWVCEVCEQAAASVTCRADAAALCVACDRDIHSANPLAQRHERVPVVPFYDPVESVVKSTAATLLVSITDTTTTTMTTTTGIAPALSKVDACIGHHDNNNDPWIPPNTITSKLPLTTEMKAMDFIFTDSENFLDFDYPVSVDTQSQPHYNSANDSVVPVQTNTAIKSLPFHHQEKHFEIDFTQSHIKSYNTPSLSVSSSSLDVGIVPDGSSISEISYPYMRTMNNSNSSIDLSNSANHQGEKLLGLDREARVLRYREKKKNRKFEKTIRYASRKAYAETRPRIKGRFAKRTDGGAGEFDDVDGIFSGAEFIAADSRYGVVPSFLT

>PGSC0003DMG400026311

MVAESWSTTAKRCDACKATPSTVFCKADMAFLCLTCDSKIHAANKLASRHARVWVCEVCEHAPASVTCKADAAALCVTCDQDIHAANPLARRHERIPVVPFYDSASARSPRGAGADENDPHQHEDETEEEEAEAESWLLQTPNSNTQGIEYKSAEYLFSDVDPYVEMDIIADQKTCTTTMDIAHNQEEYKEDCVVPHVQNNKNEIQLQGPVVDGYPTYEMDFSGGSKPFMYNFTTQSISQSVSSSSMEVGVVPDHNTMTDVSNTFVRNSAIDGLPNPVSSLDREARVLRYREKRKNRKFEKTIRYASRKAYAETRPRIKGRFAKRTENDMGDSLVASDASYGVVPSF

>PGSC0003DMG400029365

MGTENWSLTAKLCDSCKTTPATVFCRADSAFLCLGCDCKIHAANKLASRHARVWVCEVCEQAPAIVTCKADAAALCVTCDHDIHSANPLARRHERFPVVPFYDSAVAKSDGGGDADADAADDEKYFDSTSENPSQPEEEAEAASWILPTPKEGTDQYKSADYLFNDMDSYLDIDLMSCEQKPHIIHHQHGHYSSDGVVPVQNNNNNNETSTHLPGPVVDGFPTYEIDFAGSKPYMYNFTSQSISQSVSSSSIDVGVVPDHSAMTDVSNTFVMNSSAGTGTGTDAVPNAVSGLDREARVMRYREKRKNRKFEKTIRYASRKAYAETRPRIKGRFAKRTEIEIDSLIAADASYGVVPSF

>PGSC0003DMG400005997

MEMEKACEFCMLLKPVVYCEADAAHLCLSCDAKVHSANALSNRHPRTLVCECCGHNPAYIRCSDHQSFMCRDCDRCHHDLSSQHQRKVITSYMGSPSAKDLAALWGFGLKDLENATPPDQFISTSNGKANGAKVTPKKFKRSHSSPGGSSLASELDFTGLVVSAESEVGSASYYTKVLSLRSRRENTSLILQQILDLERLQLTEGSNNLTSGESRNNVSSLKNCTSWNVHNKFDCLQSSLDLGPELQDWGSTHESPVAESFPLPLPDGDSFWECKSPVQSSQLWPQNLQDLGAYAELECFDDSNMPDVDLTFQNFEELFGEEQDLNNTLLEEDTTCSSMENDSSIDRSDYSYVKKDISIASSVRTGHSTHFGQGHGEHIPTIKDCPPPIRTNFSSLSFSASRLSSESSGNEYVDSPAANDQEVSCNSQMDSKENLLAMQKEKKKARLYEKQARNTPRRARTNLKKQPIRGQVLKAHCYESDALNMSRSF

>PGSC0003DMG400001263

MDPVCDLCSEARAVVYCKSDSARLCLQCDDYVHSPNLISRRHSRSFICDKCNSQPSVVRCMDEAISLCERCDWDGNGCIGTGHRLKKLNPYTGCPSPDEFTKMLSQVLEMPIGTDTNFGSFGNSFCSSLSINENNSSLETKVNEGSFVPSKLNELASNYKFEAAWAIPLEPNYLTSYHPDLTPFSEGSALSKKDCPIKDLGLQEGDVLSKGVDFDDVTLDFNCGYEILPNSLPTGFSDENKELDCLVMEKNSSVTGSNNVETSHEATSSVQQEYMGLQSSQMSAAASSTNLLQTMSANANCMLMNSNCNGSIGALPFLPAPIHPSMSLSLSNITGESCAEQDCGLSPGFLNEAPWDLSLENCPQKRHEAKMRYNEKKKTRTFGKQIRYASRKARADTRRRVKGRFVKAGEAYDYDPSETRDF

>PGSC0003DMG400005325

MGHICEFCGEQRSIVYCRSDAACLCLSCDRNVHSANALSQRHSRTLLCERCNSQPAIVRRVEEKVSLCKNCDSIGHAGSGTGSMHNRQALSSYTGCPSAAELSTIWSFLLDNPLGGDSTCEKGMGSMSITDNRPTDSRAPQGTFSSQDASATVEVSEMHTPNKSSVLVGSSMLTLGNKLNKAEQIVGSVNFSSSKGCYSGVKGSTIYEEDPFSQDFNMDEVDLSFENYEELFSGSLDNPNQLFENDDIDGLFGTKDMSVSDSCCQDANAVEGSSIRRVRTMLPACSNAESADSLMSCKTEPTVYFARQASILSFSNLGGESNAGDYQECGASTMLLMGEPPWCHPCPESSLASSSRSDAVLRYKEKKKTRKFDKHVRYASRKARADVRRRVKGRFVKAGDAYDYDPLNETRSF

>PGSC0003DMG400025414

MSPGESRPCDFCNQQIAVLYCRADTAKLCLFCDQLVHSANALSKKHLRSQICDNCGSEPVSIRCDTDKLVLCQECDWDAHGSCAVSGAHDRSPVEGFSGCPSASDLASAWGLDIDSKKLHQQHTVLEFPSWMSKDAPPPSSVLLQDLMVPSANNSAIYSTKQTPTCGKQKQVIFKQLIELFKRDLADGVGAGAEDLVPKIPNATSDWQGNVNMLVMDGVNQKLEQQQPQNVPFTSSIMPHNPIDSDQMVERNILWRGNSFDQNTQIWDFNLGQLRSHEQSSSLEADYSESDMAYMMKSYGELIKGTSLATSKGLGLSGINCSLAHDDMTAFSNDSNNQAASQGPATSESNNLPRIKTSSDSGYVKTKCCGVSTDMNFMDQSIVVGGDNTGEETLKADMELLAKNRGNAMQRYKEKKKIRRYDKHIRYESRKARADTRKRVKGRFVKANESPDG

>PGSC0003DMG400017411

MGYICEYCGEQRSIVYCRSDSACLCLSCDRNIHSANALSQRHSRTLVCERCNSQPAIFRCVEERVSLCQNCDWMAHASSSTCLTHKRQALSCYTGCPSAVELSTIWSFLLGDPSIRDLTCEQGMGSMSITDCQPGDSQHPQVKEKSQDMSSALDEANDLHNLVKSAPFMGSSMPSLDNELHNVELLVGSTNLTWSKVRNSGTKGYDDPFSDDFNMDEVDLSIENYEELFGVSIDNRDRLFKNEDIDDFFGTKDMSTAESSFQGVNAVEGSAVGKVNTVKSACSNAASAESMMSCKTEPPTLCFAMQQSSFSFSNLTGESNAGDNQDCGASSMLSMGEPSWRPPCPESSMPSTSRSDAVLRYKEKKKKRQFDKRVRYVSRKTRAEVRKRVKGRFVKDGDAYDYDPLHQTRSY

>PGSC0003DMG400011378

MTELKKDQENHQQQQQEPQRRHLCDFCGNNTALLYCRADSAKLCFTCDREVHSTNQLFTKHTRWLLCNLCDSSPASILCCTESSVLCQNCDWESHNKQLSLHERRPLEGFSGCPSVSELLSILGFEDLGKKELLYGGDGSYGFSDWVIWDTPSVVSLDDLIANNDSEHNFQAIGVPPLPKNRNAACGKHKEEILSQLRELSKLEPNSGDDQDETVPTIGFQSMEPVQNCPLRFKGLGFMQNSDQHVVPSSEGSAFHWHGDTGEFADQGFSSSLMDCFIETKCLLPNRDSDVCDASGGANEEQSHNPPTTETFQMVPKVVHRELNSQERETAVSRYKEKKKTRRYEKHIRYESRKARAETRTRIKGRFAKMDYRDSSVHQ

>PGSC0003DMG400028818

MGYICEYCGEQRSIVYCRSDAACLCLSCDRNVHSANALSQRHSRTLICERCNSQPAVVRCVEERTSLCQNCDWSGHASSSSGSSMHKRQALSCYTGCPSAAELSNIWSFLLDDPSIGDTCEQRMGSMSINDNRPRDGQDTQGKDNSQNVCAAVEVNDMNISEKSNLLVESSMPTFDNKLHNVEPPIGSSSKGCYMGAKGSSLIEEDPYCDNLIMDAVDLSIENYEELFGDSLNYPDELFENENLDSFFGMKDIKGADYSYRGVNAAEGSSNWRVNTVQPTCSDAASADSMMSCKTDSILYFARQSSLSVSNQTGGECSAGDHQDCGVSPMLLMGEPPWCPPCTEISSTSTSRSNAVLRYKEKKKTRKFDKRVRYVSRKARADVRRRVKGRFIKAGDAYDYDPLPTRSY

>PGSC0003DMG400005633

MTSGSKTANAIGGKTARACDSCLSKRARWFCPADDAFLCQSCDVSIHSANQLASRHERVRLDTCSNKSTITKLVDKTHQPAWHQGFTRKARTPRNGKKAQIRQRKKNEENRVPEIGSDENEFENEEQLLYRVPIFDPFEAELCNVPDETGSIADLDILLNTEDACDDLNLPEFLSSDIELAEFAADVETLLGGEEEQSTRLLNADFEDNKAIKIEVEDEEMRAVVACHLDPELDMEREGLNWNFDEYYEERVEQKVMAAVTEFVASAEYSGSSTKSDEKNRLFLRLNHEAVISAWPNQSSPWTNGIRPHFNPDDYWPDFSETCGGNLGYYGGHVRSGDGGREARVLRYREKRRTRLFSKKIRYEVRKLNAEKRPRLKGRFIKRNSSSSFSVPGFPYMMNKR

>PGSC0003DMG400007749annotation

MGSEKKLANAVGAKTARACDNCIRKRARWYCAADDAFLCQSCDSSVHSANPLARRHERVRLKTSSIKSSDEFPNLGSTVSGSGSGSESVPSWHRGFTRKARTPRHGNKHAKRVKSTEEEEEDEEEMKNPIQLVVPEILSDENSHDENEEEQLLYRVPIFDPFVEDGSNYGNKYSSNKVDFNQDMNTFQGLLAPSEMELADFAADVECLLGKGLDDEESFNYMEGLGFLEKHDEKLVKVEDEGEMGFVNMVSTNNHDINQVDYSDFDMVGETFELKFDYDSQVMSNLDEDKKVEFGEINYDSGKNNNKIMLNLDYESVLKSWADKRSPWTMGERPEVDFNDCWPVCMVFKIGNIGIVNRHGGGGVDEGREARVLRYKEKRRTRLFSKKIRYEVRKLNAEKRPRMKGRFVKRTNFAPTPFPSLNK

>PGSC0003DMG400014566

MVSERKLASAMGGKTTRACDNCIKKRARWYCPADDAFLCQNCDASVHSANPLARRHERVRLKTSSLKQPSSSSSSDDYFPDLESPLSISSVSVPSWHRGFTRKVRTPRHGRKASKSAGDGDVIQRNPIHLVPEILSDENSHDENEEEQLLYRVPILDPFVAQLYSSATRNNHEMVESNATAAADADSEFKLESKEMMLQDDICNVDLNRFHEMLPSEMELAEFAADVESLLGKGLDDESFDMEGLGLLGVCNKEENSMEYSMISHEKVKIEDEGEIEVVRKTISATTHDHQYNHTHHDIDINEDTFEFKFDCDSPMNIIGEDEVIMKDENKKKILLNLDYEGVLTAWADQRSPWTNGERPELDSNDSWPHCMGNYMGIMNENVTIVDRGREARVTRYREKRRTRLFSKKIRYEVRKLNAEKRPRMKGRFVKRTNFVTTTTPNYPLVK

>PGSC0003DMG400003109

MRTLCDVCESAAAILFCAADEAALCRSCDEKVHMCNKLASRHVRVGLADPSKIQRCDICENAPAFFYCEIDGSSLCLQCDMIVHVGGKRTHGRYLLIRQRIEFPGDKLGPSNELGFPSTEQGDVRREPALPFKLPMIDNHQPNRETAMAAVENNVNNSVKMENELIDLNSRPHRMHGQTSNNQEQGMDMLGGSNHESVGVVPDGPFKREPEK

>PGSC0003DMG400030958

MKIQCDVCDKEEASVYCSADEATLCQSCDYQVHHANKLASKHLRFSLIHPSFKDSPLCDICQERRALLFCKEDRAILCKECDLPIHKANEHTKKHNRFLLSGVQLSSDVLASNSNNQNSISPTGSAASNAGTNNFKARSGNFGMKSNSISSTTESTPNYFQVDYHVQEGSVSTSSISEYLTETLPGWHVEDFLEYPSSSSYDLGYFLFCFVGDQSFCNASGIALCLLPALIIEYGIPICFKYSLPNQFFMGD

>PGSC0003DMG400003625

MRTLCDVCESAAAILFCAADEAALCRACDEKVHMCNKLASRHVRVGLAKPNEVPRCDICESSPAFFYCEVDGSSLCLQCDMMVHVGGKRTHSRYLLLRQKVEFPGDKSGPTEELARKTLDPGENKRDHSHSPKPMVKDNQQNHRGSPILISDGSADGNGKKDKMIDLNVKPNRFHGHASNQD

>PGSC0003DMG400003711

MKIQCDVCNNNEASVFCVADEAALCDSCDHRVHHANKLASKHQRFSLIQPSPKQIPVCDICQERRAFLFCQQDRAILCRECDVSIHKANEHTQKHNRFLLTGVKISANSSLYTSSESASATSCSANQDSVTNLNKSQTCTKKTLPVSGSVPQQVSVAVNIGENSYTSSISEYLEMLPGWHVEELLNSSTIPTNGFCKIGDNDVFPIWDTEIESTMNSFSPENLGIWVPQAPPPPTPQKNQNQVFPQNINFGGQIEFKNMKEVTSNKSSRKWRDDNSFAVPQIIPSSSSISFKRSRTLW

>PGSC0003DMG400027017

MKIQCDVCEKAQATVICCADEAALCAKCDIEVHAANKLASKHQRLHLQCLSNKLPPCDICQDKAAFIFCVEDRALFCKDCDEAIHSASSLAKNHQRFLATGIRVALSSSCNKEAVKNQLEPQPPQQNSQQVGLKMPTQQLSGITSPSWPVDDLLGFPDYESSDKKDLLELGEFEWLGGIDLFGEQTAAEVPELSVPQSSNTNIYRTTKYQMPYKKPRFEIPDEDEYFTVPDLG

>PGSC0003DMG400007061

MKIQCNVCEVAEANVLCCADEAALCWSCDEKVHAANKLASKHQRVPLSGSSSSMPMCDICQETVGYFFCLEDRALLCRKCDIAIHTANPHVAAHQRFLLTGVKVGLEPVDPGGISSSATSQSIQKVSEPESAPLSKRNAPVSLDAQFNKVLPTQASGVGDFAPTKSPFAGGSAAGSMPQWQFDEFIGLSDFNQNYGYMDDGSYKADNGKLGGESDSSSILRIEDEELDGDEFLGQVPDTSWAVPQVPSPPTASGLYWPKTYQNPFDSAVFVPDISYSPSSSLQQQPPIGTRLKRRRQC

>PGSC0003DMG400019025

MKIQCNVCEVAEAKVLCCADEAALCWYCDDKVHAANKLANKHQRVPLSASSSPMPKCDICQETVGFFFCLEDRALLCRKCDISIHTVNAYVSSHQRFLLTGVKVGLEPLGPSASASSGKSPSIQKVAEQESPPIPKSVAPLSSATPSEGVLPVHTSGNGNFAPSRLPMVGGSAAGIIPQWQFDQYLGMGDFNQNYGYMDYGQSKADNGKLGESASSPFLRDADAEVAGDECFSTEVADTCWAVPQIPSPPTASGLNWPTKTIQNPFDAALLESDASYFPLQNIQDQQSNGSGSKRSRHF

>PGSC0003DMG400029426

MKIQCDVCNKKEAVVFCTADEAALCDDCDHRVHHVNKLASKHQRFSLLQPSPKQAPMCDICQERRGFLFCQQDRAIMCRECDIPIHKANEHTQKHNRYLLTGIKLSANSDLYSAPSQLQSQSQSQSAISSADLCVSNLKSQNPISKSVAGSVSVSSTNTSNTKGGAVSSVVESVKVVKEKVGGCNNNVQLVNGGGNNLTSSISEYLEMLPGWHVEDFLDCSTPNVYSKNIGDEDMLSFWDTDLESQLSSFPPQNVGIWVPQAPPLPESKQETQIQFFPSQNLNFGGKIGLKESREVTNIKSSRKWTDDNSFAVPQMKPPSTSFKRSRTLW

>PGSC0003DMG400022345

MKKCELCSSIARVYCESDQASLCWDCDARVHTANFLVAKHSRILLCNSCQSLTPWSGSGSKLGPTVSVCQKCFHRNNNDGEDQDLNDHNNDDTEGEDDDEDDDDEEEDEDDGSSDGDNQVVPLSSSTTDQPNPPSLTSSSSSSEDSTSRFHRRSVIGFSPNRTEENIQSHYCGITRPEEKVERRSSWRWR

>PGSC0003DMG400026515

MEVMSSKLCELCNDQAALFCPSDSAFLCFHCDAKVHQANFLVARHLRLTLCSHCNSLTKNRFSPCSPRRPALCPSCSRNSSADSDLRSLSSSSSSTCVSSTQSSAVTQKINISFSNRKQFPEYSTNDSIGEVNSGSSNLVRSRSAKLRDPRAATCVFMHWCTKLGMNGEERVVQTACSVLAICFGRFRGLPLRVALAACFWLGLKNIEEKSKSTWQSLKKLEEISGVPAKIILATELKLRKIVKTNNRRRQGMEESWAESST

>PGSC0003DMG400026169

MCSGRREGEEETTSISYCKGPLKEGESIIRSTISCALCSSEASVYCEADNAFLCRKCDRSVHGANFLAQRHIRCLLCSVCRKTTWRFLIGTSSELILPTIAGLEQRNRRRSADSETMDYRTTPQELFLFI

>PGSC0003DMG400026181

MKNCELCKGLARMYCESDNASLCWDCDAKVHSANFLAARHSRSLLCQVCQSPTAWSAAGAKLGKTVSVCEKCVDGYYHRDEVEESESVNDEGSGTEDEETDYEEDEEDEDEIDSEGIQVVPWSNTTPPPPASSSSSENSSNGCRNVSSKRMREADPDLHSDDDSGSSSCHRKVHTSPAMDADAGDEAAAIVDSYSSKMRTPAKIQRTDRLTAPRCTSVIEFIRRNNRQRMSSGAAIAELCRLNNDSRTIDLESSETS

>PGSC0003DMG400025024

MNCELCDKNKSMMYCESDQARLCWDCDSKVHSANFLVAKHSRNLLCHSCQNSTPWTASGPKLSPTFSVCNSCLENPNTAAVRLQDPIEEIGERIEENYQTETDNDEDEDEYGSTDSEIDDDDEDEDEEDGNQVVPLSSSPLSNFSPPSPVRSVSSSGSYGDLSAAGDSGGGGGATVAVNSSPWKRVRESDCLHFEDEEAYSSSLLLEENKGTTSSIGFLRPMKVLRTNELCD

>PGSC0003DMG400013178

MCNGRREIEEEKIEQDNIIVCELCKSEAYVYCEADNAFLCRKCDKLVHTANFLAQRHIRCILCGICKRLTKRYLIGVSHEVILLKVVRWNNLDDNFDEENCSRKVKEPFLFL

>PGSC0003DMG400013753

MKNCELCNGLARIYCESDHANLCWDCDLKVHSANFLVAKHSRSLLCNVCQSPTVWSATGAKIGRTVSVCERCVNDEETDREEEEEEEEEEIDLEDIQVVPWSSTPPPQPASSSSSSDESLTPIFSLKRMRDDDDSGTSTSHREVNMLSPAAKDHGGDEPAEFVKSFSMRMSSKIQRTGEILPGRVENIGCSATMELIGRINRGEDS

>VIT_214s0083g00640.2

MLKDEGCNADAAAGGGGGWARVCDTCRSAACTIYCRADSAYLCAGCDARIHAANRVASQHERVWVCESCERAPAAFVCKADAASLCATCDADIHSANPLARRHHRVPVLPIAGCLYGPPATDPGGTVVRSAAEADNGFLGQEAEETIDEEDEDEAASWLLLNPVKNNNGSSNNQNNGLLFGGEVDEYLDLVEYNSCPENQFSDQYNQQQPPPHYSVPHKNYGGDRVVPVQCGEAKGQLHQQHQQQGFHLGMEYESSKAAYSYNPSISHSVSVSSMDVGVVPEATTMSDISISISHPRPPKGTIDLFSGPPIQMPTQLTPMDREARVLRYREKKKTRKFEKTIRYASRKAYAETRPRIKGRFAKRTDVEVEVDQMFSTTLMAESGYGIVPSF

>VIT_211s0052g01800.1

MGFQELNGKTFSAATWALAAKPCDSCKSAAALLFCRADSAFLCVGCDSKIHGANKLASRHERVWMCEVCEQAPASVTCKADAAALCVTCDRDIHSANPLARRHDRVPVVPFYDSAESLVKSTAAAVGFLVPGGAGDEEDSEAASWLLPNPKLPEGPEVKSGEVFFSDIDPFLDFDYPDAKFPHHHHHHCGGNDGVVPVQAKDPSPPVTNHPADNCFELDFSRSKLSAYNYTAQSLSQSISSSDVGVVPDGNCNSMSDTSYPSMKQVSGGGGGGSTGSQATQLSGMDREARVLRYREKRKNRKFEKTIRYASRKAYAETRPRIKGRFAKRTEMESEMVDHIYNSASAAAFMVDAGYGVVPSY

>VIT_204s0008g07340.1

MVVEVESWRMASKLCDSCKSAPPTLFCRADSAFLCVACDSKVHAANKLASRHARVWMCEVCEQAPAHVTCKADAAALCVTCDRDIHSANPLARRHERVPVVPFYDSAAAAAKSNAVNLLVDDRYYSDPDGDASREEAEAASWLLPNPNPKLAESSDLNSSHYMFSDIDPYLDLDYPSMDPKLQSQQQQQSSGTDGVVPVQNKSVQAPLVNDNCFDMDFSGSKSFYNGQSLSQSVSSSSLEVGVVPDGNAMVDVTNPFGRSMNTGSESANQTAQISSGIDREARVLRYREKRKNRKFEKTIRYASRKAYAETRPRIKGRFAKRSEIEVDYSSSGALTADSGYGVVPSF

>VIT_212s0057g01350.2

MILNFGFFVKFFQGCEDFRLNPTYSLLVNTAYTVRWRCWNWKSCIVWCSNMGYICDFCGEQRSIVYCRSDAASLCLSCDRHVHSANALSRRHSRTLLCERCNSQPAFVRCIEEKISLCQNCDWTGHGGSTTTSSHKKETINCYSGCPSSEGLSTMWPFVLDLPSTGNSTCEQGLSLMCLNETSEMNSWGPPGNSSRQDASLTVEVNDANNVDKSSILIGSSSVPELNSPSQKLDQPSGSADLTLPKLLCPGTDDLGFCEDDSLYEDFNMDEVDLNLENYEELFGVALSHSEQLFENGGIDSLFGKMDTSGADSHCQGAVIAEGSVGLANAVQPTYSNAASADSIMSSKTEPILCFTGKQAHSSLSFSGLTGESSAGDYQDCGASPTFLMGEPPWCPPGPESSLPSTSRSSAVMRYREKKKNRKFDKRVRYASRKARADVRQRVKGRFVKAGEAYDYDPISETRSF

>VIT_200s0194g00070.1

MGQLCDFCGDQRSMVYCRSDAACLCLSCDRNVHSANALSRRHSRTLLCERCNSQPATVRCVEEKISLCQNCNWIGHGSTTSASDHKRQTINCYSGCPSAAELSRIWSFVLEFLSVDDSNCEQGMVLMSLTENSVGSCSLPPVNNNINDSTVAGRTNAQENVDKSDLWMASSPVPALNPISSSVDQPTDLVNSPTTKLCCSGICDDDDLLYEDFSMADVDLSIENYEELFGVSQNHSEQLLENGGIDSLFGIGNLPGTDASARGAYVAEGSSVVHIKAMQPACSNPISADSIISSKTDPNLCFPPRQAHSSLSLSFSGLTGESSTGDYQDCGVSSMLLMGEPPWCSPCPENSLPSANRDSAVLRYKEKKKARKFEKKIRYASRKARADVRKRVKGRFVKAGDAYDYDPLNQTRSF

>VIT_207s0104g01360.1

MSDSPGDNLHHRHEEEGQKMTIQNRLCDFCGDSMALLYCRADSAKLCLSCDREVHSTNQLFTKHTRSRLCDVCDASPASILCSTDNLVLCQNCDWAKHGRSLSSAHDRRPLEGFSGQPSVTELLAFVGFEDLGKKSLFCGDESEVNEFLGCGVYESVGVDEEFSDFLVWDTPAVVNLDDLIVSTACDHNFQAMGVPPLPKNRGAPCGQHKAEIIHQLRQLAKIELSFDFDHGDAKPPIGFQSHIPKQLIQKENECNSCDHEVEFVFPTYEASAFQWCSDGSEAANQVLPSVLLGSCADEKCLVPRKHSDIGGSVSHTNGSDEGKSECPVVTKTLPALPKVSVHELNSQERDSAISRYKEKKKTRRYEKHIRYESRKARAESRIRIKGRFAKMDH

>VIT_201s0146g00360.3

MVSPKPSNGERVPCDFCSGQIAVLYCRADSAKLCLFCDQHVHSANALSRKHLRSQICDNCSSEPVSVRCSTDNMVLCQECDWDAHGSCSVSAAHDRKPVEGFSGCPSAVQLSSIWGLDIEEKKAPLPPPPSMAVDSWVYKSNTLTFQDLMVPNGNAVAFPDALGGEVSKRQSPSCGKHKQVIFKQLVELFRRDLLAGNGGGGGIVGDDEDDGGGGVCGGGENLVTGWQGHVGESIGIENGGVLDVDHQALEQQTPFTSLLMLPNRATTGGVILWDNNPSDQSTQIWDFHLGHSRGYEECGLLEAEYGVNDAGFVIKSYSELMKETSFTNTKVVGEMYDINYSMTHEDITSFNNNSNNPTASQGAATSESNNLPIARPSSGSAFAKPKSFSGSKDIELTEQSILMRGESGRTAATTKVDLEQLAQNRGNAMLRYKEKKKTRRYDKHIRYESRKARADTRKRVKGRFVKATEAPDG

>VIT_214s0068g01380.1

METTSKSRPSAAVPCDFCDSKTAVVHCRADSAKLCLLCDRHVHSANALSRKHLRSQICDNCRTEPVSFRCFTDNLALCQSCDWDSHGNCSVPSLHERTPVESFSGCPSPLELASVFRVDLKDGNWSSWNFGSVNVQDFVVPGENCYAGCGTKVEKNGISVVYEQLVDLIRSDVDVVRGDVDGDGDEGEDGAELGPGTPGRCANMGNFQGVDLDNGDDEELLRQQTPFTSLLMLPTPVDARDTGCGYGCAVEGDAMWDRGHLSYQAPQIWDFHLGRSRICKETSPEAGYDVDNSGFVIKNYSEITKGSSLTRTKALQGMYEMNCTTTHEDILSKNSHSNKALSSQGPTTAESNNIPIVGPSSESWTAEPNTNSIKSMQFKDLLIGSGTARTETTNVDMELLAQNRGHAMLRYKEKKKTRRYEKHIRYESRKARADTRKRVKGRFVKASDS

>VIT_201s0011g04240.1

MEKICEFCTSLRPVVYCKADAALLCLSCDAKVHSANALSNRHPRTLLCESCKCRPTSLRCLDHRVFLCRNCDRSLHEVSSQHHRRAIRSYVGCPSAKDFVALWGFELNEMETVAPRDQFVSTSCGPLEQGVRNSSFSRHSCQQDGVSSLAFELNSSISIFGAESEVGSSTQQSKAFYNGQGQQKTSVILQQILDLKRLQLTDRKNTSSLIRGQEQIDISSSKFSTSQRLDDDLDEHSQNSLGICSDLQQMDSPKQELKVEPFPSPFSQLEQLSSSSTVGIPLHGDPFWQCKSPIQSSQLWSQNMQDLGVCEDVNCFDDFNIPDVDLTFQNFEELFGGDQDPTRLPVDIEDVTYSSLE

>VIT_219s0014g05120.1

MDKEIVVYCHSWTKEPRPDFWLFEVGFLRGVWLMGSVEPVCEFCGVARAVVYCKQDMAALCLQCDGFVHSANFISQRHVRSLLCDKCNSQPATIQCLEDEACLCESCECNVNSCLGSEHKHQPLSFYSGSPSPEEFIEIWSSSPSCKSPVSLSTNCINSYWENRNNSRSLFWSVTSLTNRMKELETFTKMSSSPMSPPQALFFSSVPQLLKQDPSESQNLHICNVNNPCEGRMDDAELNLEKSDKVYGSLQCYINSFEDIGMSCPYMDKSLPADESNIKAVNDQEPINPCRSQSVNLNFPTGLVQSGASLSLSNITGESCVADYQDCGLSAVILNGNPTWDANLGTSCSQARVEAKKRYNKKKKTRMYVCNSTHY

>VIT_212s0059g02500.1

MEPICEFCGVVRAVVYCKSDAARLCLHCDNSVHSANALSRRHLRSLLCDKCNLQPGIYRCMDEKLCICQACDWIGNGCSAPGHRLQSLQFYMGCPSLSDFSRLWSSVLDLPSATGLKAGWGSMNSVAVDENCVSQCLEPKDNEGSLVLGCNKLNEQKPWVGASSSMPGAYSSMPPDRKFTSSYCKDQTPFLPDESNPSKVISFIFSLLINQSQSQGCSNFKDLGLNEGGDLCQGINMDDVAVNFENSDEMIGSSQGHSTCRYDNAGMDSRLMDKNLSASSLRQHNCNAFPSSCAAGSANVIEAMSGSVGCMLVNPSCSRKMGLGFPGGQVHASVSLSLSNVTGESSAADYQDCGLSPAFLAGESPWTSNLDAHCPQARDKAKMRYNEKKKTRTFGKQIRYASRKARADTRKRVRGRFVKAGEAYDYDPLTSTSN

>VIT_201s0011g03520.1

MYPIIVPFCSRFDFGSAFCLMISEKNVANAVGGKTARACDSCIRKRARFYCAADDAFLCQACDMSVHSANPLARRHERVRLKTASLKLPGADSLENSMPSWHQGFTRKARTPRHGKPAAHPAFKSDELTRNPLPFVPEIGADETSYDDNEEQLLYRVPIFDPFVAELCASTNSNEAVTTVANDTETADVTGSETKALVAGRGHDVDSLHGFLPSDMDLAEFAADVESLLGKGLDNESFGMEGLGLIDCKEKESVEYSLHSGRVKLEEEEDIGGVMACQADAEIDMTREPFELNFDYGSPATCEEEEEKVAVGAMDMNNKVDDPKKKNKILLRLDYEAIITAWASQGSPWTNGHRPELDPDDCWPDCLGTCGIQVHHPYGEFGGMGEQQAAMGDGGREARVSRYREKRRTRLFSKKIRYEVRKLNAEKRPRMKGRFVKRASFGGPAAFPLLNK

>VIT_203s0038g00690.1

MRTLCDACESAAAILFCAADEAALCRACDEKVHMCNKLASRHVRVGLADPSDVPRCDICENAPAFFYCEVDGTSLCLQCDMIVHVGGKRTHGRYLLLRQRVEFPGDKPGRLEELRLQSGEPGEARREQNWPPMMTLRETQPNHMASSVPMLENNTHGDGKMDNKLIDLNARPQRVHGQTSNNQSMDVHSGTNHESESVVPVGSFKREPEK

>VIT_204s0023g03030.1

MRTLCDVCESAAAILFCAADEAALCRVCDEKVHMCNKLASRHVRVGLADPSDVPRCDICENAPAFFYCEIDGTSLCLQCDMIVHVGGKRTHGRYLLLRQRVEFPGDKSGNLEDPALLPMEPGENRRGQNQSSKPTVVENQQNRRVSPVPTMDANADGHAKMDTKLIDLNMKPHRIHGQASNNQEACFLLDFSTKPGK

>VIT_218s0001g13520.1

MKIHCDVCSREEATVFCTADEAALCDACDHRVHHANKLASKHQRFSLLHPSPKQVPLCDVCQEKRAFLFCQQDRAILCRDCDLPIHTANEHTQKHNRFLLTGIKLSATSALYSSTTSVADSVSDHKSQSSLKKPESVPPEISHPPSITKTSSPTTAINSINKGGDASLTSEGVSTSSISEYLIEMLPGWHVEDFLDSTSAPSGFCKSAGDDVLPYLLDADLDNNLSSFSSENLGVWVPQAPTPLHPSQYSSFMGGQIGLKESKEATTMKPNSKKWGDDVFTVPQISPPSVGSKRSRSFWQYHH

>VIT_203s0038g00340.1

MKIQCSFCSKEEASVFCTADEAPLCDICDRQVHHANKLAGKHKRYSLLRPSDKDFPSCDLCQDKRAFLFCKEDRAILCRECDVSIHKANEHTRKHYRFLLTGVKLSASASEYPISASSSSPSTIDSETKPSKSSTKRPTSVSADIFCNTAIGAEIKPSKTSTKRPTSVSAGISNPTVKTAPAAASYKRDHDNQSISEYLMETLPGWRVDDFLDPSSGFSEFPDHGVGTHLSSFPYEDFAVWVPQDTPQFNHLPLYIPQTGGGNGLKASEEANTVKVSRKRIDDGFTVPEISTLPLKKSRNLW

>VIT_218s0089g01280.1

MKIPCDICGNVEAEVLCSADEAVLCWGCDERVHTANKLSQKHQRVPLLKHPPSTSSSQLPPCDICQEKSGYFFCLEDRALLCKNCDVSTHSTNSYVSSHRRFVISGIKVALQSVTNNYRTGCNSRTYPLDMPNSNSSSVNFPMDREKKPEMTTEVASTSSDMVAMFSGEIHLATGPEWTLDEILGSNDFDYYEFSDMGQSRISSQ

>VIT_219s0014g00350.1

MKIQCNVCEAAEANVLCCADEAALCWACDEKVHAANKLASKHQRVPLSTSSSQMPKCDICQETVGYFFCLEDRALLCRKCDVSIHTANTYVSAHQRFLLTGVKVGLEPTQPGSSSSMGKSNLVGKHSETESPSASRRGAPMPLTCDYNKTLSIQAGGAGDFVPTKVSFAGGSGSTESIPQWHIDELFGLNEFNQTYDCMDNGTSKADIGKLGDSDCSPTLRAADEELNFDDCLGQVPETTWMVPQISSPPTASGLYWPKSSQISSDTSVFVPDICYSPVQMQNESATKCRRQF

>VIT_205s0102g00750.1

MKIQCDVCERAPATVICCADEAALCAKCDVEVHAANKLASKHQRLLLQCLSNKLPPCDICQEKAAFIFCVEDRALFCRDCDEPIHSAGNLAANHQRFLATGIRVALSSKCAKETDKSSSEPPPNQNSQQITMKMPPQQAPNFTSSSWAVDDLLQFSDFESSDKNKQLEFGELEWLTEMGIFGDQVPQEAMAAAEVPQLPISQPSYGASYRATKSSMPYKRPRIEILDDEDEHFTVPDLG

>VIT_219s0014g03960.1

MKEMKECELCSFPARMFCESDQARLCWDCDEKVHGANFLVARHSRSLLCHACQAPTPWKASGTRLAPTVSVCEACVLRCDKDRHRSSGEDRESEGGNTDDEEDDDDDHDDDDENDGDEGDEDDDVEEEEDGENQVVPLSSSAASLLLPPDASSSSSEDGEASSRFSARGRGGVSALKRTRQNADLDLDSDQDDIGCCSWQHNHKASRTAYDEEDTSLSSRRPCYERQPTELNRSLKLEDQGETESTSTAIVGSIKRFQKSIISGDEAPAMILGICKLSKDPRAVDLRPTANPTIPSRTV

>VIT_212s0134g00400.2

MKGCELCGCPARMYCESDQASLCWDCDAKVHGANFLVARHSRSLLCHVCRSPTPWRASGAKLGHTVSVCERCVDGCGGRKCGGAAEESEGGNDDEVGMEDDDLDEDDEYDDDDDVEDEEEEEEEDDDFGGDDEDGDNQVVPWSSTPPPPEASSSSSEESLNSFSSGNRDVPGSFTMLSLKRMRETASDLRSQDDFNRPSSQPNYASARAAAARSAADDGEAISEDALRPFKDRRIEQNRPVQPQAGSRTAEIVDSLNRLHNQDITSGSDASSIIAGICNLSKQSGAVDLDCSSSRSRRI

>VIT_212s0059g02510.1

MCRGREDGKQGRLAKSEVLKEAVSDDGGGGLVSCELCSSRALLYCQADDAFLCQKCDRWVHGANFLAFRHIRCLLCSTCQNLTQRYLIGSSHEVALPTMISWTERSWCNSSDESKCPRTLKMPFLFL

>VIT_200s0203g00210.1

MKVCELCNSPAVIYCDSDQASLCCDCDAKVHSANFLVAKHSRTLLCHVCQSPTPWNGSGPKLGSTISVCQRCVNRSRTVNETPSTDEDEDEEGEEDGEDDGDDENDEENQVVPWSSTPPPPISCSSASEEESSSRFCNGEEGPSDSKSASSLKPMRQQSSFRPTSQNDLGRSVSPPAHNVRIFMKGNE

>VIT_209s0054g00530.1

MKGRVCELCNEEASLYCGSDSAFLCWSCDARVHGANFLVARHVRHTLCSECNGLAGDTFFGVGFQPHRLICRSCSSEVESETSTDHDSKSSSSPCVSTTESAPRKGGVSRRKAERTGFTSSVSAVSGVDSRFPSKLRARSSVDAKAEDILVNWCRKLGLNGSCTSVASHALGVCLVKLTVLPLRVSLVAAISCAAKLSGDRSAYTPQNLKRLVEISGVPAKLILAAESKLARVLKMERRRPRHVRDREEGWAECSGEEFR

>TCA.TCM_014441annotation

MVTYQANWARTCDTCQAAACTLYCHTDSSYLCNDCDKRIHAANPMASSHQRVWICAVCENAAAAVTCRADAASLCIKCDIEIHSVNPLARRHIRVELPDPMFDTENEIAAGTINEEIDENETDCWLLLEPDSTDNQTMSGFTYGEQVDEYMDVRDTCTEYRCQEQCSDQQQLLCVNNPEDSGSDIDVPVQTFESKKQSQQQEKQLQRQTHQQLQGIYFNTEHRGSKAAFMYTPSSTLSVPLPLITAGILPNATSNIPSTYTGFPNGATDLFPYPLPLMPLQFTPMNREAKVLRYREKRKARKFEKKIRYASRKAYAETRPRVKGRFARKTDMEIEDDQLFSKEDYGYCIVPSL

>TCA.TCM_019107

MLKEETSDASGGGGSGNNWARVCDTCRSAACTVYCQADSAYLCAGCDARVHAANRVASRHERVWVCEACERAPAAFLCKADAASLCTTCDAEIHSANPLARRHQRVPILPISGCLYGPSATELRGRKMASAAETEDGFMDPEGDETIAEDEDEAASWLLLNPGKNSTNQRNGCLFGGEVDDYLDIVEYNSSVENHITDQYNQQQQHYSVPNKSYGGDSVVPIQSGEAKDHLQQQQHQQQTLQFGLDYESSKAAYSYNGSVSHSVSLSSVDVGIVPESTMSDISISHRRPPKGTIDLFSGPSIQMPTQLTPMDREARVLRYREKKKTRKFEKTIRYASRKAYAETRPRIKGRFAKRTDVEVEVDQMFSTTLMTETGYGIVPSF

>TCA.TCM_036860

MASKLCDSCKSATATLFCRADSAFLCSNCDSKIHAANKLASRHSRVWVCEVCEQAPAHVTCKADAAALCVTCDRDIHSANPLARRHERVPVTPFYDSVNSVPAVKPNGVVNFLDERYFSEVDGDADVSREEAEAASWLLPNPNHKAVESPDVNTGQYVFSEMDPYLDLDYGHVDPKMEAQEQNSSGTDGVVPVQSKSVQAPMVNDHCFDLDFTGSKPFAYGYNAHCVSHSVSSSSLDVGVVPDGSAMTDISNPYGRGAESTHQTVQLSSADREARVLRYREKRKNRKFEKTIRYASRKAYAEMRPRIKGRFAKRSDIEVEADRRNMYGFGVVPSF

>TCA.TCM_038829

MRMGMEIASVKGIPGGWGMAAKTCDTCKSAAAAIFCRADSAFMCLNCDSRIHSGNNKLVSSRHERVWMCEVCEQAPAAVTCKADAAALCVTCDADIHSANPLARRHERVPVEPFFDSADTVVKSSPFSFLVPTDRSGACQQEDVEPGSWLLPNPNLISKLSGETNQVKTGDLFFSEIDPFIDFEYQNSFQPHNNAAMDSVVPVQAKPATIPVINNENCFDIDFCRSKLTAFSYQTPSLSQSVSSSSLEVGVVPDGNTLSEISYPFGRTMTDPSVPISATTTNNQAPQACGIDREARVLRYREKRKNRKFEKTIRYASRKAYAESRPRIKGRFAKRAEIDNEVDHMYNSPSAAAAFMSDAQYSVVPSF

>TCA.TCM_002458

MGYICDFCGDQRSMVYCRSDAACLCLSCDRNVHSANALSKRHSRTLLCERCNSQPAFVRCAEERISLCQNCDWMGHGTSTSNSTHKRQTINCYSGCPSAAELSSVWSFVLESPSAGESACEQELGLMSITENAERTSWDPTENTISQNGTGVAEVNDDLDADKGSSWGGSASVPELRSAPRLLDQPAGSTDLLPKLCCPQTKCPGLCEDDLYADFNMDEVDLNLENYEELFGVTLNHSEELFENGGIDSLFGTKDMSAADSNCQGAVAAEGSSVGLVNAIQPACSNAASADSMMSNKTDSVLCFTARQAHSSLSFSGLTGESSAGDYQDCGASSMLLMGEPPWCPPCNENSFSSATRSDAVMRYKEKKKTRKFEKRVRYASRKARADVRKRVKGRFVKAGDAYDYDPLNQTRSC

>TCA.TCM_004548

MTDQKNNNQKQRRLCDYCNQSKALLYCRADSAKLCFSCDREVHSANQLFTRHPRSQLCDACDKSPASIFCETEQSVFCSNCDWESHKCSSSSLHNRRPIEGFSGCPSVSELVNSFGIEDLGSKTLFLSEEKVGCGDGGEDDGLLDLLSWETPEFSSLDDLIVSSDFGHGFKPTDVPPLRKNRNASCGRHKEEMLLQLRELAKSEPNLSTDFENVTGFLPLLPNTYNSQPGSVHTSCKIDKDPIPFPAYELSAPQCFSDNVEMANQVFLPFSQLRGYTEESAVVPNEHLDTSRTAHVNDSLEDQLQHQIAAGTTSALPKIVVHELNSQERDSAISRYKEKRKTRRYDKHIRYESRKVRAESRTRIKGRFAKVEH

>TCA.TCM_007770

MEPLCDFCRGVRAVVYCKSDAARLCLSCDGCVHSANLLSRRHARSLLCEKCNSQPAVVRCLDEKLSLCQACDWNSNSCSSFRHRREALNCYTGCPSLAEFRRIWSSVLDASSSSAFDAGLPVGSLPANDNCVINCLNQREPGGAFGLVGTKLNEPDPCPKLEPWMGPSSLIPTNANYMPYCRNQEPLFSEESNIPKGCSDLKDFKLLDGDDLCEGINMNDVQLNFETADEIFGCSQGQNRSQFDNVGTEGLVMEKNITLTESDVPIEHTLEASSSGQKDCMAFPGSQVGGSASVMAAMTGTSNCMLMNRGCNRNINLEYPAGQIPSTIALSLSNIAGEHGAADFQDYGLSPVFLTGESPWESNLEASCPLARDKAKMRYNEKKKTRTFGKQIRYASRKARADTRKRVKGRFVKAGEAYDYDPLVARNF

>TCA.TCM_008446

MVSPKPESKEMVPCDFCNEQIAVLYCRADSAKLCLFCDQHVHSANLLSRKHLRSQICDNCSTEPVSVRCATDNLVLCQECDWDAHGSCSVSSAHDRNPVEGFSGCPSALELASAWGFDLEEKKPLAKSWNGCHQDLMMPAMESWLYKSSLQEMMVPYECFTCEETVKKQSHGSGKCKQIIFKQLVELMKRDFMAGDVADGGRGAGENLVPYVEAKGSGLARQEFIQPQPQPQTQTPFTSLLMMQTRESERTVDGGDVLWNGNPNNQTPQIWDFNLGRLRGHEECSQLEDVGYGGSDAGFMIKNFGELMKETSLSNAKMLGDMCHINYTPPQDDMASINNSSNLAASQGPATSESNNLPIARPSSVSAFGKPKGSSSSKDVHFIEHPILMRGDQVRQSAPSKADLELLTQNRGNAMQRYKEKKKTRRYDKHIRYESRKARADTRKRVKGRFVKATEAPDH

>TCA.TCM_009367

MEKICEFCTTSGPVVYCKADAAHLCLSCDAKVHSANTLSNRHLRTLLCDSCRYRPSYVRCLDHQMFMCRGCDRTFHDASSQHQRLAVSSYLGCPSAKDFAALWGFELNELENNAIQDHSLSNSCVSVNPNAVKLDDLGQSCSQIGVSSSKSCVTQAPAAVCNVGSNSQQTKVINKGQQQQNTAFILQQILDLKKLQLTERDGHLPFIGGQEQADTSSSICNFSQNLDSNRVHDIGINLHQSNNPIHEQNADPLPLSFSHLENLASSSTSGIPLYGESFWQCKSPIRGSQLWSQNMQDLGVCEDIFCQDDFNMPDIELTFPNFEDLFGADQDPIRALLGNKDVCCSSVDKDMSFNKSDIVNARPVEDASVASSIYINQSAHTENNMDPSNHIHNFQRASDSPRPIRPSYSTMSFSVSRFSAESSGIDCPDSELSPITQGEALCFSPGLGSLHSEARENAMMRYKEKKKARLHERQIRSASWKARTDVRKMVKGRFPKMEDYDSDNANGTRSY

>TCA.TCM_017297

MMEKVSKSKPRRSVLVLCDFCNSKPAVLYCIADSAKLCLFCDQRVHSANALSFKHARSQICDGCKTKPASFLCSNDNLIFCQDCDWNSHNSNCSVSALHERSPVERFSGCPSVTELASLLGFDLKPKYLMNLDPGFSLYEQELMNLEDFMVPNENSSVSSALLSSVKLNHEVYRQLVEMGKRDLVRVSGDGAELGPGTPPGRSSEKGNLGSFEVENGDDEELLHQRTSFSSLLMLQNNVALRKTDYVAEGELIWDYNSSYQASKIWDFQSGRSRDYEESCPEDAGCMNSISSRGFVIKNYDEFAKEGSLSETKVFQDMDQMSLSMKCEDTLSRSSRCNQPFSSYTATTEASNHVPVVGSSSDSKVVEPLKCDSTRYVQVMEHLVLAGGESMNEAKAKIDMELLAQSRGNAMLRYQEKKKHRRYDKHIRYESRKARADTRKRFKGRFVKASEAPDVKV

>TCA.TCM_009901

MISTKNVANAVGGKTARACDSCIKKRARWYCAADDAFLCQACDSSVHSANPLARRHERVRLKTASLQSSGHEAPLESFAPSWHKGFTRKARTRRPTKASIHQKLKAEHTKRNSNPFPLVPEIGADEISYEENEEEQLLYRVPIFDPLVAKLCTSTTSNEAAVSAVGNDVETADAAVSESKAFLACNGQDADGSHGLFPSEMDLAEFAADVESLLGKGLENESFGMEDLGLKDSKEKYFRDCSLGNGKVKIEDEESFEAVGACHVDSEIDMAREPFKLNFSCDSPGNCGEEDELVKEEVTVKSYEEYEEDTAKKKKRKILLSLNYESVITAWASQGSPWTSGGRPDFDPDECWPDCMGTGGTEVHHTYSDLIGMGAHQALGDGGREARVSRYREKRRTRLFSKKIRYEVRKLNAEKRPRMKGRFVKRASFLAGPAFPFVNK

>TCA.TCM_012333

MTTDKKAANAMGGKTARACDGCLRKRARWYCAADDAFLCQGCDTSVHSANQLASRHERVRLETASFSKFSASVTNRTNQDAPPAWHQGFTRKARTPRQNKSMLGQQKDEGKVFSLNPLDPVVPEIGSEDGSVDENEEQLLCRVPVLNPFAAELCNMVTSDGDEVAMVNEDGNFVIDDYEREGTCELDDGLHGFLPSDMDLADFAADVENLLGVGLDDDACCDTKGIELLHSKEEDGSNVLHEGKIVKVKDEEEVEGITACCFDSAFDVTRESLNWSYDYESPTIGEEEEEKVLPATETTTMNGESKAEMKRNMLLRLNYESVITAWARQGSPWTAGSRPEFNPDDFMGSNLKEGHHLSGGIGGIYSLQGRGNTDGEREARVSRYREKRRTRLFSKKIRYEVRKLNAEKRPRMKGRFVKRTSFVGTAFTYIN

>TCA.TCM_000244

MRTLCDSCESAAAIVFCAADEAALCRACDEKVHMCNKLASRHVRVGLANPSDVPLCDICENAPAFFYCEIDGSSLCLQCDMIVHVGGKRTHARYLLFRQRVEFPGDKPGNVEDPASQPVDPGETRRGQNQPAKPTVGESQQNHKVSSVQLVDANADGHVKMDTKMIDLNMKPHRIHGQASNNQEQ

>TCA.TCM_006783

MKKEEKEVEDKWRVRDLHTAAIFRSVEECLRSEERRSKREANRKKKNRMRTLCDVCESAAAILFCAADEAALCRSCDEKVHMCNKLASRHVRVGLADPSDVPRCDICENAPAFFYCEVDGSSLCLQCDMIVHVGGKRTHGRYLLLRQRVEFPGDKPGRLDELGLQTLDPNEVRKDKNQQQPKLAARENQQNHRVSPVPVLDGNSDGDGKVGNKLIDLNAKPQRVHGQASTNQEQGMDISSGNNHDSSSVVPVGSFKREPDK

>TCA.TCM_006809

MKIWCDVCDKDEAVVFCSADEAALCESCDRRVHHANKLARKHSRFSLLHPTFKESPLCDICQVRRAFLFCQEDRAILCRECDLPIHRTNEHTQKHNRFLLTGVKLSSSSSSPCLNPTSSSSNGHNATTIDSETKSSQSCKRFRSVSNNEIFSSPSIEKPLPSTTDKVEDNCTSDTVSISTSSISEYLMETLPGWRVDDFLEPSSAANGFSVWLPQVSPPQSPQLCFLIPQNDLVDGFKELKEANVLNFGCRWNGESLIVPQISSSSSLKKSRLFR

>TCA.TCM_020013

MKIQCDVCERAPATVICCADEAALCAKCDVEVHAANKLASKHQRLLLQCLSNKLPPCDICQEKAAFIFCVEDRALFCQDCDEPIHSAGSLSANHQRFLATGIRVALSSSCNKNTEKSGLEPPNKSAPQTSMKMPVQQQSNFTSSWAVDDLLQFSDIESPEKQKEQLELGELEWLADIGLFGEQLPQEALAPAEVPQLPIPQSANFNSCRPTRYSMPLKKPRIEIPEDDDDEFFTVPDLG

>TCA.TCM_032112

MKIQCNVCEAAEAKVLCCADEAALCWACDEKVHAANKLASKHQRVPLSSSSSHMPKCDICQETSGFFFCLQDRALLCRKCDLAIHTANTYVSGHQRFLLTGVKVGPETTDPGASSSNVQSPSNEKTSEAKSNSTSRRGTPMALTGGQNEVLLANAGVGNSVPTQVLYAGGSAAGSIQSWQMDDLFGLTDFNQSYGYMDNVSSKADSGRRGDSDSSSILRSAEEEVDDDECLGQVPESSCAVPQVPSPPTASGLYWPKDSHNQSDGVVFVPDICSSIVKNPFHSRCHGSRPKRQRQI

>TCA.TCM_034151

MNPHYVTAILFYSLLSTDALITLSSQLFWGSKLPQSLTQYSLSEREIGSKEIKQIKQMKIQCDVCSKEEASVFCTADEAALCDACDHRVHHANKLASKHQRFSLLHPASSKQAPLCDICQEKRAFLFCQQDRAILCRDCDVPIHAANEHTQKHNRFLLTGVKLSATSALYTSSSSSSIASLSTGCDSVPEFESQPSIKNPVSASPTNLNPFSLAKSSPVSTTAAAVTNKSGGDNLLANEGGGSTSSISEYLIEMLPGWHFEDFLDSSSPPFGFCKSDDGMLPFSDADLESNKSSFSPESLGLWVPQSPSALYPPQYSSTMGGQIGFKETKEIIGMKANRRWTDDAFTVPQISLPSTGSKRTRPLW

>TCA.TCM_007771

MCRGLQQGNPSGFCLKEGVSPNATRVSGLVNCELCSSRASLYCQADDAFLCRKCDKWVHEANFLALRHIRCFLCNTCQNLTQRYLIGASHEAMLPTMVFHGGRSGRKRFLDGCIHM

>TCA.TCM_007842

MKSCELCKLAARTYCESDQASLCWNCDAKVHGANFLVARHVRCLLCHTCQSVTPWRAAGAKLGHTVSVCERCVNGGDREESEAENDDEDDDDDDDEEEEEVDSDDDVSVDDDVEEDGDNQVVPWSTVVITPPPSSSSSSSDDSSGGEREVSESTNLFSLKRLRENASDLLSQDDPDPSPSKRKYSYRTVWGTLCRPDDDAVSVDSVRLLKDQPIQPDGSLQFQADSSPRGAASTESLGKLGPDKIQ

>TCA.TCM_030024

MGVELRCEDILELRNGTEHIGIKLSDAHGNSSLRAILSQNKKKKKMKKCELCNSLAKMYCESDLAILCWDCDSRVHGANFLVAKHLRTLLCHLCQSPTPWNGSGPKLGPTVSACDNCVNRNACREESNNEETHDEEDEDDDDDLDGEDDSEDDDGDNGDDEENQVVPWSSTPPDSSSSTSEECSTRFCSVQEGTSQSRTVLSLKRMRETEEPASRQADDPGCSFSPQHQTQNLSNESASFDPFRSLKDQKITARDSLKRLQKGVVAGAI

>TCA.TCM_031287

MKKCELCEGLARMHCESDQANLCWDCDVEVHGANFLVAKHSRTLLCRVCQNPTPWLASGRNLSPAVSVCESCVGNNNKKNNGSICEVTDQQEESSEEEYEDDEEEEEEEDHEEEEEEEEEEEEEVEDGENQVVPWSGDSSSFSMSKPVSSLDSLSSSEGGGLRLKRMREHLSFYSDDEFGCSSSHVGSGGSTNGEATSMGSSRLSKQPRLCEVNQSARNQDHSETESRSTAIISSLKRLQNHMITNDNDASATILGICRLSRDQSPLDFTSH

>Prupe.1G093900

MGIQAGGWRTMAGGLVAKPCDSCKNSAAVLFCRADSAFLCLTCDAKIHAANKLATRHERVWMCEVCEQAPASVTCKADAAALCVTCDADIHSANPLSRRHDRVPVEPFFDSAESIIVKSAAASSSVDSLNFLLPNGAVPSHTKDDENDAASTWLIPNLNFNSKLQMDIAPDIIKSSELFFPEMDSLLEFDYPNPIHHDISGSGTDSVVPVQPDPLPPPSLNINRVSAEQNCFDLEFCRSKLSSSFSYPTQSLSQSVSSSSVEVGVVPDGNSMSDISYPFGRNANHNVSDPSAQVSATTANQVATQLSGLDREARVLRYREKRKNRKFQKTIRYASRKAYAETRPRIKGRFAKRTETEIEAVDNIYGSAQGAFIPETYGVVPTF

>Prupe.1G398700

MASKLCDSCKSATATLFCRADSAFLCVNCDSKIHAANKLASRHARVWLCEVCEQAPAHVTCKADDAALCVTCDRDIHSANPLSRRHERVPVTPFYDSGNSAANSAPVVKSVVNFLDDRYFSDVDGQDAETEVSREEAEAASWLLPNPKAMENPDLNSGEYFLPEMDPYLDLDYGHVDPKLEDAQEQNSCGTDGVVPVQSKSVQPQLVNDHSFEIDFSAASKPYVYGFHAQCLSQSVSSSSMDVSVVPDGNTTMTDVCDPYTKSMSAAVESTHQAVQISSADREARVLRYREKRKNRKFEKTIRYASRKAYAETRPRIKGRFAKRTEVEIEAERLCRYGVVPSF

>Prupe.3G245100

MLKEESNGAAATKNWARVCDTCRSAACTVYCRADSAYLCSGCDATIHAANRVASRHERVWVCEACEQAPAAFLCKADAASLCTACDADIHSANPLARRHQRVPILPISGCLYGPQATDPGEIVVGAAAAETEDGLLSQEGDETIDEEDEDEAASWLLLNPMKNNNNNCNNNNNQSNGFFFGVEVDEYLDLVEYNSSCGDQNQFTDHHQSNQQQEQQQQHYGVPVPHKNYGGDSVVPVQMQKQSFHQLGLEYESSKAAYSYNGSLSNTVSVSSMDVGVVPDSTMSDISISHPRTPKGTIDLFSGPPIQMPSQLSPLDREARVLRYREKKKTRKFEKTIRYASRKAYAETRPRIKGRFAKRTEMEVEVDQMFSTTLMAENGYGIVPSF

>Prupe.1G219800

MASPKSRSGEGVPCDFCTDQPAVLYCKADSAKLCLFCDQHVHSANLLSRKHVRSQICDNCASEPVSVRCSTDNLVLCQECDWDAHGSCSVSAAHDRTPLEGFSGCPSALQLASLLGLDLHDKNVPGQPDPQLQNWDMGMPSLDSSWSGMGLQDLMVPNHNQNGVVYPNDELMVKRQSAGGISGKQKQGIQKQLVELLKRDLDGGGGGGGGGGCGDGGENLVVPRTPTSSAWQEDNGNGNVEGLEPLDLGNRNGGVDGVVVSAATSQPLLQQQAPFTSLLMMPEENRGIVDGDMLWNSNPHGQSSEIWDFHLGRSRDHEESGPLEVTYGSNDSGFMIKNFGELMKETSLTDTKMFRDLYQMNCPVGHDDIKFSNNSNNPTPSQGPATSESNNIPVGRPSLGSAFGEDKGSGASTDLNFMEQSFLMRGDSLRTVGTKADMELLAQNRGNAMLRYKEKKKTRRYDKHIRYESRKARADTRKRVKGRFVKATEAPDG

>Prupe.1G318100

MKKLCEFCSVLRPVVYCKADAAHLCLSCDAKVHSANTLFNRHLRTILCNSCKYRPAYVLCLDHKMFMCRVCDITLHIASQHQKRATSSYKGCPSAKDFAAFWGFELNELDSSAHLDGILSTSCGSSCDSTAVNLDSCEQSCSQVEGFPVANSPTLAPGVDSEVGSRSQQYKILDLKRLQLTEGRSPSLLIRGKEQSDLSSSVHHTSGRFDDNLDQSLQHSEITRFRQRDSLLQDLKVDNLPFPLSQLEHMPPSSTAGLPLDTESFWHCRSPVESCQLWAQNMQDIGVCEELVCHDDFNMPDVDMTFQNFEELFGSDQDPTRALLDDKDVPYSSVLKYISLDKSDNGHARARMEHDASEVSSIFFNKVQNLEGSMDYCPCPIQPSSSTLSFCMSRFSAESSGTDCHESRLSPYIATGSSRNSPHDQEGGRFETRANAMTRYKEKKHSRLLVRKNRYPFRKGTADNVSGKKEEL

>Prupe.1G318400annotation

MEKACELCLELRPVVYCKADSAHLCLPCDAKVHSANPIVGCHHRTILCDSCKYRSVDIECLDHLMFMCRVCDASLHRKCSPHGQHRKRTLASSFTGCPSAKDFAAFWGFKLDESDGILDLKRLQLSQDQPNQENPSRSLEHSLNHHRLQHSQDDLGRTGSQQRDSCLPVQGLMVKFDPLPLPFPEQLEQYFPSSSSTAGFPLHTESFWQGRSTVQNSRLWSQNMQDLGVCEEQAYDHHDPNTPDVDSTFRNFEESFGAKLQPRKCTTTPPPPPPMAASPPHCRSNRRLAQTTPCSLPSIPPSSIASRPSLDTTNRSPTRPQITPTANSTTLSLSKTTRRWSTAIAAPSSNSTKPPRKPTLSAARTFISAPSISSSTSISVSLSTPPSRNSSFPP

>Prupe.3G194500

MGYICDFCGDQRSMVYCRSDAACLCLSCDRNVHSANALSRRHSRTLLCERCNSQPALVRCTEERVSLCQNCDWMGHGTSTSASAHKRQTLNCYSGCPSASELSSIWSFVLDLPAAGESACEQEMGLMSIAENSTGSAWSPPESNAKQNASGTVEVNGVRSVDKSGVLVGSSSVPELNPATHVVGQQAGSENATLPKLFCHGTKGPAIPEDDDLYDDFDMDEMDLNLENYEELFGVSLNHSEELFKNGGIDSLFGAKNMSRGNSNYQDVGAAEGSSVGLVNALQPACSNAASADSVMSTKTDPIICFPAKQAQSNLSFSGVTGESSAGDCQDCGASSMLLMGEPPWCPPCPESSMQSANRSNAVMRYKEKKKARKFDKRVRYASRKARADVRKRVKGRFIKAGEAYDYDPLNQARTRSY

>Prupe.3G220900

MEPLCEFCGVVRAVVYCKSDLARLCLHCDGCVHSANILSRRHPRSLLCDKCNAQPAMVRCMDEKLSLCQSCDQWNHNNGDTTTGMVGHRTHALTCYTGCPSLSEFSRIWSSVLDGSSSSGFETNGGGGGGGGNNNWDQSLGSAQVPINDQTNCIGNALERRDNSERSFGVVTSKLSGSLDQSSCAKFEPWMARSTVISANPNGIPQGNKDQAPFLPQESSLSKDSNDLCDGLNIDDVPLNFENAEGIFSCPQGPSRYQFEDGEMDCLLMEKNLSVTESNGPNDNAVQASSSGPQDCVAFQSSCGSDNVMPAMNGSANCLLMNPTCSRNINLGFPTGQVHSSMSLSLSSMSRESNPDYQDCGLSPIFLTGEPWDSAMEASSPQARDKAKMRYKEKKKTRMFGKQIRYASRKARADTRKRVKGRFVKAGEEYDYDPLVTRNF

>Prupe.8G253900

MNGSSRNEEAEAPQPEREHQKRLCDYCGSSMALLYCRADSAKLCFLCDREVHSANQLFSKHTRSLLCDACDGSPASIFCTTESSVLCQNCDWESHNLSSSSVHDRRPLEGFSGCPCLNELLTVVGFENMDKKALLLSDESGGGGGGGDGFLGCGVDGSFDLDDGFSDFLVWDTPSVVSLDDLIVSNPAYKFQAMGVPPLPKNRNAACGRHKEEIFRQLRVLVKSEPNLFSENVDVKPLKSLASEQNMQRGSLFTGFEHDAEPTVFPAYEAHDFQYNDCGAAEKRESSPKTFIRSYLQECCVVPDKHSYNDGSASHANDGHGHGGQMNSEASSAFPKVAAHELSSQNRETALSRYKEKKKTRRYEKHIRYESRKVRAESRTRIKGRFAKMDH

>Prupe.1G310900

MSSNKKVANAVGAKTARACDSCIRKRARWYCAADDAFLCQACDSSVHSANQLARRHERVLLKTASSVKPTDKDDQVSVSDNPAAPSWHGGFTRKARTPRQGKHKSQEGARNRFPQVPEVGADDSNSYDEENEEDQQQLLYRVPVFDPFVAEMCTATTSANSNEEAVANTCDVSKVSSSSPNYNHNGRDGFLLPSDMDLAEFAADVDSMLGRGLEDDECFGMEGLGLMDSKENESNCRVKLEDEEEEEEEEGGTGGVNFMGCELGETPEIDMMREPFVLNFEDYDDSPQSCGEQDDKLGVGMMDTTYSGDHQQHEEVHTGASKSGKKKEIFLRLDYEAVITAWDGSPWTSGGRPDFSSECLPDCMGVCGTELHYPYGDLNGLGVHPAMADGGREARVSRYREKRRTRLFSKKIRYEVRKLNAEKRPRMKGRFVKRASFAGSAFPVLTK

>Prupe.5G221000

MITEKKAANAVGGNTARACDSCLIKRARWFCAADDAFLCQRCDGSVHSANQLASRHERVKLQAASSKSNQSTLLAADQSPPAWHKGFTRKARTPRHNNKPPFMAKDDKEKVSLNPPLPFVPEIGCEEANYMDDDEENEDLEQLLYRVPIYDPFAAELCNMSSHEVGNSTSAINLDDEVTEMGRGNHHEVGDDDEENNNNNYGGNELDNLPGFLPSDMELAEFAADVENLIGGGLDDEDSGDIKGVLGLLDDSKLEGNGGVDVCMKNVDINEDRKAVKVEEEAEGMEFNPSMLDWNSDYELRLSSSPLPMGGEEEKVVFLMGARSSDEVALMDNKQMMMKRKICLRLDYEAIITAWANQGSPWTTGTRPELNPDDGWLDCMGMRGSDQIHQPHGHGESARCVGGRGDDRGREARVSRYREKRRTRLFSKKIRYEVRKLNAEKRPRMKGRFVKRTSSSNFVGSSSSNSSKTAAAAFPYLIHQ

>Prupe.1G197100

MKIQCDVCEKAPATVICCADEAALCAKCDVEVHAANKLASKHQRLLLNCLSNKLPRCDICQDKAAFIFCVEDRALFCQDCDEPIHSANSLSANHQRFLATGIRVALSSSCTKEAETSSLEPPERSTQQISTKISAPQASGVLSPWGVDDLLQLSDFESSDKKESLEFGELEWIADMGLFGEQFPQEAMAAAEVPQLPASQPSNFASQRPPKSNVPYKKPRIEIADDDDEHFTVPDLGIF

>Prupe.1G371100

MKIQCDVCNKDDASVFCTADEAALCDTCDHRVHHANKLASKHQRFSLIHPSSKQFPVCDICQERRAFLFCQQDRAILCRECDLPVHAANEHTQKHSRFLLTGVKISATSTLYTSSSPPTPTISLKSADATVTDPKPQPLIKKSVSTSAPAISNPPSMSKNSTLTTNTANSNKGGGIFVAHDGVGCGSTSSISEYLIETLPGWHVEDFLDFSSGPLGFCKADNETVLPFMDADLESNLSSFSSEHMGIWVPQASNPLHQYSQMGGELIGLNKDGTNMKANNRTWRDDSFTVPQISTPSVGSRRSRPF

>Prupe.3G155900

MKIQCDVCEKAPATVICCADEAALCAKCDVEVHAANKLASKHQRLLLQCLSKLNKLPRCDICQDKAAFIFCVEDRALFCQDCDGSVHSANSHSANHQRFLATGIRVALSSSCTTKDTETSSLEPPSHGSQQISTKLPTPQPSGFSSPWGVDDLLQLSDFESSDKKGSLEFGELEWIADMGLFGEQFPEEALAAAEVPQLPVSQQPNFTSYRPPKSNNPYKKPRIVMAEDDDEHFTVPDLGDFRHVST

>Prupe.4G156600

MKIQCNVCEAAEANVLCCADEAALCWACDEKVHKANKLASKHQRVPLSASHMPKCDICQEAVGYFFCLEDRALLCRTCDVAIHAANSLVSAHRRFLLTGIKVGPEPTEPDSGGGGVGGGGVGVGASSSSVKLRSGSGSGSGSGSRCDTHNPMPVECKVAPAGVDVMPFAGGSSAGTVPQWHIDEFLGLSDFDQSFSYIENGSSKADCGKLGEYDSPALKSSEEEMEDYECIGEVPETSWMVPQVPSPPTASGLYWPRSSQISSDFAVFVPDICHSQMQNPLYSQHNGTVSKRRRQF

>Prupe.7G165700

MRTLCDSCESAAAIVFCAADEAALCPACDEKVHMCNKLASRHVRVGLATPSEVPRCDICENAPAFFYCEIDGSSLCLQCDMVVHVGGKRTHGRYLVLRQRVEFPGDKPGNVDDPASQPIDLGETRRVQHQPPRMTIGENQQNHRASPIRISDANADGHVKMDTKLIDLNMKPHRMHEQASNKED

>Prupe.8G085000

MRTLCDVCEGAAAIVFCAADEAALCRSCDEKVHMCNKLASRHIRVGLASPSDVPCCDICENAPAFFYCEVDGSSLCLQCDMIVHVGGKRTHRRYLLFRQRVEFPGDKPGRSEELGLQPLDQKEVRKDHIQPPSLSIRENQQNCSASPVAVLDNNIVGDYKMDNRLIDLNTRPQRMNGQASTSPEQGLDVQNGVNDESASVVPVGSVKR

>Prupe.8G087900

MKIRCDVCEKEEATVFCSADEAALCDVCDRRVHYANKLAGKHKRFCLLHPTFKDSPLCDICQERRGFLFCQQDRAILCRECDFSIHKTNEHTQKHNRFVLPGVKLSAAASLYPTSSSSCSGFSQLANTTDARASKSSSKRPKTVSDKALNCSPSVEQTTSSSSYKTGENCGSDNGSVSTSSISEYLMETLPGWHVEDFFDFSFAPDGF

>Prupe.3G020100

MKARVCELCDQEASLYCPSDSAFLCSRCDARVHQANFLVARHIRQYICYNCKGLTGSRNIRSFCSSCSPDNFSGHGNGDGDTQSSSSACSACVSSTDSFGGTAATKAGFDNLKSESSVTQVSGKLSNIPARFSGAKRKCVQRAQARTSTSADAKAKGSFINWCSQLGLNGNYTAAVVSTASNALGFCLGRLAGVPLRVCLAASFWFALRFCGGRSVSTRQNLRRIEELSGVPAKLILAVEAKLGRELRIRRARRDDLEEGWAEC

>Prupe.3G221100

MCRGVEESGFRQPPVSSPTGYVCCELCSKRASLYCLADDAYLCRKCDQWVHEANFLALRHVRCLLCNTCQNLTQRYVVRISVEVMLPTILSWAERKRCSSNNKRRRSTTLKRPFLFM

>Prupe.3G226900

MKNCELCQLPARTYCESDQAILCWDCDAKVHGANFLVARHSRSLLCHSCQSPTPWKASGEKLGHTFSVCESCVVRDESRDEDEDEESQGGNDNEFDSDNDPDDDDDDFDEDGDNQVVPWSSTTPPPPSPSSSSSEEAASALNNGDTEGPKTATTVSLKRIRETASDLRSQDDLDRSSSRRRYGSASAAQASRPEPEGGATSFDSSRPRKDRRIDLNGPGSPSEPVIESKMIYRREDSLRS

>Prupe.4G033600

MKKKCELCDSVAKMYCESDQASLCWECDIKVHAANFLVAKHSRTLLCHVCQSPTPWTGSGLNLGPTVSVCEKCVHTSNQEPRNEENDEQEEDDDHDHDHDDRDDGENNPEEEDDGDDDDGDRDDDDEEENQVVPWSPPPVSSSSLSSSEECFSESLSKSTTAFSCKRMRSCNSHSNFQNEEGLNWDTRTRREPR

>Prupe.4G192700

MRECELCGLRARIHCEADQAKLCWDCDEKVHGANFLVAKHPRNLLCHGCQLPTPWMGSGPKLTPTVSVCEICVERRGIKFQRYEDQESEAENEDDVDLDDEEDDDDGGHDGEDVDDDADDDDDDEDENQVVPWSSSHSAAAEPPPAVSCSSSEEEEEEEEFLVVSKRMRENADPDSDLGCDETSSLASLRLLKRARPGEENRLLSERSSEAELRSTAIVSSIESLQNQTDVISDGDDQASAAVLGICEPRRDVLVADIVFLRGTFIHDA

>Potri.017G107500

MLKEESGGSGSGGVVNNWARVCDTCRAAACTVYCRADSAYLCAGCDARVHAANRVASRHERVRVCEACERAPAALLCKADAASLCTACDADIHSANPLARRHQRVPILPISGYLYGTQVGPAAGETEDQFMTQEGEETIGEEDEDEAASWLLLNPAKNSNNQNNNGFLFGGEVDEYLDIVEYNSCAENQYSDQYNQQHYSVPPKSCGGDSVVPIQYGEGKDHQQQQQQQHHNFQLGLEYEPAKAAYSYDGSVSQGVSMSSMDVGVVPESAMSEISISHQSAPRGTIDLFSSPPIQMPSQLSPMEREARVLRYREKKKARKFEKTIRYASRKAYAETRPRIKGRFAKRTDVDVEVDQMFSSTLMAETAYGIVPSF

>Potri.018G013800

MASKLCDSCKSATATLFCRADSAFLCISCDSKIHAANKLASRHARVSVCEVCEQAPAHFTCKADAAALCVTCDRDIHSANPLASRHERVPITPFFDSSSTVHGGGEAVNLLEDRYFDEVDGGRGDVSREEAEAESWLLPNPGGGTTKGVDSMDLNTGQYVFGSEMDPYLDLDPYVDPKVEVQEQNSSGTTDGVVPVQSNKLGFQSPALVNDHCCYELDFSTGSKSFGGGYGYNSLSQSVSSSSLDVGVVPDGSGSTLTDISNPYCSRSVCNGMESANQTVQLSAVDREARVLRYREKRKNRKFEKTIRYASRKAYAETRPRIKGRFAKRTDTEVEVDRSSLYGFGVVPSF

>Potri.004G108300

MLKQESSGGGGGDNRARVCDTCRAAPCTVYCRADSAYLCAGCDARVHAANRVASRHERVSVCEACERAPAALLCKADAASLCTACDADIHSANPLARRHQRVPILPISGCLHGSPVGPAAGETEDRFTTQEGEETISEEEEDEAASWLLLNPVKNSKNQNNNGFLFGGEVDEYLDLVEYNSCTENQCSDQYNQQHYCVPPKSYGGDRAVPIQYGEGKDHQQQRQYHNFQLGLEYEPSKAACSYNGSISQSVSMSSMDVGVVPESTMSEISISQHRPPKGTMELFSSTAIQMPSQLSPMDREARVLRYREKKKTRKFEKTIRYASRKAYAETRPRIKGRFAKRKDVEVEDDQMFSSTLMAETGYGIVPSF

>Potri.006G173600

MGIEVESLKNLTGGWSVAAKRCDSCKTAAAAAFCRADSAFLCLNCDTKIHHSGVNSKIMSRHERVWMCEVCEQAPAAVTCKADAAALCVTCDADIHSANPLARRHERVPVEPFYDSAESIVKTSSAFNFLVPGDQNGVSAYDHNDEIEGVSWLLHGNHTTHDLNTKINIENPVVKTGDMFFCEMDPFLDFEYQNSMDGRYKQSHGGGGAGADSVVPVQNKPAPLPVIDHKNCFDIDFCRSKLTSFSSYPSQSLSHSVSSSSLDVGVVPDGNSMSDISYPFGRSMNTYTDPSMPISGSTTNQAAAQLAGIDREARVLRYREKRKNRKFEKTIRYASRKAYAETRPRIKGRFAKRTEMESDMDTLYNSPSSVPFLADTHYGVVPSF

>Potri.006G267700

MASKLCDSCKSATATLFCRADSAFLCVSCDSKIHAANKLASRHARVWVCEVCEQAPAHVTCKADAAALCVTCDRDIHSANPLAQRHERVPVTPFFDSSSAAHGGGAAVNFLEYRYLDDVNGGDDVSREEAEAESWLLPNPGGGNTKGVDSLDLNTGQYVFGAEMHPYLDLDRYVDQKVEVEVQEQNSSGTTDGVVPVQSNKLGFQAPALVNDNCCFELDFSAGSKTFAGGYGYNSLSHSVSSSSLDVGVVPDGSTLTDISNPYSRSVSNGMESANQTVQLSAVDREARVLRYREKRKNRKFEKTIRYASRKAYAETRPRIKGRFAKRTDSGVEVDRSSIYGFGVVPSF

>Potri.T094400

MGIEVESLKNLTGGWSVAAKRCDSCKTAAAAAFCRADSAFLCLNCDTKIHHSQVNSKIMSRHERVWMCEVCEQAPAAVTCKADAAALCVTCDADIHSANPLARRHERVPIEPFYNSAESIVKTSTAFNILIPGENGVSGYDQNDDVEGVSWLLQSNHTTHDHNSKLQIENPVVKTGDMFFSEIDPFLELEYQNSIDASYEKIHGGAGADSVVPVQTKPAPLPVINHESCFDIDFCRSKLTSFSYSSQSLSHSVSSSSLDVGVVPDGNSMSDISYPFSRSMNTTTDPSMPLSGWTANQAATQLAGIDREARVLRYRERRKNRKFEKTIRYASRKAYAETRPRIKGRFAKRTEMESDMDNLYNSPSSVPFMADTQYGVVPSF

>Potri.002G214500

MGYICDFCGEQRSMVYCRSDAACLCLSCDQIVHSANALSKRHSRTLLCERCNSQPALVRRVEERISLCQNCDWMGYGSSTSASTHKRQTINCYFGCPSVSELSSKWPFILDFPSGGGSTCEQELGLMSIAENSTKNTLGPTENTICHNASGIVGVNDRCEIAKSGVWHEASSVPESSSVPNNLDQPTRSPNSSLPKLYCPGTKCPARYEDADLYEDFNMAEMDLNLENYEELFGVTLNNSEELFENGGIDSLFGTKDMPIVDSNCQGAFAAEGSSVGLVNTIQPTCSNAPSTDSMISSKTEPILCFTAKQGHSSLSFSGLTGESNAEDYQDCGASSMLLMGEPSWCPPCLESPLPSANRSDAVKRYMEKKKTRKFEKKVRYASRKARADVRRRVKGRFVKAGDAYDYDPLSQTRSF

>Potri.008G125200

MEKVCEFCMALRPVVYCNADAAYLCLSCDAKVHSANALFNRHLRTLLCDSCRNHPAYAQCLDHRMLMCLGCDRCLHEVSSHHQKRLVSSYLGCPSAKDFASLWGFEFGDLDKSIVKDQLVSTPCSSSVQPSASKFDIPGKSCQQIGRSSRKSRVIHSTLVSGAESDVGSGNQRPELWSQNMQDLGVCEDIICHDDDYIIPDVDKTFCNFEEFFGGDQDPIGAFLDENDFSCSFIEKDMPPEKSNNSDGRARKDASVTSSVYISCSVHIDNDKDPSNQAYNFPGSLDPAQTIRSPYSRYSISSHDAESRSNEYLDSELSPYISNGEASCYSPDLEDAHTEARENAMMRYKEKKKARMQDKQIRYTSRKPKNDVRKRGNG

>Potri.001G323500

MIMEKCLNSKQKKDCFLFLPCEFCNSKAAILYCRADSAKLCLPCDQQIHSSNTLSLKHVRSQICDNCRAEPASIHCSNDNLFLCQDCDWDSHNSSFSVSSLHNRNPVEGFMGCPPVVELASLFGFDFKSDFFVDSDPGSCSFEQEAVNFQDFAVSSDDFSVLSSSGKSRQEVYKQLVEMGKRGMVRVNGDGAELGPDTPPSRCAVQWNLESLELENGDEELLHQQTPFTSLLMLPNHVDASENDCVSDLGFMWDCNYTHQGAQAWDFQLGTSLDCTIPGPQEEGYDVKDPGFMVKNYVDFTEDGAFATQKVLDDGHVTSCCSSTCEDNLSKNSCSNQQLSRYKPPTENCNNTPLIGLSPESMPGEPNAHIQVMEQPSLTWFETLNEVRQKGDAGLFAQNRGNAMLRYREKKKNRRYDKRIRYESRKARADTRKRVKGRFVKAIENY

>Potri.002G208100

MKPRQQQKRLCDYCNDTTALLYCRADSAKLCFSCDHEVHSTNQLFSKHTRSLLCDACHASPVSIFCQTEHSVFCQNCDWERHSLSSLSSTHIRRPIEGFTGCPSGNELMTILGFEDLGLKKSLFFSEESDGFLGSELDDGCSDLFLWDSPAVCIDDLIVSSDSGSNFQTLGVPPLPKNRNAACGQHKEEILCQLRELVRLKPDPYGNADIDPANILQSLDADPQPPNLNTIGDPGAFTSYKENLPDWLADYGEAANQVLFLSTLPSSNFEESCAVPDKEFNIIGSASHVHDDHAAEPQHLTIETLPALPNVVTHELNSQERDSAISRYKEKKKTRRYSKHIRYESRKVRAEGRTRIRGRFAKMDH

>Potri.014G134601

MTESLEPQHHQQQEPKQQTTTIMNPKPQQKRLCDYCNDTTALLYCRADSAKLCLSCDHEVHSTNQLFSKHTRSLLCDVCHTSPVSIFCETEHSVFCQNCDLERHNLSSFPSTHNRRPIEGFTGCPSGNELMEILGFEDMGLKQSMLFSEETDGFMGSGLDDGYSDLFVWDSTAVSIDDFIMSSDSGPNLQALGVPPLPKENGEAANQVSFPSTLPGSNFEESRAVPEKEFNISDSASHINDGHEAEPQPSTIGTLPVLPNDGTHELSSQERDSAISRYKEKKQTRRYDKRIRYESRKVRADSRTRIKGRFAKLDH

>Potri.010G125100

MICNKSLANAVGGRTARACDSCIKKRASWYCAADDAFLCQACDSSVHSANLLARRHERVRLKSASLKSSDAGSKDNSMPSWHGGFTRKARTPRHGKPVSRSKIEETTRNIPIPLVPEVGSDEISLEDNEEEHLLYRVPIFDPFAAELCTSTTVSNEAGAVVPAGGTDTDQRAADSSGTESKVLLGGSEGKDVESLHGFLPSDMDLAEFAADMESLLGRGLENESFGMEELGLMDCKEENELGVKGYPLGNGKVKVEEEEDAGMEEKAVRECHADIEIDIAKDPPFELSFDYDSPATCVEEDEKVGIEEGDLKNSDGEYEDDGGAKKKRRTLLSLDYEAVMTAWASQGSPWTNGYRPDFDADECWPDADCMGICGAQLHHPYGDVSGLGAHPAALVDGGREARVSRYREKRRTRLFSKKIRYEVRKLNAEKRPRMKGRFVKRTTFAGK

>Potri.015G054600

MISGRKAANAMGSKTARACDSCLRKRARWFCVADDAFLCQACDASVHSANQLASRHQRVRLETASSYRISSSLNTDQDYSPPAWHQGFTRKARTPRHNSNKSLLVQQLLKDDREKVLNPLPLVPEIGSEEEPNMAPDENEDDQLLCRVPVFDPFAAKMCDIVTSEDENMVVEVYGQEGACGLDNLPGFLPSDMDLAEFAADVENLLGREADEEYHDTKDLELLDCSKGEDEGQFCFADKVVKMKDEQEMETIIDCHFDQDFNMARESLLGWNFDYETLVDGDEEVEEKKVPVPETEMMNSTGYKEMKRNVSLRLDYESVIIAWANQGCPWTTGSRPELNPDDSWTDSMGACPKDVNNPYGGLGSHTRGGDGEREARVMRYKEKRRTRLFSKKIRYEVRKLNAEKRPRMKGRFVKRTSLMGTTDFP

>Potri.008G120400

MICIKSSANAVGGKTARACDSCIKKRARWYCAADDAFLCQACDSSVHSANPLARRHERVRLKTASLKSLDLCSKENSVPSWHGGFTRKARTPRHGKPVSQSKIAETIRNIPIPLVPEVGSDEISHEDNEEEHLLYRVPIFDPFVADLCASTTISNEAGAIVPAGGNDGTDQRVADSNGVESKILIGAIERRDVESLPGFLPSDMDLAEFAADMESLLGRGLENESFGMEELGLMDCKEEKEFEVKGFPLWNGKVKVEDEENASVERKAVRKCYAGIETDMAKDPIFELSFDYNSSATCGEEDEKVGIEEGDLKNTRGEYEDDDGAKRKILLSLDYEAVMTAWASQGSPWTNGNRPDFDPDECWPDCMGICGAQLHHPYGDMISGLGAHPAMVDGGREARVSRYREKRRTRLFSKKIRYEVRKLNAEKRPRMKGRFVKRTTLAGK

>Potri.011G105400

MKIQCNVCEAAEAKVLCCADEAALCWTCDEKVHAANKLASKHQRIPLSTSSPQMPKCDICQETAGFFFCLEDRALLCRKCDVAIHTANTHVSVHQRFLLTGVKVGLEPTDPGASSSSGKSPSGEKKTLETKSRPVSRRGTLLPLANPCNQVSTVNVCGVGDFGPAKLPYSGGSATSSISQWHIDEFLDLPEFNQNYGYIDNGSSKADSGKRGDSDCSAILRSTEEEVDDEECLGQVPDSSRAVPQIPSPPTASGLYWPKSFHNHSETAIFVPDICCSVVQNCHYSEQRGTVSKRQRQL

>Potri.017G028300

MKIQCDVCEKAPATVICCADEAALCAKCDIEVHAANKLASKHQRLLLQCLSNKLPPCDICQEKAAFIFCVEDRALFCRDCDEPIHSAGSLSANHQRFLATGIRVALSSSCSKDTQTNSSGPPNQSAQQTPMKIPAQQTSSFATSWAVDDLLQFSEFESSTDKKEQLELGEFEWLADMGLFGEQLPQEALAAAEVPQLPISPPTNVNSCRPTKSSMPHKKPRIEISDDDDEYLTVPDLG

>Potri.002G028200

MKIQCDVCNKEEASVFCTADEAALCDTCDHRVHHANKLASKHQRFSLLHPSSKNFPICDICQEKRAFLFCQQDRAILCRECDGPIHTANEHTQKHNRFLLTGVKLSATSAVYISSSSVTNSGGDLVPDSKSQQQQQQQQSIKKPVFDAPVNSNPPTVPSTLSTNTEVNKGGDNLVTNEGFGSTTSSTISEYLMETLPGWHVEDFLDSSTTPFGFCKIDDGLLPFMDAHDLESNMSSFSSESLGLWVPQAPSTPYTSQQYYYPQLVGQSGFKEIKETTNMKANRRLADDVFTVPQISLPANISSKRSRPLW

>Potri.001G384000

MKIQCNVCEAAEANVLCCADEAALCRACDETVHAANKLASKHQRVPLSTSSPQIPKCDICQEAAGFFFCLEDRALLCRKCDVAIHTANTHVSAHQRFLLTGVKVGLESTDPGASSSPGKSPSGEITTLKTKSCPVSRRGTSMPLASPCNQVFPANVCGVGEFVPAKLPYSGGSAASSISQWQIDEFLELAEFNQHYGYMDNGSSKADSGKHGDSDCSAILRSAEEEVDDEECLGQVPDSSWAVPQIPSPPTASGLYWPKSIHHSDTAIFVPDICGSAVQNHRHCQQRGTVSKWRQQL

>Potri.004G161000

MVMKIRCDVCDNVEATVFCCADEAALCDGCDHRVHHANKLASKHSRFSLVHPSFKESPLCDICQERRALLFCQEDRAILCRECDLPIHKANEHTQKHNRFLLTGVKLSASSSLHTASSTSTNNFDSNINTTSNRNHQPYLKNSNEILSSPSVETASATTAYTFEENHTCDLF

>Potri.004G162600

MRMLCDVCESAAAILFCAADEAALCRSCDEKVHMCNKLASRHVRVGLADPSDVPQCDICEKAPAFFYCEIDGSSLCLQCDMIVHVGGKRTHGRYLLLRQRVEFPGDKPGCTEEQGQQPLDDNETRRDQNQPPKLTARENQQNHRASPVPMVENNTDSDGKMDNKLIDLNARPQRVHGKNPTNQLTLWMVVREASSFH

>Potri.005G117100

MRTLCDACESAFAIVFCAADEAALCLACDKKVHMCNKLASRHVRVGLANPSEVPRCDICENAPAFFYCETDGSSLCLQCDMTVHVGGKRTHGRYLLLRQKIEFPGNQPQPEDPAPQPMYPGETRRGQNRPQKATSGENRQNRQASPVLMSVTNSDGHDKVDKNMIDLNMKPHRIHEHASNNQEQ

>Potri.005G234500

MKIQCDVCSKEEASVFCTADEAALCDTCDHRVHHANKLASKHQRFSLLHPSSKNFPICDICQDKRAFLFCQQDRAILCRDCDGPIHTANEHTQKHNRFLLTGVKLSATSAVYMSSSSSVTSSGDLVPDSKSQKQQQQQLIKKPVSVAPVNSNPPAVPSTLSANTVINKDGDNLVTSEGFGSTTSSTISEYLMETLPGWHVEEFLDSSSTTPFGFSKIDDGLLPYMDTHDLERNMSSFSSESLGLWVPQAPTPPLCTSQQYYYPQLVGQSGFKETKESTNMKANRRLTDDAFTVPQISPPSNIGSKRSRPLW

>Potri.007G130100

MKIQCDVCEKAPATVICCADEAALCEKCDIEVHAANKLASKHQRLLLQCLSNKLPPCDICQEKAAFIFCVEDRALFCRDCDEPIHSAGSLSANHQRFLATGIRVALSSSCSKDTQKSSLEPPNQSEQQTSKLPWQHASSFGSSWAVDDFLQFSDIEESTDKKEQLGLGEFDWLADMGLFSEQLPQEALAAAEVPQLPISPPTNVNAYRPPKFSMSHKKPRIEIDDDEYFTVPDLG

>Potri.007G015200

MRTLCDACESAAAIVFCAADEAALCLACDEKVHMCNKLASRHVRVGLANPSDVPRCDICENAPAFFYCETDGSSLCLQCDMTVHVGGKRTHGRYLLLRQRVEFPGDKPQPDDLHSQPMHPGETRKGQNQPPKATAEEKRQNRQVSPAPMSLSNSDGHDKVDKKMIDLNMKPQRTDHEQASNNQEL

>Potri.009G122000

MTMKIRCDVCDKVEATVFCCADEAALCDGCDHRVHHANTLASKHSRFSLVHPSFKESPLCDICQERRAVLFCQEDRAILCRECDLPIHKVNEHTQKHNRFLLTGVKLCGPSLYATSSSASNCDANINTTRNRNHQHYLKKPISASNEIFSSPSVATASPPTAYSYDDNHVSGGGSVSTSSISEYLETVVPGWRVDDFLDPSFTSNNSFSKGWNAPYI

>Potri.009G124400

MRTICDVCESAAAILFCAADEAALCRSCDEKVHLCNKLASRHVRVGLADPSAVPQCDICENAPAFFYCEIDGSSLCLQCDMIVHVGGKRTHGRYLLLRQRVEFPGDKPGRMEEQGQQPLDHNETRRDQNQPLKLTARENKQNHRASPVPMVENNTDSDGKMDNNLIDLNARPQRIHGQNSTNQENHESSSAVPVGSFKREPQK

>Potri.012G069600

MLDWFNNFVSIHEIDEDGAHLQIIQKIMSDEEKAFEYQIVTAAWEEIDYAKPCLLCDQHVHSTNLLLRKHVRSQICDNCTSELVSVRCVNDNLILCQECDWDVHGTYFMVVPQLLIWLPYGVLIRKRRSWGR

>Potri.014G170600

MGYVCDFCGEQRSMVYCRSDAASLCLSCDRNVHSANALSKRHSRTLLCERCNSQPALVRCAEERISLCQNCDWIGHGTSTSASTHRRQTINSYSGCPSASELSSIWSFVLDFPSRGESTCEQELGLMSIAENSTTDSWGPTENTIGQNASGVVEVNDRREIAKSGVWLGSSSIPESSSVPNNLDQTTRSANTSLPKLCCPGTKCPAPYEDADLYEDFNMDEMDLSLENYEELFGVTLNNSEELLENGGIDSLFGTKDMSGADSSCQGAVAAEGSSVGVVNAVQPASSNAASADSMMSNKTEPILCFTEKQGHSSLSFSGLNVESSAADYQDCGASSMLLMGEPPWCPPCPESPFSSANRSDAVMRYKEKKKTRMFEKKVRYASRKARADVRRRVKGRFVKAGEAYDYDPLSQTRSF

>Potri.017G039301

MESVCDFCGVARAVVYCKPDSAKLCLHCDGCVHSANFLSRRHPRSLLCDKCSSQPAMARCLDEKMSVCQGCDCRANGCSILGHQLRALNCYTGCYSLAEFPKIWSSVLQGPSSGALDSGRDSLNSAPINENCISWLEQGENEGSFGLITGKLNELESCSKLESWRGPPSVIMPNPTYMPCSTDQVPLLPEVSNLPKQGCSIFKDIGLSDGEDLCEGLNMDDIPLDFENSDGIFGCPESHNRYPFEDVGKDCMLMEKNLSVTESNGPIENAIEVSSSGQQDCVAFQSSCVSGPVSAIQNISSNANCSMFTNPSCSRNLHLGFPAGTGQVHPSMSLSLSNIIGESSAADYQDCGLSPIFLTGESPWESHLDASSPHARDKAKMRYNEKKKTRTFRKQIRYASRKARADTRKRVKGRFVKAGEA

>Potri.002G220666annotation

MGYICDFCGEQRSMVYCRSDAACLCLSCDQIVHSANALSKRHSRTLLCERCNSQPALVRRVEERISLCQNCDWMGYGSSTSASTHKRQTINCYFGCPSVSELSSKWPFILDSPSGVDLLLYCPGTKCPARYEDADLYEDFNMAEMDLNLENYEELFGVTLNNSEELFENGGIDSLFGTKDMPIVDSNCQGAVAAEGSSVGLVNTIQPTCSNAPSTDSMISSKTEPILCFTAKQGHSSLSFSGLTGESNAGDYQDCGASSMLLMGEPSWCPPCLESPLPSANRSDAVKRYMEKKKTRKFEKKVRYASRKARADVRRRVKGRFVKAGDAYDYDPLGQTRSF

>Potri.001G061800annotation

MVTHKSKTKETVPCDFCSEQTAVLYCRADSAKLCLFCDQHVHSANLLSRKHVRSQICDNCSTEPVSFRCSTDNLVLCQECDWDAHGSCSVSASHDRTTIEGFSGCPSALDLASIWGFDLEEKKPEPLIENWSNNSCGVIHDLVNEPWVYDKSSGNLTFQDLMVPNENNNNGNRNIDNVMIFGNVTKSPSCGKYKHVIYKQLFELFKRDLIGGGVEGEGCGFGDGEGCGFGGGDGGGETLVTQSRSGWRSGVKGVQFGNGNDGGFGDDNVVVCGGNGSGGNVRGEQLLQEQRPFTSLLMLPTEVDVKSNGRVVGGDITWDSNAKAHGTQVWDFHLGQLRNHDESGQLEIEYVANDAGFVIKDFGELMKETSSTSPKMLGDMYQMNCSTAHDDMTSFNVDMELLARNRGDAMQRYKEKKKNRRYDKHIRYESRKARADTRKRVKGRFVKTTEAPDS

>Potri.007G121200

MESVCDFCGVEKAVVYCKPDSAKLCVHCDGCVHSANFLSRRHRRSLLCDKCSSLPAVARCFDEKLSICQGCDCSANGCSSLGHQLRALNCYTGCYSLAEFSKIWSSVLEGSSSGGFDSGWDSLNSAPINENCISSCLEQRDNEGSFGLFTGKLNELESCSKLEPWRGPPSIIMPNPNYMPCCRDQVSMFPEVTNLPKGCSIFKDIGLPDGEDLCEGLNLDDIPLDFENSDEIFSCSETQSKYQFGDVGKDCMLMEKNLSVTGSNGPIENTIEVSSSGQLECAAFQSSCVSGPASAMQTISGNANCSIFTNPSCCKNLNLGFPAVSGQVHSSMSLPLSNIIGESSAADYQDCGLSPLFLTGESPWESHLDASSPQARDKAKMRYNEKKKTRTFSKQIRYASRKARADTRKRVKGRFVKAGEAYDYDPLLSSNF

>Potri.011G125400

MKGCELCGSSARMFCESDEASLCWDCDEKVHSANFLVAKHCRTLLCQVCQSPTPWKASVSKFAPTVSICESCFTIPNKTKETEERMKGCELCGSSARMYCESDQASLCWDCDEKVHTANFLVAKHCRTLLCQVCQSPTPWKASGSKFAPTVSVCESCFTIPKNKRHFQSENVMTSDQESQGGGNDLDESENDQESDDDDHTDDDSDEDEDEEEEDDDGDNQVVPWSGPTASSSPSPVPPVASSSSEEETSCAGRNGFLKRMRESNVDLDSDDEIGCSSSHNIGGRILSNDGGNSLSSMRPWKQARTSVNVEEDG

>Potri.011G039700

MKRCELCDSLAKVYCESDQANLCWDCDANVHSANFLVAKHSRSLLCHVCQSLTPWTGTGHKLGPTLSVCNNCVNNSVCREERGREDDEEGDNDDGDDDDDDDDDLDREEDGDEDEDEENGDGGNDHGGEDDEENQVVPWSSTPPPPVSSPSNDSEECSSRFCDSDGGISKSRRAFSSKHRRGTVP

>Potri.010G251800

MAVKVCELCRREAGVYCDSDAAYLCFDCDSNVHNANFLVARHARRVICSGCGSITGNPFSGHTPSLSRVTCCSCSPGNKELDSISCSSSSTLSSACISSTETTRFENTRKGVKATSSSSSVKNIPGRSLRDRLKRSRNLRSEGVFVNWCKRLGLNGNLVVQRATRAMALCFGRLALPFRVSLAASFWFGLRLCGDKSVTTWENLRRLEEVSGVPNKLIVTVEMKIEQALRSKRLQLQKEMEEGWAECSV

>Potri.013G150500

MKKCELCKNPARTYCESDEANLCWNCDTKVHGANFLVARHARALLCQSCQSLTPWKASGSQLGHTVSVCERCMISNENNNREIQEQEENGSSDDIEEDSDSESDSDTNDDDGGDEDDGDGDNQVVPWSPTTPPPPPSASSSCSCTSSDDNDADFVESVNVVSFKRQRLQDDLNRSYSLRMYGNTATPVECYSSSTRGSLKEKRKELDPTV

>Potri.017G039400

MCRGNQERSNQGSSCNKEAVSPNATSRFVCCELCGSRATLYCQADHAFLCQKCDGWVHGANFLALRHVRNMLCNTCQNLTQRCLIGASTEVMLSTIVSWRERRDRNSNLEKKCSGSLKKPFSFL

>Potri.001G414700

MKGCELCGSSARMFCESDQASLCWDCDEKVHSANFLVAKHCRTLLCQVCQSPTPWKSSGSKLAPTVSVCESCFAVHKNNKKQLQDLNVMASDQESREAGNDYDESENDREFDDDDTDDDSDEYDEEEDGDNQVVPWSGLTASSSPSIAPPVASSSSSEEEISCAGGNGFLKRMRDSNVDLDSDDETGCSSSHNLRGGSMSTEEGNSLSSSRPWKQARTSVHVEEDGHDGQAMSRSSAIIDSLKRLQKDLVANGENASAAILGICKLSRDQSL

>Potri.004G027100

MKKCELCDSFAQMHCESDQAILCSACDAYVHSANFLAAKHSRTLLCHVCQSHTPWIGTGPVLGATLSVCNSCINNSSCTDGKGSENDQIANNDDEIIANMMKTMMVVKTAMKTMKKIK

>Potri.004G027000

MKRCELCDCLAKLYCESYEANHCWQCDTYVHSANLLAAKHSRTLLCHVCQSLTPWTGTGPKLVPTVSGCNSCVSNLSCKEERSSEGDQVGDNIDG

>Potri.004G026900

MKRCELCDSLAKMYCESDQASLCWDCDANVHSANFLVAKHSRTLLCHVCQSLTPWTGTGPKLGPTLSVCDNCVSNSSCREERSTEDDKDVDNDDDDDDREDDDDDDDDREDDSGEDNENGDGGNDHGSEDDEENQVVPWSSTPPPPCFKFF

>Potri.008G007000

MAVKVCELCQREAGLYCDSDAAFLCFECDSNVHNANFVVSRHLRRVICSACNSLTGGSFSGTAPSLRRVTCLSCSPENKELDSISCSSSCSSTLSSACISTTETTRFENTRKGVETSCVTNIPARFSGGRLKRSRNLRSECVFVNWCERLGLNGNLVVQRATRAIALCFGRLVLPFRVSLAASFWFGVRSCGDKSVTTWQDLRRLEEVSGVPRKIISAVEMKIEHALRSRRLELHKNMEEGWADSTDCSA

>Potri.007G121100

MCRGIHQERNNQGGSCGEEVVSSKATSRLVCCELCGSRASLYCQADDAFLCQKCDKWVHGANFLAQRHVRCMLCNTCQNLTQRYLIGASTELLLPTIVSWRERRQCNSNLEKKSSGSLKMPFLFL

>Medtr1g013450

MATKLCDSCKSTKATLFCRSDSAFLCLTCDSNIHAANKLASRHHRVTLCQVCEQAPAHVTCKADAAVLCISCDHDIHSANPLARRHERVPLTTTFNHQNSQQQSFFSENDHDATTEEAEAASWLLQTPSNPKFPDLNYSHYSYPEIDDFVTVNTKTDLPEQNSPGTTADGVVPVQSHSKTATEHEHEHYSDINIDFSNSKPFTYNFNHTVSSPSMDVGVVPDGNVMTEISYCSYQTTATETAPMTVAVPMTAVEREARVMRYREKRKNRRFEKTIRYASRKAYAETRPRIKGRFAKRSDLNMNLIAEDEYGVVPSC

>Medtr3g105710

MASKLCDSCKSATATLYCRPDSAFLCGACDSKVHAANKLASRHPRVTLCEVCEQAPAHVTCKADAASLCITCDRDIHTANPLAARHERVPVTPFFESNTSHSVKSLNNNNNNYDAVKDEAEAASWLISDPKADLNSSPYLFSDSEAIPFMDLDYGVIEHKNDGVVPVHGNFDPFVSAYKNNNVHLHTELETPSQSQISQSVSSSSMDVGVVPDANTVPEISNCGYGTVAVDREARVMRYREKRKNRRFEKTIRYASRKAYAETRPRIKGRFAKRTDAVDSISGYGVVPTC

>Medtr4g128930

MGIERGGLKSLRGGWSVPPKLCDSCKLTPAALFCRSDSAFLCINCDSTIHSANKLSSRHERVWMCEVCEQAPASVTCKADAAALCVTCDSDIHSANPLARRHERVPVEPFFDSAESVVKSSSAAAAAAASFNFVVPTDDGYGQDDAEAAAWLIPNPNFGSKLNETQDIKTREMFFSDMDPFLDFDYSNNFQNNNCSNAMNDSVVPVQTKPTPAPMMNHNSEGCFDIDFCRSKLSSFNYPSHSISHSVSSSSLDVGVVPDGNTVSEISYNFGSESMVSGGVNSSNQGVQGATQLCGMDREARVMRYREKRKNRKFEKTIRYASRKAYAETRPRIKGRFAKRTEIDSDVDRLYNPADPLSVPSSMLMDCPYGVVPTF

>Medtr7g018170

MLEQDFLTTTSATATVRSAGTWARTCDTCRSAPCAVFCRADSAYLCAACDARIHAANRVASRHERVWVCEACERAPAAFLCKADAASLCSTCDADIHSANPLASRHQRVPILPISGYLYGPPATLLGAEDEGFVRGGCEVEEEEDEGVDHDMEDENEAASWLLLNPLKNNNNNSNNNISNDHNQVANNGYLFSGEVDEYLDLVDCNSCGGDENTFTTNNTHHHDEYSQQQQQQDHYGVPQKSYVGDSVVPVQQQQVQNFQLGLEFESSKAGFSYNGGSISQSVSVSSMDVGVVPESTMTYSRPPKGTIDLFSGPSIQMSSHFSPMDREARVLRYREKKKTRKFEKTIRYASRKAYAETRPRIKGRFAKRTDVEAEVDQMFSTSLITEVGYGIVPSF

>Medtr1g109350

MEKICEFCTALRPLVYCNADAAYLCLSCDAKVHWANELSGRHLRTLVCNSCCCDLAYVQCLDHKMLICRDCDQKLHDRSSPHRKRSVKSFIGCPSAKEFATLWGFEFKEIENSVSQKDQFASISSVSTDVNVIKHHFRNSVASTTSGDKHDKGSSSQLGQILYSDQERQTILQQIVDLKRFQLIEERDHSTKINGLQVDEKFNQQAQKSKYFGINLLGEDNSIGELNPETFSSAFSQLDNLSSSSVMDLPLHGELFWTAKSSLQSNQLWPQNIQDLGICEELVCRDDFNIPDVDLTFQNYEELFGGDQDPIRVMFGGKDVSYSSLEKDLSVDNSDIDNPSTMEV

>Medtr2g088900

MGLSDDRGEYIGAKTTSFKGLPQSNVEEAKGLKEAINCLGNLRFPSLSIKLDWGFHILGLDKEQSSSDYRSCVPSNPIHSDPIHSISFRFIPSGAFMEALCEFCGVARAVVYCKPDSARLCLHCDGNVHSANSLSRRHPRSLLCDKCNFDSAIVRCVDHKLSLCQVCDWNTNDCFVLGHKHVLLTFYTGCPSLAELSKIWPHLVDANSSNAAWESPSTSSLPKTESSSGRGQHLEQQPEKNGFVGLANDKLGEGDTCVKYEPWIENSPIIPSNSNCTQYYKDQPFLFNQDSNQQKDLIIHEGTSLCEGFNVDDIQLNFESADEIIFDCSQTATKYNHEDGGIECLLMDKNIPVTKCSSHIETAVEASSSVQQDCMIFPSSGAGGSTNLMQGFNNSANCALMPPSCNRSMPLEFPQSQTHSGISIQLPNINGESNVAELLDCGLPPVFHPGESHWESNLEGACPQARDKAKMRYQEKKKTRTFGKQIRYASRKARADTRKRVKGRFVKAGEAYDYDPLLSDH

>Medtr3g082630

MGYICDFCGDQRSMVYCRSDAACLCLSCDRNVHSANTLARRHSRTLLCERCSSQPALVRCSEEKVSLCQNCDWLGHGNSTSSNHKRQTINCYSGCPSSAELSSIWSFVLDIPSLSETTCEQELGFMSINENRSAWVDPKNQNVSDSDKATDLPDLDKSFAGTSSMPESSKEPRMLDRPDGSTNECVPKLYCPATNCREASDDDDDLYGDFDMDEMDINMENYDELFGMALTHSEELFENGGFNSLFGAKAMSAGDSNCQDANAAEGSSIGHVNAAQPACSTAASADSILSTKTEPNLCITAKQSQSSLSFSGINEDGGAGDYQDCGASSMLLMGEPPWLNTCPENELQLQSANRCSAVMRYKEKKKTRKFDKRVRYASRKERADVRRRVKGRFVKAGEAYDYDPLSQTRSY

>Medtr5g069480

MGSLCDFCGDQRSLVYCRSDAASLCLSCDRNVHSANELSKRHSRTLVCERCNLQPAYVRCVEEKVSLCQNCDWSAHGTNPSSSTHKRQSINCFSGCPSASELSSIWPFFSDIPSTGEACEHKLGLMSINENSDNSARVPPESKNVSGSAQVADLPSKNKSGVDTSSIPESSAKPRILDQAPGSSNEFMPKLLCPSRKAPALCEDDKLLDDFNIDEMDFELENYSELFDFALNHSEEFFENGGINSLFERKDMSASAGDSNCQGAFAAEGSSARFVSAIQPECSNAASADSILSTKTEPVIYFTERQSNLSFSGVNKDASAGDYQECGTSSMLLTGEPPWCPPCPENSIQSANRSNAVMRYKEKKKNRKFDKKVRYASRKARADVRKRVKGRFVKAGETYDYDPLSQTRSC

>Medtr7g083540

MSFPCDYCDTRSAVLYCKPDSAKLCLVCDQHVHSANALALKHVRFQICQNCKNDAASVRCFTENLVQCHRCDWNSHGGDDDDSTSSSFHHHNRRRIEGLTGCPSVHEIVSTLGLDLKPNDAVFVAEFEGPVVPVVKNRDEVYEQVVEVAKRKRNLEEDQNELRFNDCCNDVDDLLLLQQTPFTSLLNFSSEFDVGVKRNSNDYGNESGLLLWDRNPSYQPPQVWDFQLQKSRDMTYDGVENASLSIPKSLQDVHNMNCSTLGDDILSRNNQSDQSSSSHVKKKVESNKKTRDGLSSESKLIESITYSGADSVPVMEHLLSGSENVSNINAKISLEEHTRNRGDAMLRYKEKKKTRRFDKHIRYESRKARADTRKRVRGRFVKATDDIQEG

>Medtr7g108150

MGGSPRNPNPNPSHKLVRPCDYCGHSNAVIYCRADSAKLCFSCDREVHSTNQLFSKHTRSLICDSCDDSPATILCSTESSVFCQNCDWENHNLSLSSPHERRSLEGFTGCPSVTELLSILGLQDIGKKSLLLPQESVGDGFVGYEIEGLSDMFVWDAPSFVSLDDLISSSPSSHNYRAMEVPPLPKNRKAACGRHREEILNQLREMTKSEPYDPEEYIPPANLSTSFDCDVKADIVPSNEWLRESSEPMYQVVPVDTSFKAHTEEISVKHSVSSVGEPHTHCNNGGTPSEPLNHCNNGGTPSEYVKSETLSTTSKAVPPPYELASQERDSALLRYKQKKKTRRYDKHIRYESRKVRAESRTRVKGRFAKIDH

>Medtr1g110870

MTCSSKNVANAVGGKTARACDSCITKRARWYCAADDAFLCQACDSSVHSANSLARRHERVRLKTASYKSINGDEFFNCGPFSGFTKKARTPRQGKHKSKSSSSSEPARNNNNIPFHLVPELGFDEVNSNSIEENEAQLLYRVPIFDPSIADLCTSPPSVCSTEGGLGVVVVASAFAPDVKNNESESRVQLGSDNNYEMESFHGLLPSDIELAEFAADVESLLGRGLENECIGMEELGLIDTKHEESEKWKCGGKVKEEGEECYEVVEGDNMMEIGKESSFELNFDYDDSHETCEEVKEKCGEQNENNDYNKGKRKISLQLDYDAVIIAWDSQKCPWTNGDKPILDADENWPDCMGTFGTEVHYAYGEFGGYGCHPVMVDGGREARVSRYREKRRTRLFSKKIRYEVRKLNAEKRPRMKGRFVKRASFAVPTFPLLK

>Medtr8g104190

MIIDMKGDADAGALGAKTARACDSCLRRRARWFCAADDAFLCHGCDNLVHSANLLASRHERVRLQTASAKVTTTAQAWHSGFTRKARTPRHNKNSSIQQQQQRLKEKVLFNTSFLPLVPELGGEEEQGQELLVDIDEADEEQLLCRVPVFDANPFDLETCTVKNDAVDFEEMCDLDSFCEFDVDLAEFAANVESLLGVGSSEIQENSSGQVFDYKQENEMDASKSEMLKVKDEELDDLESVFDMTSDDVFHWNIDNNDVSLAQQEKEYMPLSNSSVGYSESVITKEETKRERFLRLNYEEVITEWSRQGSPSPWTTANPPKFNCDDDSWQNLLGSSGVEGEVRSLRGQLMGSGGDGGREARVSRYREKRRTRLFAKKIRYEVRKLNAEKRPRMKGRFVKRTACFAGGATSFPTNYH

>Medtr1g023260

MKIQCDVCNKNEASLFCTADEAALCIDCDHRVHHANKLASKHHRLALHNPTPKQHPLCDICQERRAFVLCKQDRAILCKDCDSSIHSVNELTQKHDRFLLTGIKISTTNSSSSSSSSTPSSATTKSNHIPSSSLIEKSTTPSPTSMEEGSGGSTISQYLIETLPGWQVDDFLDSSSVPFAFSKGDELFNAGIEENLDSFFPNNNMGIWVPQAPPPSLYSSSQIMMGQSETTKKGSNNKSTINKSRLRDDHDSNIFTVPQISPVANSKRTRYLW

>Medtr2g073370

MKIQCNVCEGAEAKVLCCADEAALCWECDEKVHAANKLASKHQRVPLSASSSHMPKCDICQEAFGYFFCLEDRALLCRKCDLAIHTANAYVSGHQRFLLTGVKVGLEATDHGTSSNSLKTDSGEKVSDTKSSSVSRKVSQMPQSSEYNEMLPTEAGGFGDFPPAKVSYGGGSNSGNMSQWTIDEFFGVNDFSQNYNYMDGSSNSRADSGKLEGSDSPILRSNEEEMEYDDYMDRVPDSSWTVPQIPSPPTASGLNWPKNPRYSFDNALFVPDIGFTSMQHPQNSSNFSRPRNHH

>Medtr2g099010

MKIQCDVCEKAPATMICCADEAALCAKCDIEVHAANKLASKHQRIHLQSLSNKLPRCDICQDKTAFIFCVEDRALFCEDCDESIHLPGSLSANHQRFLATGIQVAMKSNCAKDDEKTHLEPPKRSTHQVSLETTSQQVPDFTPPWGVDDLLELADFNSHDKKDSMQFGEMEWFTEEGLFGDDFTQEAMAAAEVPQLPVTHASNNYSSYRNSKSHMSNKKPRIELIRDYDYDDEDEYFTVPDLG

>Medtr3g113070

MKIQCDVCNKRQASLFCTADEAALCSTCDHRVHHANKLASKHRRFSLDHPNSPNHFPLCDICLERRGFVFCQEDRAIVCKECDLKVHGVNEHTKKHNRFLLSGIKLHSPAPPPTLHEETGNFTISEYLINTIPGWKFEDFLDSPSSSVPSHELQHQNHIVHDDANYHIHEENILVSFSSESMRRICVPQAPLYYSEKMDNRSNTVNFSSSLGDFTVPQITIPPN

>Medtr3g117320

MKIQCDVCEKAEATMFCPSDEAALCHGCDHTIHRANKLATKHTRFSLVHLNSKDYPLCDICQERRGYLFCQEDRAILCRECDLPIHGANQHTQKHNRFLLSGVKLSSNSLDPDSSSTSIVSEARNYSSRSKANIIPTSVSNENASSSCMVEDNMASDTGSVSTSSISEYLIETIPGYCFEDLLDASFPPNGFCKKQKQNHYSAFQYQDIHVNKFFFSSSNVCTSSSG

>Medtr4g067320

MKIQCDVCEKAPATVICCADEAALCAKCDVEVHAANKLASKHQRLLLQCLSNKLPKCDICQDKPAFIFCVEDRALFCKDCDEPIHVAGSLSGNHQRFLATGIRVALASSCTKDNEKSQVEPSNPDTQQVPVKVSPPQQVPSFASPWAVDDFLELTGFDSPDKKQSMEFGELEWLSDAGLFNDQFPQEGLAAAEVPQLPVMHASSVYPYKASKSYMSYKKPRIEVRHEDDDDEHFMVPDLG

>Medtr4g071200

MKIQCDACHKQEASLFCPADEAALCNQCDRNIHYANKVSAKHKRFTLHHPTSKDTPLCDICKERRAYLFCKEDRAILCRECDIPIHEINKLTKQHNRFLLTGVKIGASSSCSNPTISNGSELRTSSPRPSSFSSENNSCSQSSFKENMVCDTVSTSSISEYLIETIPGYCMEDLFDASFAPNNVFCNKDYYEQNQDLQVINMSDWVPQSQVRFPQLSANSNVPN

>Medtr5g021580

MRTLCDACESAAAIVFCAADEAALCRACDEKVHMCNKLASRHVRVGLASPSDVPRCDICENAPAFFYCETDGSSLCLQCDMIVHVGGKRTHGRYLLFRQRVEFPGDKPSNADNPASQPLDPGDIKRGQSPLPKQKMGEKQQNHRMPPVPTSEPNADGNSKMENKLIDLNMKPNNRIHEHASNNQVRVHGK

>Medtr2g011450

MKKKCELCKSPAKLFCESDQASLCWECDAKVHTANFIVTKHHRFLLCHICQSLTPWHGTGPKFVPTISLCNHCVAVTNNNDEDNDQDDDDDDTEDDDEENQVVPWKSTTPPPVSSCSSNSAN

>Medtr2g089310

MKKNCELCKLPARTFCESDQASLCWDCDSKVHAANFLVERHMRTLLCHACQSPTPWKASGARLGNALSLCDRCAGGRKLHADANANANAGTSAEQQDESEGDNDNEEDTDYDSDEDDDDVDGDEDGDNQVVPWSSTAAQPPPASSSSGSDESVSVSKCNDNHDEVISKLVTTISLKRRRLDHDFEVSDSKNWKNQRREVDEVDLIGCDGEPSSKAAMRRYSDLNGSEHSQPHD

>Medtr4g008050

MKMKCKECELCNQQASFYCPSDSAFLCRSCDVAVHGANFLVARHLRHILCSKCDGFTEILISGTALHHRLSSTCRSCSPENQSSGEPSSQSSSSCESCVTEKKKTKSRKIMKSFSVSNSVTDDISPAPGNKNMKKKMIGTEDAGSVAEEIFSKWRRELELDFPVNGDRVAVEAMAVCLRTWKLLPVKVAAASSFWFGLRFCGDNSFATCRNLMRLEKISGVPAKLILATHVKLARVFTQHLELQEGCDES

>Medtr4g046640

MCKATIEEKKRGGFCKRFPPKQVASLCETRSCSISCELCGLQASLYCQADNAYLCRKCDNLVHKANFLALRHVRCFLCNTCQNLTRRCLIGASLEIILPATVSTIDNLPNNNSMHRNCSKQKSGTHFQFL

>Aqcoe6G248700

MRTLCDVCESAAATLFCAADEAALCPSCDDKVDLCNKLARPHVRVGLAAPNEVPRCDICENAPPFFYCEVDGSSLCLQCDMIVHVGGKRTHGRYLLLRQRIQFPGDKAGPIEEQGMQPMDQGVIRREQNEPPKLTTKENMRVSPVPESNAKNEEHRKIENNMIDLNTKPNRTHGEVSNDKEMDTTNRSSHKSADVVPYGYFDGEAQN

>Aqcoe7G302400

MKGRLCELCNGEAWLYCVSDSAFLCLNCDARVHQANFLVARHVRRTICCKCKYFDGNSISGVGFRRHIEPICQSCLSNCSSDSNSSCTSSSTTCIISSVNQSSNQLNSSSKIVFSGGNPFLPTKYSDEVSKNNKKKKKKNRKIVETRSLNVDFKEEDILVNWYRKLGVKCNSNISITLALHAISICLQKFTNLPFRVSLASSLWLSLKLAKEKPAVATCQNLKKLEQISGVPAKLIILAEKKISIVLKKNEGKSSSKLDDDDLQEGWAECSD

>Aqcoe3G384500

MMKKEMRSGSGGGGWSRVCDSCRAAACAVFCRADSAYLCTGCDARMHGANQLVSRHERVWVCEACESAPAAFTCKADAASLCTTCDADIHSANPLARRHHRVPILPISGCLYGPSANYPSRPLGSVADMEDGFLTSEVGEELEEDDDETSSWLLLNPVNPVKNSNPSNGFLFGGEDEYLDFEEYNSCTENQYQDQYKQQQQQQQNSFSIQHNQVKNDGNDSVVPVQYGTMDQHHHQHNLHLEMDHEASSKPGFNFTTSLTHSVSMSSMDASIVPDSTMSETSNMHSRTPKGTIDLFSSPPLQMPAQFSPMDREARVLRYREKRKTRKFEKTIRYASRKAYAETRPRIKGRFARRTDVEVEVDQMFATSVMAESGYGIVPSF

>Aqcoe1G378700

MGPLCDFCGEQRSMVYCRSDAACLCLSCDRNVHSANALSRRHSRTLLCERCSSQPATIRCLEEKTSLCQNCDWDGHGASTSSSSTHKRQTIKYYSGCPSAAELSRIWSFVLDFPAMEESNCEQRTGLMSIDENVSSSCWGSPGKTGMLENANGGNEVDKFDSWMASSSMCVPNPIPSQVDQTAGSVDSSTSKKYTSETKGLGLCEDDFCEDFDVDDMELNFENYEELFGNSQTNSEQLLENGGLDSLFGVKDMSVADSSCHGGYDTEGVSAKKGKVVQPTCSNGVSTDSAMSAKTEPNLYYHARQGHSSLSFSGLTGESSAGDYQDCGVSPMLLMGDSSLYPLGAENTHPSFSRDSAVMRYKEKKKSRKFDKKIRYASRKARADVRKRVKGRFVKAGDAYDYDPLCQTKSY

>Aqcoe3G174700

MGERIPCDYCGDEKAILFCRADSAKLCLLCDLHVHSANDLSKKHLRSQICDNCNSEPVSIRCSTDNLVLCQECDWDAHGNCSVSASHERNPIEGISGCPSAIELASIWGFDLVEKNTTNNNNSSSVSSLPSIQEDSNSVFSNWSSLDSILYIDSWIGSNSIENNNLQELIAKEEEENSNSNLNRNSASFLYPNVPCAQIPQASSKKGQNPSCGKHKQLILKQLLQLFKRDLNRDAEDLSPGTPTQQGSAEGIDLQKGVCVGDLMDANQMMQQQLPYTSLLMSSVETDHKGNVRLLEDHHLWGCNPPSSESSQIWDFNLGRSRDHEAPNQLEDGYSTNSEGFMIQTYTDFTKKTPSETTKVLEDVYGINCSTTGHHMKSDYNTSKKPASSRRTLTSESNNLNLPGADINYMEQPLLLRGENARTTTKVDMELLAQNRGNAMLRYKEKKRTRRFDKHIRYESRKARADTRKRVKGRFVKASETPDVDASR

>Aqcoe7G312300

MLKKECELCSLPAKTYCESDEASLCWNCDWKIHSANFLVAKHERSLLCHICSFPTQWKSSGSKLGSTLSICEKCVKNNCKRRSLVVHQQQQDESEGDNEDTTDDEEEDSEDEEDEDADNQVVPWNSTDQTSSSSCSNSEKSSSIKPVSMKRIREHEEDLDNQDYKCCSSSSSGNKEEEVTSLDCLRPLKQRKIDEEELILSGVNESESESDLLVQSLKKFHQKMLLVQ

>Aqcoe2G434900

MKIQCDVCEKAQATVICCADEAALCAKCDVEVHEANRLASKHQRLLLDSLSTKLPRCDICQEKTAFIFCVEDRALFCQDCDEPVHSAGSIASNHQRLLATGIKVALGSKCAKNAELIRSEPPNQNSQPLISKSLPTPQPSTCTSPTSWAIDDLLQFSDFDTNDKKESIGFGDQFEWFTDIGLFNDQVPQETLAAAEVPQLPVSKPSNLPLHRSIKSYMPPNKKPRIEVSDDEEYFTVPDLG

>Aqcoe4G302300

MGHLCDFCGQQRATVYCQSDAASLCLSCDQNVHSANALSRRHSRTLLCQKCNSQPASVRCLHHQLSYCQTCDWNEHCTSTSPTTHERQTVNCYLGCPSAEQLSKILSFVPDVSSANEYACEQKMGLMSIDENTVTNYLVPPESNNTVDPCVTGKRSNDGNEGQFGVWTGSSSMPTMNPLQCTQLAGTVADSTTTKPTPLGLEGLGIIEDDFCNDFNVDDVDINFENYEELFGICQNHREQDIENGGIDNLFGMPDMSAVGSGCQGGFFTKGSLAGKDSLMQSLCRDTVSADSAMSAKTEPNFCYSAMPAHSSLSLSFSGLTGESAAGDYQDSGVSSMLLMDEPPWFPPGPDNSFPSASRDSAVMRYKEKKKARKFEKKIRYESRKARADVRKRVKGRFVKSGEAYDYDPLCQTRSC

>Aqcoe1G210500

MKIQCNVCEAAEANVLCCADEAALCWACDEKVHAANKLASKHQRVPLSNSSSQMPKCDICQDTVGYFFCLEDRALLCRKCDVSIHTANAYVSGHQRFLLTGVKVGLEATEPVGSSTKEASNSVKSPAAKVSQSVRGSSMPINGENNEALSRQVVGVVPVPKLSFSGGSTSESIPEWPLDDLLGITDFSNSYVFMEHGSSKADSGKLGDNDFSLTPKGANEDFDLDEYSGNVPEVPWIVPEIPSPPTASGLFWPKSFRNSPSDNSVFVPDMCCSSPQNFQRYQPDALSKRQRR

>Aqcoe7G333000

MKIDEGSGKVFAASWSLSAKPCDSCKSSSALLFCRADSAFLCIGCDSKIHTANKLASRHERVWMCEVCEQAPASVTCKADAAALCVTCDRDIHSANPLARRHERFPVVPFYESAASAIKSNAVNLLVPDESEHNNEDLDEDDDDEEDDDDHQNNNKNHHHAATGGVVVTCREEAEAASWLLPNPNHNNKLMETPDLNCADQYFFSDVDPYLDLDYSSPLTGRYHNSSGTGNDSVVPVQVPDHHHHQSAFIHHHEMDFCRSTKPSYNSYTTPSLSQSVSSSSMDVGVVPDGNCNAMTDISNPFVNGTTTTTTAEFNGTVPTGVVNQASQLTGMDREARVMRYREKRKNRKFEKTIRYASRKAYAETRPRIKGRFAKRTEIEAEVDRIYSSASCLMNDTSGYGIVPTF

>Aqcoe6G256000

MKIQCDVCNKEEALVFCSADEAALCNGCDHRVHHANKLASKHHRFSLLNPANKDAPRCDVCQERRAFLFCREDRAILCRECDIPIHTANEHTQKHNRFLLTGVKLASCNPTSSSNGCDDFQKVNNKSQSFIKSTPPPPTSIENSSTTSIPSKGGGGGSSGGHVASSTTTTTSGDGSTSSIAEYLIETLPGWHVEDFLDSSSVAPHGFCKNDDLMPFLDAEVETNLGSFSSDDLCMWVPQAPNPCDHPSHTINEKKTGIFKEIKDVIPIKMNNKRWSDHGFTVPSISPPSNKRGRSFW

>Aqcoe5G178200

MRTLCDVCESAAAILFCAADEAALCQACDDKVHLCNKLANRHVRVGLADPSAVSCCDICENAPAFFYCEVDGSSLCLQCDMIVHVGGKRTHGRYLLFRQRIEFPGDKPAHLDNRTLQSMNLVENRREQTHLPEQTLTEKPQNMRVPALVPKRDEHEKMDNMIDLNARPNGIHGGHPSNDQSKNVMNGYHEESASIVPRGPPERGPEK

>Aqcoe4G244800

MCKGREEEHRNLQIDLNLQPLDDFSTEEISSHRNMSTIFCELCSSKASLYCEADDAFLCRNCDKKVHGANFLAFRHVRYLLCSSCQNLTVLYFSGSSFGMVLGAGRRSSHRDSKADEECSRFAKKRTTLSI

>Aqcoe3G039600

MSSNKKTANALGGKTARACDSCMKKRARWYCTADDAFLCQTCDASVHSANSLARRHERVRLKTASLKPAEENPVENSHPAWHHGFRRKARTPRHGKPSQHQNKIEEPIVNLPLVPEIGSDEVSIDENEEHLLYRVPIFDPFVAELYASTTPNETTNTVDSATPTTTTPIGAESKIVLHDYGHDGTVDFENLPGLLPSDMDLADFAADVETLLGRGLDEDSFCMEGLGLLDTKEEIYYDHHSDKVKLEEDEEDVKNMVACHQFGSDLDLSRETLDLNFDCDSPTTGEEEEEKVNMTINYGYKGEEKISLRLDYDAVIAAWASQGSPWMNGDRPQFNPNDCWPDSLGTYPAEAHQMYGDMGSMGGHVAMGGDGGREARVSRYREKRRTRLFSKKIRYEVRKLNAEKRPRMKGRFVKRTSFAVGCGGGSSSFPF

>Aqcoe1G384600

MKECELCNKPARMYCESDQASLCWECDFKVHSANFLVAKHSRNLLCHVCQSITPWKASGSKLGPTITACERCANRCGFQENINEEEQEDDEDDQDSHGGNEEEEEEEDEEEEEEEVNEDEDGDNQVVPWSCNLKKNNNPSPPPVVSSSSSEDSDSSNRFLKRTRDNVDREN

>Aqcoe2G310900

MVDSSHSTTPQTQNNELNEEKNTPHQFKQRLCDFCGESKALLYCRADSAKLCFNCDKQVHSTNQLFTKHTRYQLCDFCDSNPSSILCSTENLVLCQNCDWDLHGKSSSSSSSVHDRRSIEGFSGCPSGVELCSIFGIEDVKDKCFMNCGGGYLEESDLYSTPPDVGVDGFSDLLVWETPPIFSIDDLIGSNDSTQSFQAIGVPSLPKNRNTTCGKYKEEILSQLREMVKLENCFIDDGEDLESLLGFQPLVPELNSQPGKVSRSSEHRVEPILFSLNEKNEDKWEKNSCDMAVQVPCPSACLKSYAVTKSSIDLVDDGTDANGDHDSYLQPVIIETPCIVPKVALHKPITQNRDSLISRYKEKKKTRRYDHHIRYESRKARAEGRSRIRGRFAKIEH

>MA_128658g0010 high_confidence

MKMKVQCDVCENAEATFLCCADEAALCSVCDNKVHAANKLASKHQRVPLINPSSQSPKCDICQEKTGYFFCLEDRALLCQCDVSIHSLNNLVAAHQRFLVTGVKVGLEPSNTISSSTNTFAQSSDATHQKPQTLKNGPREVSATSHQGIQKGAGGGGMSRKGTVSEYFSELLPLLRMDEFLNLPELDNGYSFDEAGSSRADNSNFVEEWTANSLSMEEVNAENCLAQVPEMPSPPTASGLYWPRRVIQHPKEGKRREDLAIFGIDDASLVPDIGCRSSPPDSPLSKRRRLHA

>MA_10426894g0010 medium_confidence

MKVQCDACQSADASVFCCADEAALCMKCDSKVHDANKLASKHRRLSLLEPSSSSSTDSLRCDICQERRAFFFCQADRAVLCRDCDLSIHSANELTAKHNRFLVPGTRVSLKPMETLSCPEKAVATVTKALMPPAQRKRWPLSPRH

>MA_7292g0010 high_confidence

MVKEEDCKVPKEAGIVKEFQAWTMPKPCNVCRIASASLYCRADSAYLCSGCDVKVHGANKLASRHERVWLCEVCEQAPAAVTCKADAASLCVSCDADIHSANPLARRHDRVPIAASWLLPNPKSSAEGAKNCDDGGSCFGVDAGPPVNKAAGGYFSVVDLFPDVDPYLDLDYASPLEATGGTDSVVPVQSNVSSQDGAVSTPSDCFDTEKVTYSYTTTTSLSHSVSSSSLDVGVVPDATLSDMSRPLNRGVFELANPGVMNVGIQYVQLDREARVLRYKEKRKNRKFEKTIRYASRKAYAETRPRIKGRFAKRVDADVAQMYTSAELSYGLVPSF

>MA_10433513g0010 high_confidence

MRTLCDSCEAAAAQFFCAADEAALCAKCDEKVHGCNKLAGRHVRLQLRESWSVPRCDICETAAAFLHCSIDGSSLCLQCDMEVHVGGKRTHVRYLLLGQRVELLNGNHIPNGNHIRDEQGNPKTMDTARAWQNKYCQEHRSNGDPSHKTNISNGNIHNVVSCNKENVQSNGQKD

>MA_54929g0010 high_confidence

MVKEEDKDWHTVEDLHRGSHVDHKEFLRGIGGWRMSMPKLCDVCQVSNSVLYCRAHTAQLCLVCDVKIHGGSKASLCHERVWVCEVCEQAPAVVTCKADAAALCVSCDTDIHSANPLASRHERAPVIPFYECPNMPNNNTATNANNDNLDCNVLLNEDGGGDDPLKHDYVDDDYDDYDDDENDHNNLLNHQEEDNDAEICCAEEAATASWLIPEANRNNLTNINGGNSEGEDKMVKDKLKFKAYMQSIDFLQDVENYVDLEYLGTTTTITTPTTPTAHMGADSMVPVHTPEVIEHSSTKVSVETARSLDVDAASKCNYVYRTTSLNHCVSSSSIDVGIVPDSNTTTDISTPYHDPRGVFEIPPRVVHPGGHVEVMGREARVLRYREKRKNRRFEKTIRYASRKAYAETRPRIKGRFAKRTEVEVEQIYSSSLLPDQGYGVVPSY

>MA_10192193g0020 high_confidence

MATGVGGRAARPCDACAQQRAQWYCSADDAHLCNACDTQVHTANTVACRHERVSLSTTTTAVPFPGAPLACWSSSWLRKRSLSGQRHGRSNPNGERPLGSTPATDAVLEDDADKEEEPLSWLWSSDEVDTPLSWLCNHHVKSEASEDVGYVKTELEPEGLLDLNFDFCLVNEREMEKMKVHEESKVHLRLNYEEVVSAWSDRGSPWAQAHPVMNPLDDTISDLAGDGGSSVFGLVPDLTSLGCGQSGREHVIGQVLVMIHGGDGQRPRAGGREACVMRYREKRQSRLFYKKIRYHVRKLNAENRPRMKGRFVKRTADASYESLAL

>MA_8519g0010 medium_confidence

MATVCNSCGKMRSTVYCREDAVSLCVSCDRDVHGCNAIPKRHLRTLLCDGCSVQPAAFTCKSQKLSFCHNCDRQGHSNSPQHKRTSINYYTGCPSAAELSEPWSLGLEGSLDSLLAAGVLAGFAQNLVSCREPSASPRVDLPSSSQRNYNVKTTIFDDSEGSAGPMVEMKIGSAVDISF

>MA_25074g0010 medium_confidence

MATMCELCRKIRSTVYCRAEAISLCLSCDRDVHGANGISKRHLRTLLCDRCGVESAAFNCNNHKLSFCHNCDRQSHSNSPQHERTSISYYTGCPSAAELAKLWSRELDGLGDDEKQGPDVPGIGLDGSLDSLNLSAGVHAGFKHNLGSWMEPVASPSVAVSSNGNTMVAMQGHPMDM

>MA_10006362g0010 medium_confidence

MCDCCGEFRSTVYCRADSASLCLSCDEHIHGANALSKQHLRTVLCDGCSVEPATFSCNDHNLSFCHNCDRQSHSNSPQHRRKSLSYYTGCPSAAELAELWDCELDRLGGDDQQGPSPCNR

>MA_10427343g0010 medium_confidence

MAILCEFCGKITPTVYCRSDAASLCLSCDRKVHGANALSKRHFRTLLCHGCTIEPAAFSCNNHNMCFCHNCDRQSHSSSPQHNRTSINYYSGCPSVAELAELWSCELDGLGCDVIKGSDVPGIGWVDSVGLDGALDSLNMAAGVHAGFKVNLGLWMEPFASPMDDMPLSSQRNSEVNTMIFDSEGSAVTMAKMKVSVLISCPEILPYLTIVPGSAKDIRTANQDFGLRCQEALPNLPIDPGSAKDIRTAHPAFLLWWCGALFFKLNYIIITVLA

>MA_10430891g0010 high_confidence

MFEIITGPPRDLKMAKTMKECELCELSARLYCESDEASLCWDCDAKVHSANFLVARHCRSLLCQICQSVTAWRASGAKPGLTVSVCERCAGGSRAKSDGDDDEGNNDENQVVPLASGYSGPPLSSSSADQEESSGDERHCAESPLSIAPPDAASHKRPHESLALYTNEGDCGCSSSGVNNSNNNLSGCEDDATSSRIRVLKKRKADQGLKDDLVKTLEIILCMRTNYRAPAPAIKGDEWHCAEIPFSIAPPDVASHKRPHESLALYTNEGDCGCSSYGVNNNNNNLSGCEDDTTSSRIHILKKRKADQGLKDDLVKTLE

>MA_4244045g0010 high_confidence

EFCGKITPTVYCRSDAASLCLSCDRKVHGANALSKRHFRTLLCHGCTIEPAAFSCNNHNMCFCHNCDRQSHSSSPQHNRTSINYYSGCPSVAELAELWSCELDGLGCDVIKGSDVPGIGWVDSVGLDGALDSLNMAAGVHAGFKVNLGLWMEPFASPMDDMPLSSQRNSEVNTMIFDSEGSAVTMAKMK

>MA_332069g0010 high_confidence

MFCESDQANICWACDAKVHSANFLVARHVRKLLCHVCQSPTEWQASGVSPCPSLSLCNKCFSAKNSARKHQLDENDNQVVPSTPPVQAASPSSSSGGNGGDARGQHSGA

>MA_90307g0010 high_confidence

MDNICDFCQELCPTVYCRSDRARLCLSCDRHVHNANALSRRHLRTLLCDGCNLQPSAVRCHTKNISLCENCDWNIHGSSPTGSQHKRCAINGYTGCPSAAELSGMWGLGDFPNTNVSDSYGESSKSVSGLLAINENIANSNNRWATSRNDSSLDCVVAGRTDNTGIALKFDILMETSPSAPSASSMLYAADGLDDQHPVQTSGALEQQLKQNPVVVQQLLDLQKLPPQSSSQLQKYPEVQPRICTQVQMESKLVEPSPIKQMEQDQAENQNQQRQQKDQIQVQQHNNQERQKPSVSPRQAEPQQLKSDANIERFMQGDSFWHCNPASEISQLWDPHMQDLGVCEDGDPFNDFNMSDVDLTFENYEDIFAGSQDQSVSLFGDVGAACSSTEMNDSFADSSDHIESAPKAFSVSPVGCVLPSSHFPGPVGTTYPVSGTSVQGMLGPSRCGDISMGFSARKPHSSLSLSLSGHSGESSAADYQDCGVSTRCLKGDPPWGPTSPESVFSQARGNAMMRYKEKKKLRMFEKRIRYASRKARADVRKRVKGRFVKAGEAYDYDPLSVTGNY

>MA_3117g0010

MLCQDCDESVHSPDTLAAKHQRFLATGIRVVALNAQSLDSEGLSEFNKQPTSISNSTAPAHAGPRMGSAHSSSAKPIPLGEPCWSVDELLPLSDFESKGDPDGLGYFDWEHEQEEALACRVPQLSPTGPAGGKPSFTVKGKSKTEIPTVPDFDGACIVPDSDEACIVPDMGSLDTHEFYPTPPPKRMRRSSFEAW

>kfl00103_0190_v1.1

MPKQCDACQSQKATVYCRADVAFLCNGCDRVIHEANPLARRHERVFVCEVCETLPAVVNCKADSAFICNNCDNSIHGANGLARRHERVPVTPFYECQDAVKVAHIHYPPSLVPEGLALEVPQFCREDDDSALEAASWLLNKPDEGGSRGEEGVDKPDFCPPSQDGQVPSNLCRQPCVKQTKPFELPMLMQVNGGARVKQETGMFNPMAEKHRGVKAGPRFGVNPGLMPRAEALVPSLGGSGEVLEQRVPVVQKGKSSFAVPQLAHSVSSSSFDVALVPDSSLSEVSMPSPDADVQKGDTAKGFEMPQRLLHVGGAPPMDREARVNRYKEKRKNRTYEKTIRYASRKAYAESRPRVKGRFAKRESNVSITQGIPVSHIPVSQPFPSYDYSMYGMVPDGLPYGMMPVSYY

>kfl00001_0850_v1.1

MKIQCDACENAVAAVMCCADEAALCVDCDKRVHAANRLANKHQRVPLLAQPKEGSKCDICQEGAAFFFCLEDRALLCRDCDISIHTANDLARKHTRFLLTGVRVGLENLGADEQAEAPSTSAPSKVLGSPFPPKPTSPVARVPVYQPAEQKPSKKKSATAVPSSPVPEAFSGGGSGFSGGFGGGSSGAGPSSRGGGASSSGLRRGQSVNVNELPDSTVPVWSVDEILGIPNLADGYSLNDIGSTKGMDALMGDFDWTADLGMFDDLAFAESMHEVPNFDYSPSPGLVPSNSPAVPRKGGGSALKGKARSSYDMGPAVVPDSDDLFVVPDLGLQVPPSKKRRSNLDWH

>kfl00120_0220_v1.1

MLNLGRATPGKGKAVAFVDLDSDCDDPPGQRDADSEEDAALLSVDWKSPRASWGSGWSSAATPASDHREKSGGRRGEEAGRGRSLDLDWLTGDNAPDPAELDRLVDTVTATAEPGSAPSASGRKQGGEVGGVRRVTVVVDETERQTNSNPNSVYLELHRQHEALHAVAGVALAAEKHKLETGDYLWLAETDEGSAQGGQPCDECEAQRAALECAECEQALCSDCCASLHSGGARRAHRLTSLPPPWQCEECEGASAAVSCGECEQALCASCSAKIHSKGARARHAVAPLSLAPMQPTSRVSTPPDTHDSVERSGRALMVASSSVYASLRTVDLGPRVALEQTRGERDRGPVEESLYLGLADPGGAAGRAGEREAFWVAVSGASSATCPEPDLQAAAVLFLTSLPDPPCPSAHSPGKILILENVLHHCRAMAAAAERARLNGAAGPGEDGDQELRAFLGCHDLNEEMIRWIVRIMFSRGWNVQFTGNQLESSLLFRHLAKAYVK

>kfl00451_0030_v1.1

MRALCDVCEAAPALLFCAADEAALCRPCDEKVHGCNKLASRHVRLELAEARAVPSCDVCEKAPAYFYCEIDGTSLCLQCDMDVHSSAGKKLHERFLLMGQRVEIPMAPGKPEGGDKVPVLQEQRLHQNGNAHPSHRHGKPDEEMREARVADSSGTAMATAIGGEPSGDADTSKQSLLDLNRCPPPRSKGEHSKQAAVSSGGHRNGPAAQAGQSRPPMNDQDRNEVG

>kfl00214_0150_v1.1

MARHAEEERDASKNAKQHRSAERKTDVEVEPTKKHSTKEKFHKEEGKDKRKEKSKDEKRESRHRDESEPQKTNAVEGERRKASKELMCEVCEHREATMWCGDCGDIFYCRKCADADHSRGKRKEHLPLRTAEKAPQRAATKVVKCEEHGEILRAYCETDRKAVCSLCLHIGNHKGHSAVELDKAVRAARDSIRKEAGDLILERQKVDTFLDEMEEMAESLKKNGAELREHLRTACDSFRAVLTEREAALLVQVDEAHALRIAAARRSATPATTASLDLALEADPFINLIRESLVFSSSASANRASSVIARLQQAQSQKASLTDNTVVVLPRGCLEVRNVALLESLAYSIMKVVKRNTVDYNLFWIFELDILDHRRVCCSRFSNALVKSSLAALNELRSRAR

>kfl00279_0070_v1.1annotation

MEVPCEYCGARKATVFCRADNAKLCLSCDRSVHEANALSLRHERTLLCDACGNAPASFRCSEENLSLCLECDASSHSAVPASHHKRVKFDFFTASSPALPGCPSANELAVLWGCRLGSLEDAPVKAEPRSPALPPADSHPRAAPGWGSSLPPKASRLGMELARGASQGAGWPEAKAGPPGAGSPALYGAPAAPPQHQQQQGPAATSQQQQQQQQQQQHQQQQQPPQQAFLGPPTPSYYPPLEQQQTGMPGMGGQAPFLAPQGAQPAYPGLYQQGPYPGQLYQPFPPPTMQQGYPGPPRYGEYQDPYGRQYAPQYAGQPFQHGYGMPSPRPRMQPAKAMGTPFGGRPGLESPSGMLSQVKTSASPLPPIAEAAPSAAPGPSGLDPGSGLGRAATFIPTSGQGHTPGPGPGPTPAAATFPGQIRDSAVLRYKEKKKTRRFEKTIRYESRKARADIRTRVKGRFVKAEDER

>kfl00252_0110_v1.1

MDFSATGAQDADPRGRENEGSPEFAAGGVAKDPPPKCAVCKEAEAEFSCASCVASFCSPCWEVPHKWGDNQNHAKVVVEKPTCPCEGSFSVCPHGKDAVGYCGECDQALCQVCWDLIHTKGTRVTHNLLAPLSRSDYRAEQLALQRAVKSGPLASLAKFDVKKAAPWVTSALKKSRSVADAKRFGLGGASRSTAAGSSRYAAIQLQTSKDAAAQLKAVCDEPGFGVQDDSTIKSYALDRQADVLAVASAMKVVFETSPAGTDARNRTLCDKVDDYEKAIRIARASGKGVQLSKDQRGERLERMTTTIALVQSQLDTLLKQLNWKEGVAFLIRSPPYFDVLINLQTEAERAAKLAKDLGVRIAKLGPGIVGALLEAKEKEVAKAQKGKGTGVESTKKRTVKAGMVQQAKRKKIAGGPRGKGGKAEANIQEMEELVKEMQAKLKKKKAAVQDIESSEEEPSEGDEEDEAEVEQEDNEGEREEGEEEETN

>kfl00212_0040_v1.1

MDKLPEVEEGARETEESGTADGTVPIMKLCQVCNASPAVVYCAADAACLCQPCDVEIHSANALSSRHERVSLLGVKDAVLCEAQAETGCVSAEHTVTLSSSSKSSRAKRARTQGATSSGGSGEDLGMSPSSLDAQEVPSRWPGEEYSSYGGLTENDLIAMEMPDRGAQDSILESLGDEMCEMEESMEGIMYGIDGITAIPQLDLGSLGRSSGELRNGSEHADAGAGSDSSSMGAGSMSHYDQYGPEVKYEYGLEGLQEMRHRGAEAEQQASYESSYRLDDQAAFQLGLRLGESNAGLDLAAGKGPKMGLSLDFSDILSAWSDKDVWADGRYAQLVPDTGSPLDCLAPAAYDHGVVPDLGSQRKREGGTYAGATTGDEGPVLNRKERVERYKEKRRTREFCKKIRYEVRKANAERRPRIKGRFVKRSECIAMGLSMF

>kfl00104_0060_v1.1

MFDVFRPAASGHPAAAVAPSYLAAVRPESETACCGPSTSGASKQCELCPKAARVFCDADSAYLCSACDVRVHTANFLVSRHVRTLLCIDCQGETTWRACGGRPGKEHHRCKACEETARRLKADEDQCVPSGGVDASWLTLGPILSHPLPSRGAVAEDRASGRWGRDTVREEQAAISPSGSSAVSGLSEEREATVTPSSVLIEVPRGERKRAREEGDDGPVAASHKHQKVDHAHRPQVAALPPRPQRSAPLRPNTEPAMTNAAIVERLERTLARWYVSLNLNSPETIPLSMHIFRKVLQSLQPNLLAAEAMRVTLAGCLWLATKVDTAQVKVPKASEVAVAAGVQASRLSAVELQLLTLLDWRLLDGWVGSGEVLDS

>kfl00016_0200_v1.1

MTRPQSASTRSQHPTACIECEDELATSWCEECRTSFCEECDLHQHHPKSLRREHVRTPISPPEPPAAAASRAGLLSKAFATEETAWSAEGDEDSAEDGPADSGLTSANQGAFSSGGKLESGSSHHALKSKRKHKKKRGKSRGHSKDRVEGGGVEAEGHEDGRAEPSPTKGLDCQVRKEKDGDSGGSSSVEHLRERPVWKKRKSSLKGSHEAAGEGPRQSLHVHWSPDVKDH

>Bradi3g19010

MGSEGGSSGGDTSPNAGCAVCGVAAAVLYCAADAAALCTPCDAAIHAANPLASRHHRVPLSSAAAMAASGVYDYDSLFAGDDEAGSTLTPPGQGSPNSGSSSFSSNGGSERSLFELLSDVDLMATDAAAAPLWMMMQPGMDAAWSLSEGVVVPSGAGAGFPSLVGGGAMVAADREEKVRRYREKREKRKFHKTIRYASRKAYAEARPRIKGRFVKRAAAEESDHDSTNGGAEGSKFWLSFSDADSRDGSVGFQAPPAYGVVPSSF

>Bradi3g35856

MVSNLRVTIPVRVLRNGGRRGIQKINPWVCEASPVAVTCKADAEVHRPNLLAQRHLRVPISPILGFHGGAMRAPELEEEEALTNNLHVEARGRRRRRCQAAAAFQLRLLHQLPRAQC

>Sobic.007G092000

MGSEGSTSPGGGGGAACAVCGGAAAVYCAADAAALCGPCDAAVHAANLLASRHERVPLSMAAVAAASGVYDDLFAPDDIDAASSWAAAAHASPQNGSSSTSFTTSDGGAEGRMSLFDLLSDVDLAAACVTGGGVGGYLPDGVAPVHHHGAAPLWAQPGLQAAAWTATWSPADAAAAAAVVGVPGAAAVVAAAAEREARVQRYREKRKNRKFQKTIRYASRKAYAEARPRIKGRFVKRAGTSSSSSSGAGGTSDTNDATDAASKFWLSFSDDGRDDGVGFYVDAGAYGVVPSF

>Potri.004G108320

MLKQESSGGGGGDNRARVCDTCRAAPCTVYCRADSAYLCAGCDARVHAANRVASRHERVSVCEACERAPAALLCKADAASLCTACDADIHSANPLARRHQRVPILPISGCLHGSPVGPAAGETEDRFTTQEGEETISEEEEDEAASWLLLNPVKNSKNQNNNGFLFGGEVDEYLDLVEYNSCTENQCSDQYNQQHYCVPPKSYGGDRAVPIQYGEGKDHQQQRQYHNFQLGLEYEPSKAACSYNGSISQSVSMSSMDVGVVPESTMSEISISQHRPPKGTMELFSSTAIQMPSQLSPMDREARVLRYREKKKTRKFEKTIRYASRKAYAETRPRIKGRFAKRKDVEVEDDQMFSSTLMAETGYGIVPSF

>Potri.017G028301

MKIQCDVCEKAPATVICCADEAALCAKCDIEVHAANKLASKHQRLLLQCLSNKLPPCDICQEKAAFIFCVEDRALFCRDCDEPIHSAGSLSANHQRFLATGIRVALSSSCSKDTQTNSSGPPNQSAQQTPMKIPAQQTSSFATSWAVDDLLQFSEFESSTDKKEQLELGEFEWLADMGLFGEQLPQEALAAAEVPQLPISPPTNVNSCRPTKSSMPHKKPRIEISDDDDEYLTVPDLG

>Potri.018G096084

MGIEVESLKNLTGGWSVAAKRCDSCKTAAAAAFCRADSAFLCLNCDTKIHHSQVNSKIMSRHERVWMCEVCEQAPAAVTCKADAAALCVTCDADIHSANPLARRHERVPIEPFYNSAESIVKTSTAFNILIPGENGVSGYDQNDDVEGVSWLLQSNHTTHDHNSKLQIENPVVKTGDMFFSEIDPFLELEYQNSIDASYEKIHGGAGADSVVPVQTKPAPLPVINHESCFDIDFCRSKLTSFSYSSQSLSHSVSSSSLDVGVVPDGNSMSDISYPFSRSMNTTTDPSMPLSGWTANQAATQLAGIDREARVLRYRERRKNRKFEKTIRYASRKAYAETRPRIKGRFAKRTEMESDMDNLYNSPSSVPFMADTQYGVVPSF
