## Supplementary material for "BBX transcription factor evolution in the green plant lineage": Data S2.docx

>Vocar.0002s0569|Vcarteri_v2.1

MACVVCAAQASVYCENDKALLCKDCDVRIHMSNAVAARHVRRIPCEGGCSKGASLFCRCDNAYMCEACHCANPLAATHETEPTAPLPLMEQENAVAEQPHATGPCESVAQSAASPVAWFVDDEKPSLGSFEEPIMLSPAGSEAVVPVMSAPADDFTFTEPATFKEIKDKLEFESLEFDNSWMELNFDFTDILSDGPSDVGLVPTFDLEGVDPVADATVPSVAEEVVVAESDAATEIHRKRTAEPSDEEPAAKLPAISEAATTALGLHAAFQMTQPASLFFQSAVAQPSLLPPMVPPAVISPLATTAAAPPLQPPPTAVQSKSSAAYNTALAAGANLTREQRVARYREKRKNRKFEKTIRYASRKAYAEIRPRIKGRFAKKEEIEAWKAAHGGEDAVVPEILDGEF

>PNW82267.1 hypothetical protein CHLRE_06g278159v5 [Chlamydomonas reinhardtii]

MSSCVVCAAAAVVWCQNDKALLCKDCDVRIHTSNAVAARHTRFVPCQGCNKAGAALYCKCDAAHMCEACHSSNPLAATHETEPVAPLPSVEQGAAPEPQVLNMPCESVAQSAASPAAWFVDDEKMGTTSFFDAPAVLSPSGSEAVVPVMSAPIEDEFAFAAAPATFKEIKDKLEFEALDLDNNWLDMGFDFTDILSDGPSDVGLVPTFDAVDEAADAVADAIVPTFEEEQPQLQQQEPLVLAPAPEESAASRKRAAAEEAAEEPAAKVPALTHQALLQAQAAAFQAVPQASALFFQPQMLAALPHLPLLQQPMMPAAVAPAPVPKSGSAAASAALAAGANLTREQRVARYREKRKNRSFAKTIRYASRKAYAEIRPRIKGRFAKKEEIEAWKAAHGGDDAIVPEVLDAEC

>ABO97666.1 predicted protein [Ostreococcus lucimarinus CCE9901]

MPDVMCGTCAEAPAAVVRVESQTGVALCACARCDTRQTAKRGSGRSTTQRVGLRQASGNGSDELSCDVCQMNPAYVICHEDRAFLCRVCDVSIHEANATSRKHQRFLFANTRVELEAMGAGEEAGTRMSPSDSAAEHTVPQFEQEEVGRKRKYNRQQKASVPSEDATVPSIDDLAPGVFENFMTGLLGEEEGRKHMEKSTADENNFWGDIFSENWAAMDGMMDDELAVPNFDQQVPTNAMY

>CDF36220.1 unnamed protein product [Chondrus crispus]

MVRVCELCPEGKARQAVVYCPADACFLCSACDEEVHSANRLAGRHVRRAVATDDGGDGSSINDDSENALVPDVAELDEGEHSDPSSSEDMLLMSLPIQVPSFEDAAEYDFDFEGGLGGKMPALCAIDDDALFTGTKSLGKSFYGDISWESVVPENIEHVVPDVSAPGMGFFKREAVEVDAPMKVVSSSTAASSVGDLKSQPEVTSISVVNRPGTIVRIDGAVVSKAEPTLSLSSTGSSITGQKRSREEDESSGVSGDKTNAEKDEAERAAEQRKKRRMEALARFRSKRANRSFTKKVRYECRKQLADSRPRVKGRFVRKIEMALFRKYGALYREHLDELEGAKKEVKGDHRVPAI

>CDF40713.1 unnamed protein product [Chondrus crispus]

MTPRWCAACRAAPASIYCPSDTAILCTACDILVHEANQLARRHTRVPLNLFDNMGVPQVEHSPPTHFHGAQSTDKSSSAATAFDESDESSGGGVVPDTDDLYPEHHTFADTQVADISLSPPTKHQQPSSWAHEAQSAPHPVADTNTECLKTIAGAFVKQEIQTLFESSQRQSPPKATHANGLDFLWMRDSLPPALLCDGTSTTLTGTRHGFDSKQVLSAAQIALQNSLDQSLLDDVLGSSLSTDEDISIPGADILDGLKIEGAALTSAERSREQKRLDRQAALRRFRHKRANRSFRKKVRYACRKQLADSRPRVKGRFVGRAKTTSTGRVQKKSSAMGVTG

>CDF36563.1 unnamed protein product [Chondrus crispus]

MTRLFGVRAVCDGCLSAAATVFCVNHRTVVCFSCDDEQHASPHTASHQRVDLSSAVTALPFCERCDDAPAIVYCESEGITFCDKCDCVSHASQTSPPHCRTPISSTLRHRPVEFRGLPDNRTSSVYQVSSSPSFPASTRTKLSHLDSTKNGNDGRARRRAFMCAHVRRSSATARARSQTARADACACTSYSYAGKWRMRASSGDRAPRCTSPASRCTLLTDAPYKVDGSRPSFVTPVHIAPFRTTAALMTGAQMARWRGCRDRTAAEGGFGGGPRW

>CDF32176.1 unnamed protein product [Chondrus crispus]

MTRLFGIRAVCDGCLSAAANVFCVNHGTVVCFSCDDKQHASPHTASHQRVDLSSAVTALPFCERCDDAPATLYCESEGTTFCDKCDCVSHASHTSPPHCRTPISSTLRHRQVECRGLLDNRASSVYQVSSSPSFPASTPTKTISPGFCQERKRLPDAAEGAHVRSCAALVRDGAGALPNGQG

>XP_005705261.1 zinc finger protein CONSTANS-LIKE 2 protein [Galdieria sulphuraria]

MKASQVLASQLCELCQSANSSIYCEEDDVFCCDQCDVEYHTSTADKECHLRISFLCERLSAFRGPCNPALYQVPTCSKEM

CHLREVITHMIAEHCSKNMKHDEADILSSEKVVLESPHMKPVSGNSSILDSPTFSLSSTLDSWVDEEYPFSSFSSSLEGDDMGSLFFQVANYEDSDKCANVDSFLTSEDGTDKQDIESSSNVQSIVGLEESICAVTTFPKLARPTESLSDSYEKLNRQKAMCRLREKRMRMKVNCNTGRKIRYYCRKQLAERRCRFKGRFIKNTAS

>XP_005703374.1 transcription factor [Galdieria sulphuraria]

MVKAREILEAQLCDSCQLERATAYCEMDCAFLCDHCDAVFHDNDTQARSHVRICTTAGVLEAYSGPRKQQKRNIQHTSFGERCFPFESFSLEPSTCGEENLSDLYLLDSTLFSKNVRSGRNQANSLEALEKLSPTDSLVPEYSSGVDDVCRLFLRDEESSPQTSSEVSNCHNDALEFRKKSHSRVTLPSSVDSMCSEEQTLISDISDSTRDIESPEPSFILDDRKRREEEEEEEAEACTEAKTERRRIALERFRQKRSNRCYQKKIRYECRKRLADVRPRIRGRFVKKEEFQALCLETGDATVPNVY

>ABO98031.1 predicted protein [Ostreococcus lucimarinus CCE9901]

MPACAACRARDADIFCLADEAFLCATCDARVHGANAVAARHERITVDEWYKRTLEAGLSEAKECGDWKAAASATASARREDEDGRGRGTSESLREKSFSLFKRDDAKTTTHDSTSSMDATISAWDVGVFLNLGENGEEDTSPRAPRMSSNSDTMIFDLDDDPLASLLEMPETESALLFDGDAASISAALEAVADQIQAVSPNQAAAKAYVSKSARDNFTPRPSPLGLGLPAAQRGPAETVGSYPPGAFPPIMMPMSSDLFGIPRRVVSKERQAQLDRYRAKRERRLMGLKKVVRYECRKTLADARVRVKGRFVKANPDEKTSALKSFQSCPDLSALVEDEDNAKPLSFAPMKHTTLDDQQLHQQNSKRRISDDRLSNSDASHDDKLDVQSMRYEILRDSGAPALHPPTIPETLPLPSGLRRTKQMRHCQSEINLMDLAGY

>ABO95850.1 predicted protein [Ostreococcus lucimarinus CCE9901]

MAPSDARASASDRARAEDATRDATGAACESCPEGARRAASWYCAQDEAYLCDACDARVHAANAIASKHERTALGTNGRGVGAHGAEDADSRRMSDAYGDGDDVEVTTDDVIGICDEYLSNPMMPSSSFPVETLDGAFWDENLGELDADGIDPESFLRDPLDDEDAAKDGVVNREIDGERSGSKYSDSVGMSKSEIEALRRVGEYASSSGFLGPILDDSAVRFLELNPSYGAFGSPSPESSDMGFESLAGKLSAVAVKREPESDLDNQTVASSGAGDANGHSAMADALRSIPEGPPSGSDTYAGLPQPQTRLERLKRWKEKRKNRNFNKVIRYQSRKACADNRPRVKGKFVKVSSVPDLSKIREAAGQSDDDDAREAEEERDKIAELGLDKGLRAPPSMRKMKKGLVSSASMPDFSMYNTMDD

>KAK9917630.1 hypothetical protein WJX75_006628 [Coccomyxa subellipsoidea]

MVQCDVCENAAGSIYCFADAAVMCQACDRTVHGANKLAAKHDRVDLSKAAESAQCDICQDRPAVLFCSEDRALICRRCDIMIHTANEFTAQHHRYLLSGTTLGLNSLGGDNSEAADKRSSDSKASSASALTREAMGVSTRSSAGAGPSGLPGNGSDLDRRMSSRGTPRSMSSGALVEVPATASPMGDTWISGRGIDVNSKAAMTPSQEQEQLRSEQVMAAQQGQAGSIGLMPSFHSGGLSDFLGAVPSSSSGGGSDNNYTLAHELLGLPTMSQAFSAKDIDAAYLFQDMGDLDDDLSSLLVPDLDFSSIASIPAPSVPKLPQTGGFADRYGVSTSPTSAGDHLVPDGVVPDILAPPLKRQRM

>KAK9917850.1 hypothetical protein WJX75_008912 [Coccomyxa subellipsoidea]

MKCQACQTAHAQVYCQESQAALCKGCSYVMGDITRFRLCALCECHPAKVFCHNDNAALCETCDADIHLSNPLAVRHDRVPLGPLACELTKDLFGSANESFAASDTDSCLQRQTPFANTVGQTDADASVLFVSNGKASLKEYDVFDLDNAFFGSDLLDFNDNFCQAPSSPSDGVVPTMDIMDSSASSNASSQDYSAVAESPNAISMMGSYSSQEQVSDSLSFVPAMPAIPMATQAMGSDSLSPSASSFDMPTLHLGSTVALDREARVMRYREKRKRRTFEKTIRYQSRKAYAEVRPRIKGRFATKEEVAAMKVAAAANGTSLAALCA

>XP_003056761.1 uncharacterized protein MICPUCDRAFT_56149 [Micromonas pusilla CCMP1545]

MTNKMPCDNCHAAPAEWFCAHDGANLCARCDVAIHTANKLAMRHERIPMEQKLANDALCQSINPTNDGFDTEMGVARNEQASTSSGAPTNETSNDLDWLVSGPHGNMDDPLSNLLDMHQDAHDSSNGDASHFGLDDASLWGSHGGFGNGNHLSSAAFRNSDARRELSAQPVVGDSAGFVTGNGAAMMGQGHADGMQRQQNDGKEISSTSGTSETTQAPTQMMPPHAGGQVNFSHGGGAPGGPGSPHYDQVLTRPFATSALRHAQLQRYRAKRLARHLGHKKIRYECRKTLADNRPRIKGRFAKVHSDPNLAAAALAAVQSCPDLTALNVVGEDKESSEDSNSKKGGRNAKSRLSNGSGSNESSSPKLKNPAHKASSKKPNTRGSGASSPEQWDSTAGVSLRSGGKGMPYTQSEVSLVDLGRRLC

>XP_003058534.1 uncharacterized protein MICPUCDRAFT_58157 [Micromonas pusilla CCMP1545]

MAKINRKIQQRVLVQGRDFPRTRTSHSAPPHATQSPTMKSVCEVCTTAPATLMCVADDAVMCGMCDKRCVNPVYTICHEDRAFLCRGCDVSLHSANEAVKKHRRFLYTGVTVALAPLGEKETATPREATDIVKEVPPMAAPIRPTVTQPQPQKKRKAVDIEDDFAVPTMSPDNSAGHGGVQWQSDEEFNSFVQDFVGKGEKNPDVNFDGFLDNFFDDVPLSDDFGVVPTM

>KAA8498441.1 hypothetical protein FVE85_6026 [Porphyridium purpureum]

MSKIFVSLMICEFCDWRPAAVYCESESVTMCRKCDKFFHSGHEACLHLRRPISVTQMQRSVTLLGYESPSPSSSSSADEKDYVLESDDGSVVEPPSSPPSSLVNESMDGVVERAIDVIQAPNHDLPDSARAFDAQKRAMQVMAERAAQKHRQHNAENKRHVTKKRRTDHHMRNVTATSPNIVIDEASGVVRQTRTGGAASNAGGSGGDGSEPTASSLPQRDPASSGNDTRTSGQSNSGNNAASGGSGTGGSGTGRGFGTGGSGTGGSGTGGSLATDGLETVKGSASKPPRHSLGQNHSRSGSGSGSGSGKDGGSDSGGSGSGSGSRSGRAGPAHSSGSGNSRTGSGTVSGSGSRTGSGSGSGSGSGSGRASGNGSGSASASASASASNGMSSQSRRVESLSSGAVSAKMMPPSDQAPQ

>KAA8494015.1 hypothetical protein FVE85_3990 [Porphyridium purpureum]

MMRNSTNIVQFRVVCCECGAAEARRWCVQEMAPLCERCDVARHKGAADGASGLRSQRQPHERVAIAHDSAMAVRCERCLEMPCSFYSRSQHACWCETCAAEMLQQQQQQQQQHLAEDAAVKFPHNPLQEQDAQHESGGTTSKDTINPVATVKKDLLAPKAHVQLAVFGQHTTENVVFEKMDFSAGLTGNKGTAPDPWMIAKDHRRRKAATPSARVLSSLGLPEPVHVKVARWLDSEQNLTNVAVKDTGGEQGSDSRADLGGNTPENIQEHRTGDQRKPQKPSQLEESAPASMSTDSTPLELNTRPVLSSHERSSHIDLTYFSQRVPQFVVQDTGNHFYQAQDET
